# HORmon: automated annotation of human centromeres

**DOI:** 10.1101/2021.10.12.464028

**Authors:** Olga Kunyavskaya, Tatiana Dvorkina, Andrey V. Bzikadze, Ivan A. Alexandrov, Pavel A. Pevzner

## Abstract

Recent advances in long-read sequencing opened a possibility to address the long-standing questions about the architecture and evolution of human centromeres. They also emphasized the need for centromere annotation (partitioning human centromeres into monomers and higher-order repeats (HORs)). Even though there was a half-century-long series of semi-manual studies of centromere architecture, a rigorous centromere annotation algorithm is still lacking. Moreover, an automated centromere annotation is a prerequisite for studies of genetic diseases associated with centromeres, and evolutionary studies of centromeres across multiple species. Although the monomer decomposition (transforming a centromere into a monocentromere written in the monomer alphabet) and the HOR decomposition (representing a monocentromere in the alphabet of HORs) are currently viewed as two separate problems, we demonstrate that they should be integrated into a single framework in such a way that HOR (monomer) inference affects monomer (HOR) inference. We thus developed the HORmon algorithm that integrates the monomer/HOR inference and automatically generates the human monomers/HORs that are largely consistent with the previous semi-manual inference.

## Introduction

Recent advances in long-read sequencing technologies led to rapid progress in centromere assembly in the last year (Bzikadze and Pevzner, 2020, Miga et al., 2020, Nurk et al., 2020, Nurk et al., 2021, Logsdon et al. 2021, Altemose et al., 2021) and, for the first time, opened a possibility to address the long-standing questions about the architecture and evolution of human centromeres (Rice, 2019, Thakur et al., 2021). *Alpha satellite arrays* of “live” human centromeres that organize the kinetochore (which we refer to simply as *centromeres*) are tandem DNA repeats that are formed by units repeating thousands of times with limited nucleotide-level variations but extensive variations in copy numbers in the human population (Black and Giunta, 2018). Each such unit represents a tandem repeat formed by smaller repetitive building blocks (referred to as *monomer-blocks*), thus forming a *stacked tandem repeat* (Figure cenXArchitecture). Partitioning all monomer-blocks into clusters of similar monomer-blocks defines *monomers*, where each monomer represents the consensus of all monomer-blocks in a given cluster. The emergence of centromere-specific stacked tandem repeats is a fascinating and still poorly understood evolutionary puzzle. Its studies started nearly half a century ago when the future Nobel laureate George Smith explained how the unequal crossovers could lead to the emergence of stacked repeats (Smith, 1976), and continue today (Malik and Henikoff, 2009, Rice 2019, Uralsky et al., 2019).

**Figure cenXArchitecture.**
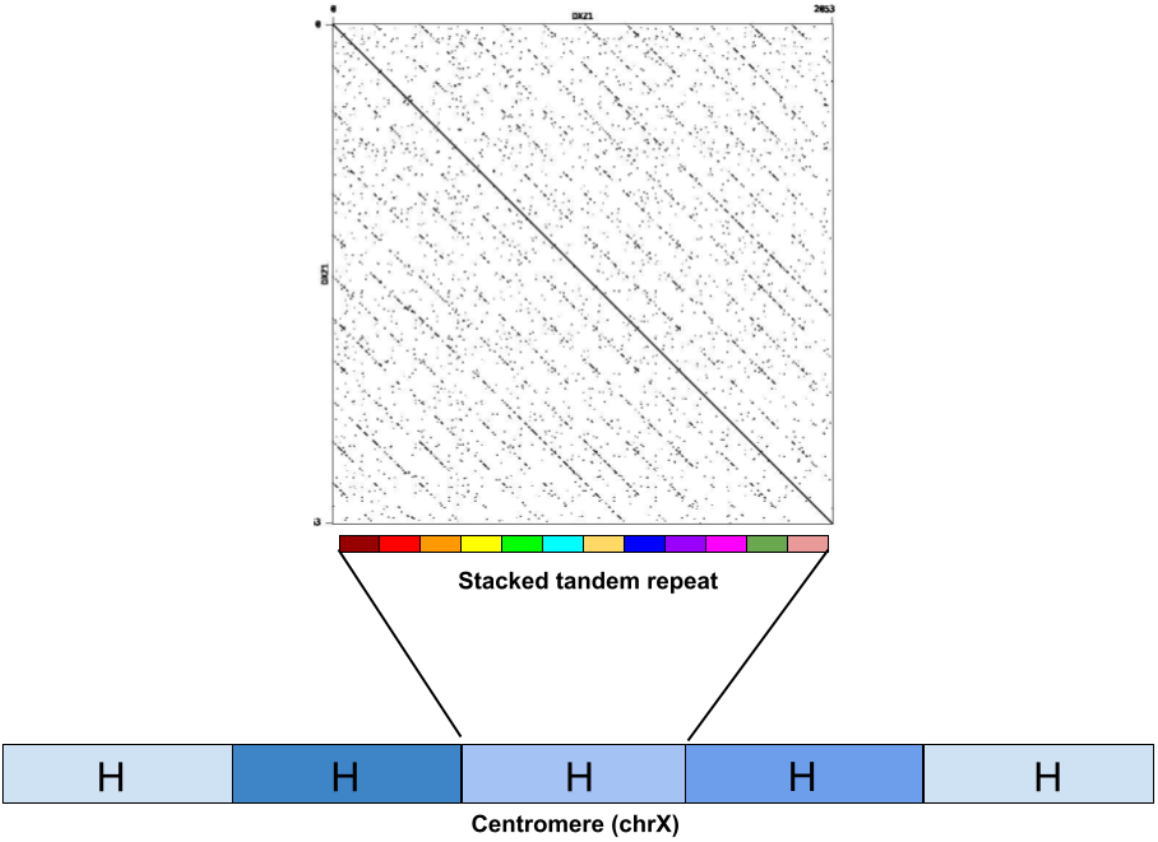
The architecture of centromere on chromosome X. The centromere of chromosome X (cenX) consists of ~18100 monomers of length ≅171 bp each based on the cenX assembly in Bzikadze and Pevzner, 2020 (the latest T2T assembly (Nurk et al., 2021) represents a minor change to this assembly). These monomers are organized into ~1500 units. Five units in the Figure are colored by five shades of blue illustrating unit variations. Each unit is a stacked tandem repeat formed by various monomers. The vast majority of units in cenX correspond to the canonical HOR which is formed by twelve monomers (shown by twelve different colors). The figure on top represents the dot plot of the nucleotide sequence for the canonical HOR that reveals twelve monomers. While the canonical units are 95–100% similar, monomers are only 65–88% similar. In addition to the canonical 12-monomer units, cenX has a small number of partial and auxiliary HORs with varying numbers of monomers.

Each human monomer is of length ≅171 bp and each higher-order unit is formed by multiple monomers that differ from each other. A monomer is *frequent* if the number of monomer-blocks in its cluster exceeds a frequency threshold, and *infrequent*, otherwise. Recently, Uralsky et al., 2019, Bzikadze and Pevzner, 2020, and Dvorkina et al., 2020, 2021 revealed still underexplored *hybrid* monomers (each hybrid monomer is a concatenate of two or even more frequent monomers) and hypothesized that such hybridization may drive the “birth” of new frequent monomers. Different human centromeres typically have different monomers and units while the number of the frequent monomers in a unit varies from 2 for chromosome 19 to 19 for chromosome 4.

A *canonical* (*cyclic*) *order of monomers* (referred to as a *higher-order repeat* or *HOR*) is specific to each centromere and is defined evolutionarily as the ancestral and chromosome-specific order of frequent non-hybrid monomers that has evolved into the complex organization of extant centromeres. This definition, however, is computationally non-constructive since the ancestral order is unknown and since no algorithm for its inference has been described yet. The current view of centromere evolution can be summarized by the following framework that we refer to as the *Centromere Evolution (CE) Postulate*:

- Each extant human centromere has evolved from a *single* ancestral HOR *M*_1_, …, *M*_*k*_ formed by *k different* monomers. Hence, each monomer occurs in a HOR only once. The parameter *k* (number of monomers in a HOR) varies between various centromeres.
- Each frequent non-hybrid monomer in a centromere has evolved from a single ancestral monomer. The number of ancestral monomers equals the number of frequent non-hybrid monomers in the extant centromere.
- Each hybrid monomer has evolved from a concatenate of two (or even more) ancestral monomers and does not participate in the ancestral HOR.
- In addition to units formed by canonical HORs, there exist units formed by *partial HORs* (substrings of canonical HORs). All other units are referred to as *auxiliary HORs*. Although the canonical HOR corresponds to the most frequent unit for most human centromeres, it is not always the case.

Although the CE postulate is universally accepted (Willard and Waye 1987, Alexandrov et al. 2001, McNulty and Sullivan 2018), we are not aware of a rigorous proof of this postulate or an algorithm that, given an extant centromere, derives its canonical HOR. Moreover, since the concept of a HOR is parameter-dependent, the CE postulate may hold for some parameters and fail for others. However, it is not clear how to select various parameters such as the frequency threshold parameter (for defining the concept of a frequent monomer), the percent identity parameter (for deciding which monomer-blocks correspond to the same monomer), and parameters for classifying a monomer as a hybrid (Dvorkina et al., 2021).

Moreover, the CE postulate implicitly assigns the inferred HOR to a particular (and unspecified) moment in the past. For example, while the HOR for centromere X (referred to as cenX) consists of 12 monomers, this 12-monomer HOR evolved from an even more ancient 5-monomer ancestral HOR (Waye and Willard 1987, reviewed in Alexandrov et al., 2001). It is thus not clear how an algorithm for HOR inference should choose between a 12-monomer HOR and a 5-monomer HOR for cenX. Further, even if the CE Postulate holds, it may be impossible to infer canonical HORs if nearly all information about the ancestral HOR was erased by millions of years of evolution, e.g., it is unclear how to derive HORs in mouse centromeres (Thakur et al., 2021).

Recent evolutionary studies of centromeres (Uralsky et al., 2019, Bzikadze and Pevzner, 2020, Suzuki et al., 2020) revealed the importance of partitioning them into monomers, the problem that was addressed by the StringDecomposer algorithm (Dvorkina et al., 2020). Given a nucleotide string *Centromere* and a monomer-set *Monomers*, StringDecomposer decomposes the *Centromere* into monomer-blocks (each block is similar to one of the monomers) and transforms *Centromere* into a *monocentromere* string *Centromere\** over the alphabet of monomers. For each monomer *M*, it generates the set of *M-blocks* in the centromere that are more similar to *M* than to other monomers (ties broken arbitrarily).

StringDecomposer opened a possibility to automatically generate all HORs and annotate human centromeres (i.e., partition them into canonical, partial, and auxiliary HORs), the problem that remains unsolved despite multiple studies in the last four decades (Waye and Willard, 1985, Alexandrov et al., 2001, Paar et al., 2005, Alkan et al., 2007, Shepelev et al., 2015, Sevim et al., 2016, McNulty and Sullivan, 2018, Uralsky et al., 2019). However, the challenge of properly defining the set of all human monomers remained outside the scope of the StringDecomposer tool. Although Sevim et al., 2016 presented a large set of human monomers, it is unclear if this set is compatible with the CE Postulate. As a result, it remains unclear how to computationally define the complete set of monomers (a prerequisite for launching StringDecomposer) and HORs in human centromeres.

Previous semi-manual studies inferred many HORs and greatly contributed to our understanding of the architecture of human centromeres (reviewed in Alexandrov 2001, McNulty and Sullivan, 2018). However, they did not specify an *algorithmically constructive definition* of a HOR. Instead, an order of monomers in a consensus HOR was implicitly defined as the “ancestral order” without specifying how to derive this order and how to prove that it is correct and unique. Although Paar et al., 2005, Alkan et al., 2007, and Sevim et al., 2016 described various HOR inference heuristics (ColorHOR, HORdetect, and Alpha-CENTAURI, respectively), these studies have not specified the exact objective function for HOR inference. As such, the concept of a HOR is highly dependent on the parameters used for generating the monomer-set. Moreover, the nucleotide sequences for human HORs of “live’’ human centromeres have been manually extracted at the dawn of the sequencing era and used reads (often sampled from a single clone from a specific centromere) rather than completely assembled centromeres, raising questions about their accuracy (Bzikadze and Pevzner, 2020; Miga and Alexandrov 2021). For example, HOR DXZ1 (S3CXH1L) on cenX, the first inferred human HOR, was derived based on limited sequencing data from a single clone (Waye and Williard, 1985). The sequence of this HOR differs from the HOR extracted from the complete cenX assembly, suggesting that either (i) reads used for deriving DXZ1 were limited to a small region of cenX that does not adequately represent the entire centromere, or (ii) HORs extracted from different individuals may be different.

These limitations prevent future evolutionary studies of centromeres across multiple species. Addressing them is important since long and accurate PacBio HiFi reads have already been used for centromere assembly in fish (Xue et al., 2021), and since various HiFi assembly projects are currently underway, opening a possibility to assemble vertebrate centromeres in the near future. On the other hand, the Telomere-to-Telomere (T2T) Consortium and the Human Pangenome Reference Consortium are now assembling centromeres from multiple humans. Since their manual annotation (including monomer and HOR inference) is prohibitively time-consuming, automated annotation is a prerequisite for any centromere analysis in the future.

Dvorkina et al., 2021 developed the CentromereArchitect tool that addressed both the monomer inference and HOR inference as two separate problems. In particular, the HOR inference was addressed as a data compression problem rather than an evolutionary problem that takes into account the CE Postulate. Thus, although CentromereArchitect enabled an automated inference of monomers, it remains unclear whether its HORs inference adequately reflects the centromere evolution. Our analysis revealed that, to generate a biologically adequate centromere annotation, the monomer and HOR inference should be viewed as two interconnected problems in such a way that HOR (monomer) generation affects monomer (HOR) generation.

In the past, the monomer generation problem was addressed as clustering of monomer-blocks without considering the follow-up inference of HORs derived from the resulting monomers (Dvorkina et al., 2021). Since this is a complex clustering problem, any clustering algorithm may merge some biologically distinct monomer-blocks into a single cluster and split a single cluster into multiple ones. Another complication is the inference of hybrid monomers that, by definition, do not participate in canonical HORs.

Below we describe the HORmon algorithm that addresses these complications by incorporating the monomer and HOR generation into a single pipeline (Figure HORmonPipeline). HORmon generated the first automated centromere annotation that is largely consistent with the CE Postulate and previous manual centromere annotations. Recognizing that HORs represent an important evolutionary concept, we show how HORmon can be used to automatically derive the currently known HORs.

**Figure HORmonPipelin.**
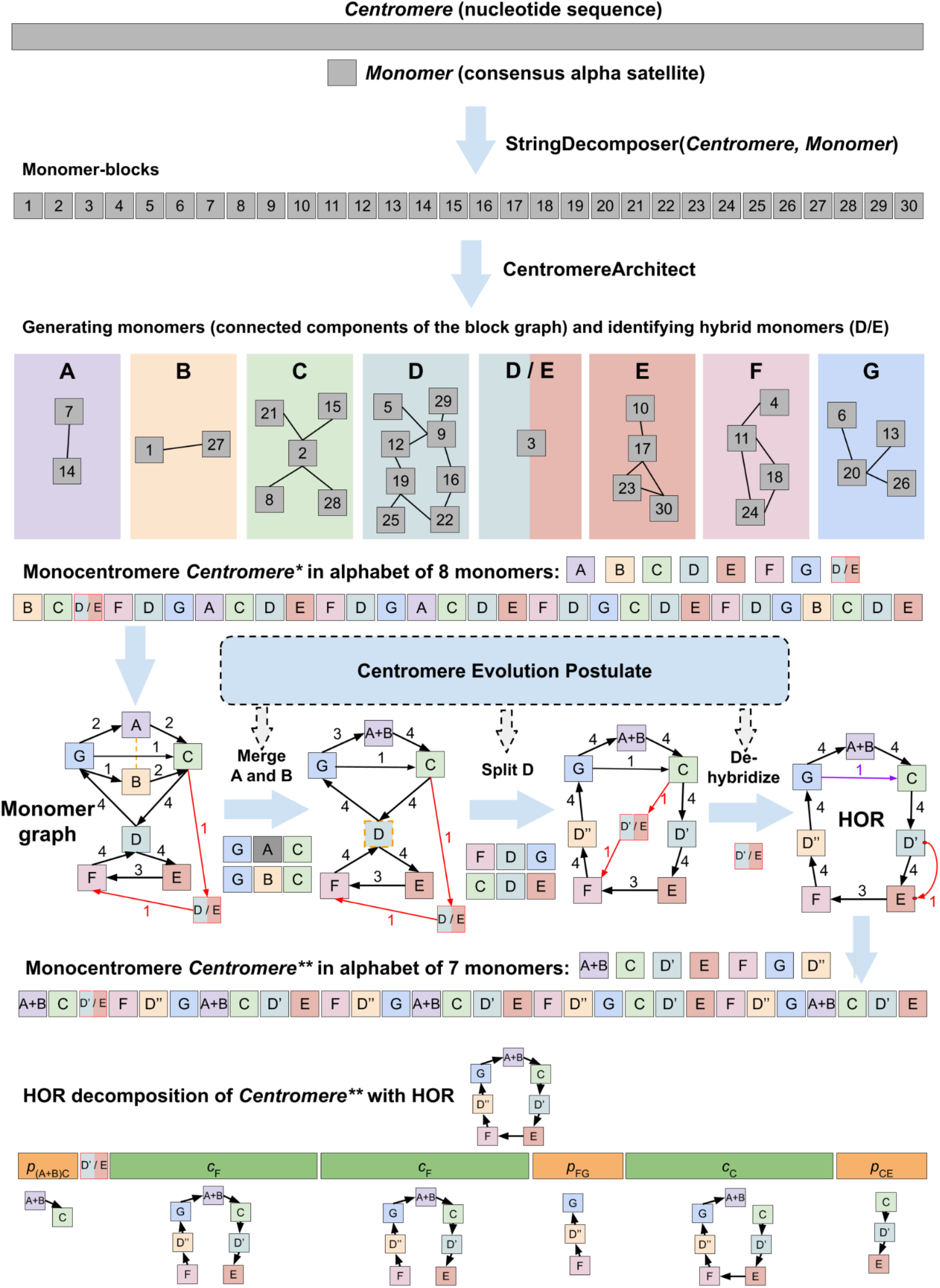
HORmon pipeline. Given the nucleotide sequence *Centromere* and a consensus alpha satellite sequence *Monomer*, HORmon iteratively launches StringDecomposer (Dvorkina et al., 2020) to partition *Centromere* into monomer-blocks. After each launch of StringDecomposer, HORmon launches CentromereArchitect (Dvorkina et al., 2021) to cluster similar monomer-blocks into monomers, identify hybrid monomers (represented by a single hybrid D/E of monomers D and E), and transform *Centromere* into the monocentromere *Centromere\**. Afterward, HORmon uses the generated monocentromere to construct a monomer-graph (red edges connect the hybrid monomer D/E with the rest of the monomer-graph). To comply with the Centromere Evolution Postulate, HORmon performs split/merge transformations and dehybridizations on the initial monomer-set. The orange dotted undirected edge connects similar monomers A and B to indicate that they represent candidates for merging. The breakable monomer D is shown as a dotted vertex to indicate that it is a candidate for splitting. The dehybridization substitutes the hybrid vertex D’/E by a single “red” edge that connects the prefix of D’ with the suffix of E. Split, merge, and dehybridization operations result in a new monomer-set and transform *Centromere* into the monocentromere *Centromere\*\**. The black cycle in the monomer-graph of *Centromere\** represents the HOR; the purple edge connecting monomers G and C is a low-frequency chord in this cycle. HORmon uses this HOR to generate the HOR decomposition of *Centromere\*\** into the canonical (*c*_F_, *c*_C_), partial (*p*_(A+B)C_, *p*_FG_, *p*_CE_), and auxiliary (the single block D’/E) HORs. *c*_F_ and *c*_C_ refer to traversing the (canonical) HOR starting from monomers F and C, respectively. *p*_(A+B)C_, *p*_FG_, and *p*_CE_ refer to partial traversals of the HOR from monomer A+B to C, from F to G, and from C to E, respectively.

## Results

### A brief description of the HORmon algorithm

Figure HORmonPipeline illustrates the various steps of the HORmon algorithm for monomer and HOR inference. Supplementary Note “HORmon terminology” summarizes the main notation that we use throughout the paper.

### Datasets

We extracted the alpha satellite arrays from the assembly (public release v1.0) of the effectively haploid CHM13 human cell line constructed by the T2T Consortium (Miga et al., 2020; Logsdon et al., 2020, Nurk et al., 2021, Altemose et al., 2021). We also extracted the alpha satellite array of the newly assembled centromere of chromosome X from HG002 cell line sequenced by the Human Pangenome Reference Consortium. For simplicity, we refer to these two genomes as the CHM13 and HG002 genomes. Supplementary Note “Information about datasets” provides information about the accession numbers and the coordinates of the extracted regions for all “live” human centromere arrays.

### Monomer inference

HORmon launches CentromereArchitect (Dvorkina et al., 2021) to generate the initial monomer-set and further modifies it by using the monomer-HOR feedback loop described in the Methods section (Figure HORmonPipeline). Supplementary Note “HORmon monomer naming” describes how HORmon assigns names to monomers and provides correspondence between these names and the traditional names described in Uralsky et al., 2019.

Since CentromereArchitect identifies many infrequent monomers, comparing its monomer-set with the previously identified monomer-sets, e.g., the monomer-set *MonomersT2T* (Altemose et al., 2021) used by the T2T consortium (based on the monomer-set derived in Shepelev et al. 2015 and Uralsky et al., 2019), is not straightforward. HORmon thus filters the monomer-set generated by CentromereArchitect as described below.

We refer to the set of frequent monomers obtained from CentromereArchitect as *MonomersNew*. Supplementary Note “Evaluating the monomer-sets’’ describes the procedure for construction of *MonomersNew* and illustrates that it provides a minor improvement over the (manually constructed) *MonomersT2T* monomer-set with respect to standard clustering metrics. However, as with any clustering approach, the parameter-dependent CentromereArchitect may both split and aggregate monomers as compared to the biologically adequate clustering. Moreover, the monomer-set *MonomersT2T* does not attempt to solve the Monomer Inference Problem that CentromereArchitect addresses (Dvorkina et al., 2021). Instead, it generates clustering that is consistent with CE Postulate which can be suboptimal with respect to standard clustering metrics that do not take into account any evolutionary assumptions.

### The challenge of monomer generation

Although it is unclear what is a biologically adequate clustering of monomer-blocks, positional information about these blocks (i.e., pairs, triples, etc., of consecutive monomers in the monocentromere) often reveals monomers that were erroneously split/aggregated. This positional information helps one to generate a more adequate monomer-set with respect to the CE Postulate, not unlike the positional information about orthologs in comparative genomics studies (Jun et al., 2009). Two monomers are called *similar* if the percent identity between them exceeds a threshold *minPI* (default value 94%). In the subsection “Positionally-similar monomers” (Figure HORmonPipeline), we define the concept of positional similarity and classify two similar monomers as *positionally-similar* if their positional similarity exceeds a threshold *minPosSim* (default value 0.4).

To illustrate the challenge of generating a biologically adequate clustering, we consider similar frequent monomers *M’* and *M****”*** from the monomer-set *Monomers* that would be merged into a single monomer if the clustering parameters were slightly relaxed. Since it is unclear how to select clustering parameters, it is also unclear whether such merging would represent a biologically adequate clustering as opposed to the clustering that separates these monomers. However, one may argue that if *M’* and *M****”*** are always flanked by the same frequent monomers *X* and *Y* in a monocentromere (resulting in triples *XM’Y* and *XM****”*** *Y*), these two monomers are likely erroneously split and should be merged into a single monomer *M*, defined as the consensus of all *M’*-blocks and *M****”*** -blocks. Such merging is justified from the perspective of the CE Postulate since each non-hybrid monomer occurs exactly once in a HOR. Specifically, unless monomers *M’* and *M****”*** are merged, the HOR cannot traverse monomers *X* and *Y* exactly once as required by the CE Postulate.

On the other hand, a frequent monomer *M* that is flanked either by frequent *X’* and *Y*’ (resulting in a triple *X’MY’*) or by frequent *X****”*** and *Y****”*** (resulting in a triple *X****”*** *MY****”***) conflicts with the CE Postulate. Since this monomer is likely erroneously aggregated from two different monomers, it can be split into monomers *M’* and *M****”***, resulting in triples *X’M’Y’* and *X****”*** *M****”*** *Y****”***, respectively. The monomers *M’* (*M****’***) can be defined as the consensus of all *M’*-blocks (*M****”*** -blocks) in triples *X’M’Y’* (*X****”*** *M****”*** *Y****”***).

Even though such transformations are not necessarily justified with respect to optimizing the standard clustering metrics, Supplementary Note “Evaluating the monomer-sets” illustrates that the monomer-set transformed by merging/splitting operations in HORmon is largely comparable to the monomer-set generated by CentromereArchitect with respect to various clustering metrics.

In addition to generating the monomer-set, CentromereArchitect includes a HOR inference algorithm based on iteratively defining the units as the “heaviest” substrings of a monocentromere (Dvorkina et al., 2021). Although this definition is adequate from the perspective of data compression, it does not necessarily reflect the evolutionary history of a centromere (even though many resulting units correspond to canonical, partial, and auxiliary HORs). Moreover, Dvorkina et al., 2021 derived monomers independently from HORs without accounting for hybrid monomers, positional information, and the CE Postulate. Below we demonstrate that positional information, as well as information about hybrid monomers, is important for both monomer and HOR inference. The Methods section describes how to identify erroneously aggregated/split monomers and split/merge them.

We further introduce the concept of a breakable monomer, i.e., a monomer that is amenable to splitting into two or more monomers in such a way that the enlarged monomer-set still adequately represents the centromere architecture. In contrast, splitting an unbreakable monomer results in an inadequate representation of the centromere architecture. We show that a series of split and merge operations results in unbreakable monomers for cen1, cen13, and cen18 that prevent HORmon from reporting HORs in these centromeres. We further describe a special procedure for splitting unbreakable monomers in these problematic centromeres (subsection “Splitting unbreakable junction monomers reveals HORs in cen1, cen13, and cen18”).

Split and merge operations on the monomer-set *MonomersNew* result in a monomer-set *MonomersNew*^*+*^, while further hybridization of hybrid monomers (Figure HORmonPipeline) and splitting of unbreakable monomers result in the monomer-set *MonomersFinal* described in Supplementary Table ModifiedMonomerSet.

### Monomer-graph

Given a monocentromere, its directed *monomer-graph* is constructed on the vertex-set of all its monomers and the edge-set formed by all pairs of its consecutive monomers. The *multiplicity* of an edge (*M,M’*) in the monomer-graph is defined as the number of times the monomer *M’* follows the monomer *M* in the monocentromere (Figure HORmonPipeline). We note that the monomer-graph of a monocentromere *Centromere\** is the *de Bruijn graph DB*(*Centromere\**, 2) (Compeau et al., 2011). Figure MonomerGraphX presents the monomer-graph for cenX in the CHM13 genome (top) and the HG002 genome (bottom) built using the monomer-set extracted by CentromereArchitect from CHM13 genome (Dvorkina et al., 2021). Both graphs reveal the cycle formed by twelve high-multiplicity edges that form the canonical twelve-monomer HOR in cenX. In addition, the monomer-graph for CHM13 reveals two infrequent hybrid monomers and ten low-multiplicity edges. In contrast, the monomer-graph for HG002 reveals only one infrequent hybrid monomer and only five low-multiplicity edges. These differences suggest that hybrid monomers represent a rather recent evolutionary innovation and illustrate large variations in centromeres across the human population.

**Figure MonomerGraphX.**
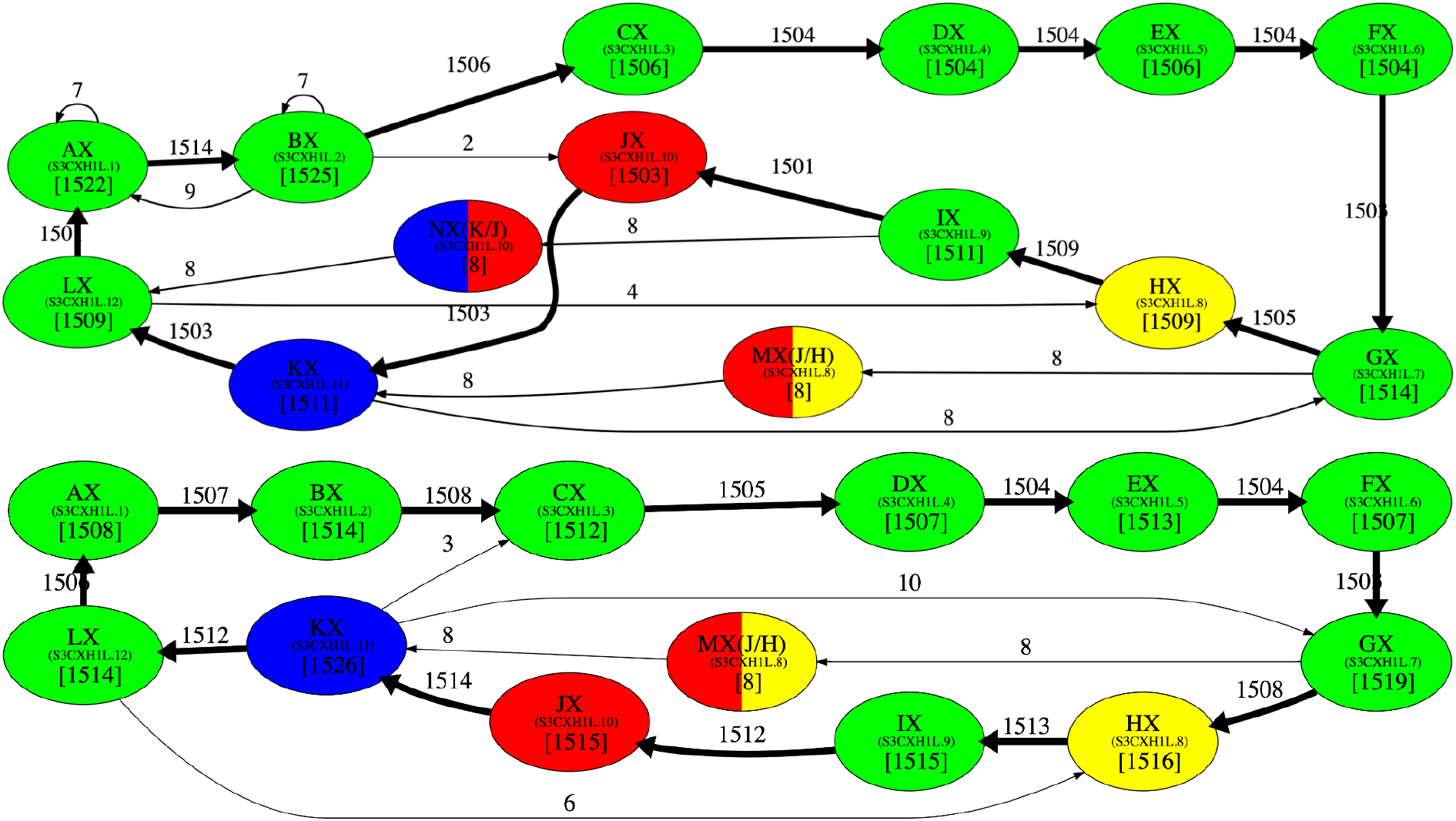
The monomer-graph of cenX in the CHM13 (top) and HG002 (bottom) genomes. The monomer-graphs of cenX were constructed on the monocentromere that was generated from the monomer-sets consisting of two infrequent hybrid monomers (labeled as MX and NX) and twelve frequent canonical monomers (labeled as AX, BX, CX, …, KX, and LX) that contribute to the canonical DXZ1 HOR in cenX (Dvorkina et al., 2021). Small font corresponds to the naming conventions introduced in Shepelev et al., 2015. The hybrid monomers M and N are inferred in Dvorkina et al., 2020. A hybrid monomer formed by frequent monomers X and Y is represented as a bicolored vertex (two colors correspond to the colors of X and Y) and is denoted as (X/Y). Only edges of the monomer-graph with multiplicity exceeding 1 are shown (edges with multiplicity exceeding 100 are shown in bold). The cycle formed by bold edges (with multiplicities above 1500) traverses the twelve most frequent monomers that form the canonical cenX HOR.

Figure MonomerGraphX creates a false impression that simply ignoring the low-multiplicity edges and hybrid monomers in the monomer-graph of a centromere would result in a graph with a single cycle that forms a HOR. Although after performing a series of split/merge transformations on the monomer-set generated by CentromereArchitect (Dvorkina et al., 2021), this is indeed true for centromeres 3, 11, 14, 16, 17, 19, 20, 21, 22, and X, the remaining human centromeres have a more complex evolutionary history, resulting in complex architectures that we analyze below.

### Monomer-graphs of human centromeres

Given a monomer-set *Monomers* and a monocentromere *Centromere\**, we define *minCount*(*Monomers*) = min ^all monomers *M* in *Monomers*^ *count*(*M,Centromere*\*). HORmon uses the set *MonomersNew*^***+***^ to generate the monocentromere *Centromere\*\** (split and merge operations on the monomer-set *MonomersNew* result in the monomer-set *MonomersNew*^*+*^), generates the monomer-graph as the de Bruijn graph *DB*(*Centromere\*\**, 2), and removes edges that have multiplicity below min(*MinEdgeMultiplicity, minCountFraction · minCount*(*MonomersNew*^***+***^)) with the default values *MinEdgeMultiplicity =* 100, *minCountFraction* = 0.9 (Figure HORmonPipeline). Supplementary Figure MonomerGraphs provides information about the generated monomer-graphs for all human centromeres.

The monomer-graphs of ten centromeres (3, 11, 14, 16, 17, 19, 20, 21, 22, and X) are formed by cycles that immediately reveal HORs. The monomer-graph for cen17 contains two cycles: the higher-multiplicity cycle corresponds to the D17Z1 HOR, while the lower-multiplicity cycle corresponds to its *sister* HOR D17Z1-B (see Miga and Alexandrov 2021 for discussion of sister HORs). The remaining monomer-graphs contain (albeit implicitly) information about HORs but represent a more detailed view of the evolutionary history of centromeres. To reveal HORs in these monomer-graphs, HORmon constructs simplified monomer-graphs described in the Methods section.

Figure SimplifiedMonomerGraphs shows that the simplified monomer-graphs represent cycles (corresponding to HORs) for all centromeres but centromeres on chromosomes 1, 5, 8, 9, 13, and 18 that do not have Hamiltonian cycles and represent special cases that we consider below.

**Figure SimplifiedMonomerGraphs.**
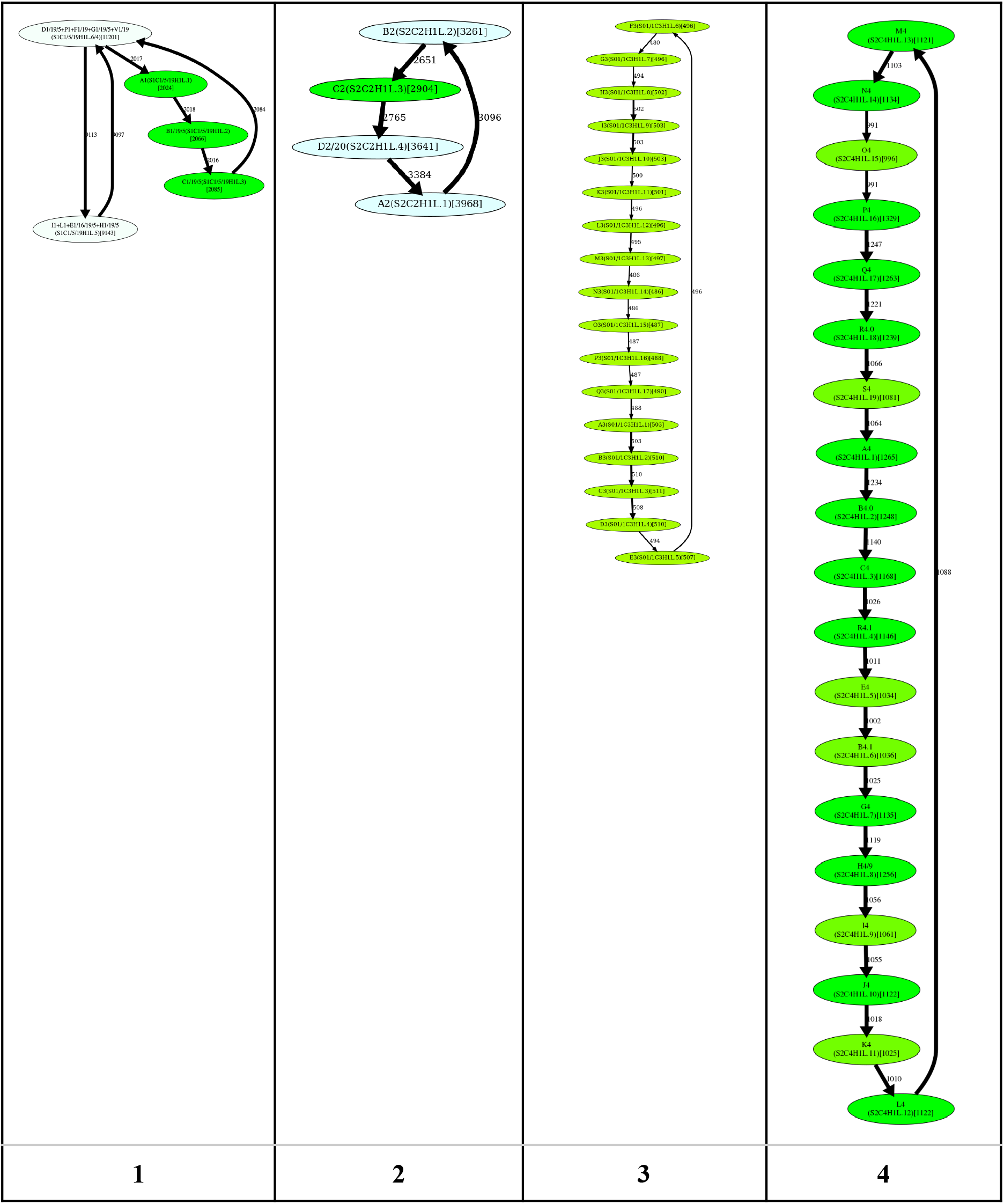
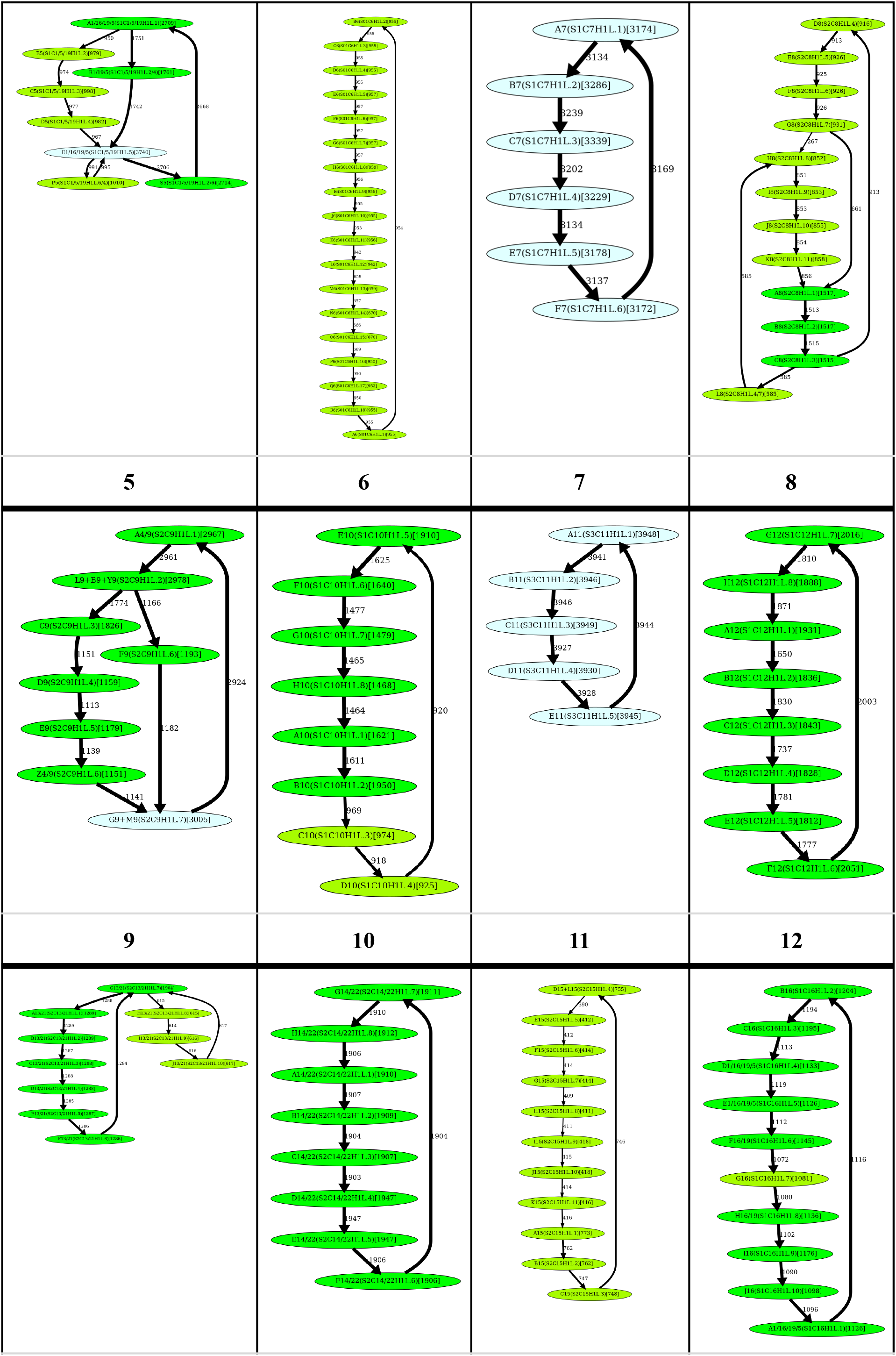
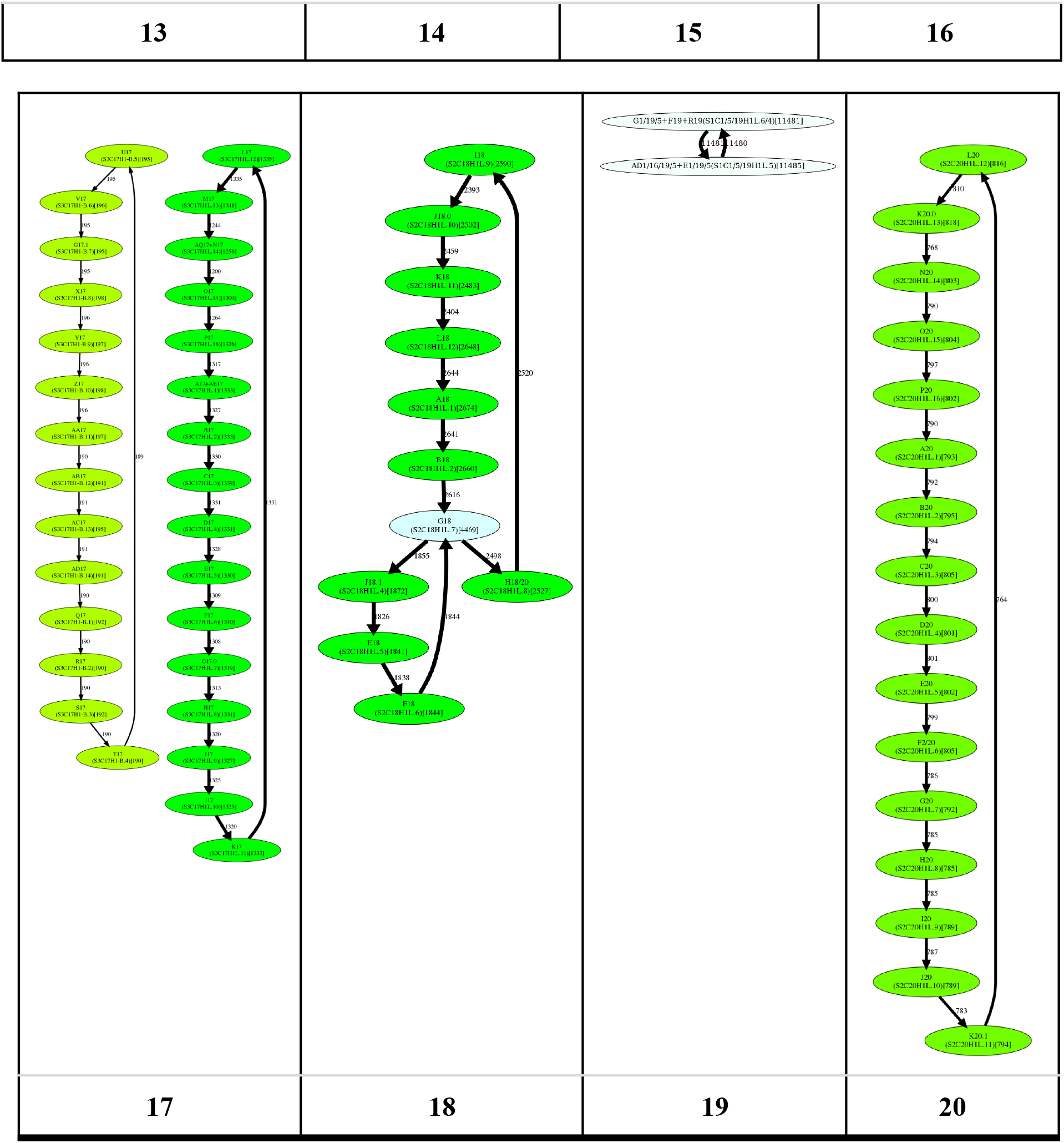
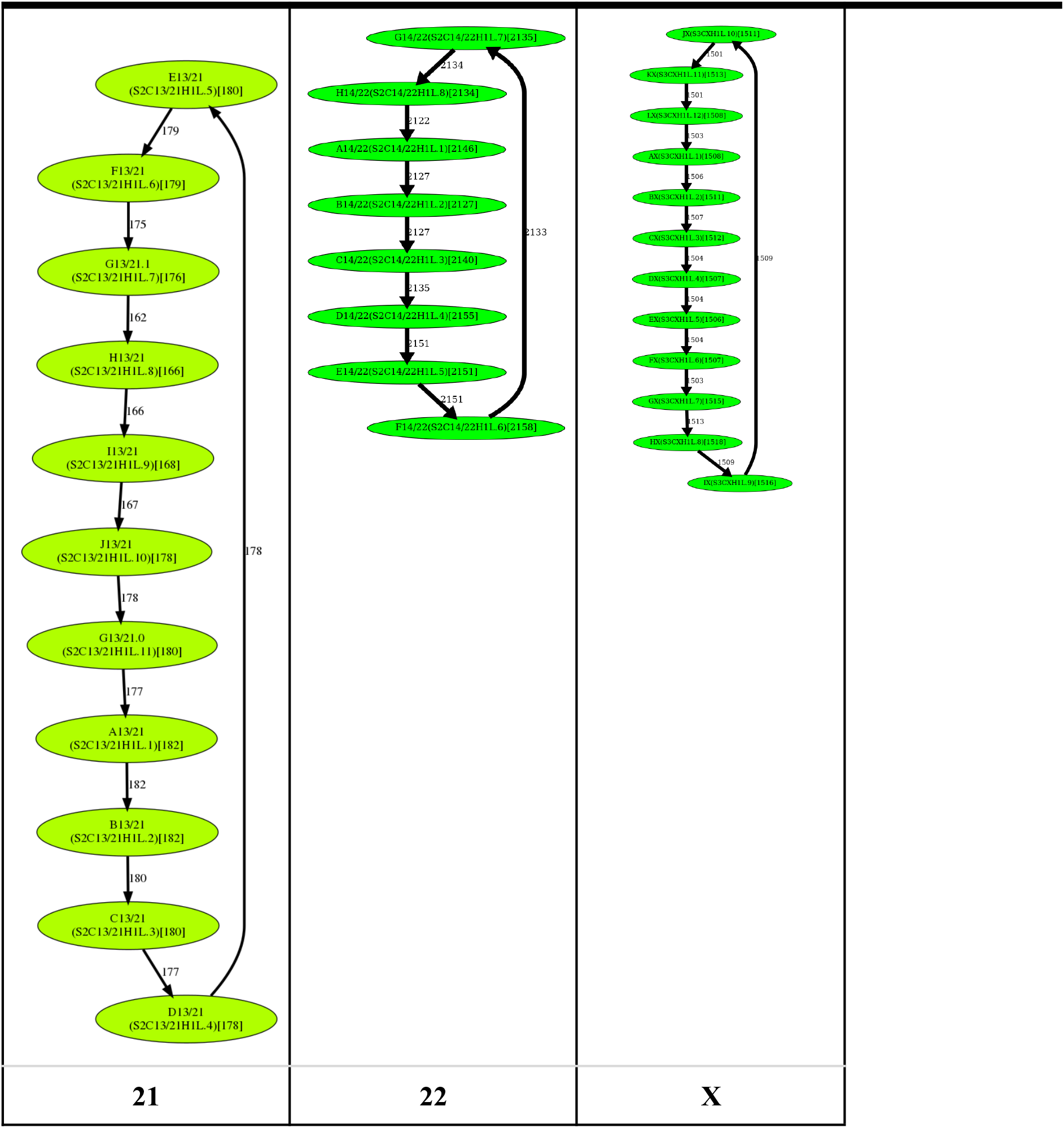
The simplified monomer-graphs of human centromeres. The label of each vertex represents the monomer ID and its count in the monocentromere (in parentheses). The label of an edge represents its multiplicity. The width of an edge (color of a vertex) reflects its multiplicity (count of a monomer). The names of the monomers follow the naming convention introduced in Shepelev et al., 2015.

### Limitations of the CE Postulate

Figure TwoMonocentromeres shows two “monocentromeres” that result in identical monomer-graphs (formed by cycles AB and BC connected via the junction vertex B), yet represent very different evolutionary scenarios. Although one can come up with a plausible “evolutionary” scenario for these centromeres, it is not clear how to find out their HORs that would be compliant with the CE Postulate. The first monocentromere can be described as two cycles (one formed by vertices A and B and another formed by vertices C and B), while the second one can be described by a single cycle (formed by vertices A, B, C, and B) in the monomer-graph (Figure TwoMonocentromeres).

**Figure TwoMonocentromeres.**
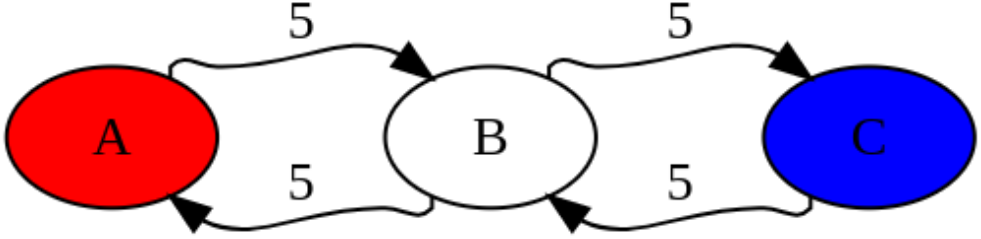
Two different “monocentromeres” BABABABABABCBCBCBCBCB and BABCBABCBABCBABCBABCB have the identical monomer-graphs.

The concept of a HOR does not allow one to adequately describe the differences between the monocentromeres shown in Figure TwoMonocentromeres as it requires that each monomer participates in a HOR once, necessitating the sequence ABC (that does not adequately reflect the centromere architecture) as the only possible HOR candidate. While this example might be considered artificial, any algorithm for centromere annotation should adequately handle such cases, even if they rarely appear in the human centromeres. As we show below, cen13 and cen18 represent an evolutionary scenario that is similar to the toy centromere described in Figure TwoMonocentromeres.

The previous approaches to centromere annotation were based on the CE Postulate and described centromeres in terms of complete and partial HORs. Given toy monocentromeres

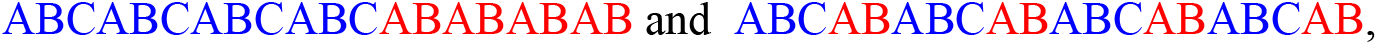

they described these very different architectures in the same way: as the complete HOR ABC and the partial HOR AB, each repeating five times. Since this representation does not distinguish these two very different centromere architectures, there is a need for a more general representation of the centromere architecture that will adequately reflect all complete and partial HORs.

### Splitting unbreakable monomers reveals HORs in cen1, cen13, and cen18

A monomer is breakable if it is amenable to splitting into two or more monomers in such a way that the enlarged monomer-set still adequately represents the centromere architecture (see Methods). In contrast, splitting an unbreakable monomer leads to conflicts and results in an inadequate representation of the centromere architecture. Even if a monomer is breakable, splitting it into two very similar monomers (e.g., monomers *M’* and *M****”*** that differ in a single position) may lead to a misclassification of monomer-blocks since all centromere decomposition tools, including StringDecomposer, often misclassify an *M’*-block as an *M****”*** -block and vice versa. Such misclassified monomer-blocks may lead to downstream challenges in analyzing centromere architecture and evolution.

Although the simplified monomer-graphs of cen1, cen13, and cen18 are formed by two cycles that share a junction vertex (that deviate from the definition of a HOR as a single cycle), these two cycles can be transformed into a single cycle by splitting the junction vertices (Figure ComplexCentromeres). However, since these junction vertices correspond to unbreakable monomers, splitting them raises concerns. Indeed, it either conflicts with some frequent traversals through the junction vertex or results in a pair of highly similar monomers that would be merged into a single monomer even under extremely stringent values of HORmon parameters.

Splitting a junction monomer in cen1 results in two monomers that differ in 11 nucleotides. This transformation results in a simplified monomer-graph that contains a cycle that corresponds to a HOR and a dimer formed by two high-multiplicity anti-parallel edges (Figure ComplexCentromeres). In fact, this dimer was originally reported as a HOR in cen1 (Carine et al., 1989, Alexandrov et al., 2001, McNulty and Sullivan 2018).

**Figure ComplexCentromeres.**
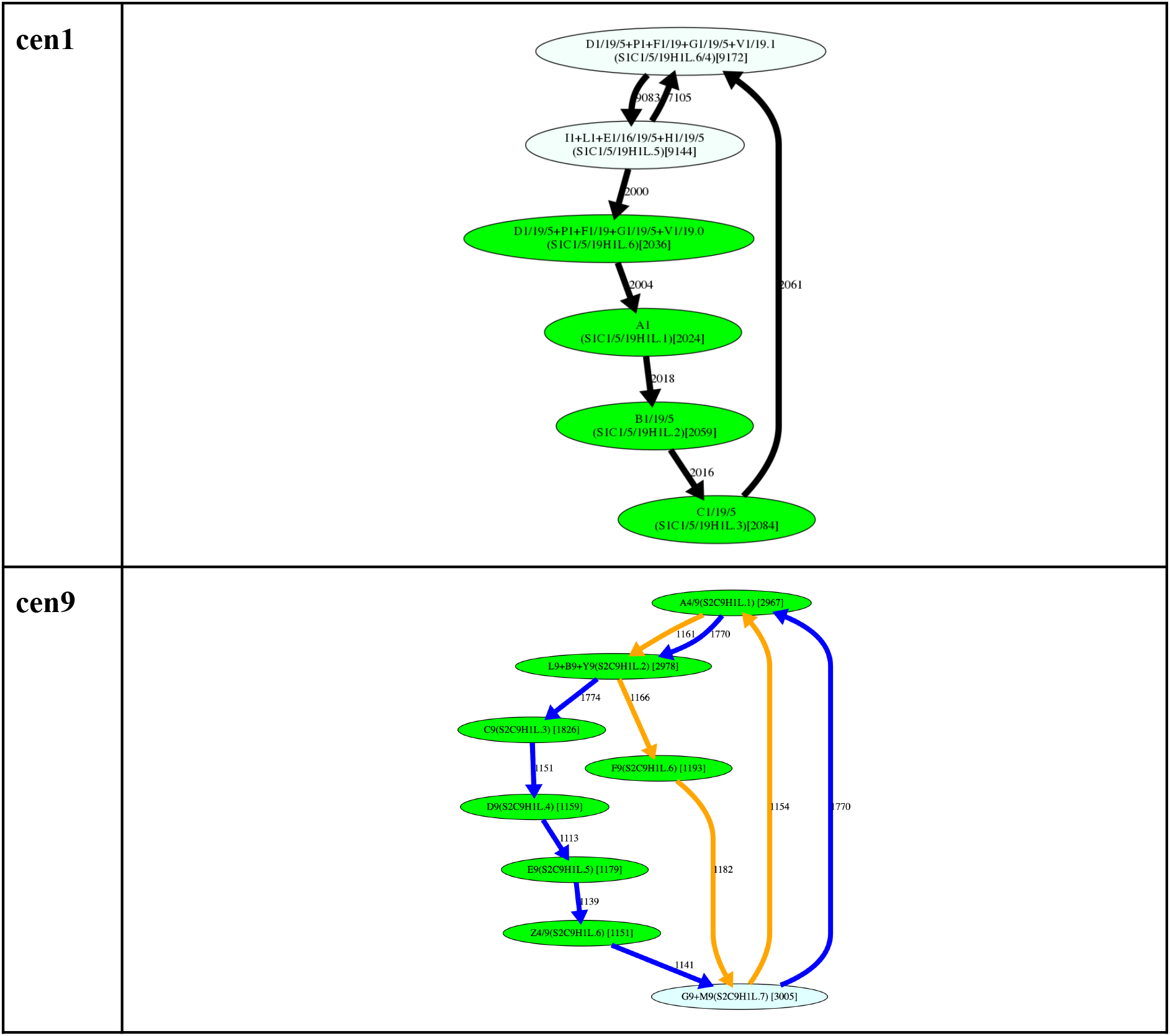
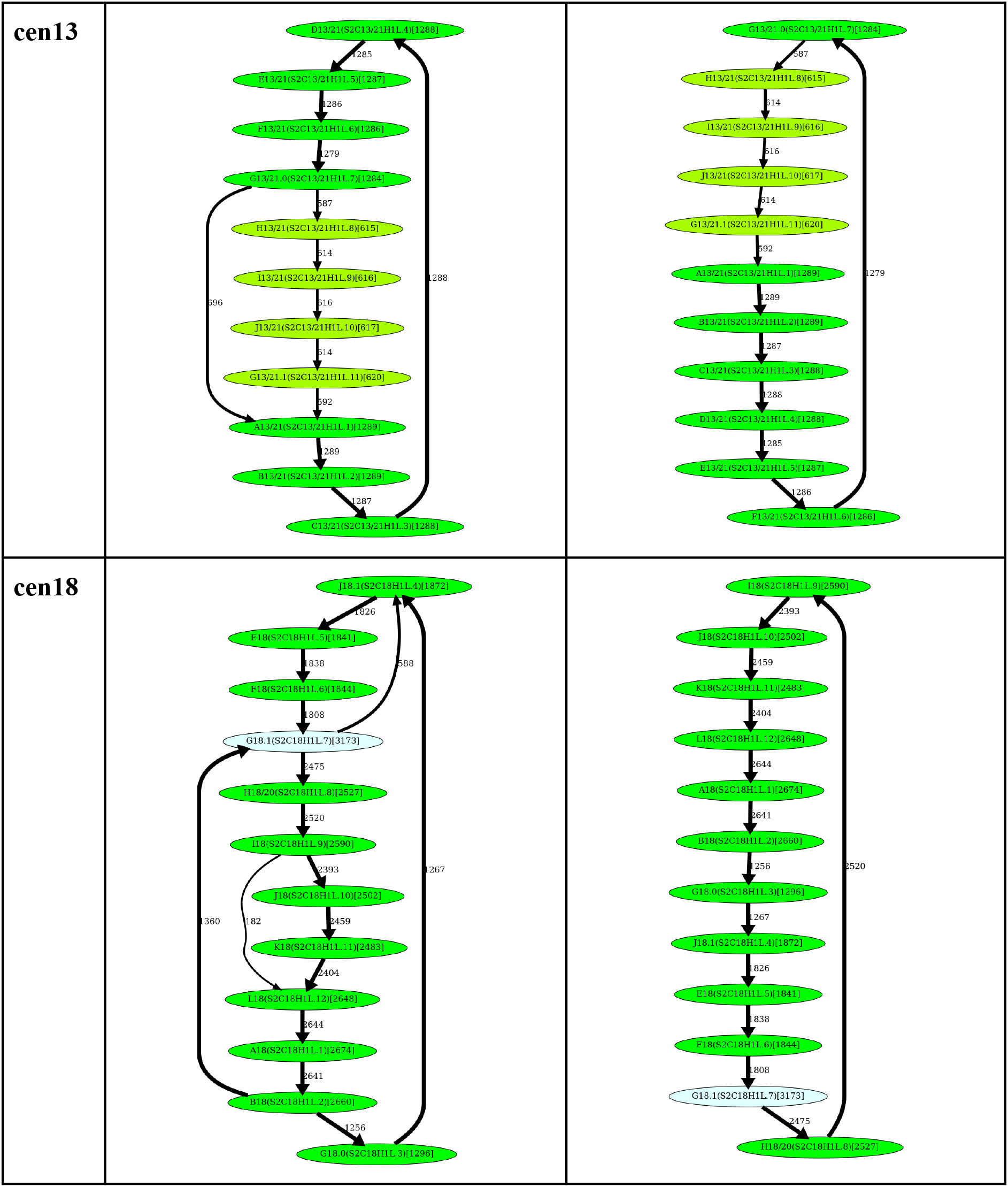
Inferring HORs for cen1, cen9, cen13, and cen18. (Top row) Splitting an unbreakable junction monomer in cen1 results in two monomers with an 11-nucleotides difference and transforms the monomer-graph of cen1 into a cycle with a single chord. **(Second row)** The manually inferred HOR of cen9 (McNulty and Sullivan 2018), shown as the blue cycle, is in conflict with the CE Postulate because the frequently traversed “yellow” cycle contains a monomer that does not belong to the “blue” cycle. **(Third row)** Splitting an unbreakable junction monomer in cen13 results in two similar monomers with an only three-nucleotides difference and transforms the monomer-graph of cen13 (Figure SimplifiedMonomerGraphs) into a cycle with a single chord shown on the left. The resulting simplified monomer-graph (shown on the right) reveals the canonical 11-monomer HOR in cen13. **(Bottom row)** Splitting an unbreakable junction monomer in cen18 results in two nearly identical monomers with only a single-nucleotide difference and transforms the simplified monomer-graph of cen18 (Figure SimplifiedMonomerGraphs) into a cycle with three chords shown on the left). The resulting simplified monomer-graph (shown on the right) reveals the canonical 12-monomer HOR in cen18.

Splitting a junction monomer in cen13 (cen18) results in two monomers that differ in only 3 (1) nucleotides. The split of the unbreakable vertex G into vertices G.0 and G.1 results in two traversals F-G.0-H and J-G.1-A (Figure ComplexCentromeres). Further launch of StringDecomposer (using monomers G.0 and G.1 instead of G) confirms that there are no traversals F-G.0-A and J-G.1-H.

Splitting a junction monomer in cen18 results in two nearly identical monomers that differ in a single nucleotide and raises a concern about the applicability of the CE Postulate to cen18. Splitting the unbreakable monomer G in cen18 should result in two traversals B-G.0-J and F-G.1-H. However, the further launch of StringDecomposer shows 547 B-G.1-J traversals and 10 F-G.0-H traversals. Importantly, in all B-G.1-J (F-G.0-H) traversals, the monomer-block G.1(G.0) is more similar (or even identical!) to the monomer G.1(G.0). Although this raises a concern about the validity of splitting the unbreakable monomer G in cen18, we proceed with the split to be consistent with the CE Postulate.

### Dehybridization reveals HORs in cen5 and cen8

We identified all hybrid monomers (among monomers in *MonomersNew*^*+*^ across all centromeres) using the approach described in “Supplementary Note: Inference of hybrid monomers”. This analysis revealed only three frequent hybrid monomers: P5, R1/5/19, and L8. Below we describe the *dehybridization* operation on monomer-graphs that reveals HORs in cen5 and cen8.

#### Dehybridization in cen5

P5 is a hybrid monomer of S5 and D5 that differs from S5(50)/D5(120) in 5 nucleotides, while R1/5/19 is a hybrid monomer of B5 and D5 that differs from B5(92)/D5(78) in 6 nucleotides. Figure cen5cen8 (top) illustrates that dehybridization of P5 and R1/5/19 resuls in a graph with a single Hamiltonian cycle that is classified as a HOR.

**Figure cen5cen8.**
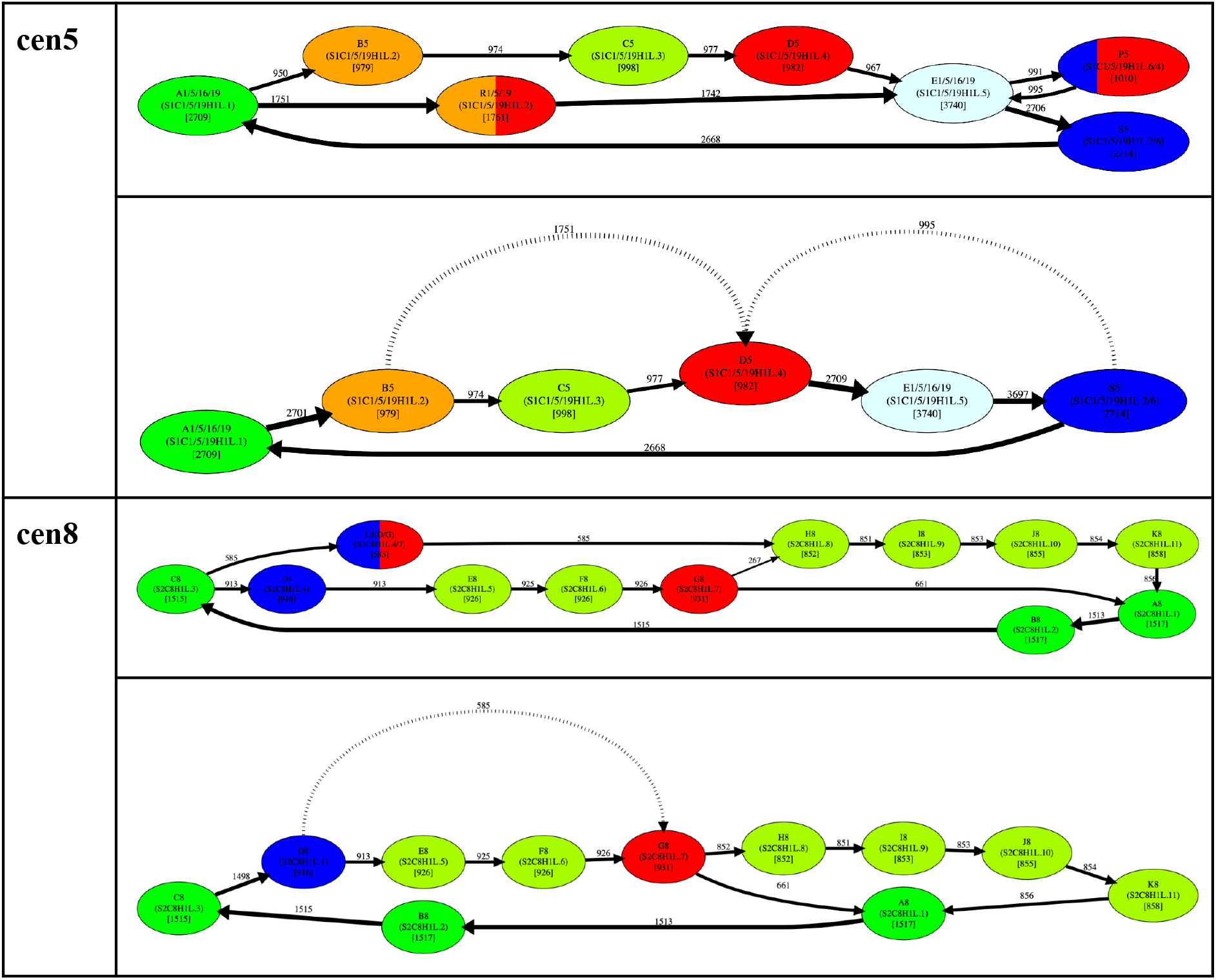
Dehybridization substitutes hybrid vertices (monomers) by hybrid edges in the monomer graph. (Top) Dehybridization of P5 and R1/5/19 in cen5. **(Right)** Dehybridization of L8 in cen8.

#### Dehybridization in cen8

L8 is a hybrid monomer of D8 and G8 which differs from the consensus D8(60)/G8(111) by only 2 nucleotides (see Supplementary Note “HORmon monomer naming”). Figure cen5cen8 (bottom) illustrates the dehybridization of L8 that models it as a hybrid edge of the monomer-graph, resulting in a graph with a single Hamiltonian cycle (and two chords) that is classified as a HOR.

### What is a HOR in cen9ã

Splitting unbreakable junction vertices (cen1, cen13, and cen18) and dehybridization (cen5 and cen8) reveal evolutionary adequate HORs for all centromeres except for cen9. This centromere represents a difficult case from the perspective of the CE Postulate since it is unclear how to infer a HOR from the monomer-graph of this centromere. The blue traversal of this graph (Figure ComplexCentromeres) corresponds to the currently known (manually inferred) HOR. The monomer F9 (that does not belong to the HOR in cen9) cannot be represented as a hybrid monomer and is quite different from its most similar monomer in cen9 (it differs from Z4/9 by 12 nucleotides). Thus, it is not clear how to automatically derive HOR for cen9.

One can argue that merging monomers F9 and Z4/9 would reveal a Hamiltonian cycle (HOR) in the resulting monomer-graph, thus extending the CE Postulate to cen9. This argument reflects the difficulty of developing an automated approach to centromere annotation and defining parameters of these approaches that work across all centromeres. Indeed, the CE Postulate is highly dependent on parameters, e.g., relaxing the parameter for monomer merging will affect the monomer-graphs for all centromeres and may “break” the CE Postulate for some of them. Although by manually fitting parameters for each centromere one can make it look consistent with the CE postulate, such an approach does not represent solid supporting evidence for this postulate. As described in Supplementary Note “HORmon parameters”, due to the limited data (only a single human genome has been completely assembled so far), it is challenging to avoid overfitting even for the default parameters of HORmon, let alone the defaults for a more complex procedure. Our approach represents the first automated analysis (with the same parameters for all centromeres) demonstrating that the CE Postulate holds for nearly all centromeres.

### Generating the centromere decomposition into HORs

HORmon decomposes each monocentromere into canonical, partial, and auxiliary HORs as described in subsection “Decomposing centromere into HORs’’ in Methods (Figure HORmonPipeline). Given a canonical HOR *H=M*_*1*_, *…, M*_*n*_, each canonical HOR *M*_*i*_, *…, M*_*n*,_,*M*_*n+1*_, *…, M*_*i-1*_, in the decomposition is labeled as *c*_*i*_. We use the notation *c*_m_ to denote *m* consecutive occurrences of a canonical HOR and refer to each such element in the HOR decomposition as a *HOR-run*. According to the CE Postulate, hybrid and infrequent monomers do not belong to the HOR.

The *length* of the HOR decomposition is defined as the total number of elements in this decomposition (each entry *x*^*y*^ is counted as a single element). Figure cenXdecomposition shows the HOR decompositions of cenX under the assumption that the monomer-set includes all HOR-monomers AB…KL, as well as hybrid monomers M and N identified in Dvorkina et al., 2020 Supplementary Table HORDecompositions and Supplementary File AllCenDecomposition provide information about the HOR decompositions for all human centromeres. Supplementary Note “Generating the nucleotide consensus of a HOR” describes how these HOR decompositions are used to generate the nucleotide consensus of each HOR. Supplementary Table MonomerNaming summarizes information about these consensus for “live’’ human centromeres. Since these consensus sequences are computed for the first time using a complete human genome assembly, they characterize the CHM13 cell line more accurately than previously inferred sequences. The question of whether they are representative for other individuals remains open.

**Figure cenXdecomposition.**
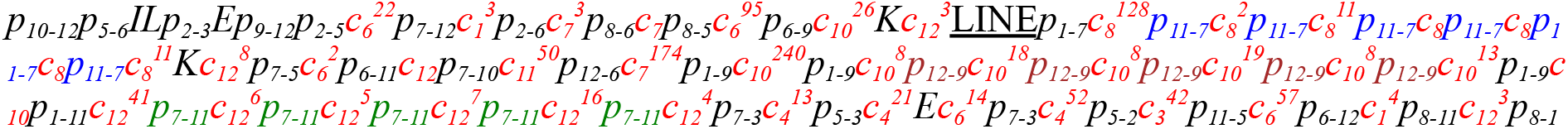
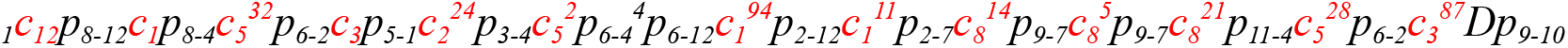
Decomposition of cenX into HORs. The 12-monomer HOR for cenX is represented as *M*_*1*_, *…, M*_*12*_=AB…KL. The monomer-set includes these 12 frequent monomers as well as hybrid monomers M (a hybrid of monomers J and H) and N (a hybrid of monomers K and J) identified in Dvorkina et al., 2020. Each occurrence of this HOR that starts from the monomer *M*_*i*_ is labeled as *c*_*i*_ (shown in red). Each occurrence of a partial HOR that includes monomers from *i* to *j* is labeled as *p*,_*j*_. The most frequent partial monomers *p*_3-7_, *p*_7-3_, and *p*_5-2_ in cenX are colored in blue, green, and brown, respectively. The HOR decomposition of cenX has a length 72 and includes 1486 complete HORs that form 34 HOR-runs. Only 257 out of 18089 (1.4%) monomer-blocks in cenX are not covered by complete HORs. The “<u>LINE</u>” entry shows the position of the LINE element. To ensure that all monomers are shown in the forward strand, we decompose the reverse complement of cenX and take reverse-complements of all monomers in cenX (Supplementary Table MonomerNaming).

Pairs of centromeres (13, 21) and (14, 22), as well as triple of centromeres (1, 5, 19), have been reported to share the same HOR (McNulty and Sullivan, 2018). Contrary to previous studies, we conclude that HORs in these centromeres are rather different, at least in CHM13 cell line. Interestingly, the edit distance between the consensus of HORs in cen13 and cen21 is rather high (20 differences, 1% divergence) while the edit distance between the consensus of HORs in cen14 and cen22 is much lower (3 differences. 0.2% divergence). Previous studies reported two frequent non-hybrid monomers for centromeres 1, 5, and 19 (McNulty and Sullivan, 2018). We report six frequent non-hybrid monomers for cen1 and cen5, and two for cen19. We hypothesize that these minor differences are due to the absence of a complete genome assembly in prior studies. Sequence comparison shows that the edit distance between the consensus of HOR in cen1 and cen5 is large (34 differences, 3.3% divergence).

## Methods

### Positionally-similar monomers

Given a monomer *M* in a monomer-set *Monomers* for a given monocentromere, we identify all triples of consecutive blocks *XMY* that appear in this monocentromere, and construct the |*Monomers*|*|*Monomers*| matrix *Triples*_*M*_, where *Triples*_*M*_(*X,Y*) is the count of the number of triples *XMY* in the monocentromere. We further construct a normalized matrix *NormalizedTriples*_*M*_(*X,Y)* by multiplying *Triples*_*M*_(*X,Y*) by a constant so that the squared sum of all its entries is equal to one.

Given two equally-sized *n*×*m* matrices *A* and *B*, we define their *similarity* as the dot-product of the *n*×*m-*dimensional vectors representing these matrices:

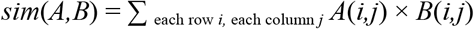

Given two monomers *M* and *M’*, we define their *positional similarity PosSim*(*M,M’*) as

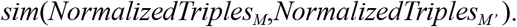

Two monomers are called *similar* if the percent identity between them exceeds a threshold *minPI* (default value 94%). Two similar monomers are called *positionally-similar* if their positional similarity exceeds a threshold *minPosSim* (default value 0.4).

### Merging positionally-similar monomers

Since two different positionally-similar monomers point to a potentially erroneous splitting of a single monomer, HORmon checks if there are positionally-similar monomer-pairs in the monomer-set *Monomers*. If such monomer-pairs exist, it iteratively identifies a pair of the most positionally-similar monomers (similar monomers with the highest positional similarity of all similar monomers), merges them into a new monomer, recomputes the consensus of the new monomer, launches StringDecomposer on the new (smaller) monomer-set, and iterates until there are no positionally-similar monomers left. Similarly to constructing the triple matrices for all triples *XMY* of a monomer *M*, HORmon constructs similar matrices for all triples *XYM* and *MXY* and merges monomers based on these two matrices in the same way it merges monomers for all triples *XMY*.

### Splitting aggregated monomers

To decide whether to split a monomer *M*, HORmon analyzes all frequent triples *XMY* in a monocentromere. Given a monomer *M*, we refer to the largest element in the matrix *NormalizedTriples*^*M*^(*X,Y)* as the *M-champion*. We classify elements (*X,Y*) and (*X’,Y’*) in the matrix *NormalizedTriples*^*M*^(*X,Y)* as *M*-*comparable* if

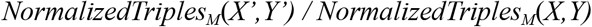

exceeds a *splitting threshold splitValue* (default value 1/8). HORmon uses the single linkage clustering to iteratively identify all monomer-pairs (*X,Y*) that are *M-*comparable with the *M*-champion and refer to them as *M-candidate-pairs*.

Monomer-pairs (*X,Y*) and (*X’,Y’*) are called *independent* if all four monomers *X, Y, X’*, and *Y*’ are different. A monomer *M* that has *M*-candidate-pairs is called *breakable* if all *M*-candidate-pairs are (pairwise) independent, and *unbreakable*, otherwise. Given a breakable monomer *M*, HORmon considers all *M*-candidate-pairs (*X*_*1*_,*Y*_*1*_), …, (*X*_*t*_,*Y*_*t*_) and splits the monomer *M* into *t* monomers *M*_*1*_, …, *M*_*t*_ by separately deriving the monomers *M*_*i*_ as the consensus of all *M*-blocks that arise from triples *X*_*i*_*MY*_*i*_ in the monocentromere for 1 ≤ *i* ≤ *t*.

Supplemental Figure SplitAndMerge describes the pseudocode of the SplitAndMerge module that HORmon uses for modifying the initial monomer-set.

### Simplified monomer-graphs

Given a monomer-graph, HORmon constructs the *complete bipartite graph* where each part represents all vertices (monomers) of the monomer-graph. A monomer *M’* in the “upper” part is connected with a monomer *M*’’ in the “bottom’’ part by an edge of the weight equal to the multiplicity of the edge (*M’,M*’’) in the monomer-graph. Afterward, HORmon solves the *assignment problem* to find the *maximum weight bipartite matching* in the bipartite graph (Ahuja et al., 1993). Edges of this bipartite matching, which also represent edges of the monomer-graph, form a set of non-overlapping cycles and paths in the monomer-graph. An edge of a monomer-graph is classified as *removable* if it forms a chord in one of these cycles/paths (a chord of a path is defined as an edge connecting two internal vertices of this path). Removal of all removable edges from the monomer-graph results in the *simplified monomer-graph*.

### Inference of hybrid monomers

HORmon’s algorithm for inferring hybrid monomers differs from the approach in Dvorkina et al., 2021. For monomers *A, B*, and *C*, we define *HybridDivergence*_*A*_(*B, C*) as the divergence between *A* and a concatenate of a prefix of *B* and a suffix of *C* that is most similar to *A*. A monomer *A* from a monomer-set *Monomers* is a *hybrid candidate* of monomers *B* and *C* if *HybridDivergence*_*A*_(*B,C*) is below the *maxResolvedDivergence* threshold and *HybridDivergence*_*A*_(*B,C*) does not exceed divergence between *A* and any another monomer from *Monomers*. HORmon first generates a set *HybridCandidates* by iterating over concatenates of all possible prefixes and suffixes for every pair of distinct monomers *B* and *C*. Afterward, if there is a single pair of monomers *B* and *C* that give rise to a hybrid candidate *A*, we classify *A* as a hybrid of *B* and *C*. If several pairs of such monomers exist, we select a pair of monomers *B* and *C* that are not hybrid candidates themselves, form a concatenate with the minimal divergence from the monomer *A*, and classify *A* as a hybrid of *B* and *C*.

### Decomposing a centromere into HORs

We defined the monomer-graph as the de Bruijn graph with low-multiplicity vertices and edges removed. We now consider the complete de Bruijn graph *DB*(*Centromere\**, 2) and classify an edge in this graph as a *HOR-edge* if it connects two consecutive monomers in a HOR, and a *non-HOR-edge*, otherwise. A monocentromere defines a traversal of edges in the de Bruijn graph (that contains both HOR-edges and non-HOR-edges) and each *non-HOR-*edge in this traversal corresponds to two consecutive monomers in the monocentromere that we refer to as *breakpoin*t. We break the monocentromere at all breakpoints defined by non-HOR-edges, resulting in multiple short substrings. These substrings, that define the HOR decomposition of a centromere, represent one of the following scenarios:

- a canonical HOR or multiple consequently traversed canonical HORs that may be followed by a partial HOR.
- a partial HOR that includes monomers from *i* to *j* denoted *p*_i_,_*j*_. Since a HOR is a cycle, *i* might exceed *j*, for example, *p*_4,2_ corresponds to the partial 4-monomer HOR *M*_4_,*M*_5_,*M*_1_,*M*_2_ for a 5-monomer HOR *M*_1_,*M*_2_,*M*_3_,*M*_4_,*M*_5_,
- an auxiliary HOR represented by a hybrid or an infrequent monomer (denoted by the identifier of this monomer).

## Discussion

Recent advances in long-read sequencing technologies and genome assembly algorithms opened new horizons for centromere genomics. For the first time, studies of human alpha satellite arrays can be based on a complete centromere assembly rather than individual reads or *satellite reference models* (Miga et al., 2014). The development of an automated centromere annotation tool is a prerequisite for future centromere research that quickly moves to the stage when the complete genomes of hundreds of individuals will be assembled. These studies include population-wide analysis of human monomers and HORs, evolutionary studies of centromeres across primates and other species, biomedical studies of diversity of human centromeres and their associations with genetic diseases.

We developed HORmon, the first annotation tool for “live” alpha satellite arrays that considers monomer and HOR inference as two interconnected problems and automatically generates the monomer- and HOR-set that mirror the four decades of centromere research. HORmon not only provides the first automatic procedure for extracting monomers and HORs in “live” alpha satellite arrays but also establishes their nucleotide consensus sequences. This is important since the currently used nucleotide sequences for many of these monomers and HORs have been extracted more than two decades ago (Alexandrov et al., 2001) in the absence of centromere assemblies. We note that since human centromeres are very divergent between individuals, it remains unclear how well the inferred nucleotide consensus of HORs in the CHM13 cell line represents other individuals.

HORmon uses a heuristic approach for monomer and HOR inference rather than popular clustering algorithms (such as *k*-means or hierarchical clustering) because the monomer inference problem differs from the classical clustering problem. E.g., the set of data points (monomer-blocks) is not explicitly given but is implicitly encoded in the centromere and depends on the selection of centers (monomers). Although the choice of the consensus alpha-satellite results in the initial set of monomer-blocks, each selection of a monomer-set affects this initial set and results in a slightly different clustering problem. Moreover, it is not clear how to select the biologically adequate function to measure the distances between data-points and centers. E.g., the sequence divergence function that HORmon uses is clearly limited (it does not take into account the positional information), necessitating the merging/splitting modules in HORmon. It is also unclear how to incorporate hybrid monomers in the framework of the classical clustering problems. To address all these complications, we have designed the HORmon heuristic instead of using the standard clustering approaches.

HORmon introduced a procedure for decomposing a centromere into HORs and generated the UCSC genome browser tracks representing this decomposition for the CHM13 genome. Even though the recently assembled CHM13 genome does not include chromosome Y, we project that HORmon will be able to generate monomers and HORs for cenY once its complete assembly becomes available. Uralsky et al., 2019 classified a HOR as *homogeneous* (*divergent*) if its copies have an average divergence below 5% (over 10%). In addition to “live” centromeres that we analyzed in this paper, human chromosomes have nearly sixty pseudocentromeric and divergent HOR arrays. Our next goal is to use HORmon for generating monomers and HORs for these HOR arrays that are still only manually annotated (inferred) in the T2T assembly.

Even though HORmon relies on the CE Postulate to rationalize the series of splits/merges and dehybridizations, computational validation of this postulate remains outside the scope of this paper. Indeed, rigorous statistical analysis of the CE Postulate (together with formulating and analyzing alternative evolutionary hypotheses) is currently lacking. Since the CE Postulate was formed implicitly at the dawn of the sequencing era, we do not rule out a possibility that it might be revised once the statistical significance of HOR extraction for all centromeres is rigorously assessed.

Since only a single human genome remains completely assembled, the selection of HORmon parameters was based on this genome only and thus may suffer from overfitting. Moreover, without a rigorous statistical assessment of the CE Postulate versus alternative models of centromere evolution (Mestrovic et al., 1998, Henikoff et al., 2001, Rice, 2019), it is unclear how to verify that the HORs extracted by HORmon represent the most likely solution of the HOR Inference Problem. To complicate the issue even further, the existing nucleotide sequences of canonical HORs have been extracted decades ago limiting the available “ground truth” to benchmark HORmon against. We anticipate that the HORmon pipeline will become an important stepping stone for the development of a fully automatic tool for the extraction of HORs and centromere annotation across the human population once more complete assemblies become available.

Since the rapidly evolving centromeres are very diverse across the human population (Miga et al., 2014, Suzuki et al., 2020), we anticipate that the concepts of the monomer-graph will assist in comparing centromeres across multiple individuals. Although HORmon proved to be useful for analyzing “live” centromeres, automatic procedures for annotating other alpha satellite domains (both HOR and monomeric) are currently not established. We project that HORmon will work just as well on all homogeneous HORs (not only the “live” ones). Other HOR arrays however are known to be more divergent than the “live” arrays, and monomeric arrays are yet more divergent, so it is currently unclear how to universally select HORmon parameters to annotate all alpha satellite arrays in the human genome.

## Supporting information

Supplementary File AllHORDecompositions

## Acknowledgments

We are grateful to Karen Miga, Aleksei Shpilman, and Cynthia Wu for many insightful comments.

## Author contributions

HORmon algorithm development: A.V.B, O.K, T.D., and P.A.P. HORmon code development: O.K and T.D. Manually-curated ground-truth data: I.A. Manuscript draft: A.V.B and P.A.P. Editing: all authors. Conceptualization: P.A.P.

## Materials & Correspondence

The codebase of HORmon is located at https://github.com/ablab/HORmon/tree/main. Monomer and HOR decompositions of alpha satellite arrays in the CHM13 cell line are available at Figshare https://doi.org/10.6084/m9.figshare.16755097.v1. Jupyter notebook that reproduces figures in this paper is available at https://github.com/TanyaDvorkina/hormon_paper/blob/dev/HORmon_paper.ipynb. A.V.B. is the corresponding author.

