## Supplementary Tables, Figures, and Notes for "HORmon: automated annotation of human centromeres"

### Supplementary Figures

```
SplitAndMerge(Centromere, Monomers, minPI, minPosSim, splitValue)
while there are positionally-similar monomers in Monomers (wrt minPI and minPosSim)
   identify the most positionally-similar monomers M' and M'' in Monomers
   identify a new monomer M as the consensus of all M'-blocks and M''-blocks in Centromere
   remove monomers M' and M'' from Monomers
   add monomer M to Monomers
for each breakable monomer M in Monomers (wrt parameter splitValue)
   for each M-candidate-pair (X,Y)
      identify a new monomer M' as the consensus of all M-blocks in triples XMY
      add M' to Monomers
   remove M from Monomers
return Monomers
```

**Supplementary Figure SplitAndMerge. The pseudocode of the split-and-merge module of HORmon.**

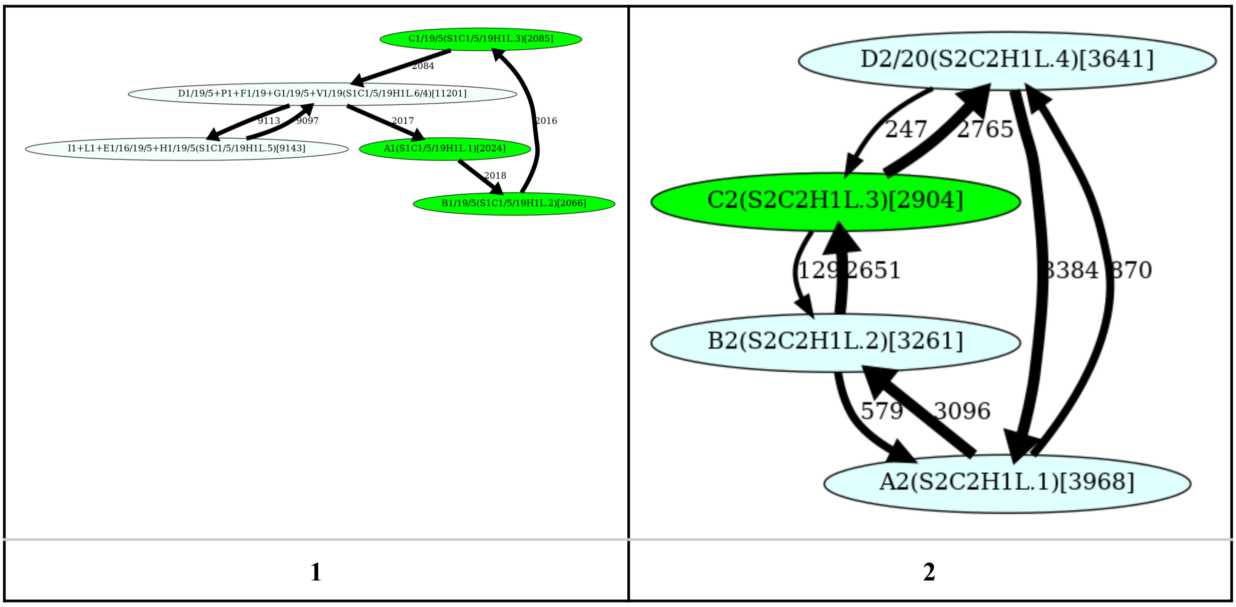

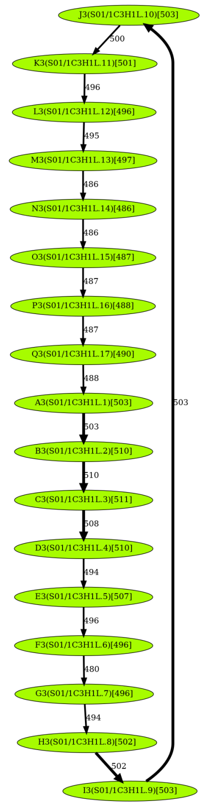
J3(S01/1C3H1L.10)[503]
K3(S01/1C3H1L.11)[501]
L3(S01/1C3H1L.12)[496]
M3(S01/1C3H1L.13)[497]
N3(S01/1C3H1L.14)[486]
O3(S01/1C3H1L.15)[487]
P3(S01/1C3H1L.16)[488]
Q3(S01/1C3H1L.17)[490]
A3(S01/1C3H1L.1)[503]
B3(S01/1C3H1L.2)[510]
C3(S01/1C3H1L.3)[511]
D3(S01/1C3H1L.4)[510]
E3(S01/1C3H1L.5)[507]
F3(S01/1C3H1L.6)[496]
G3(S01/1C3H1L.7)[496]
H3(S01/1C3H1L.8)[502]
I3(S01/1C3H1L.9)[503]

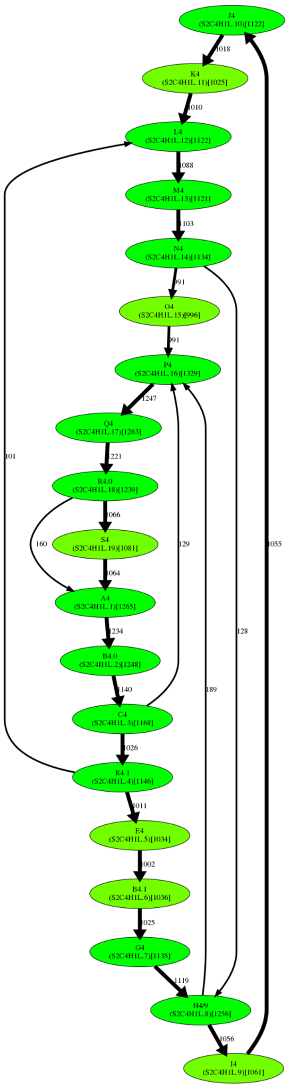

| 3 | 4 |
|---|---|

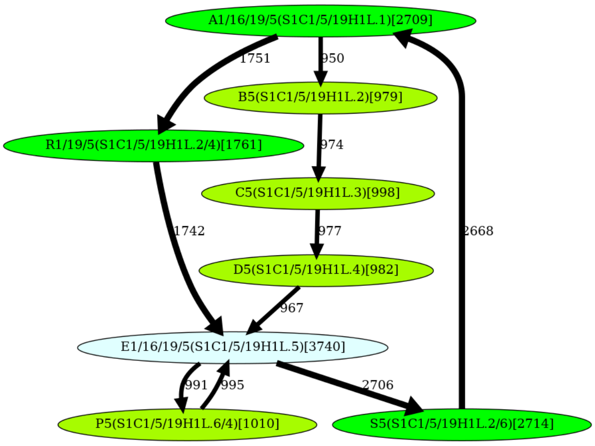
A1/16/19/5(S1C1/5/19H1L.1)[2709]
B5(S1C1/5/19H1L.2)[979]
R1/19/5(S1C1/5/19H1L.2/4)[1761]
C5(S1C1/5/19H1L.3)[998]
D5(S1C1/5/19H1L.4)[982]
E1/16/19/5(S1C1/5/19H1L.5)[3740]
P5(S1C1/5/19H1L.6/4)[1010]
S5(S1C1/5/19H1L.2/6)[2714]
1751
950
974
1742
977
2668
967
991
995
2706

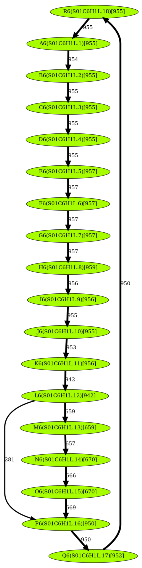

| 5 | 6 |
|---|---|
| 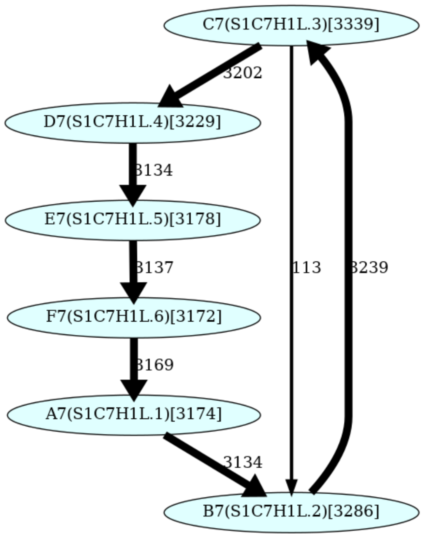C7(S1C7H1L.3)[3339]<br>3202<br>D7(S1C7H1L.4)[3229]<br>3134<br>E7(S1C7H1L.5)[3178]<br>3137<br>F7(S1C7H1L.6)[3172]<br>3169<br>A7(S1C7H1L.1)[3174]<br>3134<br>113<br>3239<br>B7(S1C7H1L.2)[3286] | 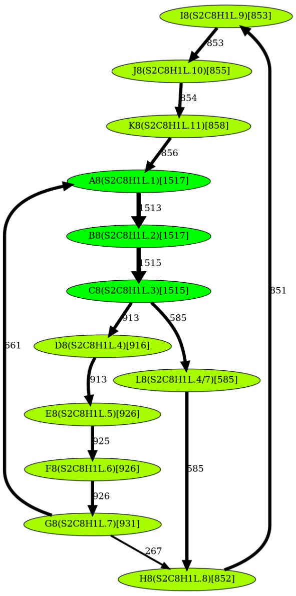I8(S2C8H1L.9)[853]<br>853<br>J8(S2C8H1L.10)[855]<br>854<br>K8(S2C8H1L.11)[858]<br>856<br>A8(S2C8H1L.1)[1517]<br>1513<br>B8(S2C8H1L.2)[1517]<br>1515<br>C8(S2C8H1L.3)[1515]<br>851<br>913<br>585<br>661<br>D8(S2C8H1L.4)[916]<br>913<br>L8(S2C8H1L.4/7)[585]<br>E8(S2C8H1L.5)[926]<br>925<br>F8(S2C8H1L.6)[926]<br>585<br>926<br>G8(S2C8H1L.7)[931]<br>267<br>H8(S2C8H1L.8)[852] |
| 7 | 8 |
| 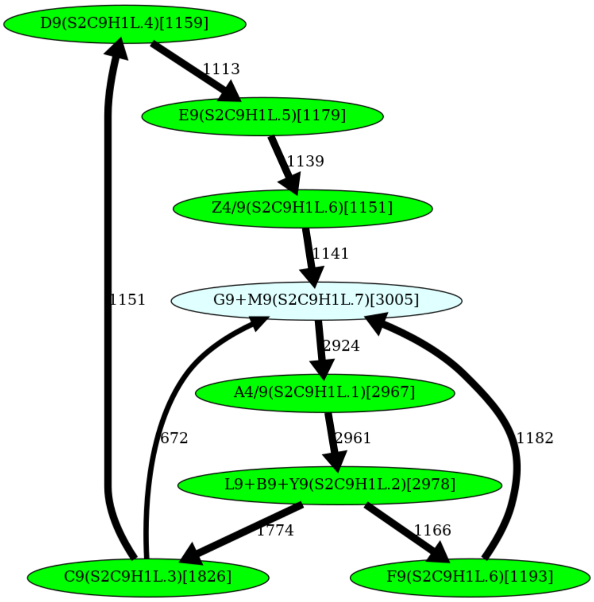D9(S2C9H1L.4)[1159]<br>1113<br>E9(S2C9H1L.5)[1179]<br>1139<br>Z4/9(S2C9H1L.6)[1151]<br>1141<br>1151<br>G9+M9(S2C9H1L.7)[3005]<br>2924<br>A4/9(S2C9H1L.1)[2967]<br>672<br>2961<br>1182<br>L9+B9+Y9(S2C9H1L.2)[2978]<br>1774<br>1166<br>C9(S2C9H1L.3)[1826]<br>F9(S2C9H1L.6)[1193] | 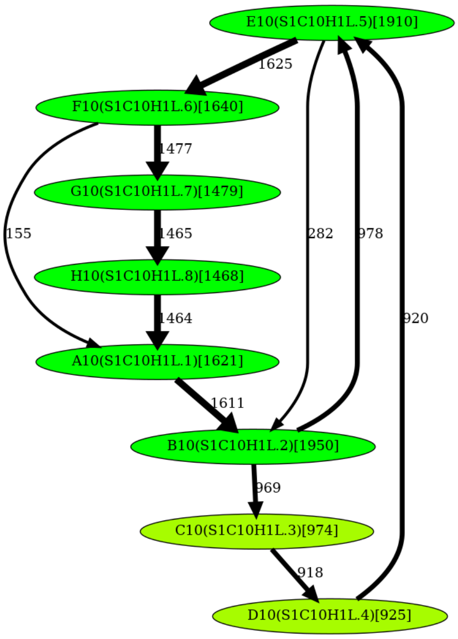E10(S1C10H1L.5)[1910]<br>1625<br>F10(S1C10H1L.6)[1640]<br>1477<br>G10(S1C10H1L.7)[1479]<br>155<br>1465<br>282<br>978<br>H10(S1C10H1L.8)[1468]<br>1464<br>920<br>A10(S1C10H1L.1)[1621]<br>1611<br>B10(S1C10H1L.2)[1950]<br>969<br>C10(S1C10H1L.3)[974]<br>918<br>D10(S1C10H1L.4)[925] |
| 9 | 10 |

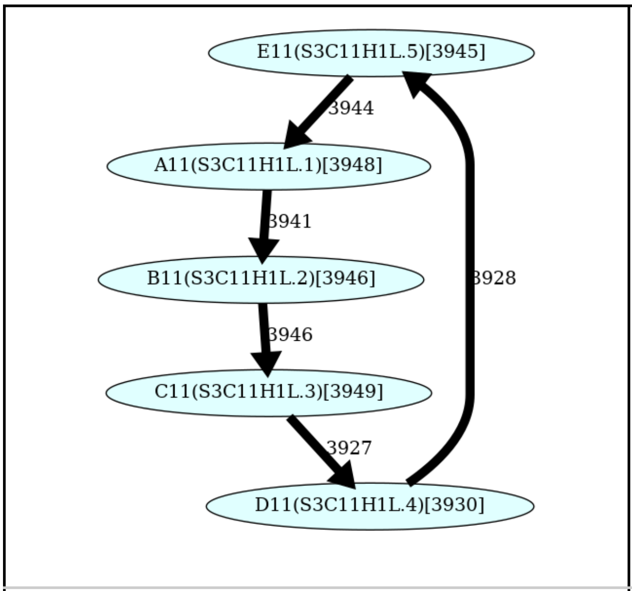

**11**

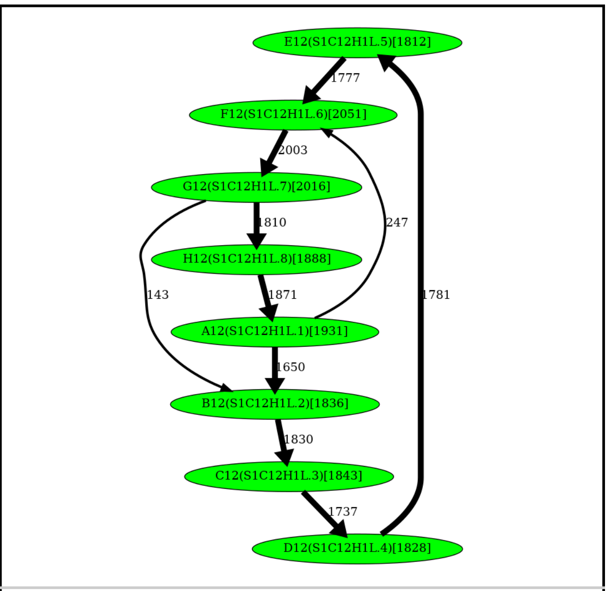

**12**

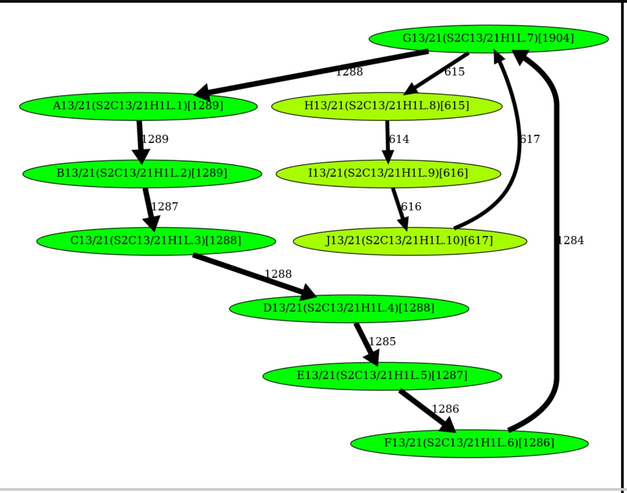

**13**

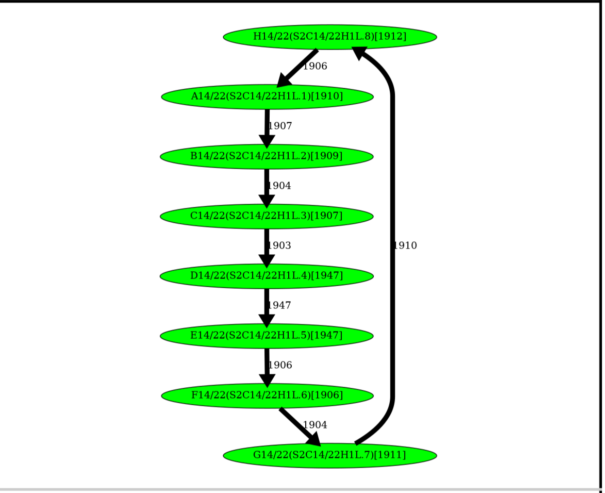

**14**

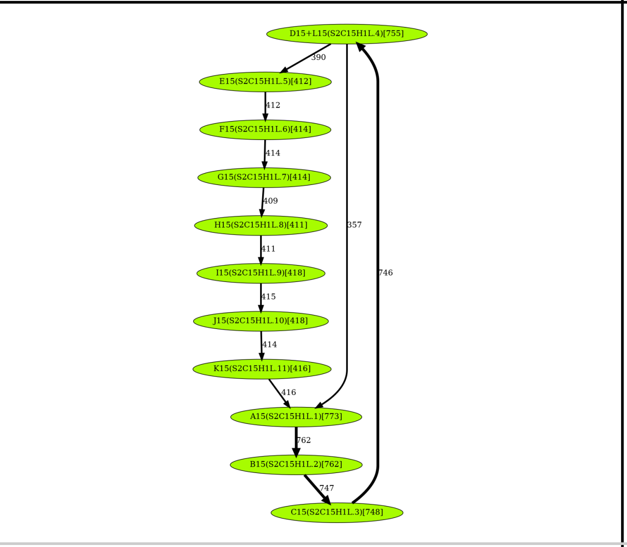

**15**

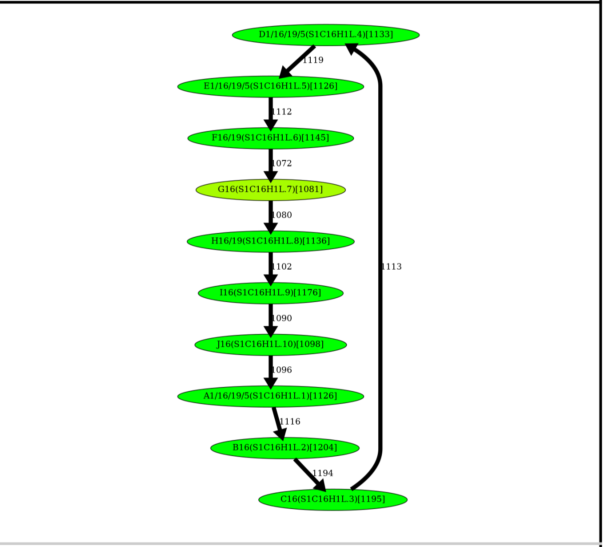

**16**

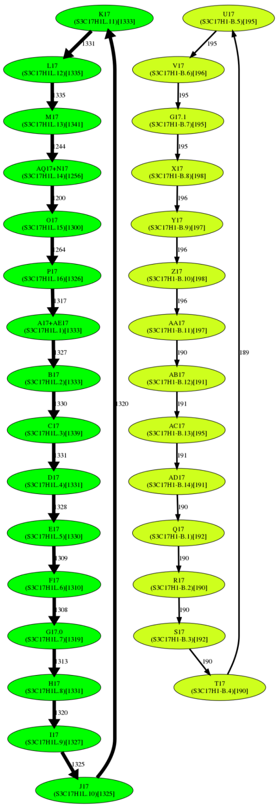

**17**

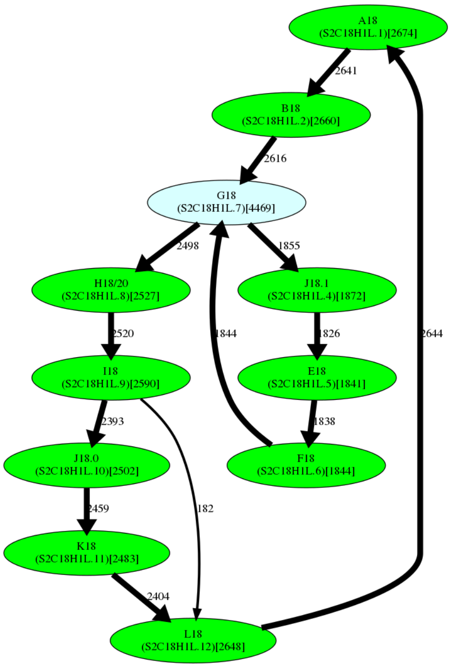
A18
(S2C18H1L.1)[2674]
2641
B18
(S2C18H1L.2)[2660]
2616
G18
(S2C18H1L.7)[4469]
2498
1855
H18/20
(S2C18H1L.8)[2527]
J18.1
(S2C18H1L.4)[1872]
2520
1844
1826
2644
I18
(S2C18H1L.9)[2590]
E18
(S2C18H1L.5)[1841]
2393
1838
J18.0
(S2C18H1L.10)[2502]
F18
(S2C18H1L.6)[1844]
2459
182
K18
(S2C18H1L.11)[2483]
2404
L18
(S2C18H1L.12)[2648]

**18**

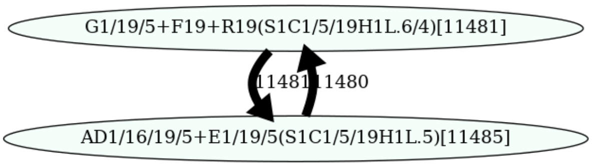
G1/19/5+F19+R19(S1C1/5/19H1L.6/4)[11481]
11481
11480
AD1/16/19/5+E1/19/5(S1C1/5/19H1L.5)[11485]

19

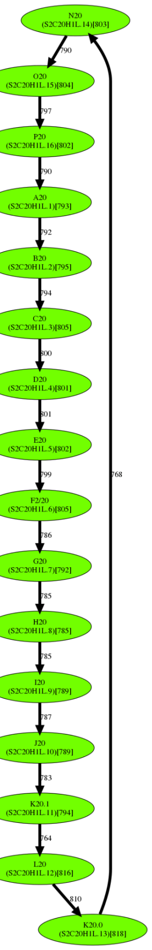
N20 (S2C20H1L.14)[803]
790
O20 (S2C20H1L.15)[804]
797
P20 (S2C20H1L.16)[802]
790
A20 (S2C20H1L.1)[793]
792
B20 (S2C20H1L.2)[795]
794
C20 (S2C20H1L.3)[805]
800
D20 (S2C20H1L.4)[801]
801
E20 (S2C20H1L.5)[802]
799
F2/20 (S2C20H1L.6)[805]
786
G20 (S2C20H1L.7)[792]
785
H20 (S2C20H1L.8)[785]
785
I20 (S2C20H1L.9)[789]
787
J20 (S2C20H1L.10)[789]
783
K20.1 (S2C20H1L.11)[794]
764
L20 (S2C20H1L.12)[816]
810
K20.0 (S2C20H1L.13)[818]
768

20

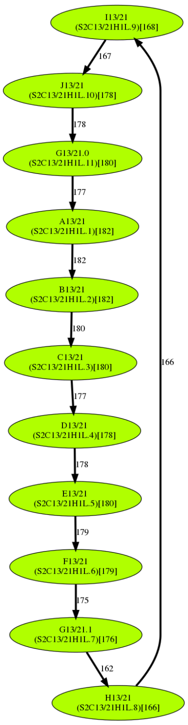
I13/21 (S2C13/21H1L.9)[168]
167
J13/21 (S2C13/21H1L.10)[178]
178
G13/21.0 (S2C13/21H1L.11)[180]
177
A13/21 (S2C13/21H1L.1)[182]
182
B13/21 (S2C13/21H1L.2)[182]
180
C13/21 (S2C13/21H1L.3)[180]
177
D13/21 (S2C13/21H1L.4)[178]
178
E13/21 (S2C13/21H1L.5)[180]
179
F13/21 (S2C13/21H1L.6)[179]
175
G13/21.1 (S2C13/21H1L.7)[176]
162
H13/21 (S2C13/21H1L.8)[166]
166

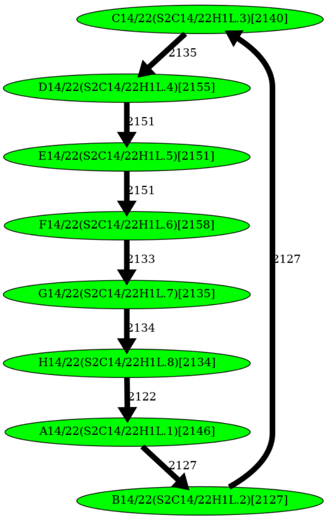
C14/22(S2C14/22H1L.3)[2140]
2135
D14/22(S2C14/22H1L.4)[2155]
2151
E14/22(S2C14/22H1L.5)[2151]
2151
F14/22(S2C14/22H1L.6)[2158]
2133
G14/22(S2C14/22H1L.7)[2135]
2134
H14/22(S2C14/22H1L.8)[2134]
2122
A14/22(S2C14/22H1L.1)[2146]
2127
B14/22(S2C14/22H1L.2)[2127]
2127

| 21 | 22 |
|---|---|
| 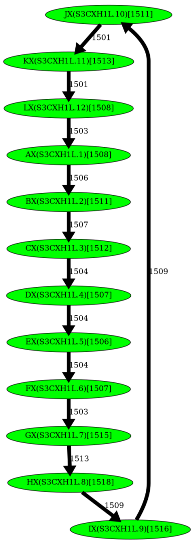  | |
| **X** | |

**Supplementary Figure MonomerGraphs. The monomer-graphs constructed by HORmon for each human centromere.** The label of each vertex represents the monomer ID (and the ID that is currently used by the T2T consortium that follows Uralsky et al., 2019) and its count in the monocentromere (in parentheses). The rules of monomer naming are described in the Supplementary Note “HORmon monomer naming.” The label of an edge in the monomer-graph represents its multiplicity. The width of an edge (color of a vertex) reflects its multiplicity (count of a monomer).

#### Supplementary Tables

| cen | #monomers in *MonomersNew/ MonomersFinal* | #edges in the monomer-graph | #merge/split operations | squared error distortion for *MonomersFinal* | Davies-Bouldin index for *MonomersFinal* |
|---|---|---|---|---|---|
| 1 | 12/5 | 6 | 7/0 | 3.15 | 3.03 |
| 2 | 4/4 | 8 | 0/0 | 2.34 | 1.93 |
| 3 | 17/17 | 17 | 0/0 | 1.79 | 1.67 |
| 4 | 17/19 | 24 | 0/2 | 1.31 | 4.96 |
| 5 | 8/8 | 10 | 0/0 | 1.62 | 1.79 |
| 6 | 18/18 | 19 | 0/0 | 0.93 | 1.57 |
| 7 | 6/6 | 7 | 0/0 | 1.40 | 6.25 |
| 8 | 12/12 | 14 | 0/0 | 1.20 | 3.34 |
| 9 | 11/8 | 10 | 3/0 | 2.48 | 2.88 |
| 10 | 8/8 | 11 | 0/0 | 1.96 | 1.65 |
| 11 | 5/5 | 5 | 0/0 | 1.67 | 0.89 |
| 12 | 8/8 | 10 | 0/0 | 1.87 | 2.66 |
| 13 | 10/10 | 11 | 0/0 | 1.01 | 1.78 |
| 14 | 8/8 | 8 | 0/0 | 1.44 | 2.55 |
| 15 | 12/11 | 12 | 1/0 | 2.02 | 2.02 |
| 16 | 10/10 | 10 | 0/0 | 1.55 | 2.23 |
| 17 | 31/30 | 30 | 2/1 | 1.37 | 2.95 |
| 18 | 10/11 | 13 | 0/1 | 1.48 | 2.60 |
| 19 | 5/2 | 2 | 3/0 | 3.33 | 1.20 |
| 20 | 15/16 | 16 | 0/1 | 1.33 | 2.97 |
| 21 | 10/11 | 11 | 0/1 | 1.55 | 1.44 |
| 22 | 8/8 | 8 | 0/0 | 1.55 | 2.70 |
| X | 12/12 | 12 | 0/0 | 1.42 | 1.60 |

**Supplementary Table ModifiedMonomerSet. Information about the monomer-set *MonomersFinal* for each human centromere.** The centromere ID (first column), number of monomers in the monomer-sets *MonomersNew* and *MonomersFinal* (second column), number of edges in the monomer-graph for the monomer-set *MonomersFinal* (third column), number of merging/splitting operations performed to generate the monomer-sets *MonomersFinal* (fourth column), the squared error distortion and (fifth column), and the Davies-Bouldin index (sixth column) for the monomer-set *MonomersFinal*. Rows highlighted in blue correspond to centromeres with the monomer-set that has been affected by the split-merge transformations.

| cen | # monomer/ monomer-bloks | # canonical HORs/ runs of canonical HORs | # / % monomers not covered by canonical HORs | 3 most frequent non-canonical HORs/ # their occurrences | length of HOR decomposition | HORs in the *MonomersT2T* alphabet |
|---|---|---|---|---|---|---|

| | | | | | | |
|---|---|---|---|---|---|---|
| **1** | 12/25770 | 1916/1824 | 12434/51 | p4-5/2794 p4-5'/1794<br>I1+L1+E1/5/16/19+H1/5/19/1271 | 6679 | S1C1/5/19H1L.123456 |
| **2** | 4/13450 | 2665/1026 | 2790/20 | A2/571<br>p4-2/323 p4-1/238 | 2681 | S2C2H1L.1234 |
| **3** | 17/8252 | 325/27 | 2727/33 | p8-6'/69<br>p6-4'/65<br>F3'/16 | 138 | S01/1C3H1L.123456789(10)(11)(12)(13)(14)(15)(16)(17) |
| **4** | 19/21429 | 766/288 | 6875/32 | H4/9/124 p1-3/116 p17-18/95 | 1407 | S2C4H1L.123456789(10)(11)(12)(13)(14)(15)(16)(17)(18)(19) |
| **5** | 8/14843 | 903/889 | 9425/63 | R1/5/19/1757 p5-1/1727 P5/999 | 6512 | S1C1/5/19H1L.123456 |
| **6** | 18/16249 | 651/181 | 4531/27 | p16-12/225 p17-12/46 p14-11/9 | 417 | S1C6H1L.123456789(10)(11)(12)(13)(14)(15)(16)(17)(18) |
| **7** | 6/19096 | 3054/209 | 772/4 | p2-4/47<br>C7/33<br>p5-2/31 | 472 | S1C7H1L.123456 |
| **8** | 12/12231 | 559/397 | 6082/49 | L8/585<br>p1-7/348 p1-3/313 | 1760 | S2C8H1L.123456789(10)(11) |
| **9** | 8/14944 | 925/897 | 8469/56 | F9/1176 p7-2/1019 p7-3/624 | 4192 | S2C9H1L.1234567 |
| **10** | 8/11388 | 820/658 | 4828/42 | p5-2/551 E10/360 p1-2/104 | 1769 | S1C10H1L.12345678 |
| **11** | 5/19668 | 3904/56 | 148/0 | p5-2/10<br>p5-1/8<br>p3-1/6 | 108 | S3C11H1L.12345 |
| **12** | 8/14694 | 1348/529 | 3910/26 | p6-1'/186 p2-7'/94<br>p6-4'/93 | 1437 | S1C12H1L.12345678 |
| **13** | 11/11466 | 601/355 | 4856/42 | p1-7'/654<br>p1-6'/9<br>p8-11'/9 | 734 | S2C13/21H1L.123456789(10)(11) |
| **14** | 8/15170 | 1835/83 | 490/3 | G14/22'/41 p4-5'/41<br>p1-5'/16 | 249 | S2C14/22H1L.12345678 |
| **15** | 11/5767 | 354/307 | 1873/32 | p1-4'/280 p9-4'/40<br>p1-6'/30 | 728 | S2C15H1L.123456789(10)(11) |
| **16** | 10/11367 | 999/165 | 1377/12 | I16/70<br>p2-6/51<br>p8-3/41 | 475 | S1C16H1L.123456<br>789(10) |
| **17** | 30/23609 | 188/11<br>1040/112 | 4337/18 | p2-16/120 p15-13/38 P17/25 | 409 | S3C17H1-B.123456789(10)(11)(12)(13)(14)<br>S3C17H1L.123456789(10)(11)(12)(13)(14)(15)(16) |
| **18** | 12/28041 | 1216/430 | 13448/47 | G18.1/623 p7-2/613 p4-2/360 | 2507 | S2C18H1L.123456789(10)(11)(12) |
| **19** | 2/21126 | 10217/802 | 692/3 | AD1/5/16/19+E1/5/19/641<br>G1/5/19+F19+R19/51 | 1430 | S1C1/5/19H1L.5(6/4) |
| **20** | 16/12744 | 728/120 | 1096/8 | p12-13/32 p14-11/8<br>p7-11/6 | 288 | S2C20H1L.123456789(10)(11)(12)(13)(14)(15)(16) |
| **21** | 11/1941 | 159/22 | 192/9 | p10-7'/11<br>p1-3'/2<br>p4-6'/2 | 52 | S2C13/21H1L.123456789(10)(11) |
| **22** | 8/16897 | 1955/164 | 1257/7 | p1-7'/49<br>p7-5'/47 G14/22'/35 | 389 | S2C14/22H1L.12345678 |
| **X** | 12/18030 | 1464/54 | 462/2 | p11-7'/6<br>p7-11'/5 | 118 | S3CXH1L.123456789(10)(11)(12) |

| | | | | p12-9'/5 |
|---|---|---|---|---|

**Supplementary Table HORDecompositions. Information about HOR decompositions generated by HORmon for all human centromeres.** Each row presents information about the HOR decomposition of a specific monocentromere. The second column presents information about the number of monomers/monomer-blocks. The third column presents information about the number of canonical HORs/HOR-runs of canonical HORs. The fourth column presents information about the number/percentage of monomers not covered by canonical HORs. The fifth column presents information about the three most frequent non-canonical (partial or auxiliary) HORs / the number of their occurrences (HORs with an ' character refer to reverse complemented HORs for centromeres assembled in reverse complemented strand, i.e. 12, 13, 14, 15, 21, 22, and X, and for centromere cen1 that has a reverse complemented substring). The sixth column presents information about the length of the HOR decomposition. The seventh column specifies the HORs for the corresponding monocentromere. A single HOR corresponds to each ("live") centromere, except cen17 where the *sister* HOR (14-mer) resides on the live array (an epiallele) in a fraction of individuals (Supplementary Table SN:HC-1).

| **cen** | **# monomers in *MonomersT2T/ MonomersNew* *** | **# blocks for *MonomersT2T** / *MonomersNew** / *SharedBlocks*** | **squared error distortion for *MonomersT2T** / *MonomersNew**** | **Davies-Bouldin index for *MonomersT2T** / *MonomersNew**** | **# monomers reported by Centromere Architect** | **# monomers in *MonomersT2T*** | **# monomers in *MonomersNew*** |
|---|---|---|---|---|---|---|---|
| 1 | 12 | 26504/26504/26486 | 2.37/**1.57** | **4.22**/7.07 | 23 | 15 | 12 |
| 2 | 4 | 13744/13744/13744 | **2.33/2.33** | 1.96/**1.95** | 10 | 6 | 4 |
| 3 | 17 | 8485/8485/8485 | 1.82/**1.77** | **1.42**/1.44 | 24 | 19 | 17 |
| 4 | 17 | 21715/21715/21711 | 1.72/**1.65** | **2.56**/2.70 | 25 | 19 | 17 |
| 5 | 8 | 14893/14893/14893 | 1.69/**1.62** | **1.73**/1.80 | 19 | 14 | 8 |
| 6 | 18 | 16315/16315/16313 | **0.87/0.87** | 1.49/**1.47** | 19 | 18 | 18 |
| 7 | 6 | 19375/19375/19373 | **1.34/1.34** | **4.23/4.23** | 17 | 6 | 6 |
| 8 | 12 | 12247/12247/12243 | **1.03/1.03** | **1.19/1.19** | 12 | 16 | 12 |
| 9 | 10 | 15456/15456/15455 | 3.51/**2.15** | **2.72**/3.36 | 26 | 10 | 11 |
| 10 | 8 | 11967/11967/11967 | **1.95/1.95** | **1.67**/1.69 | 36 | 19 | 8 |
| 11 | 5 | 19718/19718/19718 | 1.70/**1.67** | **0.89/0.89** | 14 | 6 | 5 |
| 12 | 8 | 15204/15204/15204 | **1.86/1.86** | 2.72/**2.71** | 21 | 8 | 8 |
| 13 | 10 | 11478/11478/11476 | 1.01/**0.92** | **1.01**/1.05 | 14 | 11 | 10 |
| 14 | 8 | 15349/15349/15349 | 1.44/**1.43** | **2.55**/2.57 | 14 | 8 | 8 |
| 15 | 11 | 5941/5941/5941 | **1.76**/1.87 | **2.11**/2.19 | 16 | 11 | 12 |
| 16 | 10 | 11393/11393/11391 | **0.99**/1.47 | **1.45**/1.65 | 16 | 10 | 10 |
| 17 | 16 | 23849/23849/23846 | **2.68/2.68** | **1.96/1.96** | 43 | 16 | 31 |
| 18 | 10 | 28110/28110/28110 | 1.65/**1.59** | 2.28/**2.11** | 20 | 13 | 10 |
| 19 | 5 | 22964/22964/22963 | 2.87/**2.82** | 11.36/**10.81** | 31 | 12 | 5 |
| 20 | 15 | 12793/12793/12786 | **1.48**/1.53 | **2.46**/2.53 | 17 | 16 | 15 |
| 21 | 10 | 1948/1948/1948 | **1.26**/1.67 | **1.03**/1.11 | 14 | 11 | 10 |
| 22 | 8 | 17146/17146/17146 | **1.54/1.54** | 2.85/**2.76** | 13 | 8 | 8 |
| X | 12 | 18095/18095/18095 | **1.41/1.41** | **1.58/1.58** | 14 | 17 | 12 |

**Supplementary Table MonomerComparison1. Comparison of the monomer-set identified by CentromereArchitect (Dvorkina et al., 2021) with the previously inferred monomer-set (Shepelev et al., 2015, Uralsky et al., 2019).** The centromere ID (first column), the number of monomers in the monomer-sets *MonomersT2T** and *MonomersNew** (second column), # blocks for *MonomersT2T*/MonomersNew*/SharedBlocks* (third column), the squared error distortion for the monomer-sets *MonomersT2T** and *MonomersNew** (fourth column), the Davies-Bouldin index for the monomer-sets *MonomersT2T** and *MonomersNew** (fifth columns), the number of monomers in the monomer-set identified by CentromereArchitect (sixth column), the number of monomers in the monomer-set *MonomersT2T* that were inferred by Uralsky et al., 2019 (seventh column), the number of monomers in the monomer-set *MonomersNew* that represent frequent monomers in the CentromereArchitect output (eight column).

| cen | # monomers in *MonomersNew*/ MonomersFinal** | # blocks for *MonomersNew*/ MonomersFinal*/ SharedBlocks* | squared error distortion for *MonomersNew/ MonomersFinal** | Davies-Bouldin index for *MonomersNew* / MonomersFinal** | # monomers in *MonomersFinal* |
|---|---|---|---|---|---|
| 1 | 6 | 26504/26504/26486 | **2.55**/2.60 | 3.44/**3.15** | 6 |
| 4 | 19 | 21715/21715/21711 | 1.55/**1.25** | **2.63**/3.05 | 19 |
| 9 | 8 | 15456/15456/15456 | 2.70/**2.49** | 2.91/**2.89** | 8 |
| 15 | 11 | 5941/5941/5941 | 1.87/**1.77** | 2.19/**2.07** | 11 |
| 17 | 30 | 23849/23849/23846 | 1.37/**1.34** | 2.34/**2.07** | 30 |
| 18 | 12 | 28110/28110/28110 | **1.39**/1.46 | **2.32**/4.58 | 12 |
| 19 | 2 | 22964/22964/22963 | **3.28**/3.32 | **1.22/1.22** | 2 |
| 20 | 16 | 12793/12793/12786 | 1.52/**1.29** | **2.56**/2.83 | 16 |
| 21 | 11 | 1948/1948/1948 | 1.67/**1.55** | **1.08**/1.30 | 11 |

**Supplementary Table MonomerComparison2. Comparison of the monomer-set identified by CentromereArchitect (Dvorkina et al., 2021) with the monomer-set generated by HORmon.** The centromere ID (first column), the number of monomers in the monomer-sets *MonomersNew** and *MonomersFinal** (second column), # blocks for *MonomersNew*/MonomersFinal*/SharedBlocks* (third column), the squared error distortion for the monomer-sets *MonomersNew** and *MonomersFinal** (fourth column), the Davies-Bouldin index for the monomer-sets *MonomersNew** and *MonomersFinal** (fifth columns), the number of monomers in the monomer-set identified by HORmon (sixth column). Since the monomer-sets for centromeres 2, 3, 5, 6, 7, 8, 10, 11, 12, 13, 14, 16, 22, and X were not affected by HOR-guided split-merge transformation, we omit the corresponding rows.

#### Supplementary Notes

1. **HORmon terminology**
2. **Information about datasets**
3. **Evaluating the monomer-sets**
4. **HORmon parameters**
5. **Generating the nucleotide consensus of a HOR**
6. **HORmon monomer naming**

#### Supplementary Note 1: HORmon terminology

Below we summarize the main terminology used in the paper.

*Monomer-block* — repetitive nucleotide sequence that forms repeats of a higher order (*stacked tandem repeats*).

*Monomer* — sequence consensus of a cluster of similar monomer-blocks.

*HOR* (*higher-order repeat*) — a canonical (cyclic) order of monomers specific to each centromere. It is evolutionarily defined as the ancestral and chromosome-specific order of frequent non-hybrid monomers that has evolved into the complex organization of extant centromeres.

*Hybrid monomer* — a monomer obtained by concatenation of a prefix of one monomer with a suffix of another.

*MonomersNew* — set of frequent monomers generated by CentromereArchitect, the input set of monomers for HORmon.

*MonomersNew$^+$* — set of monomers generated by HORmon after the split and merge operations.

*MonomersFinal* — the final set of monomers generated by HORmon.

*MonomersT2T* — set of monomers semi-manually generated by the T2T Consortium.

*Centromere* — nucleotide sequence of a "live" HOR array which hosts a kinetochore.

*Monocentromere* — sequence of monomer-blocks that represent a decomposition of *Centromere* into the input monomer-set.

*Centromere** — monocentromere obtained by decomposition of *Centromere* into the *MonomersNew* monomer-set.

*Centromere*** — monocentromere obtained by decomposition of *Centromere* into the *MonomersFinal* monomer-set.

*Monomer-graph* — a directed graph of a monocentromere constructed on the vertex-set of all monomers and the edge-set formed by all pairs of consecutive monomers in this centromere. The *multiplicity* of an edge (*M,M'*) in the monomer-graph is defined as the number of times the monomer *M'* follows the monomer *M* in the centromere.

*Simplified monomer-graph* — a graph obtained from a monomer-graph by removing all *removable* edges.

#### Supplementary Note 2: Information about datasets

Information about the alpha satellite arrays from the assembly (public release v1.0) of the effectively haploid CHM13 human cell line is available at (https://github.com/nanopore-wgs-consortium/chm13#v10 (NCBI accession number GCA_009914755.2). Supplementary Table SN:HC-1 presents the coordinates of the extracted regions for all "live" human centromere arrays. The alpha satellite array of the newly assembled centromere of chromosome X from HG002 cell line sequenced is available at https://github.com/marbl/CHM13#hg002-chromosome-x (accession number: CP074113).

| chromosome | start | end |
|---|---|---|
| 1 | 121 796 218 | 126 300 656 |
| 2 | 92 333 539 | 94 673 018 |
| 3 | 91 738 494 | 92 596 313 |
| 3 | 92 869 954 | 92 903 597 |
| 3 | 95 863 962 | 96 415 434 |
| 4 | 49 705 249 | 50 433 651 |
| 4 | 52 115 581 | 54 870 604 |

| | | |
|---|---|---|
| 4 | 54 980 385 | 55 199 889 |
| 5 | 47 039 130 | 47 049 658 |
| 5 | 47 077 198 | 49 596 620 |
| 6 | 58 286 939 | 61 058 622 |
| 7 | 60 414 370 | 63 714 496 |
| 8 | 44 243 543 | 46 325 076 |
| 9 | 44 952 789 | 47 582 587 |
| 10 | 39 633 785 | 41 664 580 |
| 11 | 51 061 950 | 54 413 485 |
| 12 | 34 620 831 | 37 202 143 |
| 13 | 16 220 361 | 18 171 058 |
| 14 | 10 149 798 | 12 766 096 |
| 15 | 17 263 917 | 18 275 855 |
| 16 | 35 854 534 | 37 793 358 |
| 17 S3C17H1-B+S3C17H1L | 23 433 664 | 27 487 230 |
| 18 | 15 971 634 | 20 740 248 |
| 19 | 25 846 349 | 29 749 516 |
| 20 | 26 925 846 | 29 099 648 |
| 21 | 11 699 867 | 12 031 015 |
| 22 | 12 816 949 | 15 739 833 |
| X | 57 820 108 | 60 927 196 |

**Supplementary Table SN:HC-1. Coordinates of the alpha satellite arrays in human chromosomes.** Chromosomes 3 (4; 5) contains three (three; two) alpha satellite arrays that are separated by non-monomeric regions of lengths 274 kb and 2960 kb (1682 kb and 110 kb; 27 kb). The coordinates are modified from Dvorkina et al., 2021 to include only the live HOR arrays (without sister HORs). A single exception is the S3C117H1L (D17Z1) array where the sister HOR S3C17H1-B (D17Z1-B) was included. This sister HOR has been shown to represent the live array (an epiallele) in a fraction of individuals (McNulty and Sullivan, 2018).

#### Supplementary Note 3: Evaluating the monomer-sets

Since HORmon utilizes the monomer-set constructed by CentromereArchitect, it is important to compare this monomer-set with the manually-derived monomer-sets. Below we show that the monomer-set automatically constructed by CentromereArchitect marginally improves on the currently known (manually constructed) monomer-set.

Given a monomer-set *Monomers,* we partition a centromere *Centromere* into monomer-blocks *Blocks*. We define the *average radius* of a monomer *M* (denoted as *r*(*M*)) as the average distance between *M* and all *M*-blocks in *Blocks*. Given strings *S'* and *S"*, we denote the edit distances between them as *distance*(*S',S"*).

From the clustering perspective, the monomer-set represents the *centers* of the *data points* formed by the monomer-blocks. We use the *squared error distortion* and the *Davies-Bouldin* index (Davies and Bouldin, 1979) to evaluate the clustering quality. The squared error distortion is defined as follows:

$$distortion(Monomers, Blocks) = 1/|Blocks| * \sum_{\text{each monomer } M \text{ in } Monomers} \sum_{\text{each } M\text{-block } Block \text{ in } Blocks} distance(M, Block)^2.$$

The *Davies-Bouldin* index is defined as follows:

$$DBI(Monomers,Blocks) = 1/|Monomers| * \sum_{\text{each monomer } M} \max_{\text{each monomer } M' \neq M} (r(M)+r(M')) / distance(M, M').$$

The *count* of a monomer *M* (referred to as *count*(*M*)=*count*(*M,Centromere**)) is the number of its occurrences in the monocentromere *Centromere**. HORmon orders all monomers by their decreasing counts and refers to the *i*-th most frequent monomer in *Centromere** as $M_i$. Given the set $Monomers_i = \{M_1, \dots M_i\}$ of *i* most frequent monomers in *Centromere**, we define $count(Monomers_i)$ as the total count of all monomers in this monomer-set. We identify the *imin* as the minimum value of *i* such that $count(Monomers_i)$ exceeds the threshold *MinFraction* · |*Centromere**|, where |*Centromere**| is the length of the monocentromere (the default value *MinFraction* = 0.9). HORmon constructs the set of *frequent* monomers as $Monomers_{imin}$ complemented by monomers $M_{imin+1}$, $M_{imin+2}$, ... with counts exceeding *MinExtension* · $count(M_{imin})$ (the default value *MinExtension* = 0.7). The resulting human monomer-set is referred to as *MonomersNew*.

The sets *MonomersNew* turned out to contain more monomers than the set *MonomersT2T* for all centromeres except for centromeres 3, 4, 10, 13, 17, 18, 20, and 21. Since two clustering solutions of the same set of data points are usually compared for the case when these solutions have the same number of centers, we attempted to select a subset of monomers from the set *MonomersNew* to make it comparable with the set *MonomersT2T* (for each centromere). For each monomer in *MonomersT2T*, we thus identified the closest monomer in *MonomersNew* and constructed the monomer-set *MonomersNew** of the same size as *MonomersT2T* (in this case, we define *MonomersT2T** = *MonomersT2T*). Similarly, if the set *MonomersT2T* contains more monomers than the set *MonomersNew*, for each monomer in *MonomersNew,* we identified the closest monomer in *MonomersT2T* and constructed the monomer-set *MonomersT2T** of the same size as *MonomersNew* (in this case, we define *MonomersNew** = *MonomersNew).*

Supplementary Table MonomerComparison1 compares the monomer-sets *MonomersT2T** and *MonomersNew**. To ensure that this comparison is adequate (i.e., compares two equally-sized sets of centers for the same data points), we define the set of monomer-blocks *SharedBlocks* that are shared between both monomer-sets. For each monomer-block *B'* (in the centromere decomposition into monomers from *MonomersNew**), and an overlapping monomer-block *B"* (in the centromere decomposition into monomers from *MonomersT2T**), a new block *B* is formed by taking the overlap between *B'* and *B"*. We add *B* to the set *SharedBlocks* if its length exceeds the threshold *MinSharedLength* (the default value *MinSharedLength* = 150). Since the block *B* in *SharedBlocks* is typically shorter than blocks *B'* or *B"*, we modify the definition of the distance between *B* and any monomer *M* as the minimum distance between *B* and all substrings of the *M*-consensus, where the *M-consensus* is defined as the consensus of the multiple alignment of all *M*-blocks.

Supplementary Table MonomerComparison1 illustrates that the set *MonomersNew** results in a marginally better clustering of monomer-blocks (with respect to the squared error distortion) than the set *MonomersT2T** (*MonomersT2T** resulted in a lower squared error distortion for centromeres 15, 16, 20, and 21). Both sets demonstrate similar performance with respect to the Davies-Bouldin index.

In order to compare the set *MonomersNew* generated by CentromereArchitect to the final monomer-set *MonomersFinal* generated by HORmon after merging/splitting and hybrid monomer decomposition, we similarly define the monomer-sets *MonomersNew** and *MonomersFinal* of the same size. Supplementary Table MonomerComparison2 illustrates that these sets result in a comparable clustering of monomer-blocks with respect to the analyzed clustering metrics. Specifically, the Davies-Bouldin index (squared error distortion) for *MonomersFinal** does not exceed the same metric for *MonomersNew** for all centromeres except centromeres 4, 18, 20, 21 (1, 18, 19). These results are not surprising since the CE Postulate-guided monomer transformations that HORmon conducts over the set *MonomersNew* are not coordinated with the objective clustering metrics, but are rather dictated by the biological model of a canonical HOR as an ancestral unit.

#### Supplementary Note 4: HORmon parameters

Selecting HORmon parameters is a complex challenge since only a single human genome has been assembled up to date. Moreover, the choice of "ground truth" to benchmark HORmon is limited since (1) the concept of a HOR is computationally poorly defined, (2) the CE Postulate has not been statistically assessed yet, and (3) the nucleotide sequences of the manually extracted HORs have been generated decades ago at the dawn of the sequencing era. Below we describe the rationale behind selecting HORmon parameters and limitations for their selection.

- *Generating the monomer-set.* HORmon transforms the monomer-set generated by CentromereArchitect into the set *MonomersNew.* This transformation relies on parameters *MinFraction* (default value = 0.9) and *MinExtension* (default value = 0.7) that were chosen by analyzing the counts of monomers in the set monomer-set generated by CentromereArchitect. Filtering out infrequent monomers was straightforward: even using a single parameter *MinFraction* (formally, setting *MinExtension* = 1 under the assumption that no two monomers share the same count) was sufficient to generate a reasonable monomer-set *MonomersNew* for most centromeres. However, in rare cases, we observed that HORmon filters out monomers that are included in the manually constructed monomer-sets. To ensure comparability with previous studies on all centromeres, we introduced an additional parameter *MinExtension* that allowed us to include these monomers, otherwise, filtered monomers in the monomer-set *MonomersNew*.

- *Split and merge transformations.* Splitting a monomer relies on a single parameter *splitValue* (default value 1/8). Merging monomers relies on parameters *minPI* (default value 94%) and *minPosSim* (default value 0.4). Since the length of all monomer-blocks is close to 171 bp, *minPI* corresponds to the maximum edit distance 10 between two monomers. In selecting all these parameters, we tried to ensure that the resulting monomer graphs are topologically close to "a cycle with a few chords". Typically, a lower (higher) value of *splitValue* ensures more relaxed (strict) conditions for splitting monomers. Setting an extreme value *splitValue* = 0 results in splitting each monomer with in-degree $n$ and out-degree $m$ into $n \cdot m$ monomers. On the other hand, setting an extreme value *splitValue* = 1 does not alter the initial monomer-set. A similar rationale is applicable to parameters *minPI* and *minPosSim*. Extensive benchmarking on additional assemblies, once they are available, is required to rule out potential overfit.

- *Monomer graphs*. Construction of the monomer-graphs relies on parameters the *MinEdgeMultiplicity* (default value 100) and *minCountFraction* (default value 0.9). It is possible to eliminate these parameters (formally, select *MinEdgeMultiplicity* = *minCountFraction* = 0) and include edges of all multiplicities into the monomer-graph. However, since the resulting monomer graphs often contain many low multiplicity edges, the included low-multiplicity edges will complicate the analysis of the main "trends" in the centromere architecture (Figure MonomerGraphX). Introduction of both parameters rather than a single parameter *MinEdgeMultiplicity* is necessary for shorter centromeres. For example, cen21, which is only 331,148 bp long, contains only ~2000 monomer-blocks. Selection of these parameters does not substantially affect HORmon's ability to extract HORs from monomer-graphs, and is mostly needed to ensure reasonable filtering of low multiplicity edges.

- *Comparison of monomer-sets.* The selection of a single parameter *MinSharedLength* (default value 150 bp) does not substantially affect the comparison of the monomer-sets. Typically, a higher (lower) value of *MinSharedLength* results in a stricter (looser) selection of blocks for the set *SharedBlocks.*

#### Supplementary Note 5: Generating the nucleotide consensus of a HOR

Given a pair of consecutive monomers $M_i$ and $M_{i+1}$ in a $t$-monomer (cyclic) HOR $H = M_1 ,..., M_t$, HORmon identifies all pairs formed by the consecutive $M_i$-block and $M_{i+1}$-block in the centromere (we assume that $M_{t+1} = M_1$). It further extracts all nucleotide strings formed by these pairs of monomer-blocks, constructs their multiple alignment, and computes its consensus $C_i$. Ideally, the resulting strings $C_i$ and $C_{i+1}$ should perfectly overlap over the monomer $M_{i+1}$. However, in practice, a short suffix of $C_i$ and a short prefix of $C_{i+1}$ may suffer from a somewhat lower nucleotide accuracy due to artifacts of multiple alignment. To overcome this issue, HORmon constructs an *overlap alignment* of the suffix of $C_i$ and prefix of $C_{i+1}$ using the fast Edlib tool (Šošić and Šikić, 2017). We denote the ending (starting) coordinate in $C_i$ ($C_{i+1}$) of the *longest match* in the constructed alignment as $right_i$ ($left_{i+1}$). HORmon concatenates the prefix of $C_i$ ending at position $right_i$ with the suffix of $C_{i+1}$ starting at position $left_{i+1}$ resulting in a more accurate circular nucleotide consensus of $H$. Since $M$-consensus for each monomer $M$ is constructed as a multiple alignment of all $M$-blocks, its short prefix and suffix can also suffer from a lower nucleotide accuracy. Thus, concatenation of $M_1$-consensus, $M_2$-consensus, …, $M_t$-consensus does not necessarily coincide with the nucleotide consensus of that HOR $H$. HORmon improves the $M$-consensus for each monomer $M$ by launching StringDecomposer on monomers $M_1 ,..., M_t$ and the constructed consensus of $H$. This partitioning ensures that the concatenate of all $M$-consensus coincides with the nucleotide sequence of $H$.

#### Supplementary Note 6: HORmon monomer naming

Below we describe the HORmon rules for monomer naming. Supplementary Table MonomerNaming provides information about the monomer naming generated by HORmon, the classical naming, and naming in Altemose et al., 2021.

- Each monomer from the monomer-set *MonomersNew,* that is not affected by heuristics inspired by the CE Postulate (split/merge transformations or a dehybridization), is named as $\mathbf{SC_1/../C_n}$, where the identifier **S** is typically a letter from A to Z (if there are more than 26 monomers, HORmon names them using 2-letter strings from AA to ZZ), and $\mathbf{C_1/../C_n}$ is the list of chromosomes, where this monomer was found by CentromereArchitect. For example, **A1** is a monomer A found in cen1, **CX** is a monomer C from cenX, and **R1/5/19** is a monomer R found in three centromeres 1, 5, and 19. No two monomers within a single centromere can share the same identifier.

- Hybrid monomers have notation $\mathbf{SC_1/../C_n(M_1/M_2)}$. The notation $\mathbf{SC_1/../C_n}$ is described above, and $M_1$ and $M_2$ are monomers that constitute the hybrid monomer **S**. For example, **NX(K/J)** is a hybrid monomer N from cenX, constructed as a concatenate of a prefix of the monomer KX with a suffix of the monomer JX. In order to describe a hybrid monomer

**$SC_1$/../$C_n$($M_1$/$M_2$)** in more detail, we occasionally represent it as **$M_1C_1$/../$C_n$(*r*)/$M_2C_1$/../$C_k$(*l*)** to show that **S** was constructed as a concatenate of monomers **$M_1C_1$/../$C_n$** and **$M_2C_1$/../$C_k$** using the prefix of **$M_1C_1$/../$C_n$** of length *r* and the suffix of **$M_2C_1$/../$C_k$** of length *l*. Note that Dvorkina et al., 2021 used "+" sign instead of "/" to denote hybrid. We decided to reassign the hybrid operation to "/" and reserve "+" for the merge operation.

- Monomers obtained by a merge operation are represented as a sum of merged monomers, i.e. **$S_1C_1$/../$C_n$ +..+ $S_mC_1$/../$C_n$**. For example, monomer **I1+L1+E1/5/16/19+H1/5/19** is a result of the merge operation of four initial monomers **I1**, **L1**, **E1/5/16/19**, and **H1/5/19**.

Monomers obtained by a split operation of a single monomer **$SC_1$/../$C_n$** are represented as **$SC_1$/../$C_n$.0** and **$SC_1$/../$C_n$.1**. For example, the initial monomer H18 was split into monomers **H18.0** and **H18.1**.

| cen | classical HOR name | Altemose et al., 2021 HOR name | HOR length (bp) | Altemose et al., 2021 monomer name | HORmon monomer name |
|---|---|---|---|---|---|
| 1 | D1Z7 | S1C1/5/19H1L | 1019 | 1 | A1 |
| | | | | 2 | B1/5/19 |
| | | | | 3 | C1/5/19 |
| | | | | 4 | D1/19/5+P1+F1/19+G1/5/19+V1/19.1 |
| | | | | 5 | I1+L1+E1/5/16/19+H1/5/19 |
| | | | | 6 | D1/5/19+P1+F1/19+G1/5/19+V1/19.0 |
| 2 | D2Z1 | S2C2H1L | 680 | 1 | A2 |
| | | | | 2 | B2 |
| | | | | 3 | C2 |
| | | | | 4 | D2/20 |
| 3 | D3Z1 | S01/1C3H1L | 2891 | 1 | A3 |
| | | | | 2 | B3 |
| | | | | 3 | C3 |
| | | | | 4 | D3 |
| | | | | 5 | E3 |
| | | | | 6 | F3 |
| | | | | 7 | G3 |
| | | | | 8 | H3 |
| | | | | 9 | I3 |
| | | | | 10 | J3 |
| | | | | 11 | K3 |
| | | | | 12 | L3 |
| | | | | 13 | M3 |
| | | | | 14 | N3 |
| | | | | 15 | O3 |
| | | | | 16 | P3 |
| | | | | 17 | R3 |
| 4 | D4Z1 | S2C4H1L | 3232 | 1 | A4 |
| | | | | 2 | B4.0 |
| | | | | 3 | C4 |
| | | | | 4 | R4.1 |
| | | | | 5 | E4 |
| | | | | 6 | B4.1 |
| | | | | 7 | G4 |
| | | | | 8 | H4/9 |
| | | | | 9 | I4 |
| | | | | 10 | J4 |
| | | | | 11 | K4 |
| | | | | 12 | L4 |
| | | | | 13 | M4 |
| | | | | 14 | N4 |
| | | | | 15 | O4 |
| | | | | 16 | P4 |
| | | | | 17 | Q4 |
| | | | | 18 | R4.0 |

| | | | | | |
|---|---|---|---|---|---|
| | | | | 19 | S4 |
| 5 | D5Z2 | S1C1/5/19H1L | 1019/1020 | 1 | A1/5/16/19 |
| | | | | 2 | B5 |
| | | | | 3 | C5 |
| | | | | 4 | D5 |
| | | | | 5 | E1/5/16/19 |
| | | | | 6 | F5 |
| 6 | D6Z1 | S01C6H1L | 3057 | 1 | A6 |
| | | | | 2 | B6 |
| | | | | 3 | C6 |
| | | | | 4 | D6 |
| | | | | 5 | E6 |
| | | | | 6 | F6 |
| | | | | 7 | G6 |
| | | | | 8 | H6 |
| | | | | 9 | I6 |
| | | | | 10 | J6 |
| | | | | 11 | K6 |
| | | | | 12 | L6 |
| | | | | 13 | M6 |
| | | | | 14 | N6 |
| | | | | 15 | O6 |
| | | | | 16 | P6 |
| | | | | 17 | R6 |
| 7 | D7Z1 | S1C7H1L | 1022 | 1 | A7 |
| | | | | 2 | B7 |
| | | | | 3 | C7 |
| | | | | 4 | D7 |
| | | | | 5 | E7 |
| | | | | 6 | F7 |
| 8 | D8Z2 | S2C8H1L | 1868 | 1 | A8 |
| | | | | 2 | B8 |
| | | | | 3 | C8 |
| | | | | 4 | D8 |
| | | | | 5 | E8 |
| | | | | 6 | F8 |
| | | | | 7 | G8 |
| | | | | 8 | H8 |
| | | | | 9 | I8 |
| | | | | 10 | J8 |
| | | | | 11 | K8 |
| 9 | D9Z4 | S2C9H1L | 1194/1192 | 1 | A4/9 |
| | | | | 2 | L9+B9+Y9 |
| | | | | 3 | C9 |
| | | | | 4 | D9 |
| | | | | 5 | E9 |
| | | | | 6 | Z4/9 |
| | | | | 7 | G9+M9 |
| 10 | D10Z1 | S1C10H1L | 1357 | 1 | A10 |
| | | | | 2 | B10 |
| | | | | 3 | C10 |
| | | | | 4 | D10 |
| | | | | 5 | E10 |
| | | | | 6 | F10 |
| | | | | 7 | G10 |
| | | | | 8 | H10 |
| 11 | D11Z1 | S3C11H1L | 850 | 1 | A11 |
| | | | | 2 | B11 |
| | | | | 3 | C11 |
| | | | | 4 | D11 |

| | | | | | |
|---|---|---|---|---|---|
| | | | | 5 | E11 |
| 12 | D12Z3 | S1C12H1L | 1359 | 1 | A12 |
| | | | | 2 | B12 |
| | | | | 3 | C12 |
| | | | | 4 | D12 |
| | | | | 5 | E12 |
| | | | | 6 | F12 |
| | | | | 7 | G12 |
| | | | | 8 | H12 |
| 13 | D13Z1 | S2C13/21H1L | 1870 | 1 | A13/21 |
| | | | | 2 | B13/21 |
| | | | | 3 | C13/21 |
| | | | | 4 | D13/21 |
| | | | | 5 | E13/21 |
| | | | | 6 | F13/21 |
| | | | | 7 | G13/21.0 |
| | | | | 8 | H13/21 |
| | | | | 9 | I13/21 |
| | | | | 10 | J13/21 |
| | | | | 11 | G13/21.1 |
| 14 | D14Z9 | S2C14/22H1L | 1364 | 1 | A14/22 |
| | | | | 2 | B14/22 |
| | | | | 3 | C14/22 |
| | | | | 4 | D14/22 |
| | | | | 5 | E14/22 |
| | | | | 6 | F14/22 |
| | | | | 7 | G14/22 |
| | | | | 8 | H14/22 |
| 15 | D15Z3 | S2C15H1L | 1877 | 1 | A15 |
| | | | | 2 | B15 |
| | | | | 3 | C15 |
| | | | | 4 | D15+L15 |
| | | | | 5 | E15 |
| | | | | 6 | F15 |
| | | | | 7 | G15 |
| | | | | 8 | H15 |
| | | | | 9 | I15 |
| | | | | 10 | J15 |
| | | | | 11 | K15 |
| 16 | D16Z2 | S1C16H1L | 1699 | 1 | A1/5/16/19 |
| | | | | 2 | B16 |
| | | | | 3 | C16 |
| | | | | 4 | D1/5/16/19 |
| | | | | 5 | E1/5/16/19 |
| | | | | 6 | F16/19 |
| | | | | 7 | G16 |
| | | | | 8 | H16/19 |
| | | | | 9 | I16 |
| | | | | 10 | J16 |
| 17 | D17Z1 | S3C17H1L | 2715 | 1 | A17+AE17 |
| | | | | 2 | B17 |
| | | | | 3 | C17 |
| | | | | 4 | D17 |
| | | | | 5 | E17 |
| | | | | 6 | F17 |
| | | | | 7 | G17.0 |
| | | | | 8 | H17 |
| | | | | 9 | I17 |
| | | | | 10 | J17 |
| | | | | 11 | K17 |

| | | | | | |
|---|---|---|---|---|---|
| | | | | 12 | L17 |
| | | | | 13 | M17 |
| | | | | 14 | AQ17+N17 |
| | | | | 15 | O17 |
| | | | | 16 | P17 |
| 17 | D17Z1B | S3C17H1-B | 2379 | 1 | Q17 |
| | | | | 2 | R17 |
| | | | | 3 | S17 |
| | | | | 4 | T17 |
| | | | | 5 | U17 |
| | | | | 6 | V17 |
| | | | | 7 | G17.1 |
| | | | | 8 | X17 |
| | | | | 9 | Y17 |
| | | | | 10 | Z17 |
| | | | | 11 | AA17 |
| | | | | 12 | AB17 |
| | | | | 13 | AC17 |
| | | | | 14 | AD17 |
| 18 | D18Z1 | S2C18H1L | 2035 | 1 | A18 |
| | | | | 2 | B18 |
| | | | | 3 | G18.0 |
| | | | | 4 | J18.1 |
| | | | | 5 | E18 |
| | | | | 6 | F18 |
| | | | | 7 | G18.1 |
| | | | | 8 | H18/20 |
| | | | | 9 | I18 |
| | | | | 10 | J18.0 |
| | | | | 11 | K18 |
| | | | | 12 | L18 |
| 19 | D19Z3 | S1C1/5/19H1L | 340 | 5 | AD1/5/16/19+E1/5/19 |
| | | | | 6/4 | G1/5/19+F19+R19 |
| 20 | D20Z2 | S2C20H1L | 2719 | 1 | A20 |
| | | | | 2 | B20 |
| | | | | 3 | C20 |
| | | | | 4 | D20 |
| | | | | 5 | E20 |
| | | | | 6 | F2/20 |
| | | | | 7 | G20 |
| | | | | 8 | H20 |
| | | | | 9 | I20 |
| | | | | 10 | J20 |
| | | | | 11 | K20.1 |
| | | | | 12 | L20 |
| | | | | 13 | K20.0 |
| | | | | 14 | N20 |
| | | | | 15 | O20 |
| | | | | 16 | P20 |
| 21 | D21Z1 | S2C13/21H1L | 1870 | 1 | A13/21 |
| | | | | 2 | B13/21 |
| | | | | 3 | C13/21 |
| | | | | 4 | D13/21 |
| | | | | 5 | E13/21 |
| | | | | 6 | F13/21 |
| | | | | 7 | G13/21.1 |
| | | | | 8 | H13/21 |
| | | | | 9 | I13/21 |
| | | | | 10 | J13/21 |
| | | | | 11 | G13/21.0 |

| | | | | | |
|---|---|---|---|---|---|
| 22 | D22Z1 | S2C14/22H1L | 1364/1365 | 1 | A14/22 |
| | | | | 2 | B14/22 |
| | | | | 3 | C14/22 |
| | | | | 4 | D14/22 |
| | | | | 5 | E14/22 |
| | | | | 6 | F14/22 |
| | | | | 7 | G14/22 |
| | | | | 8 | H14/22 |
| X | DXZ1 | S3CXH1L | 2057 | 1 | AX |
| | | | | 2 | BX |
| | | | | 3 | CX |
| | | | | 4 | DX |
| | | | | 5 | EX |
| | | | | 6 | FX |
| | | | | 7 | GX |
| | | | | 8 | HX |
| | | | | 9 | IX |
| | | | | 10 | JX |
| | | | | 11 | KX |
| | | | | 12 | LX |

**Supplementary Table MonomerNaming. Information about the monomer naming generated by HORmon, the classical naming, and naming in Altemose et al., 2021.** Each row corresponds to a frequent non-hybrid monomer for a specific centromere. The first column corresponds to the centromere. The second column corresponds to the classical HOR naming based on Alexandrov et al., 2001, Shepelev et al., 2015, and McNulty and Sullivan 2018. The third column corresponds to the HOR naming in Altemose et al., 2021. The fourth column shows the length of the nucleotide consensus of HOR. The fifth column corresponds to the monomer numbering with respect to the rules specified in Uralsky et al., 2019 (see also Altemose et al., 2021). The sixth column provides the monomer names generated by HORmon. In centromeres 1, 2, 5, and 15, we report a different number of monomers than McNulty and Sullivan 2018. We hypothesize that these minor differences are due to the absence of a complete genome assembly in prior studies.
