## Supplementary File AllHORDecompositions for "HORmon: automated annotation of human centromeres"

### cen1

$c_5$I1+L1+E1/16/19/5+H1/19/5$p_{4-5}^2$$p_{2-3}$$c_6$I1+L1+E1/16/19/5+H1/19/5$p_{4-5}^2$$p_{2-3}$$p_{1-5}$
I1+L1+E1/16/19/5+H1/19/5$p_{4-5}$I1+L1+E1/16/19/5+H1/19/5$p_{4-5}$
I1+L1+E1/16/19/5+H1/19/5$^2$D1/19/5+P1+F1/19+G1/19/5+V1/19.1I1+L1+E1/16/19/5+H1/19/5$^3$D
1/19/5+P1+F1/19+G1/19/5+V1/19.1I1+L1+E1/16/19/5+H1/19/5$^2$$p_{4-5}^2$
I1+L1+E1/16/19/5+H1/19/5$^2$A1$p_{4-5}$I1+L1+E1/16/19/5+H1/19/5$^3$$p_{4-5}^2$
I1+L1+E1/16/19/5+H1/19/5$^5$$p_{4-5}$I1+L1+E1/16/19/5+H1/19/5$^2$$p_{4-5}$I1+L1+E1/16/19/5+H1/19/5$p_{4-5}^2$
I1+L1+E1/16/19/5+H1/19/5$^3$$p_{4-5}$I1+L1+E1/16/19/5+H1/19/5$^2$$p_{4-5}^2$I1+L1+E1/16/19/5+H1/19/5$^7$$p_{4-5}$
I1+L1+E1/16/19/5+H1/19/5$^2$$p_{4-5}^4$I1+L1+E1/16/19/5+H1/19/5$^2$$p_{4-5}$I1+L1+E1/16/19/5+H1/19/5$p_{4-5}^2$
I1+L1+E1/16/19/5+H1/19/5$^2$D1/19/5+P1+F1/19+G1/19/5+V1/19.1I1+L1+E1/16/19/5+H1/19/5$^2$
$c_5$I1+L1+E1/16/19/5+H1/19/5$c_4$$p_{4-5}^2$
I1+L1+E1/16/19/5+H1/19/5D1/19/5+P1+F1/19+G1/19/5+V1/19.1$p_{4-6}$$p_{2-5}$
I1+L1+E1/16/19/5+H1/19/5$p_{2-5}$$p_{4-5}$$c_4$$p_{4-5}^3$I1+L1+E1/16/19/5+H1/19/5$^2$$p_{4-6}$$p_{2-5}$
I1+L1+E1/16/19/5+H1/19/5$p_{3-5}$$p_{4-5}$$c_4$$p_{4-5}^3$I1+L1+E1/16/19/5+H1/19/5$^3$$p_{1-5}$
A1$c_5^2$I1+L1+E1/16/19/5+H1/19/5$p_{4-5}$$c_4$$p_{4-5}$I1+L1+E1/16/19/5+H1/19/5$^2$$p_{4-5}^2$$c_4$$p_{4-5}$
D1/19/5+P1+F1/19+G1/19/5+V1/19.1$p_{6-3}$$p_{6-1}$$c_6$$p_{4-1}$$p_{6-2}$$p_{4-5}$I1+L1+E1/16/19/5+H1/19/5$^2$$c_4$$p_{4-5}$
$c_5$I1+L1+E1/16/19/5+H1/19/5$^2$$p_{4-5}$I1+L1+E1/16/19/5+H1/19/5$c_4$$p_{4-5}$
$c_5$I1+L1+E1/16/19/5+H1/19/5$^2$$p_{4-5}$$p_{2-5}$D1/19/5+P1+F1/19+G1/19/5+V1/19.1$p_{5-3}$
D1/19/5+P1+F1/19+G1/19/5+V1/19.0I1+L1+E1/16/19/5+H1/19/5$p_{4-6}$$c_5$$p_{5-3}$
I1+L1+E1/16/19/5+H1/19/5D1/19/5+P1+F1/19+G1/19/5+V1/19.1$c_5$$p_{5-3}$
D1/19/5+P1+F1/19+G1/19/5+V1/19.0I1+L1+E1/16/19/5+H1/19/5D1/19/5+P1+F1/19+G1/19/5+
V1/19.1$c_5$$p_{5-3}$I1+L1+E1/16/19/5+H1/19/5D1/19/5+P1+F1/19+G1/19/5+V1/19.1$c_5$$p_{5-3}$
D1/19/5+P1+F1/19+G1/19/5+V1/19.0I1+L1+E1/16/19/5+H1/19/5D1/19/5+P1+F1/19+G1/19/5+
V1/19.1$c_5$$p_{5-3}$D1/19/5+P1+F1/19+G1/19/5+V1/19.0I1+L1+E1/16/19/5+H1/19/5$p_{4-5}$$p_{4-6}$$c_5$$p_{5-3}$
D1/19/5+P1+F1/19+G1/19/5+V1/19.0I1+L1+E1/16/19/5+H1/19/5$p_{4-5}$$p_{4-6}$$c_5$$p_{5-3}$
D1/19/5+P1+F1/19+G1/19/5+V1/19.0I1+L1+E1/16/19/5+H1/19/5$p_{4-5}$$p_{4-6}$$p_{5-3}$$p_{6-1}$$p_{6-2}$$p_{4-5}$
I1+L1+E1/16/19/5+H1/19/5$p_{4-5}$I1+L1+E1/16/19/5+H1/19/5$c_4$$p_{4-5}$I1+L1+E1/16/19/5+H1/19/5$p_{4-5}$
$p_{2-5}$$c_4$D1/19/5+P1+F1/19+G1/19/5+V1/19.1$p_{5-3}$
D1/19/5+P1+F1/19+G1/19/5+V1/19.0I1+L1+E1/16/19/5+H1/19/5$p_{4-6}$$c_5$$p_{5-3}$
D1/19/5+P1+F1/19+G1/19/5+V1/19.0I1+L1+E1/16/19/5+H1/19/5$p_{4-5}$$p_{4-6}$
$c_5$I1+L1+E1/16/19/5+H1/19/5$p_{5-6}$
$c_5$$c_4$D1/19/5+P1+F1/19+G1/19/5+V1/19.0I1+L1+E1/16/19/5+H1/19/5$p_{4-5}$$p_{5-6}$
$c_5$$c_4$D1/19/5+P1+F1/19+G1/19/5+V1/19.0I1+L1+E1/16/19/5+H1/19/5$p_{4-5}$$p_{4-6}$
$c_5$$c_4$D1/19/5+P1+F1/19+G1/19/5+V1/19.0I1+L1+E1/16/19/5+H1/19/5$p_{4-5}$$p_{4-6}$
$c_5$I1+L1+E1/16/19/5+H1/19/5$p_{4-1}$I1+L1+E1/16/19/5+H1/19/5$^3$$p_{4-6}$I1+L1+E1/16/19/5+H1/19/5$^2$$p_{4-5}^3$
$p_{4-1}$$c_6$I1+L1+E1/16/19/5+H1/19/5$p_{4-5}^3$$c_4$$p_{4-5}$I1+L1+E1/16/19/5+H1/19/5$p_{4-5}$$p_{4-2}$
I1+L1+E1/16/19/5+H1/19/5$c_4$$p_{4-5}^2$$c_4$$p_{4-5}^4$I1+L1+E1/16/19/5+H1/19/5$p_{4-5}$
D1/19/5+P1+F1/19+G1/19/5+V1/19.1$c_6$$p_{4-5}^2$$c_4$$p_{4-5}$I1+L1+E1/16/19/5+H1/19/5$^2$$c_4$$p_{4-5}$
I1+L1+E1/16/19/5+H1/19/5$p_{4-5}^3$$c_4$$p_{4-5}$
I1+L1+E1/16/19/5+H1/19/5$c_4$I1+L1+E1/16/19/5+H1/19/5$p_{4-5}^3$$c_4$$p_{4-5}$$c_4$$p_{4-5}$
I1+L1+E1/16/19/5+H1/19/5$c_4$$p_{4-5}$
D1/19/5+P1+F1/19+G1/19/5+V1/19.1$c_6$I1+L1+E1/16/19/5+H1/19/5$p_{4-5}$$c_4$$p_{4-5}^2$$c_4$$p_{4-5}$
I1+L1+E1/16/19/5+H1/19/5$p_{4-5}$$c_4$$p_{4-5}^2$$c_4$$p_{4-5}^2$D1/19/5+P1+F1/19+G1/19/5+V1/19.1$p_{1-4}$$p_{1-5}$$p_{4-5}^2$$p_{4-1}$
$p_{1-5}$$p_{4-6}$I1+L1+E1/16/19/5+H1/19/5$p_{4-5}$$c_4$D1/19/5+P1+F1/19+G1/19/5+V1/19.1$p_{1-5}$$p_{4-5}^2$$p_{4-1}$$p_{6-1}$$c_6$$p_{4-1}$
$c_6$$p_{4-5}^2$$p_{4-1}$$p_{6-1}$$c_6$$p_{4-1}$$c_6^2$$p_{4-5}$I1+L1+E1/16/19/5+H1/19/5$c_4$$p_{4-5}^2$$c_6$$p_{4-5}^2$$p_{1-5}$$p_{4-5}^3$
I1+L1+E1/16/19/5+H1/19/5$^3$$p_{4-5}$$c_5$I1+L1+E1/16/19/5+H1/19/5$^2$$p_{4-5}^2$$c_4$$p_{4-5}$
I1+L1+E1/16/19/5+H1/19/5$p_{4-5}^3$D1/19/5+P1+F1/19+G1/19/5+V1/19.1$p_{3-5}$
I1+L1+E1/16/19/5+H1/19/5$c_4$$p_{4-5}$I1+L1+E1/16/19/5+H1/19/5$p_{4-5}^2$$c_4$$p_{4-5}$
I1+L1+E1/16/19/5+H1/19/5$p_{4-5}$$p_{4-}$

$_6$I1+L1+E1/16/19/5+H1/19/5D1/19/5+P1+F1/19+G1/19/5+V1/19.1p$_{3-}$
$_5$I1+L1+E1/16/19/5+H1/19/5c$_4$p$_{4-5}$p$_{5-1}$c$_6$p$_{4-5}^3$D1/19/5+P1+F1/19+G1/19/5+V1/19.1p$_{3-}$
$_5$I1+L1+E1/16/19/5+H1/19/5c$_1$p$_{1-5}$p$_{4-5}^2$I1+L1+E1/16/19/5+H1/19/5p$_{4-}$
$_5^2$I1+L1+E1/16/19/5+H1/19/5$^2$c$_5$I1+L1+E1/16/19/5+H1/19/5p$_{4-5}^2$p$_{1-5}$p$_{4-}$
$_5^4$D1/19/5+P1+F1/19+G1/19/5+V1/19.1I1+L1+E1/16/19/5+H1/19/5p$_{4-5}$c$_4^3$p$_{4-}$
$_5^3$c$_5$I1+L1+E1/16/19/5+H1/19/5c$_4$p$_{4-5}^3$p$_{1-5}$p$_{4-}$
$_5^4$D1/19/5+P1+F1/19+G1/19/5+V1/19.1I1+L1+E1/16/19/5+H1/19/5p$_{4-5}$c$_4^4$p$_{4-}$
$_5^2$I1+L1+E1/16/19/5+H1/19/5c$_4^2$p$_{4-5}^2$c$_4$p$_{4-5}$I1+L1+E1/16/19/5+H1/19/5$^3$p$_{4-}$
$_5$I1+L1+E1/16/19/5+H1/19/5p$_{4-5}$c$_4$p$_{4-5}$c$_4$p$_{4-5}$I1+L1+E1/16/19/5+H1/19/5p$_{4-}$
$_5$I1+L1+E1/16/19/5+H1/19/5D1/19/5+P1+F1/19+G1/19/5+V1/19.1p$_{3-5}$p$_{5-1}$p$_{6-4}$p$_{4-}$
$_5$I1+L1+E1/16/19/5+H1/19/5c$_4$D1/19/5+P1+F1/19+G1/19/5+V1/19.1p$_{4-}$
$_5$I1+L1+E1/16/19/5+H1/19/5c$_4$p$_{4-5}$c$_4$p$_{4-5}^2$I1+L1+E1/16/19/5+H1/19/5c$_4$p$_{4-}$
$_6$I1+L1+E1/16/19/5+H1/19/5p$_{4-5}$c$_4^2$p$_{4-5}$c$_4$D1/19/5+P1+F1/19+G1/19/5+V1/19.1p$_{4-}$
$_5^2$c$_4$D1/19/5+P1+F1/19+G1/19/5+V1/19.1p$_{4-}$
$_5^2$D1/19/5+P1+F1/19+G1/19/5+V1/19.1c$_6$I1+L1+E1/16/19/5+H1/19/5c$_4^2$p$_{4-}$
$_5$c$_5$I1+L1+E1/16/19/5+H1/19/5$^2$c$_4$p$_{4-5}$c$_4$p$_{4-5}^3$c$_4^2$p$_{4-5}$c$_5$I1+L1+E1/16/19/5+H1/19/5$^2$c$_4$p$_{4-}$
$_5$D1/19/5+P1+F1/19+G1/19/5+V1/19.1c$_6$p$_{4-5}^2$c$_4^2$p$_{4-5}$c$_5$I1+L1+E1/16/19/5+H1/19/5p$_{4-5}$c$_4$p$_{4-5}^2$c$_4$p$_{4-}$
$_5$c$_4$p$_{4-5}^3$c$_4$p$_{4-5}$I1+L1+E1/16/19/5+H1/19/5c$_4$p$_{4-6}$I1+L1+E1/16/19/5+H1/19/5p$_{4-}$
$_5^2$c$_5^2$I1+L1+E1/16/19/5+H1/19/5p$_{4-5}^2$c$_5$I1+L1+E1/16/19/5+H1/19/5p$_{4-}$
$_5$I1+L1+E1/16/19/5+H1/19/5c$_4$p$_{4-5}$D1/19/5+P1+F1/19+G1/19/5+V1/19.1p$_{4-5}$c$_4$p$_{4-}$
$_5^2$c$_5$I1+L1+E1/16/19/5+H1/19/5c$_4$p$_{4-5}$c$_5$I1+L1+E1/16/19/5+H1/19/5$^2$c$_4$p$_{4-}$
$_5$D1/19/5+P1+F1/19+G1/19/5+V1/19.1p$_{4-5}$c$_4$p$_{4-5}^2$c$_5$I1+L1+E1/16/19/5+H1/19/5c$_4$p$_{4-}$
$_5$c$_5$I1+L1+E1/16/19/5+H1/19/5$^2$p$_{4-5}^3$c$_5^2$I1+L1+E1/16/19/5+H1/19/5p$_{4-5}$c$_4$p$_{4-5}$p$_{4-1}$c$_6$c$_4$p$_{4-}$
$_5$I1+L1+E1/16/19/5+H1/19/5c$_4$p$_{4-5}$D1/19/5+P1+F1/19+G1/19/5+V1/19.1p$_{4-5}$c$_4$p$_{4-}$
$_5^2$c$_5$I1+L1+E1/16/19/5+H1/19/5c$_4$p$_{4-5}$c$_5$I1+L1+E1/16/19/5+H1/19/5$^2$p$_{4-}$
$_6$I1+L1+E1/16/19/5+H1/19/5p$_{4-5}$c$_5$I1+L1+E1/16/19/5+H1/19/5p$_{4-}$
$_5$c$_5$I1+L1+E1/16/19/5+H1/19/5c$_6$p$_{4-5}$c$_4$p$_{4-5}^2$c$_4$p$_{4-}$
$_5$D1/19/5+P1+F1/19+G1/19/5+V1/19.1c$_6$D1/19/5+P1+F1/19+G1/19/5+V1/19.1D1/19/5+P1+F1/1
9+G1/19/5+V1/19.0I1+L1+E1/16/19/5+H1/19/5c$_4$p$_{4-5}$c$_4$p$_{4-}$
$_5^3$D1/19/5+P1+F1/19+G1/19/5+V1/19.1p$_{3-5}$c$_4$p$_{4-5}^3$D1/19/5+P1+F1/19+G1/19/5+V1/19.1p$_{3-}$
$_5$I1+L1+E1/16/19/5+H1/19/5D1/19/5+P1+F1/19+G1/19/5+V1/19.1p$_{3-5}$p$_{4-}$
$_5^3$I1+L1+E1/16/19/5+H1/19/5D1/19/5+P1+F1/19+G1/19/5+V1/19.1c$_3$p$_{3-4}$p$_{4-}$
$_5$D1/19/5+P1+F1/19+G1/19/5+V1/19.1p$_{6-4}$D1/19/5+P1+F1/19+G1/19/5+V1/19.1p$_{6-4}$p$_{4-5}$c$_5$p$_{4-}$
$_5$I1+L1+E1/16/19/5+H1/19/5D1/19/5+P1+F1/19+G1/19/5+V1/19.1p$_{3-}$
$_5$I1+L1+E1/16/19/5+H1/19/5p$_{4-5}$c$_5$p$_{4-}$
$_5$I1+L1+E1/16/19/5+H1/19/5D1/19/5+P1+F1/19+G1/19/5+V1/19.1p$_{3-}$
$_5$I1+L1+E1/16/19/5+H1/19/5p$_{4-5}$I1+L1+E1/16/19/5+H1/19/5p$_{4-}$
$_5$I1+L1+E1/16/19/5+H1/19/5D1/19/5+P1+F1/19+G1/19/5+V1/19.1p$_{3-5}$p$_{4-}$
$_5^3$I1+L1+E1/16/19/5+H1/19/5D1/19/5+P1+F1/19+G1/19/5+V1/19.1c$_3$p$_{3-4}$p$_{4-}$
$_5$D1/19/5+P1+F1/19+G1/19/5+V1/19.1p$_{6-4}$p$_{4-5}$c$_5$I1+L1+E1/16/19/5+H1/19/5p$_{4-}$
$_5$c$_4$D1/19/5+P1+F1/19+G1/19/5+V1/19.1p$_{4-5}$I1+L1+E1/16/19/5+H1/19/5p$_{4-5}$c$_4$p$_{4-5}$c$_6$p$_{4-}$
$_5$I1+L1+E1/16/19/5+H1/19/5p$_{4-5}^2$c$_4$p$_{4-5}$c$_6$p$_{4-5}$I1+L1+E1/16/19/5+H1/19/5p$_{4-}$
$_5$D1/19/5+P1+F1/19+G1/19/5+V1/19.1p$_{3-5}$I1+L1+E1/16/19/5+H1/19/5p$_{4-}$
$_5$D1/19/5+P1+F1/19+G1/19/5+V1/19.1p$_{3-}$
$_5$I1+L1+E1/16/19/5+H1/19/5$^2$D1/19/5+P1+F1/19+G1/19/5+V1/19.1p$_{3-}$
$_5$I1+L1+E1/16/19/5+H1/19/5p$_{4-5}$I1+L1+E1/16/19/5+H1/19/5p$_{4-}$
$_5$I1+L1+E1/16/19/5+H1/19/5D1/19/5+P1+F1/19+G1/19/5+V1/19.1p$_{3-5}$p$_{4-}$
$_5^3$D1/19/5+P1+F1/19+G1/19/5+V1/19.1c$_3$p$_{3-4}$p$_{4-5}$D1/19/5+P1+F1/19+G1/19/5+V1/19.1p$_{6-4}$p$_{4-}$
$_5$c$_5$I1+L1+E1/16/19/5+H1/19/5p$_{4-5}$I1+L1+E1/16/19/5+H1/19/5p$_{4-5}$c$_4$p$_{4-}$
$_5^2$I1+L1+E1/16/19/5+H1/19/5p$_{4-}$
$_5$c$_4$I1+L1+E1/16/19/5+H1/19/5$^2$D1/19/5+P1+F1/19+G1/19/5+V1/19.1p$_{3-4}$p$_{4-}$

$_{5}$I1+L1+E1/16/19/5+H1/19/5$c_4p_{4-5}$D1/19/5+P1+F1/19+G1/19/5+V1/19.1$c_6{}^2p_{4-}$
$_{2}$I1+L1+E1/16/19/5+H1/19/5$p_{4-5}c_4p_{4-5}{}^2$I1+L1+E1/16/19/5+H1/19/5$p_{4-5}c_4p_{4-5}c_6p_{4-}$
$_{5}$I1+L1+E1/16/19/5+H1/19/5$p_{4-5}c_4$I1+L1+E1/16/19/5+H1/19/5$p_{4-}$
$_{5}$D1/19/5+P1+F1/19+G1/19/5+V1/19.1$c_6p_{4-5}$I1+L1+E1/16/19/5+H1/19/5$p_{4-}$
$_{5}c_4$I1+L1+E1/16/19/5+H1/19/5$p_{4-5}c_4p_{4-5}c_4p_{4-5}c_4p_{4-5}p_{3-5}p_{4-2}$I1+L1+E1/16/19/5+H1/19/5$p_{4-}$
$_{5}{}^2$I1+L1+E1/16/19/5+H1/19/5$p_{4-5}c_4$I1+L1+E1/16/19/5+H1/19/5$^2c_4$I1+L1+E1/16/19/5+H1/19/5$p_{4-}$
$_{5}{}^4c_4p_{4-5}p_{3-5}p_{4-2}$I1+L1+E1/16/19/5+H1/19/5$p_{4-5}{}^2$I1+L1+E1/16/19/5+H1/19/5$p_{4-}$
$_{5}c_4$I1+L1+E1/16/19/5+H1/19/5$^2c_4p_{4-5}p_{3-5}p_{4-2}$I1+L1+E1/16/19/5+H1/19/5$p_{4-5}{}^2p_{4-}$
$_{6}$I1+L1+E1/16/19/5+H1/19/5$^2c_4$I1+L1+E1/16/19/5+H1/19/5$p_{4-5}{}^5c_4p_{4-5}p_{3-5}p_{4-}$
$_{2}$I1+L1+E1/16/19/5+H1/19/5$p_{4-5}{}^2$I1+L1+E1/16/19/5+H1/19/5$p_{4-5}c_4$I1+L1+E1/16/19/5+H1/19/5$p_{4-}$
$_{5}c_4p_{4-5}c_4p_{4-5}p_{3-5}p_{4-2}$I1+L1+E1/16/19/5+H1/19/5$p_{4-}$
$_{5}$I1+L1+E1/16/19/5+H1/19/5$c_4$I1+L1+E1/16/19/5+H1/19/5$p_{4-5}{}^5c_4p_{4-5}p_{3-5}p_{4-}$
$_{2}$I1+L1+E1/16/19/5+H1/19/5$p_{4-5}{}^2$I1+L1+E1/16/19/5+H1/19/5$p_{4-5}c_4$I1+L1+E1/16/19/5+H1/19/5$p_{4-}$
$_{5}c_4p_{4-5}c_4p_{4-5}{}^2c_4p_{4-5}$I1+L1+E1/16/19/5+H1/19/5$p_{4-5}c_4p_{4-5}p_{4-2}$I1+L1+E1/16/19/5+H1/19/5$c_4p_{4-}$
$_{5}{}^2$I1+L1+E1/16/19/5+H1/19/5$p_{4-5}$I1+L1+E1/16/19/5+H1/19/5$p_{4-5}c_4p_{4-5}{}^6c_4{}^2p_{4-}$
$_{5}$I1+L1+E1/16/19/5+H1/19/5$^2c_4{}^2p_{4-5}$D1/19/5+P1+F1/19+G1/19/5+V1/19.1$c_6{}^2p_{4-}$
$_{5}{}^2c_4$I1+L1+E1/16/19/5+H1/19/5$p_{4-5}{}^2$I1+L1+E1/16/19/5+H1/19/5$p_{4-5}{}^2c_4p_{4-}$
$_{5}$I1+L1+E1/16/19/5+H1/19/5$p_{4-5}{}^2c_4p_{4-}$
$_{5}$I1+L1+E1/16/19/5+H1/19/5$c_4$I1+L1+E1/16/19/5+H1/19/5$p_{4-5}{}^4c_4p_{4-}$
$_{5}$I1+L1+E1/16/19/5+H1/19/5$p_{4-5}{}^2c_4p_{4-}$
$_{5}$I1+L1+E1/16/19/5+H1/19/5$c_4$I1+L1+E1/16/19/5+H1/19/5$p_{4-5}{}^4c_4p_{4-}$
$_{5}$I1+L1+E1/16/19/5+H1/19/5$p_{4-5}{}^3$I1+L1+E1/16/19/5+H1/19/5$p_{4-5}{}^3$I1+L1+E1/16/19/5+H1/19/5$p_{4-}$
$_{5}{}^2$I1+L1+E1/16/19/5+H1/19/5$p_{4-5}c_4p_{4-5}$I1+L1+E1/16/19/5+H1/19/5$^2p_{4-5}c_4p_{4-}$
$_{5}$I1+L1+E1/16/19/5+H1/19/5$^2p_{4-5}c_4p_{4-5}{}^3p_{4-1}$I1+L1+E1/16/19/5+H1/19/5$^4p_{4-5}c_4p_{4-5}{}^2c_4p_{4-}$
$_{5}$I1+L1+E1/16/19/5+H1/19/5$p_{4-5}{}^2c_4p_{4-5}$D1/19/5+P1+F1/19+G1/19/5+V1/19.1$c_4p_{4-}$
$_{5}$I1+L1+E1/16/19/5+H1/19/5$p_{4-5}{}^3p_{4-1}$I1+L1+E1/16/19/5+H1/19/5$^2p_{4-5}{}^3p_{4-}$
$_{1}$I1+L1+E1/16/19/5+H1/19/5$^2p_{4-}$
$_{5}{}^4$D1/19/5+P1+F1/19+G1/19/5+V1/19.1$c_6$D1/19/5+P1+F1/19+G1/19/5+V1/19.0I1+L1+E1/16/19/
5+H1/19/5$p_{4-5}{}^4p_{4-6}$I1+L1+E1/16/19/5+H1/19/5$p_{4-5}{}^3p_{4-6}$I1+L1+E1/16/19/5+H1/19/5$p_{4-5}{}^3c_4p_{4-5}c_4p_{4-}$
$_{5}$I1+L1+E1/16/19/5+H1/19/5$p_{4-5}{}^2$I1+L1+E1/16/19/5+H1/19/5$p_{4-5}{}^3c_4p_{4-}$
$_{5}{}^2$I1+L1+E1/16/19/5+H1/19/5$^2$D1/19/5+P1+F1/19+G1/19/5+V1/19.1$c_6$I1+L1+E1/16/19/5+H1/19/
5$p_{4-5}p_{5-3}$I1+L1+E1/16/19/5+H1/19/5$p_{4-5}{}^4p_{4-6}$I1+L1+E1/16/19/5+H1/19/5$p_{4-5}{}^4p_{4-}$
$_{6}$I1+L1+E1/16/19/5+H1/19/5$p_{4-5}p_{4-6}$I1+L1+E1/16/19/5+H1/19/5$p_{4-5}{}^3c_4p_{4-5}c_4p_{4-}$
$_{5}$I1+L1+E1/16/19/5+H1/19/5$p_{4-5}{}^9c_4p_{4-}$
$_{5}$D1/19/5+P1+F1/19+G1/19/5+V1/19.1I1+L1+E1/16/19/5+H1/19/5$p_{4-5}{}^3c_4p_{4-}$
$_{5}{}^2$I1+L1+E1/16/19/5+H1/19/5$c_4p_{4-5}$I1+L1+E1/16/19/5+H1/19/5$p_{4-5}c_4p_{4-5}{}^3p_{5-}$
$_{3}$I1+L1+E1/16/19/5+H1/19/5$p_{4-5}{}^3c_4c_5$I1+L1+E1/16/19/5+H1/19/5$^2p_{4-5}{}^3c_4{}^2p_{4-}$
$_{5}$I1+L1+E1/16/19/5+H1/19/5$p_{4-5}{}^2c_4p_{4-5}$I1+L1+E1/16/19/5+H1/19/5$p_{4-}$
$_{5}{}^3$I1+L1+E1/16/19/5+H1/19/5$p_{4-5}{}^2c_4p_{4-5}{}^3$I1+L1+E1/16/19/5+H1/19/5$c_4p_{4-}$
$_{5}$I1+L1+E1/16/19/5+H1/19/5$p_{4-5}c_4p_{4-5}$I1+L1+E1/16/19/5+H1/19/5$p_{4-5}{}^5c_4p_{4-5}{}^8c_4p_{4-5}{}^4c_4p_{4-}$
$_{5}{}^3$I1+L1+E1/16/19/5+H1/19/5$c_4{}^2p_{4-5}$I1+L1+E1/16/19/5+H1/19/5$p_{4-5}{}^5c_4{}^2p_{4-5}c_4p_{4-5}{}^5c_4p_{4-}$
$_{5}$I1+L1+E1/16/19/5+H1/19/5$p_{4-5}{}^2$I1+L1+E1/16/19/5+H1/19/5$p_{4-5}c_4p_{4-}$
$_{5}{}^2$I1+L1+E1/16/19/5+H1/19/5$^2p_{4-5}c_4p_{4-5}$I1+L1+E1/16/19/5+H1/19/5$p_{4-}$
$_{5}{}^2$I1+L1+E1/16/19/5+H1/19/5$p_{4-5}c_4p_{4-5}{}^2$I1+L1+E1/16/19/5+H1/19/5$^2p_{4-5}c_4{}^2p_{4-}$
$_{5}$I1+L1+E1/16/19/5+H1/19/5$p_{4-5}{}^2c_4p_{4-5}$I1+L1+E1/16/19/5+H1/19/5$p_{4-5}{}^2c_4p_{4-5}c_4p_{4-}$
$_{5}$I1+L1+E1/16/19/5+H1/19/5$p_{4-5}c_4p_{4-5}$I1+L1+E1/16/19/5+H1/19/5$p_{4-5}c_4p_{4-}$
$_{5}$I1+L1+E1/16/19/5+H1/19/5$p_{4-5}{}^3c_4p_{4-5}c_4p_{4-5}{}^3c_4p_{4-5}$I1+L1+E1/16/19/5+H1/19/5$p_{4-}$
$_{5}{}^2$I1+L1+E1/16/19/5+H1/19/5$p_{4-5}c_4p_{4-5}{}^2$I1+L1+E1/16/19/5+H1/19/5$^2p_{4-5}c_4{}^2p_{4-}$
$_{5}$I1+L1+E1/16/19/5+H1/19/5$p_{4-5}{}^2c_4p_{4-5}$I1+L1+E1/16/19/5+H1/19/5$p_{4-5}{}^2c_4p_{4-5}c_4p_{4-}$
$_{5}$I1+L1+E1/16/19/5+H1/19/5$p_{4-5}c_4p_{4-5}$I1+L1+E1/16/19/5+H1/19/5$p_{4-5}c_4p_{4-}$
$_{5}$I1+L1+E1/16/19/5+H1/19/5$p_{4-5}{}^3c_4p_{4-5}c_4p_{4-5}{}^3$I1+L1+E1/16/19/5+H1/19/5$^2c_4p_{4-}$

5I1+L1+E1/16/19/5+H1/19/5^2p4-5D1/19/5+P1+F1/19+G1/19/5+V1/19.1c4p4-5^3c4p4-
5^2c4D1/19/5+P1+F1/19+G1/19/5+V1/19.1c6c5I1+L1+E1/16/19/5+H1/19/5p4-
5^3c4D1/19/5+P1+F1/19+G1/19/5+V1/19.1c6I1+L1+E1/16/19/5+H1/19/5p4-5^3c4p4-5^3c4p4-
5I1+L1+E1/16/19/5+H1/19/5p4-5^3c4p4-5I1+L1+E1/16/19/5+H1/19/5p4-5^3c4p4-5^3c4p4-5^2c4p4-5^3c4p4-
5I1+L1+E1/16/19/5+H1/19/5p4-5^3c4p4-5c4p4-5^2c4p4-5^4c4D1/19/5+P1+F1/19+G1/19/5+V1/19.1c6c4p4-
5I1+L1+E1/16/19/5+H1/19/5p4-5c4p4-5c4^2p4-5I1+L1+E1/16/19/5+H1/19/5p4-5^3c4p4-
5I1+L1+E1/16/19/5+H1/19/5p4-5c4^2p4-5I1+L1+E1/16/19/5+H1/19/5p4-5^3c4p4-
5I1+L1+E1/16/19/5+H1/19/5p4-5^3c4^2p4-5I1+L1+E1/16/19/5+H1/19/5p4-5c4p4-
5I1+L1+E1/16/19/5+H1/19/5p4-5c4p4-5^4c4p4-5^6c4p4-5I1+L1+E1/16/19/5+H1/19/5p4-5c4p4-
5I1+L1+E1/16/19/5+H1/19/5p4-5c4p4-5^4c4p4-5^6c4p4-5c4p4-5c4p4-5^2c4p4-
5^4c4D1/19/5+P1+F1/19+G1/19/5+V1/19.1c6c4p4-5I1+L1+E1/16/19/5+H1/19/5p4-5c4p4-5c4^2p4-
5I1+L1+E1/16/19/5+H1/19/5p4-5^3c4p4-5I1+L1+E1/16/19/5+H1/19/5p4-5c4^2p4-
5I1+L1+E1/16/19/5+H1/19/5p4-5^3c4p4-5I1+L1+E1/16/19/5+H1/19/5p4-5^3c4^2p4-
5I1+L1+E1/16/19/5+H1/19/5p4-5c4p4-5I1+L1+E1/16/19/5+H1/19/5p4-5c4p4-5^4c4p4-5^6c4p4-
5I1+L1+E1/16/19/5+H1/19/5p4-5c4p4-5I1+L1+E1/16/19/5+H1/19/5p4-5c4p4-5^4c4p4-5^6c4p4-
5I1+L1+E1/16/19/5+H1/19/5p4-5^2c4^2p4-5I1+L1+E1/16/19/5+H1/19/5p4-5^4c4p4-
5I1+L1+E1/16/19/5+H1/19/5p4-5I1+L1+E1/16/19/5+H1/19/5p4-
5^3B1/19/5I1+L1+E1/16/19/5+H1/19/5c4p4-5^2c4^2p4-5I1+L1+E1/16/19/5+H1/19/5p4-5^6c4^2p4-
5I1+L1+E1/16/19/5+H1/19/5p4-5^4c4p4-5I1+L1+E1/16/19/5+H1/19/5p4-
5I1+L1+E1/16/19/5+H1/19/5p4-5^3p2-5p4-5^2c4p4-5^4c4p4-5I1+L1+E1/16/19/5+H1/19/5p4-
5I1+L1+E1/16/19/5+H1/19/5p4-5^3p2-5p4-5^2c4p4-5^4c4p4-5I1+L1+E1/16/19/5+H1/19/5p4-
5I1+L1+E1/16/19/5+H1/19/5p4-5I1+L1+E1/16/19/5+H1/19/5p4-5c4p4-
5I1+L1+E1/16/19/5+H1/19/5p4-5I1+L1+E1/16/19/5+H1/19/5p4-5I1+L1+E1/16/19/5+H1/19/5p4-
5I1+L1+E1/16/19/5+H1/19/5p4-5^3p2-5p4-5^3c4I1+L1+E1/16/19/5+H1/19/5p4-5c4p4-
5I1+L1+E1/16/19/5+H1/19/5c4p4-5^2c4p4-5I1+L1+E1/16/19/5+H1/19/5p4-5c4p4-
5I1+L1+E1/16/19/5+H1/19/5c4p4-5^2c4p4-5I1+L1+E1/16/19/5+H1/19/5p4-5c4p4-
5I1+L1+E1/16/19/5+H1/19/5p4-5c4p4-5I1+L1+E1/16/19/5+H1/19/5c4p4-5^2c4p4-
5I1+L1+E1/16/19/5+H1/19/5p4-5c4p4-5I1+L1+E1/16/19/5+H1/19/5p4-
5c4D1/19/5+P1+F1/19+G1/19/5+V1/19.1I1+L1+E1/16/19/5+H1/19/5p4-5c4p4-
5I1+L1+E1/16/19/5+H1/19/5p4-5c4p4-5^2c4p4-5I1+L1+E1/16/19/5+H1/19/5p4-5c4p4-
5I1+L1+E1/16/19/5+H1/19/5c4D1/19/5+P1+F1/19+G1/19/5+V1/19.1I1+L1+E1/16/19/5+H1/19/5p
4-5c4p4-5I1+L1+E1/16/19/5+H1/19/5p4-
5c4D1/19/5+P1+F1/19+G1/19/5+V1/19.1I1+L1+E1/16/19/5+H1/19/5p4-5c4p4-
5I1+L1+E1/16/19/5+H1/19/5p4-5c4I1+L1+E1/16/19/5+H1/19/5p4-5I1+L1+E1/16/19/5+H1/19/5p4-
5c4p4-
5I1+L1+E1/16/19/5+H1/19/5D1/19/5+P1+F1/19+G1/19/5+V1/19.1D1/19/5+P1+F1/19+G1/19/5+
V1/19.0p2-5I1+L1+E1/16/19/5+H1/19/5p4-5^3c4p4-5I1+L1+E1/16/19/5+H1/19/5p4-5^3c4^2p4-
5I1+L1+E1/16/19/5+H1/19/5p4-5c4p4-5I1+L1+E1/16/19/5+H1/19/5p4-5c4p4-5^3c4p4-5^4c4p4-5^6c4p4-
5^4c4^2p4-5I1+L1+E1/16/19/5+H1/19/5p4-5^4c4p4-5I1+L1+E1/16/19/5+H1/19/5p4-
5I1+L1+E1/16/19/5+H1/19/5p4-5^3B1/19/5I1+L1+E1/16/19/5+H1/19/5c4p4-5^4c4^2p4-
5I1+L1+E1/16/19/5+H1/19/5p4-5^4B1/19/5p4-5^3c4p4-5^4c4p4-5I1+L1+E1/16/19/5+H1/19/5p4-
5I1+L1+E1/16/19/5+H1/19/5p4-5^3p2-5p4-5^2c4p4-5^4c4p4-5I1+L1+E1/16/19/5+H1/19/5p4-
5I1+L1+E1/16/19/5+H1/19/5p4-5I1+L1+E1/16/19/5+H1/19/5p4-5I1+L1+E1/16/19/5+H1/19/5p4-5^3p2-
5p4-5^3c4I1+L1+E1/16/19/5+H1/19/5p4-5c4p4-
5I1+L1+E1/16/19/5+H1/19/5c4c6^2I1+L1+E1/16/19/5+H1/19/5p4-5^4c4p4-
5I1+L1+E1/16/19/5+H1/19/5p4-5I1+L1+E1/16/19/5+H1/19/5p4-5^3B1/19/5p4-5^3c4p4-5^4c4p4-
5I1+L1+E1/16/19/5+H1/19/5p4-5I1+L1+E1/16/19/5+H1/19/5p4-5^3p2-5p4-5^2c4p4-5^4c4p4-
5I1+L1+E1/16/19/5+H1/19/5p4-5I1+L1+E1/16/19/5+H1/19/5p4-5I1+L1+E1/16/19/5+H1/19/5p4-
5I1+L1+E1/16/19/5+H1/19/5p4-5^3p2-5p4-5^5c4p4-5I1+L1+E1/16/19/5+H1/19/5p4-
5I1+L1+E1/16/19/5+H1/19/5p4-5I1+L1+E1/16/19/5+H1/19/5p4-5I1+L1+E1/16/19/5+H1/19/5p4-5^3p2-
5p4-5^3c4p4-5^2c4p4-5I1+L1+E1/16/19/5+H1/19/5p4-5c4p4-5I1+L1+E1/16/19/5+H1/19/5c4p4-5^2c4p4-

$_5$I1+L1+E1/16/19/5+H1/19/5$c_4p_{4-5}^2c_4p_{4-5}$I1+L1+E1/16/19/5+H1/19/5$p_{4-5}c_4p_{4-}$
$_5$I1+L1+E1/16/19/5+H1/19/5$c_4p_{4-5}^2c_4p_{4-5}c_4p_{4-5}^2c_4p_{4-5}$I1+L1+E1/16/19/5+H1/19/5$p_{4-5}c_4p_{4-}$
$_5$I1+L1+E1/16/19/5+H1/19/5$c_4p_{4-5}^2c_4p_{4-5}$I1+L1+E1/16/19/5+H1/19/5$c_4p_{4-5}^2c_4p_{4-}$
$_5$I1+L1+E1/16/19/5+H1/19/5$p_{4-5}c_4p_{4-5}$I1+L1+E1/16/19/5+H1/19/5$c_4p_{4-5}^2c_4p_{4-}$
$_5$I1+L1+E1/16/19/5+H1/19/5$p_{4-5}c_4p_{4-5}$I1+L1+E1/16/19/5+H1/19/5$p_{4-}$
$_5c_4$D1/19/5+P1+F1/19+G1/19/5+V1/19.1I1+L1+E1/16/19/5+H1/19/5$p_{4-5}c_4p_{4-}$
$_5$I1+L1+E1/16/19/5+H1/19/5$p_{4-5}c_4p_{4-5}^2c_4p_{4-5}$I1+L1+E1/16/19/5+H1/19/5$p_{4-}$
$_5c_4$D1/19/5+P1+F1/19+G1/19/5+V1/19.1I1+L1+E1/16/19/5+H1/19/5$p_{4-5}c_4p_{4-}$
$_5$I1+L1+E1/16/19/5+H1/19/5$p_{4-5}c_4p_{4-5}$I1+L1+E1/16/19/5+H1/19/5$p_{4-5}c_4p_{4-}$
$_5$I1+L1+E1/16/19/5+H1/19/5$p_{4-5}c_4$I1+L1+E1/16/19/5+H1/19/5$p_{4-5}$I1+L1+E1/16/19/5+H1/19/5$p_{4-}$
$_5c_4p_{4-5}$I1+L1+E1/16/19/5+H1/19/5$p_{4-6}p_{2-5}$I1+L1+E1/16/19/5+H1/19/5$p_{4-}$
$_5^2$I1+L1+E1/16/19/5+H1/19/5$p_{3-5}c_4p_{4-5}^2c_4p_{4-5}$I1+L1+E1/16/19/5+H1/19/5$p_{4-}$
$_5^2$I1+L1+E1/16/19/5+H1/19/5$p_{3-5}$I1+L1+E1/16/19/5+H1/19/5$p_{4-5}c_4p_{4-5}^2c_4p_{4-}$
$_5$I1+L1+E1/16/19/5+H1/19/5$p_{4-5}c_4p_{4-5}$I1+L1+E1/16/19/5+H1/19/5$p_{4-}$
$_5^2$I1+L1+E1/16/19/5+H1/19/5$p_{3-5}c_4p_{4-5}$I1+L1+E1/16/19/5+H1/19/5$p_{4-5}c_4p_{4-}$
$_5$I1+L1+E1/16/19/5+H1/19/5$p_{4-}$
$_5c_4$D1/19/5+P1+F1/19+G1/19/5+V1/19.1I1+L1+E1/16/19/5+H1/19/5$p_{4-5}c_4p_{4-}$
$_5$I1+L1+E1/16/19/5+H1/19/$5^2p_{4-5}c_4p_{4-5}$I1+L1+E1/16/19/5+H1/19/5$p_{4-}$
$_5^2$I1+L1+E1/16/19/5+H1/19/5$p_{3-5}p_{4-5}c_4p_{4-5}$I1+L1+E1/16/19/5+H1/19/5$p_{4-}$
$_5^2$I1+L1+E1/16/19/5+H1/19/5$p_{3-5}$I1+L1+E1/16/19/5+H1/19/5$p_{4-5}c_4p_{4-}$
$_5^2$I1+L1+E1/16/19/5+H1/19/5D1/19/5+P1+F1/19+G1/19/5+V1/19.1$p_{1-5}p_{4-}$
$_5$I1+L1+E1/16/19/5+H1/19/5$p_{4-5}c_4p_{4-5}^2c_4p_{4-5}^2$I1+L1+E1/16/19/5+H1/19/$5^2c_4p_{4-}$
$_5^3$I1+L1+E1/16/19/5+H1/19/5$p_{3-5}$I1+L1+E1/16/19/5+H1/19/5$c_4p_{4-5}^2c_4^2p_{4-}$
$_5$I1+L1+E1/16/19/5+H1/19/5$p_{4-5}c_4p_{4-5}$I1+L1+E1/16/19/5+H1/19/5$p_{4-5}c_4p_{4-}$
$_5$I1+L1+E1/16/19/5+H1/19/5$c_4p_{4-5}^2c_4^2p_{4-5}$I1+L1+E1/16/19/5+H1/19/5$p_{4-5}c_4p_{4-}$
$_5$I1+L1+E1/16/19/5+H1/19/5$p_{4-5}c_4p_{4-5}$I1+L1+E1/16/19/5+H1/19/5$p_{4-5}^2c_4p_{4-}$
$_5$I1+L1+E1/16/19/5+H1/19/5$p_{4-}$
$_5$D1/19/5+P1+F1/19+G1/19/5+V1/19.1I1+L1+E1/16/19/5+H1/19/5$p_{4-5}^2c_4p_{4-}$
$_5$I1+L1+E1/16/19/5+H1/19/5$p_{4-}$
$_5$D1/19/5+P1+F1/19+G1/19/5+V1/19.1I1+L1+E1/16/19/5+H1/19/5$p_{4-5}c_4p_{4-5}^3c_4p_{4-}$
$_5$I1+L1+E1/16/19/5+H1/19/5$p_{4-5}c_4p_{4-5}$I1+L1+E1/16/19/5+H1/19/5$p_{4-5}c_4p_{4-}$
$_5$I1+L1+E1/16/19/5+H1/19/5$p_{4-5}c_4p_{4-5}$I1+L1+E1/16/19/5+H1/19/5$p_{4-5}^2c_4p_{4-}$
$_5$I1+L1+E1/16/19/5+H1/19/5$p_{4-}$
$_5$D1/19/5+P1+F1/19+G1/19/5+V1/19.1I1+L1+E1/16/19/5+H1/19/5$p_{4-5}c_4p_{4-5}^3c_4p_{4-}$
$_5^2$I1+L1+E1/16/19/5+H1/19/5$p_{4-5}c_4p_{4-5}$I1+L1+E1/16/19/5+H1/19/5$p_{4-5}c_4p_{4-}$
$_5^2$I1+L1+E1/16/19/5+H1/19/5$p_{4-5}c_4p_{4-}$
$_5$D1/19/5+P1+F1/19+G1/19/5+V1/19.1I1+L1+E1/16/19/5+H1/19/5$p_{4-5}c_4p_{4-}$
$_5$I1+L1+E1/16/19/5+H1/19/5$p_{4-5}c_4p_{4-5}$I1+L1+E1/16/19/5+H1/19/5$p_{4-5}c_4p_{4-}$
$_5$I1+L1+E1/16/19/5+H1/19/5$p_{4-5}c_4p_{4-5}$I1+L1+E1/16/19/5+H1/19/5$p_{4-5}c_4p_{4-}$
$_5$I1+L1+E1/16/19/5+H1/19/5$p_{4-5}^2c_4p_{4-5}$I1+L1+E1/16/19/5+H1/19/5$p_{4-}$
$_5$D1/19/5+P1+F1/19+G1/19/5+V1/19.1I1+L1+E1/16/19/5+H1/19/5$p_{4-5}c_4p_{4-5}^3c_4p_{4-}$
$_5^2$I1+L1+E1/16/19/5+H1/19/5$p_{4-5}c_4p_{4-5}$I1+L1+E1/16/19/5+H1/19/5$p_{4-5}c_4p_{4-}$
$_5$I1+L1+E1/16/19/5+H1/19/5$p_{4-5}c_4p_{4-5}^2$I1+L1+E1/16/19/5+H1/19/5$p_{4-5}c_4p_{4-}$
$_5^2$I1+L1+E1/16/19/5+H1/19/5$p_{4-5}c_4p_{4-5}$I1+L1+E1/16/19/5+H1/19/5$p_{4-5}c_4p_{4-}$
$_5$I1+L1+E1/16/19/5+H1/19/5$p_{4-5}c_4p_{4-5}^2$I1+L1+E1/16/19/5+H1/19/5$p_{4-5}c_4p_{4-}$
$_5$I1+L1+E1/16/19/5+H1/19/5$p_{4-5}c_4p_{4-5}$I1+L1+E1/16/19/5+H1/19/5$p_{4-5}c_4p_{4-}$
$_5$I1+L1+E1/16/19/5+H1/19/5$p_{4-5}^2c_4p_{4-5}^3$I1+L1+E1/16/19/5+H1/19/5$p_{4-5}c_4p_{4-}$
$_5^3c_5$I1+L1+E1/16/19/5+H1/19/$5^2p_{4-5}c_4p_{4-5}c_5$I1+L1+E1/16/19/5+H1/19/$5^2p_{4-5}c_4p_{4-5}^2c_4p_{4-5}^2c_4p_{4-}$
$_5$I1+L1+E1/16/19/5+H1/19/5$p_{4-5}c_4p_{4-5}$I1+L1+E1/16/19/5+H1/19/5$p_{4-5}c_4p_{4-}$
$_5$I1+L1+E1/16/19/5+H1/19/5$p_{4-5}c_4p_{4-5}^2c_4p_{4-5}^2c_4p_{4-5}$I1+L1+E1/16/19/5+H1/19/5$p_{4-5}c_4p_{4-5}^2c_4p_{4-5}^2c_4p_{4-}$
$_5c_4p_{4-5}^2c_4p_{4-5}$I1+L1+E1/16/19/5+H1/19/5$p_{4-5}c_4p_{4-5}$I1+L1+E1/16/19/5+H1/19/5$p_{4-5}c_4p_{4-}$

$_5$I1+L1+E1/16/19/5+H1/19/5$p_{4-5}c_4p_{4-5}$I1+L1+E1/16/19/5+H1/19/5$p_{4-5}c_4p_{4-}$

$_5$I1+L1+E1/16/19/5+H1/19/5$p_{4-5}c_4p_{4-5}{}^2c_4p_{4-5}{}^2c_4p_{4-5}{}^2c_4p_{4-5}$I1+L1+E1/16/19/5+H1/19/5$p_{4-5}c_4p_{4-5}{}^2c_4p_{4-}$

$_5$I1+L1+E1/16/19/5+H1/19/5$p_{4-5}c_4p_{4-5}{}^2c_4p_{4-5}$I1+L1+E1/16/19/5+H1/19/5$p_{4-5}c_4p_{4-}$

$_5$I1+L1+E1/16/19/5+H1/19/5$c_4p_{4-5}$I1+L1+E1/16/19/5+H1/19/5$p_{4-5}c_4p_{4-5}{}^3c_4p_{4-}$

$_5$I1+L1+E1/16/19/5+H1/19/5$p_{4-5}{}^3c_4p_{4-5}$I1+L1+E1/16/19/5+H1/19/5$p_{4-5}{}^3c_4p_{4-}$

$_5$I1+L1+E1/16/19/5+H1/19/5$p_{4-5}{}^3c_4p_{4-5}$I1+L1+E1/16/19/5+H1/19/5$p_{4-5}{}^3c_4p_{4-}$

$_5$I1+L1+E1/16/19/5+H1/19/5$p_{4-5}{}^3c_4p_{4-5}$I1+L1+E1/16/19/5+H1/19/5$p_{4-5}{}^3c_4p_{4-}$

$_5$I1+L1+E1/16/19/5+H1/19/5$p_{4-5}{}^3c_4p_{4-5}$I1+L1+E1/16/19/5+H1/19/5$p_{4-5}{}^3c_4p_{4-}$

$_5$I1+L1+E1/16/19/5+H1/19/5$p_{4-5}{}^3c_4p_{4-5}{}^2c_4p_{4-5}$I1+L1+E1/16/19/5+H1/19/5$p_{4-5}c_4p_{4-}$

$_5$I1+L1+E1/16/19/5+H1/19/5$p_{4-5}c_4p_{4-5}$I1+L1+E1/16/19/5+H1/19/5$p_{4-5}c_4p_{4-}$

$_5$I1+L1+E1/16/19/5+H1/19/5$p_{4-5}c_4p_{4-5}$I1+L1+E1/16/19/5+H1/19/5$p_{4-}$

$_5{}^3$I1+L1+E1/16/19/5+H1/19/5$p_{4-5}c_4p_{4-5}{}^2c_4p_{4-5}$I1+L1+E1/16/19/5+H1/19/5$p_{4-5}c_4p_{4-}$

$_5$I1+L1+E1/16/19/5+H1/19/5$p_{4-5}c_4p_{4-5}$I1+L1+E1/16/19/5+H1/19/5$p_{4-5}c_4p_{4-}$

$_5$I1+L1+E1/16/19/5+H1/19/5$p_{4-5}{}^3$I1+L1+E1/16/19/5+H1/19/5$p_{4-5}c_4p_{4-}$

$_5$I1+L1+E1/16/19/5+H1/19/5$p_{4-5}{}^3c_4p_{4-5}$I1+L1+E1/16/19/5+H1/19/5$p_{4-}$

$_5{}^3$I1+L1+E1/16/19/5+H1/19/5$p_{4-5}c_4p_{4-5}$I1+L1+E1/16/19/5+H1/19/5$p_{4-5}{}^2c_4p_{4-}$

$_5$I1+L1+E1/16/19/5+H1/19/5$p_{4-5}c_4p_{4-5}$I1+L1+E1/16/19/5+H1/19/5$p_{4-}$

$_5c_4$D1/19/5+P1+F1/19+G1/19/5+V1/19.1I1+L1+E1/16/19/5+H1/19/5$p_{4-}$

$_5c_4$I1+L1+E1/16/19/5+H1/19/5$p_{4-5}$I1+L1+E1/16/19/5+H1/19/5$p_{4-6}p_{2-}$

$_5$I1+L1+E1/16/19/5+H1/19/5$p_{4-5}{}^2$I1+L1+E1/16/19/5+H1/19/5$p_{3-5}c_4p_{4-5}{}^2c_4p_{4-}$

$_5$I1+L1+E1/16/19/5+H1/19/5$p_{4-5}{}^2$I1+L1+E1/16/19/5+H1/19/5$p_{3-5}$I1+L1+E1/16/19/5+H1/19/5$p_{4-}$

$_5c_4p_{4-5}{}^2c_4p_{4-5}$I1+L1+E1/16/19/5+H1/19/5$p_{4-5}c_4p_{4-5}$I1+L1+E1/16/19/5+H1/19/5$p_{4-}$

$_5{}^2$I1+L1+E1/16/19/5+H1/19/5$p_{3-5}$I1+L1+E1/16/19/5+H1/19/5$p_{4-5}c_4p_{4-5}{}^2c_4p_{4-}$

$_5$I1+L1+E1/16/19/5+H1/19/5$p_{4-5}c_4p_{4-5}$I1+L1+E1/16/19/5+H1/19/5$p_{4-}$

$_5c_4$D1/19/5+P1+F1/19+G1/19/5+V1/19.1I1+L1+E1/16/19/5+H1/19/5$p_{4-5}c_4p_{4-}$

$_5$I1+L1+E1/16/19/5+H1/19/5$p_{4-5}{}^2c_4p_{4-5}$I1+L1+E1/16/19/5+H1/19/5$p_{4-}$

$_5{}^2$I1+L1+E1/16/19/5+H1/19/5$p_{3-5}$I1+L1+E1/16/19/5+H1/19/5$p_{4-5}c_4p_{4-}$

$_5{}^2$I1+L1+E1/16/19/5+H1/19/5$p_{4-5}c_4p_{4-5}{}^2c_4p_{4-5}$I1+L1+E1/16/19/5+H1/19/5$p_{4-}$

$_5{}^2$I1+L1+E1/16/19/5+H1/19/5$p_{3-5}$I1+L1+E1/16/19/5+H1/19/5$p_{4-5}c_4p_{4-}$

$_5$I1+L1+E1/16/19/5+H1/19/5$^2p_{4-5}c_4p_{4-5}{}^3$I1+L1+E1/16/19/5+H1/19/5$p_{3-}$

$_5$I1+L1+E1/16/19/5+H1/19/5$c_4p_{4-5}{}^2c_4{}^2p_{4-5}$I1+L1+E1/16/19/5+H1/19/5$p_{4-5}c_4p_{4-}$

$_5$I1+L1+E1/16/19/5+H1/19/5$p_{4-5}c_4p_{4-5}$I1+L1+E1/16/19/5+H1/19/5$p_{4-5}c_4p_{4-}$

$_5$I1+L1+E1/16/19/5+H1/19/5$p_{4-5}{}^2c_4p_{4-5}$I1+L1+E1/16/19/5+H1/19/5$p_{4-}$

$_5$D1/19/5+P1+F1/19+G1/19/5+V1/19.1I1+L1+E1/16/19/5+H1/19/5$p_{4-5}{}^2c_4p_{4-}$

$_5$I1+L1+E1/16/19/5+H1/19/5$p_{4-}$

$_5$D1/19/5+P1+F1/19+G1/19/5+V1/19.1I1+L1+E1/16/19/5+H1/19/5$p_{4-5}c_4p_{4-5}{}^3c_4p_{4-}$

$_5$I1+L1+E1/16/19/5+H1/19/5$p_{4-5}{}^4$I1+L1+E1/16/19/5+H1/19/5$p_{3-5}$I1+L1+E1/16/19/5+H1/19/5$c_4p_{4-}$

$_5{}^2c_4p_{4-6}$I1+L1+E1/16/19/5+H1/19/5$^2p_{4-5}c_4p_{4-5}$I1+L1+E1/16/19/5+H1/19/5$p_{4-5}c_4p_{4-}$

$_5$I1+L1+E1/16/19/5+H1/19/5$p_{4-5}c_4p_{4-5}$I1+L1+E1/16/19/5+H1/19/5$p_{4-5}{}^2c_4p_{4-}$

$_5$I1+L1+E1/16/19/5+H1/19/5$p_{4-}$

$_5$D1/19/5+P1+F1/19+G1/19/5+V1/19.1I1+L1+E1/16/19/5+H1/19/5$p_{4-5}{}^2c_4p_{4-}$

$_5$I1+L1+E1/16/19/5+H1/19/5$p_{4-}$

$_5$D1/19/5+P1+F1/19+G1/19/5+V1/19.1I1+L1+E1/16/19/5+H1/19/5$p_{4-5}c_4p_{4-5}{}^3c_4p_{4-}$

$_5$I1+L1+E1/16/19/5+H1/19/5$p_{4-5}c_4p_{4-5}$I1+L1+E1/16/19/5+H1/19/5$p_{4-5}c_4p_{4-}$

$_5$I1+L1+E1/16/19/5+H1/19/5$p_{4-5}c_4p_{4-5}$I1+L1+E1/16/19/5+H1/19/5$p_{4-5}{}^2c_4p_{4-}$

$_5$I1+L1+E1/16/19/5+H1/19/5$p_{4-}$

$_5$D1/19/5+P1+F1/19+G1/19/5+V1/19.1I1+L1+E1/16/19/5+H1/19/5$p_{4-5}c_4p_{4-}$

$_5{}^3c_4$I1+L1+E1/16/19/5+H1/19/5$p_{4-5}$I1+L1+E1/16/19/5+H1/19/5$c_4p_{4-}$

$_5$I1+L1+E1/16/19/5+H1/19/5$p_{4-5}c_4p_{4-5}$I1+L1+E1/16/19/5+H1/19/5$p_{4-5}{}^2c_4p_{4-}$

$_5$I1+L1+E1/16/19/5+H1/19/5$p_{4-}$

$_5$D1/19/5+P1+F1/19+G1/19/5+V1/19.1I1+L1+E1/16/19/5+H1/19/5$p_{4-5}c_4p_{4-5}{}^3c_4p_{4-}$

$_5^2$I1+L1+E1/16/19/5+H1/19/5p$_{4-5}$c$_4$p$_{4-5}$I1+L1+E1/16/19/5+H1/19/5p$_{4-5}$c$_4$p$_{4-}$

$_5$I1+L1+E1/16/19/5+H1/19/5p$_{4-5}$c$_4$p$_{4-5}$$^2$I1+L1+E1/16/19/5+H1/19/5p$_{4-5}$c$_4$p$_{4-}$

$_5^2$I1+L1+E1/16/19/5+H1/19/5p$_{4-5}$c$_4$p$_{4-5}$I1+L1+E1/16/19/5+H1/19/5p$_{4-5}$c$_4$p$_{4-}$

$_5$I1+L1+E1/16/19/5+H1/19/5p$_{4-5}$c$_4$p$_{4-5}$$^2$I1+L1+E1/16/19/5+H1/19/5p$_{4-5}$c$_4$p$_{4-}$

$_5$I1+L1+E1/16/19/5+H1/19/5p$_{4-5}$c$_4$p$_{4-5}$I1+L1+E1/16/19/5+H1/19/5p$_{4-5}$c$_4$p$_{4-}$

$_5$I1+L1+E1/16/19/5+H1/19/5p$_{4-5}$$^2$c$_4$p$_{4-5}$I1+L1+E1/16/19/5+H1/19/5p$_{4-}$

$_5$D1/19/5+P1+F1/19+G1/19/5+V1/19.1I1+L1+E1/16/19/5+H1/19/5p$_{4-5}$c$_4$p$_{4-}$

$_5$I1+L1+E1/16/19/5+H1/19/5p$_{4-5}$D1/19/5+P1+F1/19+G1/19/5+V1/19.1c$_6$p$_{4-5}$$^2$c$_4$p$_{4-}$

$_5^2$I1+L1+E1/16/19/5+H1/19/5p$_{4-5}$c$_4$p$_{4-5}$I1+L1+E1/16/19/5+H1/19/5p$_{4-5}$c$_4$p$_{4-}$

$_5$I1+L1+E1/16/19/5+H1/19/5p$_{4-5}$c$_4$p$_{4-5}$$^2$I1+L1+E1/16/19/5+H1/19/5p$_{4-5}$c$_4$p$_{4-}$

$_5^2$I1+L1+E1/16/19/5+H1/19/5p$_{4-5}$c$_4$p$_{4-5}$I1+L1+E1/16/19/5+H1/19/5p$_{4-}$

$_5^2$I1+L1+E1/16/19/5+H1/19/5p$_{4-5}$c$_4$p$_{4-5}$I1+L1+E1/16/19/5+H1/19/5p$_{4-5}$c$_4$p$_{4-}$

$_5^2$I1+L1+E1/16/19/5+H1/19/5p$_{4-5}$c$_4$p$_{4-5}$$^2$I1+L1+E1/16/19/5+H1/19/5p$_{4-5}$c$_4$p$_{4-}$

$_5$I1+L1+E1/16/19/5+H1/19/5p$_{4-5}$c$_4$p$_{4-5}$I1+L1+E1/16/19/5+H1/19/5p$_{4-}$

$_5^3$I1+L1+E1/16/19/5+H1/19/5$^2$c$_4$p$_{4-5}$I1+L1+E1/16/19/5+H1/19/5p$_{4-5}$$^2$c$_4$p$_{4-}$

$_5^3$I1+L1+E1/16/19/5+H1/19/5p$_{4-5}$c$_4$p$_{4-5}$$^3$c$_5$I1+L1+E1/16/19/5+H1/19/5$^2$p$_{4-5}$c$_4$p$_{4-}$

$_5$I1+L1+E1/16/19/5+H1/19/5p$_{4-5}$$^2$c$_4$p$_{4-5}$I1+L1+E1/16/19/5+H1/19/5p$_{4-5}$$^2$c$_4$p$_{4-}$

$_5$I1+L1+E1/16/19/5+H1/19/5p$_{4-5}$c$_4$p$_{4-5}$I1+L1+E1/16/19/5+H1/19/5p$_{4-5}$c$_4$p$_{4-5}$$^2$c$_4$p$_{4-5}$$^2$c$_4$p$_{4-}$

$_5$I1+L1+E1/16/19/5+H1/19/5p$_{4-5}$c$_4$p$_{4-5}$$^2$c$_4$p$_{4-5}$I1+L1+E1/16/19/5+H1/19/5$^2$c$_4$p$_{4-}$

$_5$I1+L1+E1/16/19/5+H1/19/5c$_4$p$_{4-5}$I1+L1+E1/16/19/5+H1/19/5p$_{4-5}$c$_4$p$_{4-5}$$^3$c$_4$p$_{4-}$

$_5$I1+L1+E1/16/19/5+H1/19/5p$_{4-}$

$_5^3$c$_4$D1/19/5+P1+F1/19+G1/19/5+V1/19.1I1+L1+E1/16/19/5+H1/19/5p$_{4-5}$$^3$c$_4$p$_{4-}$

$_5$I1+L1+E1/16/19/5+H1/19/5p$_{4-5}$$^3$c$_4$p$_{4-5}$I1+L1+E1/16/19/5+H1/19/5p$_{4-5}$$^3$c$_4$p$_{4-}$

$_5$I1+L1+E1/16/19/5+H1/19/5p$_{4-5}$$^3$c$_4$p$_{4-5}$I1+L1+E1/16/19/5+H1/19/5p$_{4-5}$$^3$c$_4$p$_{4-}$

$_5$I1+L1+E1/16/19/5+H1/19/5p$_{4-5}$$^3$c$_4$p$_{4-5}$I1+L1+E1/16/19/5+H1/19/5p$_{4-5}$$^3$c$_4$p$_{4-}$

$_5$I1+L1+E1/16/19/5+H1/19/5p$_{4-5}$$^3$c$_4$p$_{4-5}$$^2$c$_4$p$_{4-5}$I1+L1+E1/16/19/5+H1/19/5p$_{4-5}$c$_4$p$_{4-}$

$_5$I1+L1+E1/16/19/5+H1/19/5p$_{4-5}$c$_4$p$_{4-5}$I1+L1+E1/16/19/5+H1/19/5p$_{4-5}$c$_4$p$_{4-}$

$_5$I1+L1+E1/16/19/5+H1/19/5p$_{4-5}$c$_4$p$_{4-5}$I1+L1+E1/16/19/5+H1/19/5p$_{4-}$

$_5^3$I1+L1+E1/16/19/5+H1/19/5p$_{4-5}$c$_4$p$_{4-5}$$^2$c$_4$p$_{4-5}$I1+L1+E1/16/19/5+H1/19/5p$_{4-5}$c$_4$p$_{4-}$

$_5$I1+L1+E1/16/19/5+H1/19/5p$_{4-5}$c$_4$p$_{4-5}$I1+L1+E1/16/19/5+H1/19/5p$_{4-5}$c$_4$p$_{4-}$

$_5$I1+L1+E1/16/19/5+H1/19/5p$_{4-5}$$^3$I1+L1+E1/16/19/5+H1/19/5p$_{4-5}$c$_4$p$_{4-}$

$_5$I1+L1+E1/16/19/5+H1/19/5p$_{4-5}$$^2$I1+L1+E1/16/19/5+H1/19/5c$_4$p$_{4-5}$I1+L1+E1/16/19/5+H1/19/5p$_{4-}$

$_5^3$I1+L1+E1/16/19/5+H1/19/5p$_{4-5}$c$_4$p$_{4-5}$I1+L1+E1/16/19/5+H1/19/5p$_{4-5}$$^2$c$_4$p$_{4-}$

$_5$I1+L1+E1/16/19/5+H1/19/5p$_{4-5}$c$_4$p$_{4-5}$I1+L1+E1/16/19/5+H1/19/5p$_{4-}$

$_5^2$I1+L1+E1/16/19/5+H1/19/5c$_4$p$_{4-5}$I1+L1+E1/16/19/5+H1/19/5p$_{4-5}$c$_4$p$_{4-}$

$_5$I1+L1+E1/16/19/5+H1/19/5p$_{4-}$

$_5^3$c$_4$D1/19/5+P1+F1/19+G1/19/5+V1/19.1I1+L1+E1/16/19/5+H1/19/5p$_{4-5}$c$_4$p$_{4-}$

$_5$I1+L1+E1/16/19/5+H1/19/5p$_{4-}$

$_5$c$_4$D1/19/5+P1+F1/19+G1/19/5+V1/19.1I1+L1+E1/16/19/5+H1/19/5p$_{4-}$

$_5^2$I1+L1+E1/16/19/5+H1/19/5c$_4$p$_{4-5}$$^3$c$_4$p$_{4-5}$I1+L1+E1/16/19/5+H1/19/5c$_4$p$_{4-}$

$_5$I1+L1+E1/16/19/5+H1/19/5p$_{4-5}$c$_3$p$_{3-5}$c$_4$p$_{4-5}$I1+L1+E1/16/19/5+H1/19/5p$_{4-5}$$^2$c$_4$p$_{4-}$

$_5$I1+L1+E1/16/19/5+H1/19/5p$_{4-5}$$^2$I1+L1+E1/16/19/5+H1/19/5p$_{4-}$

$_5$c$_4$I1+L1+E1/16/19/5+H1/19/5c$_4$p$_{4-5}$I1+L1+E1/16/19/5+H1/19/5c$_4$p$_{4-}$

$_5$I1+L1+E1/16/19/5+H1/19/5p$_{4-5}$$^2$I1+L1+E1/16/19/5+H1/19/5p$_{4-5}$$^3$I1+L1+E1/16/19/5+H1/19/5p$_{4-}$

$_5$c$_4$p$_{4-5}$I1+L1+E1/16/19/5+H1/19/5p$_{4-5}$c$_4$p$_{4-5}$$^2$I1+L1+E1/16/19/5+H1/19/5p$_{4-}$

$_5^2$I1+L1+E1/16/19/5+H1/19/5p$_{4-5}$p$_{5-3}$p$_{2-5}$p$_{4-5}$I1+L1+E1/16/19/5+H1/19/5c$_4$p$_{4-}$

$_5$I1+L1+E1/16/19/5+H1/19/5p$_{4-5}$c$_4$p$_{4-5}$I1+L1+E1/16/19/5+H1/19/5c$_4$p$_{4-}$

$_5$I1+L1+E1/16/19/5+H1/19/5p$_{4-5}$$^2$I1+L1+E1/16/19/5+H1/19/5p$_{4-5}$$^3$I1+L1+E1/16/19/5+H1/19/5p$_{4-}$

$_5$c$_4$p$_{4-5}$I1+L1+E1/16/19/5+H1/19/5p$_{4-5}$c$_4$p$_{4-5}$$^2$I1+L1+E1/16/19/5+H1/19/5p$_{4-}$

$_5^2$I1+L1+E1/16/19/5+H1/19/5p$_{4-5}$c$_5$I1+L1+E1/16/19/5+H1/19/5p$_{4-}$

$_5$I1+L1+E1/16/19/5+H1/19/5c$_4$p$_{4-5}$I1+L1+E1/16/19/5+H1/19/5p$_{4-5}$c$_4$I1+L1+E1/16/19/5+H1/19/5p$_{4-}$

$_5^2$I1+L1+E1/16/19/5+H1/19/5$p_{4-}$
$_5c_4$D1/19/5+P1+F1/19+G1/19/5+V1/19.1I1+L1+E1/16/19/5+H1/19/5$p_{4-}$
$_5^2$I1+L1+E1/16/19/5+H1/19/5$c_4^2$D1/19/5+P1+F1/19+G1/19/5+V1/19.1I1+L1+E1/16/19/5+H1/19/
5$p_{4-5}^2$I1+L1+E1/16/19/5+H1/19/5$c_4^2p_{4-5}^2c_4^2p_{4-5}^2$I1+L1+E1/16/19/5+H1/19/5$c_4p_{4-}$
$_5$I1+L1+E1/16/19/5+H1/19/5$p_{4-5}^4$I1+L1+E1/16/19/5+H1/19/5$p_{4-}$
$_5c_4$D1/19/5+P1+F1/19+G1/19/5+V1/19.1I1+L1+E1/16/19/5+H1/19/5$p_{4-5}^3c_4^2p_{4-}$
$_5c_4$I1+L1+E1/16/19/5+H1/19/5$p_{4-5}^4$I1+L1+E1/16/19/5+H1/19/5$p_{4-}$
$_5c_4$D1/19/5+P1+F1/19+G1/19/5+V1/19.1I1+L1+E1/16/19/5+H1/19/5$p_{4-}$
$_5^3c_4^2$D1/19/5+P1+F1/19+G1/19/5+V1/19.1I1+L1+E1/16/19/5+H1/19/5$p_{4-}$
$_5^4$I1+L1+E1/16/19/5+H1/19/5$^2p_{4-5}c_4p_{4-5}^2c_4p_{4-5}$I1+L1+E1/16/19/5+H1/19/5$c_4p_{4-}$
$_5^2$I1+L1+E1/16/19/5+H1/19/5$c_4$D1/19/5+P1+F1/19+G1/19/5+V1/19.1$c_4$I1+L1+E1/16/19/5+H1/19
/5$c_5$I1+L1+E1/16/19/5+H1/19/5$p_{4-5}$I1+L1+E1/16/19/5+H1/19/5$p_{4-}$
$_5^2$I1+L1+E1/16/19/5+H1/19/5$^2p_{4-5}c_4p_{4-5}^2c_4p_{4-5}$I1+L1+E1/16/19/5+H1/19/5$c_4p_{4-}$
$_5^2$I1+L1+E1/16/19/5+H1/19/5$c_4$D1/19/5+P1+F1/19+G1/19/5+V1/19.1$c_4$I1+L1+E1/16/19/5+H1/19
/5$c_5$I1+L1+E1/16/19/5+H1/19/5$p_{4-5}$I1+L1+E1/16/19/5+H1/19/5$c_4$I1+L1+E1/16/19/5+H1/19/5$p_{4-}$
$_5c_4$D1/19/5+P1+F1/19+G1/19/5+V1/19.1I1+L1+E1/16/19/5+H1/19/5$p_{4-}$
$_5^3c_4$D1/19/5+P1+F1/19+G1/19/5+V1/19.1I1+L1+E1/16/19/5+H1/19/5$p_{4-}$
$_5^2$I1+L1+E1/16/19/5+H1/19/5$p_{4-5}$I1+L1+E1/16/19/5+H1/19/5$^2p_{4-5}c_4p_{4-5}^2c_4p_{4-}$
$_5$D1/19/5+P1+F1/19+G1/19/5+V1/19.1I1+L1+E1/16/19/5+H1/19/5$p_{4-5}c_4p_{4-}$
$_5$I1+L1+E1/16/19/5+H1/19/5$p_{4-5}^3$I1+L1+E1/16/19/5+H1/19/5$p_{4-5}^2$I1+L1+E1/16/19/5+H1/19/5$p_{4-}$
$_5c_4p_{4-5}^3c_4p_{4-5}$I1+L1+E1/16/19/5+H1/19/5$p_{4-}$
$_5$I1+L1+E1/16/19/5+H1/19/5$c_4$D1/19/5+P1+F1/19+G1/19/5+V1/19.1$c_4p_{4-}$
$_5c_5$I1+L1+E1/16/19/5+H1/19/5D1/19/5+P1+F1/19+G1/19/5+V1/19.1I1+L1+E1/16/19/5+H1/19/5c
$_4$I1+L1+E1/16/19/5+H1/19/5$p_{4-5}^2$I1+L1+E1/16/19/5+H1/19/5$p_{4-}$
$_5c_4$D1/19/5+P1+F1/19+G1/19/5+V1/19.1I1+L1+E1/16/19/5+H1/19/5$^2p_{4-}$
$_5^2c_4$D1/19/5+P1+F1/19+G1/19/5+V1/19.1$c_6p_{4-5}^4$I1+L1+E1/16/19/5+H1/19/5$p_{4-}$
$_5c_4$D1/19/5+P1+F1/19+G1/19/5+V1/19.1I1+L1+E1/16/19/5+H1/19/5$p_{4-}$
$_5^3c_4^2$I1+L1+E1/16/19/5+H1/19/5$p_{4-5}$I1+L1+E1/16/19/5+H1/19/5$^3c_4p_{4-}$
$_5^5$I1+L1+E1/16/19/5+H1/19/5$p_{4-}$
$_5c_4$D1/19/5+P1+F1/19+G1/19/5+V1/19.1I1+L1+E1/16/19/5+H1/19/5$p_{4-}$
$_5^3c_4^2$I1+L1+E1/16/19/5+H1/19/5$p_{4-5}$I1+L1+E1/16/19/5+H1/19/5$^3c_4p_{4-}$
$_5$I1+L1+E1/16/19/5+H1/19/5$p_{4-5}c_4p_{4-5}$I1+L1+E1/16/19/5+H1/19/5$p_{4-}$
$_5$I1+L1+E1/16/19/5+H1/19/5$p_{4-5}c_4p_{4-5}$I1+L1+E1/16/19/5+H1/19/5$p_{4-}$
$_5^2$I1+L1+E1/16/19/5+H1/19/5$p_{4-5}$D1/19/5+P1+F1/19+G1/19/5+V1/19.1$p_{4-}$
$_5^2$I1+L1+E1/16/19/5+H1/19/5$c_4p_{4-5}$I1+L1+E1/16/19/5+H1/19/5$p_{4-5}c_4p_{4-}$
$_5^4$I1+L1+E1/16/19/5+H1/19/5$p_{4-5}c_4p_{4-}$
$_5$I1+L1+E1/16/19/5+H1/19/5$^2c_5$I1+L1+E1/16/19/5+H1/19/5$p_{2-5}p_{4-}$
$_5^2$I1+L1+E1/16/19/5+H1/19/5$^2c_4p_{4-}$
$_5^4c_4$I1+L1+E1/16/19/5+H1/19/5D1/19/5+P1+F1/19+G1/19/5+V1/19.1$c_6$I1+L1+E1/16/19/5+H1/19
/5$^2p_{4-5}p_{2-4}p_{1-5}$I1+L1+E1/16/19/5+H1/19/5$p_{4-5}$I1+L1+E1/16/19/5+H1/19/5$p_{4-5}c_4p_{4-}$
$_5^3$I1+L1+E1/16/19/5+H1/19/5$^2$D1/19/5+P1+F1/19+G1/19/5+V1/19.1$p_{1-}$
$_5c_4$D1/19/5+P1+F1/19+G1/19/5+V1/19.1I1+L1+E1/16/19/5+H1/19/5$p_{4-}$
$_5^2$D1/19/5+P1+F1/19+G1/19/5+V1/19.1$p_{1-5}p_{4-}$
$_5c_4$D1/19/5+P1+F1/19+G1/19/5+V1/19.1I1+L1+E1/16/19/5+H1/19/5$p_{4-}$
$_5$I1+L1+E1/16/19/5+H1/19/5$p_{4-5}c_4p_{4-5}^3c_4p_{4-5}^5c_4p_{4-5}$I1+L1+E1/16/19/5+H1/19/5$p_{4-5}^2c_4p_{4-5}^3c_4p_{4-}$
$_5^3c_4p_{4-5}^3c_4p_{4-5}^3c_4p_{4-5}$I1+L1+E1/16/19/5+H1/19/5$p_{4-5}$I1+L1+E1/16/19/5+H1/19/5$p_{4-5}c_4p_{4-5}^3c_4p_{4-}$
$_5^4c_4p_{4-5}^3$D1/19/5+P1+F1/19+G1/19/5+V1/19.1$p_{1-5}p_{4-5}c_4p_{4-5}$I1+L1+E1/16/19/5+H1/19/5$p_{4-}$
$_5^2$D1/19/5+P1+F1/19+G1/19/5+V1/19.1$p_{1-5}p_{4-5}c_4p_{4-5}$I1+L1+E1/16/19/5+H1/19/5$p_{4-}$
$_5$I1+L1+E1/16/19/5+H1/19/5$p_{4-5}c_4p_{4-5}$I1+L1+E1/16/19/5+H1/19/5$p_{4-}$
$_5$I1+L1+E1/16/19/5+H1/19/5$p_{4-5}c_4p_{4-5}^2c_4p_{4-5}$I1+L1+E1/16/19/5+H1/19/5$p_{4-5}^2c_4p_{4-5}^3c_4p_{4-5}^2c_4p_{4-}$
$_5$I1+L1+E1/16/19/5+H1/19/5$p_{4-5}^2c_4p_{4-5}^3c_4p_{4-5}^3c_4p_{4-5}^3c_4p_{4-5}^3c_4p_{4-5}$I1+L1+E1/16/19/5+H1/19/5$p_{4-}$

$_5$I1+L1+E1/16/19/5+H1/19/5$p_{4-5}c_4p_{4-5}^{3}c_4p_{4-5}^{3}c_4p_{4-5}^{4}c_4p_{4-5}^{3}c_4p_{4-5}^{3}c_4p_{4-5}^{3}c_4p_{4-}$

$_5$I1+L1+E1/16/19/5+H1/19/5$^{2}p_{4-5}^{2}c_4p_{4-5}^{3}c_4p_{4-5}$I1+L1+E1/16/19/5+H1/19/5$p_{4-5}^{3}c_4p_{4-}$

$_5$I1+L1+E1/16/19/5+H1/19/5$p_{4-5}^{4}c_4p_{4-5}^{3}c_4p_{4-5}$I1+L1+E1/16/19/5+H1/19/5$p_{4-5}^{3}c_4p_{4-}$

$_5$I1+L1+E1/16/19/5+H1/19/5$p_{4-5}^{2}c_4p_{4-}$

$_5^{3}c_4$D1/19/5+P1+F1/19+G1/19/5+V1/19.1I1+L1+E1/16/19/5+H1/19/5$p_{4-5}^{3}c_4p_{4-5}^{3}c_4p_{4-}$

$_5$I1+L1+E1/16/19/5+H1/19/5$^{3}p_{4-5}^{3}c_4p_{4-5}^{3}c_4p_{4-5}$I1+L1+E1/16/19/5+H1/19/5$p_{4-5}^{2}c_4p_{4-5}^{3}c_4p_{4-}$

$_5$I1+L1+E1/16/19/5+H1/19/5$p_{4-5}^{3}c_4p_{4-5}^{3}c_4p_{4-5}^{3}c_4p_{4-5}^{3}c_4p_{4-5}$I1+L1+E1/16/19/5+H1/19/5$p_{4-5}^{3}c_4p_{4-}$

$_5$I1+L1+E1/16/19/5+H1/19/5$p_{4-5}c_4^{2}p_{4-5}^{3}c_4p_{4-5}$I1+L1+E1/16/19/5+H1/19/5$p_{4-5}c_4^{2}p_{4-}$

$_5$I1+L1+E1/16/19/5+H1/19/5$p_{4-5}^{2}$I1+L1+E1/16/19/5+H1/19/5$^{2}c_4p_{4-5}$I1+L1+E1/16/19/5+H1/19/5$p_{4-}$

$_5^{2}$I1+L1+E1/16/19/5+H1/19/5$p_{4-5}c_4p_{4-5}$I1+L1+E1/16/19/5+H1/19/5$p_{4-}$

$_5^{2}$I1+L1+E1/16/19/5+H1/19/5$p_{4-5}c_4p_{4-5}$I1+L1+E1/16/19/5+H1/19/5$p_{4-}$

$_5$I1+L1+E1/16/19/5+H1/19/5$c_4p_{4-5}$I1+L1+E1/16/19/5+H1/19/5$p_{4-}$

$_5^{4}$I1+L1+E1/16/19/5+H1/19/5$c_4^{2}p_{4-}$

$_5^{3}c_4$D1/19/5+P1+F1/19+G1/19/5+V1/19.1I1+L1+E1/16/19/5+H1/19/5$p_{4-5}c_4p_{4-}$

$_5$I1+L1+E1/16/19/5+H1/19/5$p_{4-5}c_4^{2}p_{4-5}$I1+L1+E1/16/19/5+H1/19/5$p_{4-}$

$_5^{2}$I1+L1+E1/16/19/5+H1/19/5$^{2}c_4p_{4-5}$I1+L1+E1/16/19/5+H1/19/5$p_{4-5}^{2}$I1+L1+E1/16/19/5+H1/19/5$p_{4-}$

$_5c_4p_{4-5}$I1+L1+E1/16/19/5+H1/19/5$p_{4-5}^{3}c_4p_{4-5}^{2}c_4p_{4-5}$I1+L1+E1/16/19/5+H1/19/5$p_{4-5}^{3}c_4p_{4-}$

$_5$I1+L1+E1/16/19/5+H1/19/5$p_{4-5}c_4^{2}p_{4-5}^{3}c_4p_{4-5}$I1+L1+E1/16/19/5+H1/19/5$p_{4-5}c_4p_{4-}$

$_5$I1+L1+E1/16/19/5+H1/19/5$p_{4-5}^{3}$I1+L1+E1/16/19/5+H1/19/5$p_{4-5}c_4^{2}p_{4-5}^{2}c_4p_{4-}$

$_5$I1+L1+E1/16/19/5+H1/19/5$p_{4-5}c_4p_{4-5}$I1+L1+E1/16/19/5+H1/19/5$p_{4-5}c_4p_{4-}$

$_5$I1+L1+E1/16/19/5+H1/19/5$p_{4-5}^{2}$I1+L1+E1/16/19/5+H1/19/5$p_{4-5}c_4p_{4-}$

$_5$I1+L1+E1/16/19/5+H1/19/5$p_{4-5}^{2}$I1+L1+E1/16/19/5+H1/19/5$^{2}c_4p_{4-}$

$_5$I1+L1+E1/16/19/5+H1/19/5$^{2}c_4p_{4-5}$I1+L1+E1/16/19/5+H1/19/5$p_{4-5}^{3}$I1+L1+E1/16/19/5+H1/19/5$p_{4-}$

$_5^{2}$I1+L1+E1/16/19/5+H1/19/5$p_{4-5}c_4p_{4-5}$I1+L1+E1/16/19/5+H1/19/5$p_{4-5}^{2}c_4^{2}p_{4-}$

$_5$I1+L1+E1/16/19/5+H1/19/5$p_{4-5}^{3}$I1+L1+E1/16/19/5+H1/19/5$p_{4-5}^{4}c_4p_{4-5}^{3}c_4p_{4-}$

$_5$I1+L1+E1/16/19/5+H1/19/5$p_{4-5}^{3}c_4p_{4-5}$I1+L1+E1/16/19/5+H1/19/5$p_{4-5}^{3}c_4p_{4-5}^{3}c_4p_{4-}$

$_5$I1+L1+E1/16/19/5+H1/19/5$p_{4-5}c_4p_{4-5}$I1+L1+E1/16/19/5+H1/19/5$p_{4-5}c_4^{2}p_{4-}$

$_5$I1+L1+E1/16/19/5+H1/19/5$p_{4-5}^{2}$I1+L1+E1/16/19/5+H1/19/5$^{2}c_4p_{4-5}$I1+L1+E1/16/19/5+H1/19/5$p_{4-}$

$_5c_4^{2}p_{4-5}$I1+L1+E1/16/19/5+H1/19/5$p_{4-5}^{2}$I1+L1+E1/16/19/5+H1/19/5$^{2}c_4p_{4-}$

$_5$I1+L1+E1/16/19/5+H1/19/5$p_{4-5}$I1+L1+E1/16/19/5+H1/19/5$p_{4-5}$I1+L1+E1/16/19/5+H1/19/5$p_{4-}$

$_5c_4p_{4-5}$I1+L1+E1/16/19/5+H1/19/5$^{2}c_4p_{4-5}$I1+L1+E1/16/19/5+H1/19/5$p_{4-}$

$_5$I1+L1+E1/16/19/5+H1/19/5$c_4p_{4-5}$I1+L1+E1/16/19/5+H1/19/5$p_{4-5}c_4p_{4-5}c_4p_{4-}$

$_5$I1+L1+E1/16/19/5+H1/19/5$p_{4-5}c_4p_{4-}$

$_5$I1+L1+E1/16/19/5+H1/19/5$c_4$D1/19/5+P1+F1/19+G1/19/5+V1/19.1I1+L1+E1/16/19/5+H1/19/5p

$_{4-5}c_5$I1+L1+E1/16/19/5+H1/19/5$^{2}c_4p_{4-5}c_4p_{4-5}$I1+L1+E1/16/19/5+H1/19/5$c_4p_{4-}$

$_5$I1+L1+E1/16/19/5+H1/19/5$p_{4-5}^{2}c_4p_{4-5}$I1+L1+E1/16/19/5+H1/19/5$p_{4-5}c_4p_{4-}$

$_5$I1+L1+E1/16/19/5+H1/19/5$p_{4-5}c_5$I1+L1+E1/16/19/5+H1/19/5$^{2}c_4p_{4-5}c_4p_{4-}$

$_5$I1+L1+E1/16/19/5+H1/19/5$c_4p_{4-5}$I1+L1+E1/16/19/5+H1/19/5$p_{4-5}^{2}c_4p_{4-}$

$_5$I1+L1+E1/16/19/5+H1/19/5$p_{4-5}^{3}c_4p_{4-5}$I1+L1+E1/16/19/5+H1/19/5$p_{4-5}^{2}c_4p_{4-}$

$_5$I1+L1+E1/16/19/5+H1/19/5$c_4p_{4-5}$I1+L1+E1/16/19/5+H1/19/5$p_{4-5}c_4p_{4-}$

$_5$I1+L1+E1/16/19/5+H1/19/5$c_4p_{4-5}^{2}$I1+L1+E1/16/19/5+H1/19/5$p_{4-}$

$_5^{3}$I1+L1+E1/16/19/5+H1/19/5$c_4p_{4-5}^{2}$I1+L1+E1/16/19/5+H1/19/5$p_{4-5}^{2}c_4p_{4-}$

$_5$I1+L1+E1/16/19/5+H1/19/5$c_4$I1+L1+E1/16/19/5+H1/19/5$c_4p_{4-5}$I1+L1+E1/16/19/5+H1/19/5$c_4p_{4-}$

$_5$I1+L1+E1/16/19/5+H1/19/5$p_{4-}$

$_5^{3}$D1/19/5+P1+F1/19+G1/19/5+V1/19.1$c_6$I1+L1+E1/16/19/5+H1/19/5$p_{4-5}^{3}c_4p_{4-}$

$_5$I1+L1+E1/16/19/5+H1/19/5$p_{4-5}^{4}c_4p_{4-5}$I1+L1+E1/16/19/5+H1/19/5$p_{4-5}c_4p_{4-2}p_{4-}$

$_5$I1+L1+E1/16/19/5+H1/19/5$p_{4-}$

$_5c_4$D1/19/5+P1+F1/19+G1/19/5+V1/19.1I1+L1+E1/16/19/5+H1/19/5$p_{4-5}c_4p_{4-}$

$_5$I1+L1+E1/16/19/5+H1/19/5$c_4$D1/19/5+P1+F1/19+G1/19/5+V1/19.1I1+L1+E1/16/19/5+H1/19/5p

$_{4-5}^{2}c_4p_{4-5}^{2}c_4p_{4-5}$I1+L1+E1/16/19/5+H1/19/5$c_4p_{4-5}$I1+L1+E1/16/19/5+H1/19/5$p_{4-5}^{2}c_4p_{4-}$

$_5$I1+L1+E1/16/19/5+H1/19/5$p_{4-5}^{4}c_4p_{4-5}$I1+L1+E1/16/19/5+H1/19/5$p_{4-5}c_4p_{4-}$

$_5$I1+L1+E1/16/19/5+H1/19/5$c_4p_{4-5}$D1/19/5+P1+F1/19+G1/19/5+V1/19.1$c_4p_{4-}$
$_5$I1+L1+E1/16/19/5+H1/19/5$^3c_4p_{4-5}$I1+L1+E1/16/19/5+H1/19/5$c_4p_{4-}$
$_5$I1+L1+E1/16/19/5+H1/19/5$c_4$D1/19/5+P1+F1/19+G1/19/5+V1/19.1I1+L1+E1/16/19/5+H1/19/5$^3$
$p_{4-5}$I1+L1+E1/16/19/5+H1/19/5$c_4p_{4-5}^4$I1+L1+E1/16/19/5+H1/19/5$p_{4-}$
$_5^3c_4$D1/19/5+P1+F1/19+G1/19/5+V1/19.1$p_{1-5}p_{4-5}c_4p_{4-}$
$_5$I1+L1+E1/16/19/5+H1/19/5$c_4$I1+L1+E1/16/19/5+H1/19/5$^2p_{4-5}c_4p_{4-}$
$_5^2$I1+L1+E1/16/19/5+H1/19/5$c_4p_{4-5}$I1+L1+E1/16/19/5+H1/19/5$c_4p_{4-}$
$_5^2$D1/19/5+P1+F1/19+G1/19/5+V1/19.1I1+L1+E1/16/19/5+H1/19/5$p_{4-}$
$_5$I1+L1+E1/16/19/5+H1/19/5$c_4p_{4-5}c_4p_{4-}$
$_5$I1+L1+E1/16/19/5+H1/19/5$c_4$I1+L1+E1/16/19/5+H1/19/5$^2p_{4-5}c_4p_{4-}$
$_5^2$I1+L1+E1/16/19/5+H1/19/5$c_4p_{4-5}$I1+L1+E1/16/19/5+H1/19/5$c_4p_{4-}$
$_5^2$D1/19/5+P1+F1/19+G1/19/5+V1/19.1I1+L1+E1/16/19/5+H1/19/5$p_{4-}$
$_5^2$D1/19/5+P1+F1/19+G1/19/5+V1/19.1I1+L1+E1/16/19/5+H1/19/5$^2p_{4-}$
$_5$I1+L1+E1/16/19/5+H1/19/5$c_4p_{4-5}^7c_4p_{4-5}^8c_4p_{4-5}^3c_4p_{4-5}$I1+L1+E1/16/19/5+H1/19/5$c_4p_{4-}$
$_5$I1+L1+E1/16/19/5+H1/19/5$p_{4-5}c_4p_{4-5}^2$I1+L1+E1/16/19/5+H1/19/5$c_4p_{4-}$
$_5^2$I1+L1+E1/16/19/5+H1/19/5$c_4p_{4-5}$I1+L1+E1/16/19/5+H1/19/5$c_5$I1+L1+E1/16/19/5+H1/19/5$p_{4-}$
$_5^{10}c_4p_{4-5}^8c_4p_{4-5}^3c_4p_{4-5}^2$I1+L1+E1/16/19/5+H1/19/5$p_{4-5}^5c_4p_{4-5}^3c_4p_{4-5}^8c_4p_{4-}$
$_5^6$I1+L1+E1/16/19/5+H1/19/5$c_4p_{4-5}^6$D1/19/5+P1+F1/19+G1/19/5+V1/19.1$c_6p_{4-}$
$_5$I1+L1+E1/16/19/5+H1/19/5$p_{4-5}^{13}c_4p_{4-5}^6$I1+L1+E1/16/19/5+H1/19/5$c_4p_{4-5}^8c_4p_{4-}$
$_5^2$I1+L1+E1/16/19/5+H1/19/5$p_{4-5}^6c_4p_{4-5}^5c_4p_{4-5}$I1+L1+E1/16/19/5+H1/19/5$p_{4-}$
$_5c_5$I1+L1+E1/16/19/5+H1/19/5$p_{4-5}^4c_4p_{4-5}^8c_4p_{4-5}^2$I1+L1+E1/16/19/5+H1/19/5$p_{4-5}^2c_4p_{4-}$
$_5^2$I1+L1+E1/16/19/5+H1/19/5$p_{4-5}^6c_4p_{4-5}^8c_4p_{4-5}^2$I1+L1+E1/16/19/5+H1/19/5$c_4p_{4-}$
$_5^2$I1+L1+E1/16/19/5+H1/19/5$p_{4-5}^2c_4p_{4-5}$I1+L1+E1/16/19/5+H1/19/5$p_{4-}$
$_5$I1+L1+E1/16/19/5+H1/19/5$^2p_{4-5}c_4p_{4-5}$I1+L1+E1/16/19/5+H1/19/5$c_4p_{4-}$
$_5$I1+L1+E1/16/19/5+H1/19/5$c_4p_{4-5}^3$I1+L1+E1/16/19/5+H1/19/5$p_{4-5}^4c_4p_{4-}$
$_5^4$I1+L1+E1/16/19/5+H1/19/5$p_{4-5}$I1+L1+E1/16/19/5+H1/19/5$c_4p_{4-5}^2$I1+L1+E1/16/19/5+H1/19/5$p_{4-}$
$_5$I1+L1+E1/16/19/5+H1/19/5$p_{4-5}^3c_4p_{4-5}^3$I1+L1+E1/16/19/5+H1/19/5$p_{4-}$
$_5$I1+L1+E1/16/19/5+H1/19/5$c_4p_{4-5}^4$I1+L1+E1/16/19/5+H1/19/5$c_4p_{4-}$
$_5^5c_5$I1+L1+E1/16/19/5+H1/19/5$p_{4-5}c_5$I1+L1+E1/16/19/5+H1/19/5$p_{4-}$
$_5^5c_5$I1+L1+E1/16/19/5+H1/19/5$p_{4-5}^4c_4p_{4-5}^4$I1+L1+E1/16/19/5+H1/19/5$p_{4-}$
$_5^2$I1+L1+E1/16/19/5+H1/19/5$p_{4-5}c_4p_{4-}$
$_5^3$I1+L1+E1/16/19/5+H1/19/5$c_5$I1+L1+E1/16/19/5+H1/19/5$^2p_{4-5}^2c_4p_{4-}$
$_5^2$I1+L1+E1/16/19/5+H1/19/5$p_{4-5}$I1+L1+E1/16/19/5+H1/19/5$p_{4-5}c_4p_{4-}$
$_5^3$I1+L1+E1/16/19/5+H1/19/5$p_{4-}$
$_5^2$I1+L1+E1/16/19/5+H1/19/5$c_5$I1+L1+E1/16/19/5+H1/19/5D1/19/5+P1+F1/19+G1/19/5+V1/19.1
$c_6p_{4-5}^2c_4p_{4-5}^5c_4p_{4-5}^5c_4p_{4-5}$I1+L1+E1/16/19/5+H1/19/5$^2p_{4-}$
$_5$I1+L1+E1/16/19/5+H1/19/5$c_5$I1+L1+E1/16/19/5+H1/19/5$c_5$I1+L1+E1/16/19/5+H1/19/5$p_{4-5}^2c_4p_{4-}$
$_5c_4p_{4-5}c_4c_6$I1+L1+E1/16/19/5+H1/19/5$c_4p_{4-5}^4$I1+L1+E1/16/19/5+H1/19/5$c_4p_{4-}$
$_5$I1+L1+E1/16/19/5+H1/19/5$p_{4-5}c_4p_{4-5}$I1+L1+E1/16/19/5+H1/19/5$p_{4-5}^2c_4p_{4-}$
$_5$I1+L1+E1/16/19/5+H1/19/5$p_{4-5}c_4p_{4-5}$I1+L1+E1/16/19/5+H1/19/5$p_{4-5}^2c_4p_{4-}$
$_5^4$D1/19/5+P1+F1/19+G1/19/5+V1/19.1$c_6$I1+L1+E1/16/19/5+H1/19/5$p_{4-5}c_4p_{4-}$
$_5$I1+L1+E1/16/19/5+H1/19/5$^4p_{4-5}p_{4-6}p_{3-5}p_{4-5}c_4p_{4-5}^2p_{4-6}p_{3-5}p_{4-5}c_4^2p_{4-}$
$_5^4$I1+L1+E1/16/19/5+H1/19/5$c_4p_{4-5}$I1+L1+E1/16/19/5+H1/19/5$p_{4-5}c_4p_{4-}$
$_5$I1+L1+E1/16/19/5+H1/19/5$p_{4-5}^2c_4p_{4-5}^2c_4p_{4-5}^2$I1+L1+E1/16/19/5+H1/19/5$^2p_{4-5}c_4p_{4-}$
$_5$I1+L1+E1/16/19/5+H1/19/5$^2p_{4-5}c_4p_{4-5}$I1+L1+E1/16/19/5+H1/19/5$^2p_{4-5}c_4p_{4-}$
$_5$I1+L1+E1/16/19/5+H1/19/5$p_{4-5}$I1+L1+E1/16/19/5+H1/19/5$p_{4-5}^2c_4p_{4-}$
$_5$I1+L1+E1/16/19/5+H1/19/5$^3p_{4-5}c_4p_{4-5}$I1+L1+E1/16/19/5+H1/19/5$p_{4-}$
$_5$I1+L1+E1/16/19/5+H1/19/5$^2p_{4-5}c_4p_{4-5}$I1+L1+E1/16/19/5+H1/19/5$p_{4-5}^2c_4p_{4-}$
$_5$I1+L1+E1/16/19/5+H1/19/5$^3p_{4-5}c_4p_{4-5}$I1+L1+E1/16/19/5+H1/19/5$p_{4-5}^2c_4p_{4-}$
$_5$I1+L1+E1/16/19/5+H1/19/5$^3p_{4-5}c_4p_{4-5}$I1+L1+E1/16/19/5+H1/19/5$p_{4-5}^2c_4p_{4-}$
$_5$I1+L1+E1/16/19/5+H1/19/5$^2p_{4-5}c_5$I1+L1+E1/16/19/5+H1/19/5$^2p_{4-5}^2c_4p_{4-}$

$_5$I1+L1+E1/16/19/5+H1/19/5$^2$p$_{4-5}$c$_5$I1+L1+E1/16/19/5+H1/19/5p$_{4-5}^{3}$c$_4$p$_{4-5}^{2}$c$_4$p$_{4-5}$c$_4$p$_{4-}$

$_5$I1+L1+E1/16/19/5+H1/19/5p$_{4-5}^{2}$c$_{-3}$I1+L1+E1/16/19/5+H1/19/5'p$_{-5--4}^{2}$c$_{-5}$p$_{-5--4}$c$_{-5}$p$_{-5--}$

$_4^3$I1+L1+E1/16/19/5+H1/19/5'$^2$p$_{-5--4}^{4}$c$_{-5}$p$_{-5--4}$c$_{-5}$p$_{-5--4}^{3}$I1+L1+E1/16/19/5+H1/19/5'$^2$p$_{-5--4}^{4}$c$_{-5}$p$_{-5--4}$c$_{-5}$p$_{-5--}$

$_4^2$I1+L1+E1/16/19/5+H1/19/5'p$_{-5--4}$I1+L1+E1/16/19/5+H1/19/5'p$_{-5--4}^{4}$I1+L1+E1/16/19/5+H1/19/5'p$_{-}$

$_{2--4}$I1+L1+E1/16/19/5+H1/19/5'p$_{-5--4}$c$_{-5}^{2}$I1+L1+E1/16/19/5+H1/19/5'p$_{-5--}$

$_4^2$I1+L1+E1/16/19/5+H1/19/5'p$_{-5--4}$I1+L1+E1/16/19/5+H1/19/5'p$_{-5--4}$I1+L1+E1/16/19/5+H1/19/5'c$_{-}$

$_5$p$_{-5--4}^{2}$I1+L1+E1/16/19/5+H1/19/5'p$_{-5--4}$c$_{-5}$I1+L1+E1/16/19/5+H1/19/5'p$_{-5--4}^{2}$c$_{-}$

$_5$I1+L1+E1/16/19/5+H1/19/5'$^3$p$_{-5--4}$c$_{-5}$p$_{-5--4}$I1+L1+E1/16/19/5+H1/19/5'p$_{-5--}$

$_4$I1+L1+E1/16/19/5+H1/19/5'p$_{-5--4}^{2}$c$_{-5}$p$_{-5--4}$I1+L1+E1/16/19/5+H1/19/5'$^2$p$_{-5--}$

$_4^2$I1+L1+E1/16/19/5+H1/19/5'c$_{-5}$I1+L1+E1/16/19/5+H1/19/5'c$_{-5}$I1+L1+E1/16/19/5+H1/19/5'c$_{-5}$p$_{-5--}$

$_4^2$c$_{-5}$p$_{-5--4}^{3}$c$_{-5}$p$_{-5--4}^{2}$c$_{-5}$p$_{-5--4}^{2}$c$_{-5}$p$_{-5--4}^{2}$c$_{-5}$p$_{-5--4}^{2}$I1+L1+E1/16/19/5+H1/19/5'p$_{-5--}$

$_4$I1+L1+E1/16/19/5+H1/19/5'p$_{-5--4}$c$_{-5}$I1+L1+E1/16/19/5+H1/19/5'p$_{-5--}$

$_4^2$I1+L1+E1/16/19/5+H1/19/5'c$_{-5}$I1+L1+E1/16/19/5+H1/19/5'$^3$p$_{-5--4}$c$_{-}$

$_5$I1+L1+E1/16/19/5+H1/19/5'$^2$p$_{-5--4}$I1+L1+E1/16/19/5+H1/19/5'c$_{-5}^{2}$I1+L1+E1/16/19/5+H1/19/5'c$_{-}$

$_5$p$_{-5--4}^{3}$c$_{-5}$p$_{-5--4}$c$_{-5}$p$_{-5--4}$I1+L1+E1/16/19/5+H1/19/5'p$_{-5--4}$I1+L1+E1/16/19/5+H1/19/5'p$_{-5--4}$c$_{-5}$p$_{-5--}$

$_4$I1+L1+E1/16/19/5+H1/19/5'p$_{-5--4}$c$_{-5}$p$_{-5--4}$c$_{-5}$I1+L1+E1/16/19/5+H1/19/5'$^2$p$_{-5--}$

$_4$I1+L1+E1/16/19/5+H1/19/5'p$_{-5--4}$I1+L1+E1/16/19/5+H1/19/5'p$_{-5--4}$c$_{-5}$p$_{-5--}$

$_4$I1+L1+E1/16/19/5+H1/19/5'p$_{-5--4}^{3}$c$_{-5}$I1+L1+E1/16/19/5+H1/19/5'p$_{-5--4}$c$_{-5}$p$_{-5--}$

$_4^3$I1+L1+E1/16/19/5+H1/19/5'$^2$c$_{-5}$p$_{-5--4}$I1+L1+E1/16/19/5+H1/19/5'p$_{-5--}$

$_4^2$I1+L1+E1/16/19/5+H1/19/5'$^3$c$_{-5}$p$_{-5--4}$I1+L1+E1/16/19/5+H1/19/5'p$_{-5--}$

$_4^3$I1+L1+E1/16/19/5+H1/19/5'$^2$c$_{-5}$p$_{-5--4}$I1+L1+E1/16/19/5+H1/19/5'p$_{-5--4}^{3}$c$_{-5}$p$_{-5--4}$c$_{-5}$p$_{-5--4}$c$_{-5}^{2}$p$_{-5--}$

$_4$I1+L1+E1/16/19/5+H1/19/5'p$_{-5--4}$I1+L1+E1/16/19/5+H1/19/5'p$_{-5--4}$c$_{-5}^{2}$p$_{-5--4}$c$_{-5}$p$_{-5--4}$c$_{-5}$p$_{-5--}$

$_4^2$I1+L1+E1/16/19/5+H1/19/5'p$_{-5--4}^{4}$c$_{-5}$p$_{-5--4}$c$_{-5}$p$_{-5--4}$I1+L1+E1/16/19/5+H1/19/5'p$_{-5--}$

$_4$I1+L1+E1/16/19/5+H1/19/5'p$_{-5--4}$I1+L1+E1/16/19/5+H1/19/5'p$_{-5--4}^{4}$c$_{-5}$p$_{-5--4}$c$_{-}$

$_4$D1/19/5+P1+F1/19+G1/19/5+V1/19.1'I1+L1+E1/16/19/5+H1/19/5'p$_{-5--}$

$_4$I1+L1+E1/16/19/5+H1/19/5'$^3$p$_{-5--4}^{4}$c$_{-5}$p$_{-5--4}$c$_{-5}$p$_{-5--4}$I1+L1+E1/16/19/5+H1/19/5'p$_{-5--}$

$_4$I1+L1+E1/16/19/5+H1/19/5'p$_{-5--4}$I1+L1+E1/16/19/5+H1/19/5'p$_{-5--4}^{2}$c$_{-5}$p$_{-5--4}^{3}$c$_{-5}$p$_{-5--4}$c$_{-5}$p$_{-5--}$

$_4$I1+L1+E1/16/19/5+H1/19/5'p$_{-5--4}$I1+L1+E1/16/19/5+H1/19/5'p$_{-5--4}$I1+L1+E1/16/19/5+H1/19/5'p$_{-5--}$

$_4^2$c$_{-5}$p$_{-5--4}^{3}$c$_{-5}$p$_{-5--4}$c$_{-5}$p$_{-5--4}$c$_{-5}$p$_{-5--4}$I1+L1+E1/16/19/5+H1/19/5'p$_{-5--4}$c$_{-5}$p$_{-5--4}$c$_{-}$

$_4$D1/19/5+P1+F1/19+G1/19/5+V1/19.1'c$_{-}$

$_4$D1/19/5+P1+F1/19+G1/19/5+V1/19.1'I1+L1+E1/16/19/5+H1/19/5'p$_{-5--4}$c$_{-5}$p$_{-5--}$

$_4^3$I1+L1+E1/16/19/5+H1/19/5'D1/19/5+P1+F1/19+G1/19/5+V1/19.1'p$_{-5--}$

$_4^2$I1+L1+E1/16/19/5+H1/19/5'c$_{-4}$D1/19/5+P1+F1/19+G1/19/5+V1/19.1'p$_{-5--}$

$_4^2$I1+L1+E1/16/19/5+H1/19/5'c$_{-5}^{2}$I1+L1+E1/16/19/5+H1/19/5'c$_{-5}$p$_{-5--4}^{2}$c$_{-5}$p$_{-5--}$

$_4^3$I1+L1+E1/16/19/5+H1/19/5'c$_{-5}$p$_{-5--4}$I1+L1+E1/16/19/5+H1/19/5'c$_{-5}$p$_{-5--}$

$_4$I1+L1+E1/16/19/5+H1/19/5'$^2$p$_{-5--4}^{2}$c$_{-5}$p$_{-5--4}$I1+L1+E1/16/19/5+H1/19/5'$^2$p$_{-5--4}$c$_{-5}$p$_{-5--}$

$_4$I1+L1+E1/16/19/5+H1/19/5'p$_{-5--4}$c$_{-5}$p$_{-5--4}^{2}$c$_{-5}$p$_{-5--4}^{3}$I1+L1+E1/16/19/5+H1/19/5'c$_{-}$

$_4$D1/19/5+P1+F1/19+G1/19/5+V1/19.1'p$_{-5--4}^{4}$I1+L1+E1/16/19/5+H1/19/5'c$_{-}$

$_4$D1/19/5+P1+F1/19+G1/19/5+V1/19.1'p$_{-5--4}^{2}$I1+L1+E1/16/19/5+H1/19/5'c$_{-}$

$_4$D1/19/5+P1+F1/19+G1/19/5+V1/19.1'p$_{-5--4}$I1+L1+E1/16/19/5+H1/19/5'c$_{-}$

$_4$D1/19/5+P1+F1/19+G1/19/5+V1/19.1'p$_{-5--4}^{2}$I1+L1+E1/16/19/5+H1/19/5'c$_{-5}$p$_{-5--}$

$_4$I1+L1+E1/16/19/5+H1/19/5'$^2$p$_{-5--4}$I1+L1+E1/16/19/5+H1/19/5'c$_{-5}$p$_{-5--}$

$_4$I1+L1+E1/16/19/5+H1/19/5'$^2$p$_{-5--4}$I1+L1+E1/16/19/5+H1/19/5'c$_{-}$

$_5$I1+L1+E1/16/19/5+H1/19/5'D1/19/5+P1+F1/19+G1/19/5+V1/19.1'p$_{-5--}$

$_4^2$I1+L1+E1/16/19/5+H1/19/5'c$_{-5}$p$_{-5--4}^{3}$I1+L1+E1/16/19/5+H1/19/5'c$_{-5}$p$_{-5--}$

$_4^3$I1+L1+E1/16/19/5+H1/19/5'c$_{-5}$p$_{-5--4}$I1+L1+E1/16/19/5+H1/19/5'c$_{-5}$p$_{-5--}$

$_4$I1+L1+E1/16/19/5+H1/19/5'c$_{-5}$p$_{-5--4}$I1+L1+E1/16/19/5+H1/19/5'c$_{-5}$p$_{-5--}$

$_4^3$I1+L1+E1/16/19/5+H1/19/5'c$_{-5}$p$_{-5--4}$I1+L1+E1/16/19/5+H1/19/5'p$_{-5--}$

$_4$I1+L1+E1/16/19/5+H1/19/5'c$_{-5}$p$_{-5--4}$I1+L1+E1/16/19/5+H1/19/5'$^3$c$_{-5}$p$_{-5--}$

$_4^2$I1+L1+E1/16/19/5+H1/19/5'c$_{-5}$p$_{-5--4}$I1+L1+E1/16/19/5+H1/19/5'c$_{-5}$p$_{-5--}$

$_4$I1+L1+E1/16/19/5+H1/19/5'p$_{-5--4}$I1+L1+E1/16/19/5+H1/19/5'c$_{-5}$p$_{-5--}$

43I1+L1+E1/16/19/5+H1/19/5'c-4D1/19/5+P1+F1/19+G1/19/5+V1/19.1'p-5--
43I1+L1+E1/16/19/5+H1/19/5'c-5p-5--4I1+L1+E1/16/19/5+H1/19/5'p-5--
42I1+L1+E1/16/19/5+H1/19/5'c-
5I1+L1+E1/16/19/5+H1/19/5'D1/19/5+P1+F1/19+G1/19/5+V1/19.1'p-5--43c-
4D1/19/5+P1+F1/19+G1/19/5+V1/19.1'p-5--43I1+L1+E1/16/19/5+H1/19/5'c-5p-5--
4I1+L1+E1/16/19/5+H1/19/5'2c-5p-5--43I1+L1+E1/16/19/5+H1/19/5'p-5--4c-5p-5--
43I1+L1+E1/16/19/5+H1/19/5'c-5p-5--4I1+L1+E1/16/19/5+H1/19/5'p-5--
4I1+L1+E1/16/19/5+H1/19/5'c-5p-5--43I1+L1+E1/16/19/5+H1/19/5'c-5p-5--
4I1+L1+E1/16/19/5+H1/19/5'p-5--4I1+L1+E1/16/19/5+H1/19/5'c-5p-5--
4I1+L1+E1/16/19/5+H1/19/5'p-5--42I1+L1+E1/16/19/5+H1/19/5'c-5p-5--
4I1+L1+E1/16/19/5+H1/19/5'c-5p-5--4I1+L1+E1/16/19/5+H1/19/5'p-5--
4I1+L1+E1/16/19/5+H1/19/5'c-
4D1/19/5+P1+F1/19+G1/19/5+V1/19.1'I1+L1+E1/16/19/5+H1/19/5'p-5--
4I1+L1+E1/16/19/5+H1/19/5'c-5p-5--4I1+L1+E1/16/19/5+H1/19/5'p-5--
4I1+L1+E1/16/19/5+H1/19/5'c-5p-5--4I1+L1+E1/16/19/5+H1/19/5'p-5--
4I1+L1+E1/16/19/5+H1/19/5'c-5p-5--4I1+L1+E1/16/19/5+H1/19/5'c-5p-5--
44I1+L1+E1/16/19/5+H1/19/5'c-5p-5--43c-5p-5--42c-5p-5--42c-5p-5--4I1+L1+E1/16/19/5+H1/19/5'c-5p-5--42c-
5p-5--42c-5p-5--47c-5p-5--42c-5p-5--4c-5p-5--42I1+L1+E1/16/19/5+H1/19/5'c-3I1+L1+E1/16/19/5+H1/19/5'c-
4D1/19/5+P1+F1/19+G1/19/5+V1/19.1'p-5--42c-5p-5--4I1+L1+E1/16/19/5+H1/19/5'c-
4D1/19/5+P1+F1/19+G1/19/5+V1/19.1'p-5--42c-5p-5--4I1+L1+E1/16/19/5+H1/19/5'c-
4D1/19/5+P1+F1/19+G1/19/5+V1/19.1'p-5--4I1+L1+E1/16/19/5+H1/19/5'c-
4D1/19/5+P1+F1/19+G1/19/5+V1/19.1'p-5--42c-5p-5--4I1+L1+E1/16/19/5+H1/19/5'c-
4D1/19/5+P1+F1/19+G1/19/5+V1/19.1'p-5--4I1+L1+E1/16/19/5+H1/19/5'c-
4D1/19/5+P1+F1/19+G1/19/5+V1/19.1'p-5--42c-5p-5--4I1+L1+E1/16/19/5+H1/19/5'c-
4D1/19/5+P1+F1/19+G1/19/5+V1/19.1'p-5--4c-5p-5--44I1+L1+E1/16/19/5+H1/19/5'c-
5I1+L1+E1/16/19/5+H1/19/5'p-5--4I1+L1+E1/16/19/5+H1/19/5'c-5p-5--4c-5p-5--
44I1+L1+E1/16/19/5+H1/19/5'2p-5--4I1+L1+E1/16/19/5+H1/19/5'c-5p-5--4c-5p-5--42c-5p-5--42c-5p-5--
43I1+L1+E1/16/19/5+H1/19/5'2p-5--4I1+L1+E1/16/19/5+H1/19/5'c-5p-5--
44I1+L1+E1/16/19/5+H1/19/5'c-5p-5--44I1+L1+E1/16/19/5+H1/19/5'c-5p-5--
44I1+L1+E1/16/19/5+H1/19/5'p-5--42I1+L1+E1/16/19/5+H1/19/5'c-5p-5--
4I1+L1+E1/16/19/5+H1/19/5'p-5--4I1+L1+E1/16/19/5+H1/19/5'c-5p-5--
4I1+L1+E1/16/19/5+H1/19/5'2p-5--
4I1+L1+E1/16/19/5+H1/19/5'2D1/19/5+P1+F1/19+G1/19/5+V1/19.1'p-5--
42I1+L1+E1/16/19/5+H1/19/5'c-5p-5--43I1+L1+E1/16/19/5+H1/19/5'p-5--
42I1+L1+E1/16/19/5+H1/19/5'2c-5p-5--42I1+L1+E1/16/19/5+H1/19/5'c-5p-5--
4I1+L1+E1/16/19/5+H1/19/5'p-5--4I1+L1+E1/16/19/5+H1/19/5'p-5--44I1+L1+E1/16/19/5+H1/19/5'c-
5p-5--4I1+L1+E1/16/19/5+H1/19/5'p-5--4I1+L1+E1/16/19/5+H1/19/5'3c-5p-5--
4I1+L1+E1/16/19/5+H1/19/5'p-5--42I1+L1+E1/16/19/5+H1/19/5'c-5p-5--
4I1+L1+E1/16/19/5+H1/19/5'p-5--4I1+L1+E1/16/19/5+H1/19/5'2c-5p-5--
4I1+L1+E1/16/19/5+H1/19/5'p-5--42I1+L1+E1/16/19/5+H1/19/5'c-5p-5--
4I1+L1+E1/16/19/5+H1/19/5'p-5--4c-5p-5--4I1+L1+E1/16/19/5+H1/19/5'c-5p-5--42c-5p-5--
44I1+L1+E1/16/19/5+H1/19/5'c-5p-5--42c-5p-5--44I1+L1+E1/16/19/5+H1/19/5'c-
5D1/19/5+P1+F1/19+G1/19/5+V1/19.1'p-5--42I1+L1+E1/16/19/5+H1/19/5'c-5p-5--
42I1+L1+E1/16/19/5+H1/19/5'p-5--44I1+L1+E1/16/19/5+H1/19/5'c-
5D1/19/5+P1+F1/19+G1/19/5+V1/19.1'p-5--42I1+L1+E1/16/19/5+H1/19/5'c-5p-5--
42I1+L1+E1/16/19/5+H1/19/5'p-5--44I1+L1+E1/16/19/5+H1/19/5'c-
5D1/19/5+P1+F1/19+G1/19/5+V1/19.1'p-5--42I1+L1+E1/16/19/5+H1/19/5'c-5p-5--
42I1+L1+E1/16/19/5+H1/19/5'c-5p-5--4c-5p-5--42c-5p-5--44I1+L1+E1/16/19/5+H1/19/5'c-5p-5--
46I1+L1+E1/16/19/5+H1/19/5'c-5I1+L1+E1/16/19/5+H1/19/5'p-5--4I1+L1+E1/16/19/5+H1/19/5'c-5p-
5--42c-5p-5--42c-5p-5--4c-5p-5--4I1+L1+E1/16/19/5+H1/19/5'2c-5p-5--42I1+L1+E1/16/19/5+H1/19/5'c-5p-5--
42D1/19/5+P1+F1/19+G1/19/5+V1/19.1'p-5--4I1+L1+E1/16/19/5+H1/19/5'c-5p-5--

44I1+L1+E1/16/19/5+H1/19/5'c-5p-5--44I1+L1+E1/16/19/5+H1/19/5'c-5p-5--
44I1+L1+E1/16/19/5+H1/19/5'c-5p-5--43I1+L1+E1/16/19/5+H1/19/5'c-5p-5--
42I1+L1+E1/16/19/5+H1/19/5'c-5p-5--4I1+L1+E1/16/19/5+H1/19/5'4p-5--
43I1+L1+E1/16/19/5+H1/19/5'c-5p-5--43I1+L1+E1/16/19/5+H1/19/5'p-5--
42I1+L1+E1/16/19/5+H1/19/5'c-5p-5--43I1+L1+E1/16/19/5+H1/19/5'c-5p-5--
43I1+L1+E1/16/19/5+H1/19/5'c-5p-5--42I1+L1+E1/16/19/5+H1/19/5'c-5p-5--
4I1+L1+E1/16/19/5+H1/19/5'p-5--42I1+L1+E1/16/19/5+H1/19/5'c-5p-5--
4I1+L1+E1/16/19/5+H1/19/5'p-5--42I1+L1+E1/16/19/5+H1/19/5'c-5p-5--
4I1+L1+E1/16/19/5+H1/19/5'p-5--42I1+L1+E1/16/19/5+H1/19/5'c-5p-5--
44I1+L1+E1/16/19/5+H1/19/5'c-5p-5--42I1+L1+E1/16/19/5+H1/19/5'c-5p-5--
4I1+L1+E1/16/19/5+H1/19/5'c-5p-5--44I1+L1+E1/16/19/5+H1/19/5'c-5p-5--
42I1+L1+E1/16/19/5+H1/19/5'c-5p-5--4I1+L1+E1/16/19/5+H1/19/5'c-5p-5--
4I1+L1+E1/16/19/5+H1/19/5'p-5--42I1+L1+E1/16/19/5+H1/19/5'c-5p-5--
42I1+L1+E1/16/19/5+H1/19/5'c-5p-5--43I1+L1+E1/16/19/5+H1/19/5'2c-5p-5--
4I1+L1+E1/16/19/5+H1/19/5'p-5--4I1+L1+E1/16/19/5+H1/19/5'3c-5p-5--4c-5p-5--
4I1+L1+E1/16/19/5+H1/19/5'p-5--4I1+L1+E1/16/19/5+H1/19/5'3c-5p-5--
43I1+L1+E1/16/19/5+H1/19/5'c-5p-5--4I1+L1+E1/16/19/5+H1/19/5'c-5p-5--
42I1+L1+E1/16/19/5+H1/19/5'c-5p-5--4I1+L1+E1/16/19/5+H1/19/5'p-5--4c-5p-5--
4I1+L1+E1/16/19/5+H1/19/5'p-5--4c-5p-5--4I1+L1+E1/16/19/5+H1/19/5'c-5p-5--42c-5p-5--
43I1+L1+E1/16/19/5+H1/19/5'c-5p-5--4c-5p-5--42c-5p-5--43c-5I1+L1+E1/16/19/5+H1/19/5'p-5--43c-5p-5--
42I1+L1+E1/16/19/5+H1/19/5'c-5I1+L1+E1/16/19/5+H1/19/5'p-5--42I1+L1+E1/16/19/5+H1/19/5'c-
5p-5--42I1+L1+E1/16/19/5+H1/19/5'c-5p-5--4I1+L1+E1/16/19/5+H1/19/5'2c-5p-5--
4I1+L1+E1/16/19/5+H1/19/5'2c-5p-5--4I1+L1+E1/16/19/5+H1/19/5'2c-5p-5--
4I1+L1+E1/16/19/5+H1/19/5'2c-5p-5--4I1+L1+E1/16/19/5+H1/19/5'2c-5p-5--
4I1+L1+E1/16/19/5+H1/19/5'2c-5p-5--4I1+L1+E1/16/19/5+H1/19/5'2c-5p-5--
4I1+L1+E1/16/19/5+H1/19/5'2c-5p-5--4I1+L1+E1/16/19/5+H1/19/5'2c-5p-5--
4I1+L1+E1/16/19/5+H1/19/5'2c-5p-5--4I1+L1+E1/16/19/5+H1/19/5'2c-5p-5--
4I1+L1+E1/16/19/5+H1/19/5'2c-5p-5--4I1+L1+E1/16/19/5+H1/19/5'2c-5p-5--
4I1+L1+E1/16/19/5+H1/19/5'2c-5p-5--4I1+L1+E1/16/19/5+H1/19/5'2c-5p-5--
4I1+L1+E1/16/19/5+H1/19/5'2c-5p-5--4I1+L1+E1/16/19/5+H1/19/5'2c-5p-5--
4I1+L1+E1/16/19/5+H1/19/5'2c-5p-5--4I1+L1+E1/16/19/5+H1/19/5'2c-5p-5--
4I1+L1+E1/16/19/5+H1/19/5'2c-5p-5--4I1+L1+E1/16/19/5+H1/19/5'2c-5p-5--
4I1+L1+E1/16/19/5+H1/19/5'2c-5p-5--4I1+L1+E1/16/19/5+H1/19/5'2c-5p-5--
4I1+L1+E1/16/19/5+H1/19/5'2c-5p-5--4I1+L1+E1/16/19/5+H1/19/5'2c-5p-5--
4I1+L1+E1/16/19/5+H1/19/5'2c-5p-5--4I1+L1+E1/16/19/5+H1/19/5'2c-5p-5--
4I1+L1+E1/16/19/5+H1/19/5'2c-5p-5--44I1+L1+E1/16/19/5+H1/19/5'c-5p-5--
44I1+L1+E1/16/19/5+H1/19/5'c-5p-5--4I1+L1+E1/16/19/5+H1/19/5'c-5I1+L1+E1/16/19/5+H1/19/5'p-
5--43I1+L1+E1/16/19/5+H1/19/5'c-5p-5--44I1+L1+E1/16/19/5+H1/19/5'c-52p-5--42c-5p-5--
44I1+L1+E1/16/19/5+H1/19/5'c-5p-5--42I1+L1+E1/16/19/5+H1/19/5'2c-5p-5--
44I1+L1+E1/16/19/5+H1/19/5'c-5p-5--4I1+L1+E1/16/19/5+H1/19/5'c-5I1+L1+E1/16/19/5+H1/19/5'p-
5--43I1+L1+E1/16/19/5+H1/19/5'c-5p-5--43I1+L1+E1/16/19/5+H1/19/5'2c-5p-5--
42I1+L1+E1/16/19/5+H1/19/5'c-5p-5--42I1+L1+E1/16/19/5+H1/19/5'c-5p-5--
4I1+L1+E1/16/19/5+H1/19/5'p-5--4c-5p-5--42c-4D1/19/5+P1+F1/19+G1/19/5+V1/19.1'p-5--
4I1+L1+E1/16/19/5+H1/19/5'c-3p-5--4c-52p-5--4I1+L1+E1/16/19/5+H1/19/5'p-5--
43I1+L1+E1/16/19/5+H1/19/5'p-5--4I1+L1+E1/16/19/5+H1/19/5'2c-5p-5--
42I1+L1+E1/16/19/5+H1/19/5'c-5p-5--4I1+L1+E1/16/19/5+H1/19/5'p-5--4c-5p-5--42c-
4D1/19/5+P1+F1/19+G1/19/5+V1/19.1'p-5--4c-5p-5--42c-52p-5--4I1+L1+E1/16/19/5+H1/19/5'p-5--
43I1+L1+E1/16/19/5+H1/19/5'p-5--4I1+L1+E1/16/19/5+H1/19/5'2c-5p-5--
43I1+L1+E1/16/19/5+H1/19/5'p-5--4I1+L1+E1/16/19/5+H1/19/5'c-5p-5--
4I1+L1+E1/16/19/5+H1/19/5'p-5--4I1+L1+E1/16/19/5+H1/19/5'c-5p-5--44c-5p-5--4c-5p-5--42c-5p-5--42c-5p-5--
44c-5p-5--44c-5p-5--44c-5I1+L1+E1/16/19/5+H1/19/5'3c-5p-5--4I1+L1+E1/16/19/5+H1/19/5'2p-5--44c-

$_{4}$D1/19/5+P1+F1/19+G1/19/5+V1/19.1'p$_{-5--4}^{3}$I1+L1+E1/16/19/5+H1/19/5'c$_{-5}$p$_{-5--}$

$_{4}^{3}$I1+L1+E1/16/19/5+H1/19/5'$^{2}$c$_{-5}$p$_{-5--4}$I1+L1+E1/16/19/5+H1/19/5'p$_{-5--4}^{2}$c$_{-5}$p$_{-5--4}^{2}$c$_{-5}$p$_{-5--}$

$_{4}$D1/19/5+P1+F1/19+G1/19/5+V1/19.1'p$_{-5--4}$I1+L1+E1/16/19/5+H1/19/5'$^{2}$c$_{-5}$p$_{-5--}$

$_{4}$D1/19/5+P1+F1/19+G1/19/5+V1/19.1'p$_{-5--4}$I1+L1+E1/16/19/5+H1/19/5'$^{2}$p$_{-5--}$

$_{4}^{2}$I1+L1+E1/16/19/5+H1/19/5'c$_{-5}$p$_{-5--4}$I1+L1+E1/16/19/5+H1/19/5'c$_{-5}$p$_{-5--4}^{4}$c$_{-5}$p$_{-5--}$

$_{4}$I1+L1+E1/16/19/5+H1/19/5'p$_{-5--4}$I1+L1+E1/16/19/5+H1/19/5'p$_{-5--4}$I1+L1+E1/16/19/5+H1/19/5'p$_{-5--}$

$_{4}$I1+L1+E1/16/19/5+H1/19/5'$^{2}$c$_{-5}$p$_{-5--4}$I1+L1+E1/16/19/5+H1/19/5'p$_{-5--4}^{2}$c$_{-5}$p$_{-5--}$

$_{4}$I1+L1+E1/16/19/5+H1/19/5'p$_{-5--4}$I1+L1+E1/16/19/5+H1/19/5'p$_{-5--4}$I1+L1+E1/16/19/5+H1/19/5'p$_{-5--}$

$_{4}$I1+L1+E1/16/19/5+H1/19/5'c$_{-5}$p$_{-5--4}$D1/19/5+P1+F1/19+G1/19/5+V1/19.1'p$_{-5--}$

$_{4}$I1+L1+E1/16/19/5+H1/19/5'$^{2}$c$_{-5}$p$_{-5--4}^{2}$c$_{-5}$p$_{-5--4}$I1+L1+E1/16/19/5+H1/19/5'p$_{-5--}$

$_{4}$I1+L1+E1/16/19/5+H1/19/5'p$_{-5--4}$I1+L1+E1/16/19/5+H1/19/5'p$_{-5--4}^{2}$I1+L1+E1/16/19/5+H1/19/5'c$_{-}$

$_{5}$p$_{-5--4}^{4}$I1+L1+E1/16/19/5+H1/19/5'c$_{-5}$p$_{-5--4}^{4}$I1+L1+E1/16/19/5+H1/19/5'p$_{-5--}$

$_{4}$I1+L1+E1/16/19/5+H1/19/5'c$_{-5}$p$_{-5--4}$I1+L1+E1/16/19/5+H1/19/5'c$_{-5}$p$_{-5--}$

$_{4}^{3}$I1+L1+E1/16/19/5+H1/19/5'c$_{-5}$D1/19/5+P1+F1/19+G1/19/5+V1/19.1'p$_{-5--4}^{2}$c$_{-5}$p$_{-5--}$

$_{4}$I1+L1+E1/16/19/5+H1/19/5'c$_{-5}$p$_{-5--4}$I1+L1+E1/16/19/5+H1/19/5'$^{2}$p$_{-5--}$

$_{4}$I1+L1+E1/16/19/5+H1/19/5'c$_{-5}$p$_{-5--4}$I1+L1+E1/16/19/5+H1/19/5'c$_{-5}$p$_{-5--4}^{2}$c$_{-5}$p$_{-5--}$

$_{4}^{3}$I1+L1+E1/16/19/5+H1/19/5'c$_{-5}$p$_{-5--4}^{2}$I1+L1+E1/16/19/5+H1/19/5'c$_{-}$

$_{3}$I1+L1+E1/16/19/5+H1/19/5'D1/19/5+P1+F1/19+G1/19/5+V1/19.1'p$_{-5--}$

$_{4}$I1+L1+E1/16/19/5+H1/19/5'c$_{-5}$p$_{-5--4}^{2}$I1+L1+E1/16/19/5+H1/19/5'c$_{-5}$p$_{-5--}$

$_{4}$I1+L1+E1/16/19/5+H1/19/5'p$_{-5--4}^{2}$I1+L1+E1/16/19/5+H1/19/5'p$_{-5--4}^{3}$I1+L1+E1/16/19/5+H1/19/5'c$_{-}$

$_{5}$p$_{-5--4}^{3}$I1+L1+E1/16/19/5+H1/19/5'c$_{-5}$D1/19/5+P1+F1/19+G1/19/5+V1/19.1'p$_{-5--}$

$_{4}$I1+L1+E1/16/19/5+H1/19/5'c$_{-5}$p$_{-5--4}^{2}$c$_{-5}$p$_{-5--4}^{2}$D1/19/5+P1+F1/19+G1/19/5+V1/19.1'p$_{-5--}$

$_{4}$I1+L1+E1/16/19/5+H1/19/5'c$_{-5}$p$_{-5--4}^{3}$I1+L1+E1/16/19/5+H1/19/5'p$_{-5--}$

$_{4}^{3}$I1+L1+E1/16/19/5+H1/19/5'c$_{-3}$p$_{-5--4}$I1+L1+E1/16/19/5+H1/19/5'c$_{-5}$p$_{-5--}$

$_{4}^{3}$I1+L1+E1/16/19/5+H1/19/5'c$_{-5}$p$_{-5--4}$I1+L1+E1/16/19/5+H1/19/5'c$_{-5}^{2}$p$_{-5--}$

$_{4}^{2}$I1+L1+E1/16/19/5+H1/19/5'c$_{-5}$D1/19/5+P1+F1/19+G1/19/5+V1/19.1'p$_{-5--4}^{2}$c$_{-5}$p$_{-5--4}^{4}$c$_{-5}$p$_{-5--}$

$_{4}^{4}$I1+L1+E1/16/19/5+H1/19/5'c$_{-5}$p$_{-5--4}^{4}$c$_{-}$

$_{4}$D1/19/5+P1+F1/19+G1/19/5+V1/19.1'I1+L1+E1/16/19/5+H1/19/5'p$_{-5--}$

$_{4}^{2}$I1+L1+E1/16/19/5+H1/19/5'c$_{-5}$p$_{-5--4}^{2}$I1+L1+E1/16/19/5+H1/19/5'c$_{-5}$p$_{-5--}$

$_{4}^{4}$I1+L1+E1/16/19/5+H1/19/5'c$_{-5}$p$_{-5--4}^{2}$I1+L1+E1/16/19/5+H1/19/5'c$_{-5}$p$_{-5--4}^{2}$c$_{-5}$p$_{-5--}$

$_{4}^{2}$D1/19/5+P1+F1/19+G1/19/5+V1/19.1'p$_{-5--4}$I1+L1+E1/16/19/5+H1/19/5'c$_{-5}$p$_{-5--}$

$_{4}^{3}$I1+L1+E1/16/19/5+H1/19/5'p$_{-5--4}^{3}$I1+L1+E1/16/19/5+H1/19/5'c$_{-5}$p$_{-5--}$

$_{4}^{3}$I1+L1+E1/16/19/5+H1/19/5'c$_{-5}$p$_{-5--4}$I1+L1+E1/16/19/5+H1/19/5'c$_{-5}$p$_{-5--}$

$_{4}^{3}$I1+L1+E1/16/19/5+H1/19/5'c$_{-5}$p$_{-5--4}^{2}$c$_{-5}$p$_{-5--4}^{4}$I1+L1+E1/16/19/5+H1/19/5'c$_{-5}$p$_{-5--}$

$_{4}$I1+L1+E1/16/19/5+H1/19/5'c$_{-5}$I1+L1+E1/16/19/5+H1/19/5'c$_{-5}$p$_{-5--4}^{2}$I1+L1+E1/16/19/5+H1/19/5'c$_{-}$

$_{5}$p$_{-5--4}^{2}$I1+L1+E1/16/19/5+H1/19/5'c$_{-5}$p$_{-5--4}^{2}$I1+L1+E1/16/19/5+H1/19/5'c$_{-5}$p$_{-5--4}^{2}$c$_{-}$

$_{5}$D1/19/5+P1+F1/19+G1/19/5+V1/19.1'p$_{-5--4}^{2}$I1+L1+E1/16/19/5+H1/19/5'p$_{-5--4}$c$_{-5}$p$_{-5--}$

$_{4}$I1+L1+E1/16/19/5+H1/19/5'p$_{-5--4}$I1+L1+E1/16/19/5+H1/19/5'p$_{-3--2}$p$_{-5--}$

$_{4}^{2}$I1+L1+E1/16/19/5+H1/19/5'p$_{-5--4}^{2}$I1+L1+E1/16/19/5+H1/19/5'p$_{-3--2}$p$_{-5--}$

$_{4}^{2}$I1+L1+E1/16/19/5+H1/19/5'p$_{-5--4}^{2}$I1+L1+E1/16/19/5+H1/19/5'p$_{-5--4}^{3}$c$_{-}$

$_{5}$I1+L1+E1/16/19/5+H1/19/5'c$_{-5}$I1+L1+E1/16/19/5+H1/19/5'p$_{-5--4}^{2}$c$_{-5}$p$_{-5--4}$c$_{-5}$p$_{-5--}$

$_{4}^{5}$I1+L1+E1/16/19/5+H1/19/5'c$_{-5}$p$_{-5--4}^{2}$I1+L1+E1/16/19/5+H1/19/5'c$_{-5}$p$_{-5--}$

$_{4}^{2}$I1+L1+E1/16/19/5+H1/19/5'p$_{-5--3}$D1/19/5+P1+F1/19+G1/19/5+V1/19.1'c$_{-5}$p$_{-5--}$

$_{4}^{2}$I1+L1+E1/16/19/5+H1/19/5'c$_{-5}$p$_{-5--4}$I1+L1+E1/16/19/5+H1/19/5'p$_{-5--4}^{3}$c$_{-5}$p$_{-5--}$

$_{4}$I1+L1+E1/16/19/5+H1/19/5'p$_{-5--4}$I1+L1+E1/16/19/5+H1/19/5'p$_{-4--}$

$_{2}$D1/19/5+P1+F1/19+G1/19/5+V1/19.1'c$_{-5}$p$_{-5--4}$c$_{-5}$I1+L1+E1/16/19/5+H1/19/5'$^{2}$p$_{-5--}$

$_{4}$I1+L1+E1/16/19/5+H1/19/5'c$_{-5}$p$_{-5--4}^{2}$I1+L1+E1/16/19/5+H1/19/5'p$_{-5--}$

$_{4}$I1+L1+E1/16/19/5+H1/19/5'$^{2}$p$_{-3--5}$p$_{-5--4}$I1+L1+E1/16/19/5+H1/19/5'p$_{-5--4}^{4}$p$_{-5--}$

$_{2}$D1/19/5+P1+F1/19+G1/19/5+V1/19.1'p$_{-5--4}^{2}$c$_{-5}$p$_{-5--4}$c$_{-5}$I1+L1+E1/16/19/5+H1/19/5'$^{2}$p$_{-5--}$

$_{4}$I1+L1+E1/16/19/5+H1/19/5'c$_{-5}$p$_{-5--4}$c$_{-5}$I1+L1+E1/16/19/5+H1/19/5'$^{2}$p$_{-5--}$

$_{4}$I1+L1+E1/16/19/5+H1/19/5'c$_{-5}$p$_{-5--4}^{4}$c$_{-5}$p$_{-5--4}^{2}$c$_{-5}$p$_{-5--4}$c$_{-5}$p$_{-5--4}^{2}$c$_{-5}$p$_{-5--4}^{2}$c$_{-5}$p$_{-5--4}$c$_{-5}$p$_{-5--4}^{2}$c$_{-5}$p$_{-5--4}$c$_{-5}$p$_{-5--}$

$_4$I1+L1+E1/16/19/5+H1/19/5'p$_{-5--4}$c$_{-5}$p$_{-5--4}^{4}$c$_{-5}$p$_{-5--4}$c$_{-5}$p$_{-5--4}^{3}$c$_{-5}$p$_{-5--4}$c$_{-5}$p$_{-5--}$

$_4$I1+L1+E1/16/19/5+H1/19/5'p$_{-5--4}$c$_{-5}$p$_{-5--4}^{4}$c$_{-5}$p$_{-5--4}$c$_{-5}$p$_{-5--4}^{4}$c$_{-5}$p$_{-5--4}^{3}$c$_{-5}^{2}$p$_{-5--}$

$_4$I1+L1+E1/16/19/5+H1/19/5'$^{2}$c$_{-5}$p$_{-5--4}^{4}$c$_{-5}$p$_{-5--4}$I1+L1+E1/16/19/5+H1/19/5'$^{2}$c$_{-5}$p$_{-5--4}^{4}$c$_{-5}$p$_{-5--}$

$_4^{3}$I1+L1+E1/16/19/5+H1/19/5'c$_{-2}$p$_{-2--4}$p$_{-5--4}^{2}$I1+L1+E1/16/19/5+H1/19/5'p$_{-5--4}$c$_{-5}$p$_{-5--4}^{5}$c$_{-5}$p$_{-5--}$

$_4^{3}$I1+L1+E1/16/19/5+H1/19/5'c$_{-5}$p$_{-5--4}^{3}$I1+L1+E1/16/19/5+H1/19/5'c$_{-5}$p$_{-5--}$

$_4^{4}$I1+L1+E1/16/19/5+H1/19/5'c$_{-1}$p$_{-1--4}$c$_{-5}$p$_{-5--4}$I1+L1+E1/16/19/5+H1/19/5'p$_{-5--}$

$_4$I1+L1+E1/16/19/5+H1/19/5'p$_{-5--4}$c$_{-5}$p$_{-5--4}^{2}$I1+L1+E1/16/19/5+H1/19/5'c$_{-5}$p$_{-5--}$

$_4^{2}$I1+L1+E1/16/19/5+H1/19/5'p$_{-5--4}^{2}$I1+L1+E1/16/19/5+H1/19/5'c$_{-5}$p$_{-5--}$

$_4$I1+L1+E1/16/19/5+H1/19/5'p$_{-5--4}^{3}$I1+L1+E1/16/19/5+H1/19/5'c-

$_4$D1/19/5+P1+F1/19+G1/19/5+V1/19.1'p$_{-5--4}^{3}$I1+L1+E1/16/19/5+H1/19/5'c$_{-5}$p$_{-5--4}$c$_{-5}$p$_{-5--}$

$_4^{2}$I1+L1+E1/16/19/5+H1/19/5'$^{2}$c$_{-5}$p$_{-5--4}$I1+L1+E1/16/19/5+H1/19/5'c$_{-5}$p$_{-5--}$

$_4^{4}$I1+L1+E1/16/19/5+H1/19/5'c$_{-5}$p$_{-5--4}^{3}$I1+L1+E1/16/19/5+H1/19/5'c$_{-5}$p$_{-5--4}^{2}$c$_{-5}$p$_{-5--}$

$_4^{3}$I1+L1+E1/16/19/5+H1/19/5'c$_{-5}$p$_{-5--4}^{2}$I1+L1+E1/16/19/5+H1/19/5'c$_{-5}$p$_{-5--}$

$_4^{3}$I1+L1+E1/16/19/5+H1/19/5'p$_{-5--4}^{2}$c$_{-5}$p$_{-5--4}^{2}$I1+L1+E1/16/19/5+H1/19/5'c$_{-5}$p$_{-5--}$

$_4^{2}$I1+L1+E1/16/19/5+H1/19/5'c$_{-5}$p$_{-5--4}^{5}$I1+L1+E1/16/19/5+H1/19/5'$^{2}$c$_{-5}$p$_{-5--}$

$_4^{5}$I1+L1+E1/16/19/5+H1/19/5'$^{2}$c$_{-5}$p$_{-5--4}^{2}$I1+L1+E1/16/19/5+H1/19/5'$^{2}$c$_{-5}$p$_{-5--}$

$_4^{4}$I1+L1+E1/16/19/5+H1/19/5'c$_{-5}$p$_{-5--4}^{2}$I1+L1+E1/16/19/5+H1/19/5'$^{2}$c$_{-5}$p$_{-5--}$

$_4$I1+L1+E1/16/19/5+H1/19/5'p$_{-5--4}$I1+L1+E1/16/19/5+H1/19/5'c$_{-5}$p$_{-5--}$

$_4^{2}$I1+L1+E1/16/19/5+H1/19/5'c$_{-5}$p$_{-5--4}^{3}$I1+L1+E1/16/19/5+H1/19/5'c$_{-5}$p$_{-5--}$

$_4^{2}$I1+L1+E1/16/19/5+H1/19/5'c$_{-5}$p$_{-5--4}^{4}$c$_{-5}$p$_{-5--4}^{2}$I1+L1+E1/16/19/5+H1/19/5'c$_{-5}$p$_{-5--}$

$_4^{2}$I1+L1+E1/16/19/5+H1/19/5'c$_{-5}$p$_{-5--4}^{4}$c$_{-5}$p$_{-5--4}^{3}$c$_{-5}$p$_{-5--4}^{3}$c$_{-5}^{2}$p$_{-5--4}$I1+L1+E1/16/19/5+H1/19/5'c$_{-5}$p$_{-5--}$

$_4$I1+L1+E1/16/19/5+H1/19/5'p$_{-5--4}^{3}$I1+L1+E1/16/19/5+H1/19/5'c$_{-5}$p$_{-5--}$

$_4$I1+L1+E1/16/19/5+H1/19/5'p$_{-5--4}$c$_{-5}$p$_{-5--4}$I1+L1+E1/16/19/5+H1/19/5'p$_{-5--}$

$_4^{3}$I1+L1+E1/16/19/5+H1/19/5'c$_{-5}$p$_{-5--4}$I1+L1+E1/16/19/5+H1/19/5'p$_{-5--4}$c-

$_5$I1+L1+E1/16/19/5+H1/19/5'p$_{-5--4}^{2}$I1+L1+E1/16/19/5+H1/19/5'c$_{-5}$p$_{-5--}$

$_4^{6}$I1+L1+E1/16/19/5+H1/19/5'c$_{-5}$p$_{-5--4}^{3}$c$_{-5}$p$_{-5--4}^{4}$I1+L1+E1/16/19/5+H1/19/5'c$_{-5}$p$_{-5--}$

$_4^{2}$I1+L1+E1/16/19/5+H1/19/5'c$_{-5}$p$_{-5--4}^{2}$c$_{-5}$p$_{-5--4}^{2}$I1+L1+E1/16/19/5+H1/19/5'$^{2}$c$_{-5}$p$_{-5--}$

$_4$I1+L1+E1/16/19/5+H1/19/5'c$_{-5}$p$_{-5--4}^{4}$I1+L1+E1/16/19/5+H1/19/5'p$_{-5--2}$p$_{-5--}$

$_4$I1+L1+E1/16/19/5+H1/19/5'c$_{-5}$p$_{-5--4}^{3}$I1+L1+E1/16/19/5+H1/19/5'c$_{-5}$p$_{-5--}$

$_4^{4}$I1+L1+E1/16/19/5+H1/19/5'p$_{-5--2}$p$_{-5--4}$I1+L1+E1/16/19/5+H1/19/5'c$_{-5}$p$_{-5--}$

$_4^{3}$I1+L1+E1/16/19/5+H1/19/5'c$_{-5}$p$_{-5--4}$I1+L1+E1/16/19/5+H1/19/5'c$_{-5}$p$_{-5--}$

$_4^{3}$I1+L1+E1/16/19/5+H1/19/5'c$_{-5}$p$_{-5--4}^{3}$I1+L1+E1/16/19/5+H1/19/5'c$_{-5}$p$_{-5--}$

$_4^{2}$I1+L1+E1/16/19/5+H1/19/5'$^{2}$c$_{-5}$p$_{-5--4}^{2}$I1+L1+E1/16/19/5+H1/19/5'$^{2}$c$_{-5}$p$_{-5--}$

$_4^{4}$I1+L1+E1/16/19/5+H1/19/5'c$_{-5}$p$_{-5--4}^{2}$I1+L1+E1/16/19/5+H1/19/5'c$_{-5}$p$_{-5--}$

$_4^{3}$I1+L1+E1/16/19/5+H1/19/5'c$_{-5}$p$_{-5--4}^{2}$I1+L1+E1/16/19/5+H1/19/5'$^{2}$c$_{-5}$p$_{-5--}$

$_4^{3}$I1+L1+E1/16/19/5+H1/19/5'c$_{-5}$p$_{-5--4}^{2}$I1+L1+E1/16/19/5+H1/19/5'c$_{-5}$p$_{-5--4}^{3}$c$_{-5}$p$_{-5--}$

$_4^{3}$I1+L1+E1/16/19/5+H1/19/5'c$_{-5}$p$_{-5--4}$I1+L1+E1/16/19/5+H1/19/5'c$_{-5}$p$_{-5--}$

$_4$I1+L1+E1/16/19/5+H1/19/5'p$_{-5--4}^{3}$I1+L1+E1/16/19/5+H1/19/5'c$_{-5}$p$_{-5--}$

$_4$I1+L1+E1/16/19/5+H1/19/5'p$_{-5--4}^{3}$I1+L1+E1/16/19/5+H1/19/5'c$_{-5}$p$_{-5--}$

$_4^{3}$I1+L1+E1/16/19/5+H1/19/5'c$_{-5}$p$_{-5--4}^{2}$I1+L1+E1/16/19/5+H1/19/5'c$_{-5}$p$_{-5--}$

$_4^{3}$I1+L1+E1/16/19/5+H1/19/5'c$_{-5}$p$_{-5--4}^{2}$I1+L1+E1/16/19/5+H1/19/5'$^{2}$c$_{-5}$p$_{-5--}$

$_4^{2}$I1+L1+E1/16/19/5+H1/19/5'$^{2}$c$_{-5}$p$_{-5--4}^{2}$I1+L1+E1/16/19/5+H1/19/5'$^{2}$c$_{-5}$p$_{-5--}$

$_4^{2}$I1+L1+E1/16/19/5+H1/19/5'$^{2}$c$_{-5}$p$_{-5--4}^{3}$c$_{-5}$p$_{-5--4}^{3}$I1+L1+E1/16/19/5+H1/19/5'c$_{-5}$p$_{-5--}$

$_4$I1+L1+E1/16/19/5+H1/19/5'c$_{-5}$p$_{-5--4}$I1+L1+E1/16/19/5+H1/19/5'c$_{-5}$p$_{-5--}$

$_4$I1+L1+E1/16/19/5+H1/19/5'c$_{-5}$p$_{-5--4}$I1+L1+E1/16/19/5+H1/19/5'p$_{-5--}$

$_4^{3}$I1+L1+E1/16/19/5+H1/19/5'c$_{-5}$p$_{-5--4}^{2}$I1+L1+E1/16/19/5+H1/19/5'c$_{-5}$p$_{-5--}$

$_4$I1+L1+E1/16/19/5+H1/19/5'c$_{-5}$D1/19/5+P1+F1/19+G1/19/5+V1/19.1'p$_{-5--}$

$_4$I1+L1+E1/16/19/5+H1/19/5'c$_{-5}$p$_{-5--4}^{2}$I1+L1+E1/16/19/5+H1/19/5'p$_{-5--4}$c$_{-5}$p$_{-5--}$

$_4^{2}$I1+L1+E1/16/19/5+H1/19/5'c$_{-5}$p$_{-5--4}^{2}$I1+L1+E1/16/19/5+H1/19/5'c$_{-5}$p$_{-5--}$

$_4^{2}$I1+L1+E1/16/19/5+H1/19/5'c$_{-5}$p$_{-5--4}^{2}$I1+L1+E1/16/19/5+H1/19/5'c$_{-5}$p$_{-5--4}$c$_{-5}$p$_{-5--}$

$_4^{2}$I1+L1+E1/16/19/5+H1/19/5'c$_{-5}$p$_{-5--4}^{2}$I1+L1+E1/16/19/5+H1/19/5'c$_{-5}$p$_{-5--4}$p$_{-5--2}$c$_{-5}$p$_{-5--4}^{3}$p$_{-5--2}$p$_{-5--4}$c$_{-5}$p-

5--4c-5I1+L1+E1/16/19/5+H1/19/5'c-3p-5--4^3I1+L1+E1/16/19/5+H1/19/5'p-5--
4I1+L1+E1/16/19/5+H1/19/5'p-5--4c-5I1+L1+E1/16/19/5+H1/19/5'p-5--4^5c-
5I1+L1+E1/16/19/5+H1/19/5'^2p-5--4c-5p-5--4I1+L1+E1/16/19/5+H1/19/5'p-5--4c-5p-5--
4^4I1+L1+E1/16/19/5+H1/19/5'c-5p-5--4I1+L1+E1/16/19/5+H1/19/5'c-5p-5--
4I1+L1+E1/16/19/5+H1/19/5'c-5p-5--4I1+L1+E1/16/19/5+H1/19/5'p-5--4c-5p-5--4^2c-5p-5--4c-5p-5--4c-5p-5--
4I1+L1+E1/16/19/5+H1/19/5'p-5--4^2I1+L1+E1/16/19/5+H1/19/5'c-5I1+L1+E1/16/19/5+H1/19/5'p-5--
4^2I1+L1+E1/16/19/5+H1/19/5'D1/19/5+P1+F1/19+G1/19/5+V1/19.1'p-5--
4^2I1+L1+E1/16/19/5+H1/19/5'c-5I1+L1+E1/16/19/5+H1/19/5'c-5p-5--4I1+L1+E1/16/19/5+H1/19/5'p-
5--4^2I1+L1+E1/16/19/5+H1/19/5'c-5I1+L1+E1/16/19/5+H1/19/5'p-5--
4^2I1+L1+E1/16/19/5+H1/19/5'D1/19/5+P1+F1/19+G1/19/5+V1/19.1'I1+L1+E1/16/19/5+H1/19/5'c
-5I1+L1+E1/16/19/5+H1/19/5'p-5--4c-4D1/19/5+P1+F1/19+G1/19/5+V1/19.1'p-5--
4^2I1+L1+E1/16/19/5+H1/19/5'c-
4D1/19/5+P1+F1/19+G1/19/5+V1/19.1'I1+L1+E1/16/19/5+H1/19/5'c-5p-5--
4I1+L1+E1/16/19/5+H1/19/5'c-5p-5--4c-5p-5--4c-5p-5--4c-5p-5--4c-5p-5--4I1+L1+E1/16/19/5+H1/19/5'c-5p-5--
4c-5p-5--4c-5p-5--4^3I1+L1+E1/16/19/5+H1/19/5'p-5--4c-5p-5--4^2I1+L1+E1/16/19/5+H1/19/5'c-5p-5--
4^2I1+L1+E1/16/19/5+H1/19/5'c-5p-5--4^2I1+L1+E1/16/19/5+H1/19/5'c-5p-5--
4^2I1+L1+E1/16/19/5+H1/19/5'c-5p-5--4^2I1+L1+E1/16/19/5+H1/19/5'c-5p-5--
4^2I1+L1+E1/16/19/5+H1/19/5'c-5p-5--4^4I1+L1+E1/16/19/5+H1/19/5'c-
4D1/19/5+P1+F1/19+G1/19/5+V1/19.1'p-5--4I1+L1+E1/16/19/5+H1/19/5'c-5^2p-5--
4^2I1+L1+E1/16/19/5+H1/19/5'c-5^2p-5--4^2I1+L1+E1/16/19/5+H1/19/5'c-5p-5--4c-
4D1/19/5+P1+F1/19+G1/19/5+V1/19.1'p-5--4I1+L1+E1/16/19/5+H1/19/5'c-5^2p-5--
4^2I1+L1+E1/16/19/5+H1/19/5'c-5p-5--4^2I1+L1+E1/16/19/5+H1/19/5'c-5^2p-5--
4^2I1+L1+E1/16/19/5+H1/19/5'c-5p-5--4^3I1+L1+E1/16/19/5+H1/19/5'p-4--
1D1/19/5+P1+F1/19+G1/19/5+V1/19.1'p-5--4I1+L1+E1/16/19/5+H1/19/5'c-5p-5--
4^3I1+L1+E1/16/19/5+H1/19/5'c-5p-5--4^3I1+L1+E1/16/19/5+H1/19/5'c-5p-5--
4^2I1+L1+E1/16/19/5+H1/19/5'c-5p-5--4^2I1+L1+E1/16/19/5+H1/19/5'c-5p-5--
4^3I1+L1+E1/16/19/5+H1/19/5'c-5p-5--4^2I1+L1+E1/16/19/5+H1/19/5'c-5p-5--
4^2I1+L1+E1/16/19/5+H1/19/5'c-5p-5--4^2I1+L1+E1/16/19/5+H1/19/5'c-5p-5--
4I1+L1+E1/16/19/5+H1/19/5'p-5--4^2c-5p-5--4p-5--2p-5--4^3c-5p-5--4p-5--2I1+L1+E1/16/19/5+H1/19/5'c-5p-5--
4^3I1+L1+E1/16/19/5+H1/19/5'c-5p-5--4^4I1+L1+E1/16/19/5+H1/19/5'c-5p-5--
4^3I1+L1+E1/16/19/5+H1/19/5'c-5p-5--4^3I1+L1+E1/16/19/5+H1/19/5'^2p-5--4^3c-
4D1/19/5+P1+F1/19+G1/19/5+V1/19.1'p-5--4I1+L1+E1/16/19/5+H1/19/5'c-5p-5--4^3c-
5I1+L1+E1/16/19/5+H1/19/5'p-5--4^4I1+L1+E1/16/19/5+H1/19/5'c-5I1+L1+E1/16/19/5+H1/19/5'p-5--
4I1+L1+E1/16/19/5+H1/19/5'p-3--5c-5p-5--4^5I1+L1+E1/16/19/5+H1/19/5'c-5p-5--
4I1+L1+E1/16/19/5+H1/19/5'p-5--4^4c-5p-5--4^2c-5I1+L1+E1/16/19/5+H1/19/5'p-5--4^3c-5p-5--4c-
5I1+L1+E1/16/19/5+H1/19/5'p-5--4^3c-5p-5--4c-5^2p-5--4^2p-5--2p-5--4p-5--
3D1/19/5+P1+F1/19+G1/19/5+V1/19.1'c-5p-5--4^2p-5--3D1/19/5+P1+F1/19+G1/19/5+V1/19.1'c-5p-5--
4D1/19/5+P1+F1/19+G1/19/5+V1/19.1'p-5--3D1/19/5+P1+F1/19+G1/19/5+V1/19.1'c-
5I1+L1+E1/16/19/5+H1/19/5'p-5--4D1/19/5+P1+F1/19+G1/19/5+V1/19.1'c-5p-5--4c-5p-5--
4I1+L1+E1/16/19/5+H1/19/5'c-5p-5--4^2p-5--2D1/19/5+P1+F1/19+G1/19/5+V1/19.1'c-5p-5--4^4c-5p-5--4c-
5p-5--4I1+L1+E1/16/19/5+H1/19/5'c-3c-5p-5--4^3I1+L1+E1/16/19/5+H1/19/5'^2c-5p-5--4^4c-5p-5--4^4c-5p-5--
4I1+L1+E1/16/19/5+H1/19/5'p-5--4^2c-5p-5--4I1+L1+E1/16/19/5+H1/19/5'p-5--
4I1+L1+E1/16/19/5+H1/19/5'p-5--4^4I1+L1+E1/16/19/5+H1/19/5'c-5p-5--
4D1/19/5+P1+F1/19+G1/19/5+V1/19.1'p-5--4I1+L1+E1/16/19/5+H1/19/5'^2c-5p-5--
4I1+L1+E1/16/19/5+H1/19/5'p-5--4I1+L1+E1/16/19/5+H1/19/5'^2c-5p-5--
4^4I1+L1+E1/16/19/5+H1/19/5'c-5p-5--4^4I1+L1+E1/16/19/5+H1/19/5'c-5p-5--
4^3I1+L1+E1/16/19/5+H1/19/5'c-5p-5--4I1+L1+E1/16/19/5+H1/19/5'p-5--
4I1+L1+E1/16/19/5+H1/19/5'p-5--4I1+L1+E1/16/19/5+H1/19/5'p-5--4^2I1+L1+E1/16/19/5+H1/19/5'c-
5p-5--4I1+L1+E1/16/19/5+H1/19/5'p-5--4^2I1+L1+E1/16/19/5+H1/19/5'c-5p-5--
4I1+L1+E1/16/19/5+H1/19/5'p-5--4^2I1+L1+E1/16/19/5+H1/19/5'c-5p-5--
4^3I1+L1+E1/16/19/5+H1/19/5'c-5p-5--4I1+L1+E1/16/19/5+H1/19/5'p-5--

4I1+L1+E1/16/19/5+H1/19/5'c-5p-5--4I1+L1+E1/16/19/5+H1/19/5'p-5--
42I1+L1+E1/16/19/5+H1/19/5'c-5p-5--4I1+L1+E1/16/19/5+H1/19/5'p-5--
4I1+L1+E1/16/19/5+H1/19/5'c-5p-5--4I1+L1+E1/16/19/5+H1/19/5'c-52p-5--
4I1+L1+E1/16/19/5+H1/19/5'c-5p-5--42c-5p-5--42c-5p-5--42c-5p-5--42c-5p-5--4I1+L1+E1/16/19/5+H1/19/5'c-
5p-5--42I1+L1+E1/16/19/5+H1/19/5'c-5p-5--42c-5p-5--42I1+L1+E1/16/19/5+H1/19/5'c-
4D1/19/5+P1+F1/19+G1/19/5+V1/19.1'p-5--42c-5p-5--42I1+L1+E1/16/19/5+H1/19/5'c-
4D1/19/5+P1+F1/19+G1/19/5+V1/19.1'p-5--4I1+L1+E1/16/19/5+H1/19/5'c-5p-5--48c-5p-5--42c-5p-5--
44I1+L1+E1/16/19/5+H1/19/5'c-5p-5--46I1+L1+E1/16/19/5+H1/19/5'c-5p-5--
4I1+L1+E1/16/19/5+H1/19/5'c-5p-5--42c-5p-5--43c-5p-5--4I1+L1+E1/16/19/5+H1/19/5'2c-5p-5--
44I1+L1+E1/16/19/5+H1/19/5'c-5p-5--43I1+L1+E1/16/19/5+H1/19/5'c-5p-5--
42D1/19/5+P1+F1/19+G1/19/5+V1/19.1'p-5--4I1+L1+E1/16/19/5+H1/19/5'c-5p-5--
44I1+L1+E1/16/19/5+H1/19/5'c-5p-5--44I1+L1+E1/16/19/5+H1/19/5'c-5p-5--
44I1+L1+E1/16/19/5+H1/19/5'c-5p-5--42I1+L1+E1/16/19/5+H1/19/5'c-5p-5--
43I1+L1+E1/16/19/5+H1/19/5'c-5p-5--43I1+L1+E1/16/19/5+H1/19/5'2p-5--
4I1+L1+E1/16/19/5+H1/19/5'c-5p-5--42I1+L1+E1/16/19/5+H1/19/5'p-5--
4I1+L1+E1/16/19/5+H1/19/5'2p-5--43I1+L1+E1/16/19/5+H1/19/5'c-5p-5--
43I1+L1+E1/16/19/5+H1/19/5'p-5--4I1+L1+E1/16/19/5+H1/19/5'2c-5p-5--
42I1+L1+E1/16/19/5+H1/19/5'c-5p-5--4I1+L1+E1/16/19/5+H1/19/5'p-5--
42I1+L1+E1/16/19/5+H1/19/5'c-5p-5--42I1+L1+E1/16/19/5+H1/19/5'c-5p-5--
44I1+L1+E1/16/19/5+H1/19/5'c-4D1/19/5+P1+F1/19+G1/19/5+V1/19.1'p-5--
4I1+L1+E1/16/19/5+H1/19/5'c-5p-5--4I1+L1+E1/16/19/5+H1/19/5'p-5--
42I1+L1+E1/16/19/5+H1/19/5'c-5p-5--42I1+L1+E1/16/19/5+H1/19/5'c-5p-5--
44I1+L1+E1/16/19/5+H1/19/5'c-5p-5--4I1+L1+E1/16/19/5+H1/19/5'p-5--
4I1+L1+E1/16/19/5+H1/19/5'3c-5p-5--4c-5p-5--43I1+L1+E1/16/19/5+H1/19/5'c-5p-5--
4I1+L1+E1/16/19/5+H1/19/5'p-5--4I1+L1+E1/16/19/5+H1/19/5'3c-5p-5--4c-5p-5--
4I1+L1+E1/16/19/5+H1/19/5'3c-5p-5--44I1+L1+E1/16/19/5+H1/19/5'c-5p-5--
4I1+L1+E1/16/19/5+H1/19/5'p-5--4I1+L1+E1/16/19/5+H1/19/5'2c-5p-5--
42I1+L1+E1/16/19/5+H1/19/5'c-5p-5--4I1+L1+E1/16/19/5+H1/19/5'p-5--4c-5p-5--
4I1+L1+E1/16/19/5+H1/19/5'p-5--4c-5p-5--4I1+L1+E1/16/19/5+H1/19/5'c-5p-5--42c-5p-5--42c-5p-5--42c-5p-5--
43c-5p-5--42c-5p-5--42c-5p-5--44I1+L1+E1/16/19/5+H1/19/5'c-5D1/19/5+P1+F1/19+G1/19/5+V1/19.1'p-5--
4c-5p-5--42c-5p-5--46c-5p-5--42c-5p-5--42I1+L1+E1/16/19/5+H1/19/5'c-5I1+L1+E1/16/19/5+H1/19/5'p-5--
42I1+L1+E1/16/19/5+H1/19/5'c-5p-5--4I1+L1+E1/16/19/5+H1/19/5'2c-5p-5--
4I1+L1+E1/16/19/5+H1/19/5'2c-5p-5--4I1+L1+E1/16/19/5+H1/19/5'2c-5p-5--
4I1+L1+E1/16/19/5+H1/19/5'2c-5p-5--4I1+L1+E1/16/19/5+H1/19/5'2c-5p-5--
4I1+L1+E1/16/19/5+H1/19/5'2c-5p-5--4I1+L1+E1/16/19/5+H1/19/5'2c-5p-5--
4I1+L1+E1/16/19/5+H1/19/5'2c-5p-5--4I1+L1+E1/16/19/5+H1/19/5'2c-5p-5--
4I1+L1+E1/16/19/5+H1/19/5'2c-5p-5--4I1+L1+E1/16/19/5+H1/19/5'2c-5p-5--
4I1+L1+E1/16/19/5+H1/19/5'2c-5p-5--4I1+L1+E1/16/19/5+H1/19/5'2c-5p-5--
4I1+L1+E1/16/19/5+H1/19/5'2c-5p-5--4I1+L1+E1/16/19/5+H1/19/5'2c-5p-5--
4I1+L1+E1/16/19/5+H1/19/5'2c-5p-5--4I1+L1+E1/16/19/5+H1/19/5'2c-5p-5--
4I1+L1+E1/16/19/5+H1/19/5'2c-5p-5--4I1+L1+E1/16/19/5+H1/19/5'2c-5p-5--
4I1+L1+E1/16/19/5+H1/19/5'2c-5p-5--4I1+L1+E1/16/19/5+H1/19/5'2c-5p-5--
4I1+L1+E1/16/19/5+H1/19/5'2c-5p-5--4I1+L1+E1/16/19/5+H1/19/5'2c-5p-5--
43I1+L1+E1/16/19/5+H1/19/5'c-5p-5--44c-5p-5--44I1+L1+E1/16/19/5+H1/19/5'c-5p-5--
4I1+L1+E1/16/19/5+H1/19/5'p-5--42I1+L1+E1/16/19/5+H1/19/5'c-52p-5--
42I1+L1+E1/16/19/5+H1/19/5'c-5p-5--44I1+L1+E1/16/19/5+H1/19/5'c-5p-5--
42I1+L1+E1/16/19/5+H1/19/5'2c-5p-5--44I1+L1+E1/16/19/5+H1/19/5'c-5p-5--
4I1+L1+E1/16/19/5+H1/19/5'c-5I1+L1+E1/16/19/5+H1/19/5'p-5--43I1+L1+E1/16/19/5+H1/19/5'c-5p-
5--43I1+L1+E1/16/19/5+H1/19/5'2c-5p-5--42I1+L1+E1/16/19/5+H1/19/5'c-5p-5--
42I1+L1+E1/16/19/5+H1/19/5'c-5c-3I1+L1+E1/16/19/5+H1/19/5'p-5--4I1+L1+E1/16/19/5+H1/19/5'c-
5p-5--44c-5p-5--4c-5p-5--42c-5p-5--42c-5p-5--42c-5p-5--44c-5p-5--44c-5p-5--42c-5p-5--42c-5p-5--42c-5p-5--44c-5p-5--44c-5p-5--
44c-5I1+L1+E1/16/19/5+H1/19/5'3p-5--44c-

4D1/19/5+P1+F1/19+G1/19/5+V1/19.1'I1+L1+E1/16/19/5+H1/19/5'2p-5--44c-
4D1/19/5+P1+F1/19+G1/19/5+V1/19.1'p-5--43c-5p-5--44c-5I1+L1+E1/16/19/5+H1/19/5'3c-5p-5--
4I1+L1+E1/16/19/5+H1/19/5'2p-5--44c-4D1/19/5+P1+F1/19+G1/19/5+V1/19.1'p-5--43c-
5I1+L1+E1/16/19/5+H1/19/5'3c-5p-5--4c-5p-5--44I1+L1+E1/16/19/5+H1/19/5'c-5p-5--
43I1+L1+E1/16/19/5+H1/19/5'2c-5p-5--4I1+L1+E1/16/19/5+H1/19/5'p-5--
42I1+L1+E1/16/19/5+H1/19/5'c-
5D1/19/5+P1+F1/19+G1/19/5+V1/19.1'I1+L1+E1/16/19/5+H1/19/5'p-5--
42I1+L1+E1/16/19/5+H1/19/5'c-5p-5--4I1+L1+E1/16/19/5+H1/19/5'p-5--42c-5p-5--
4I1+L1+E1/16/19/5+H1/19/5'p-5--42c-5p-5--42c-5p-5--4I1+L1+E1/16/19/5+H1/19/5'p-5--
4I1+L1+E1/16/19/5+H1/19/5'p-5--42I1+L1+E1/16/19/5+H1/19/5'2c-
4D1/19/5+P1+F1/19+G1/19/5+V1/19.1'2p-5--4I1+L1+E1/16/19/5+H1/19/5'2p-5--
42I1+L1+E1/16/19/5+H1/19/5'c-5p-5--4I1+L1+E1/16/19/5+H1/19/5'c-5p-5--45c-5p-5--42c-5p-5--
4I1+L1+E1/16/19/5+H1/19/5'p-5--4I1+L1+E1/16/19/5+H1/19/5'p-5--44I1+L1+E1/16/19/5+H1/19/5'c-
5p-5--44I1+L1+E1/16/19/5+H1/19/5'c-5p-5--43I1+L1+E1/16/19/5+H1/19/5'p-5--
43I1+L1+E1/16/19/5+H1/19/5'c-5p-5--4I1+L1+E1/16/19/5+H1/19/5'c-5p-5--
42I1+L1+E1/16/19/5+H1/19/5'c-5p-5--43c-5p-5--4I1+L1+E1/16/19/5+H1/19/5'c-5p-5--
4I1+L1+E1/16/19/5+H1/19/5'2p-5--4I1+L1+E1/16/19/5+H1/19/5'c-5p-5--
4I1+L1+E1/16/19/5+H1/19/5'c-5p-5--42c-5p-5--43I1+L1+E1/16/19/5+H1/19/5'c-5p-5--
4I1+L1+E1/16/19/5+H1/19/5'c-5p-5--43I1+L1+E1/16/19/5+H1/19/5'c-3I1+L1+E1/16/19/5+H1/19/5'p-
5--42I1+L1+E1/16/19/5+H1/19/5'c-5p-5--44I1+L1+E1/16/19/5+H1/19/5'c-
5D1/19/5+P1+F1/19+G1/19/5+V1/19.1'p-5--4c-5p-5--42D1/19/5+P1+F1/19+G1/19/5+V1/19.1'p-5--
4I1+L1+E1/16/19/5+H1/19/5'c-5p-5--43I1+L1+E1/16/19/5+H1/19/5'p-5--
43I1+L1+E1/16/19/5+H1/19/5'c-3p-5--4I1+L1+E1/16/19/5+H1/19/5'c-5p-5--
4I1+L1+E1/16/19/5+H1/19/5'c-3c-5p-5--42c-52I1+L1+E1/16/19/5+H1/19/5'p-5--4c-
52I1+L1+E1/16/19/5+H1/19/5'p-5--4c-5p-5--4c-5p-5--42c-5p-5--44c-5p-5--4c-5p-5--42c-5p-5--42c-5p-5--4c-52p-5--4p-5--
1D1/19/5+P1+F1/19+G1/19/5+V1/19.1'p-5--44c-5I1+L1+E1/16/19/5+H1/19/5'p-5--43c-
5I1+L1+E1/16/19/5+H1/19/5'p-5--44c-5p-5--44c-5p-5--
43I1+L1+E1/16/19/5+H1/19/5'I1+L1+E1/16/19/5+H1/19/5c4p4-
5I1+L1+E1/16/19/5+H1/19/5c5I1+L1+E1/16/19/5+H1/19/5p4-5I1+L1+E1/16/19/5+H1/19/5c4p4-
53I1+L1+E1/16/19/5+H1/19/5p4-5I1+L1+E1/16/19/5+H1/19/5c4p4-53I1+L1+E1/16/19/5+H1/19/5p4-
6c5I1+L1+E1/16/19/5+H1/19/5p4-54c5I1+L1+E1/16/19/5+H1/19/5p4-
5I1+L1+E1/16/19/5+H1/19/52c42p4-53c4p4-52I1+L1+E1/16/19/5+H1/19/52c4p4-
52I1+L1+E1/16/19/5+H1/19/5c4p4-5I1+L1+E1/16/19/5+H1/19/5p4-5I1+L1+E1/16/19/5+H1/19/5p4-
52c42p4-5c42p4-54I1+L1+E1/16/19/5+H1/19/52p4-5c42p4-53I1+L1+E1/16/19/5+H1/19/5p4-
5I1+L1+E1/16/19/5+H1/19/5p4-5c5I1+L1+E1/16/19/5+H1/19/5p4-
5c4D1/19/5+P1+F1/19+G1/19/5+V1/19.1c42p4-5I1+L1+E1/16/19/5+H1/19/5p4-
5c5I1+L1+E1/16/19/5+H1/19/5p4-52c42p4-5I1+L1+E1/16/19/5+H1/19/5p4-5c42p4-
5I1+L1+E1/16/19/5+H1/19/5p4-52c4p4-52c4p4-52I1+L1+E1/16/19/5+H1/19/52c4p4-
52I1+L1+E1/16/19/5+H1/19/5c4p4-53c4p4-53c4p4-53c4p4-5c4p4-52p3-5I1+L1+E1/16/19/5+H1/19/5p4-
5I1+L1+E1/16/19/5+H1/19/5c43p4-5c42p4-5I1+L1+E1/16/19/5+H1/19/5p4-5c4p4-
5I1+L1+E1/16/19/5+H1/19/52p4-52c42D1/19/5+P1+F1/19+G1/19/5+V1/19.1p4-
52I1+L1+E1/16/19/5+H1/19/5c4p4-5c4p4-5I1+L1+E1/16/19/5+H1/19/5p4-
53I1+L1+E1/16/19/5+H1/19/5p4-52I1+L1+E1/16/19/5+H1/19/5p4-5c4p4-
5I1+L1+E1/16/19/5+H1/19/5p4-5c4p4-52I1+L1+E1/16/19/5+H1/19/5p4-
52I1+L1+E1/16/19/5+H1/19/5c4p4-
5I1+L1+E1/16/19/5+H1/19/5D1/19/5+P1+F1/19+G1/19/5+V1/19.1c6c5I1+L1+E1/16/19/5+H1/19/
5p4-5D1/19/5+P1+F1/19+G1/19/5+V1/19.1p3-5p4-
52I1+L1+E1/16/19/5+H1/19/54D1/19/5+P1+F1/19+G1/19/5+V1/19.1c6c5I1+L1+E1/16/19/5+H1/1
9/5p4-5D1/19/5+P1+F1/19+G1/19/5+V1/19.1p3-5p4-52I1+L1+E1/16/19/5+H1/19/5p4-
52I1+L1+E1/16/19/5+H1/19/5c4p4-
5I1+L1+E1/16/19/5+H1/19/5D1/19/5+P1+F1/19+G1/19/5+V1/19.1c6c5I1+L1+E1/16/19/5+H1/19/

5$^{2}$p$_{4-5}^{2}$c$_{5}$I1+L1+E1/16/19/5+H1/19/5c$_{4}$p$_{4-5}$c$_{5}$I1+L1+E1/16/19/5+H1/19/5$^{2}$p$_{4-}$
$_{5}^{2}$c$_{5}$I1+L1+E1/16/19/5+H1/19/5$^{2}$p$_{3-5}$p$_{4-}$
$_{5}$c$_{5}$I1+L1+E1/16/19/5+H1/19/5c$_{5}$I1+L1+E1/16/19/5+H1/19/5p$_{4-}$
$_{5}$I1+L1+E1/16/19/5+H1/19/5c$_{5}$I1+L1+E1/16/19/5+H1/19/5p$_{4-}$
$_{5}$I1+L1+E1/16/19/5+H1/19/5D1/19/5+P1+F1/19+G1/19/5+V1/19.1c$_{6}$I1+L1+E1/16/19/5+H1/19/5p
$_{4-5}$I1+L1+E1/16/19/5+H1/19/5p$_{4-5}$I1+L1+E1/16/19/5+H1/19/5p$_{4-5}$I1+L1+E1/16/19/5+H1/19/5p$_{4-}$
$_{5}$I1+L1+E1/16/19/5+H1/19/5$^{2}$c$_{2}$p$_{2-5}^{2}$I1+L1+E1/16/19/5+H1/19/5$^{2}$p$_{3-5}$p$_{4-}$
$_{5}^{3}$c$_{4}$D1/19/5+P1+F1/19+G1/19/5+V1/19.1p$_{1-5}$I1+L1+E1/16/19/5+H1/19/5$^{3}$p$_{4-}$
$_{5}^{5}$c$_{5}$I1+L1+E1/16/19/5+H1/19/5p$_{4-5}$c$_{4}$p$_{4-5}^{2}$I1+L1+E1/16/19/5+H1/19/5p$_{4-}$
$_{5}^{2}$I1+L1+E1/16/19/5+H1/19/5c$_{4}$p$_{4-5}^{2}$I1+L1+E1/16/19/5+H1/19/5c$_{5}$I1+L1+E1/16/19/5+H1/19/5p$_{4-}$
$_{5}$I1+L1+E1/16/19/5+H1/19/5$^{2}$p$_{4-5}$c$_{5}$I1+L1+E1/16/19/5+H1/19/5c$_{4}$p$_{4-}$
$_{5}$c$_{5}$I1+L1+E1/16/19/5+H1/19/5p$_{4-5}^{2}$I1+L1+E1/16/19/5+H1/19/5c$_{5}$I1+L1+E1/16/19/5+H1/19/5p$_{4-}$
$_{5}$c$_{5}$I1+L1+E1/16/19/5+H1/19/5p$_{4-5}^{7}$I1+L1+E1/16/19/5+H1/19/5$^{2}$c$_{4}$p$_{4-}$
$_{5}^{2}$I1+L1+E1/16/19/5+H1/19/5p$_{3-5}$p$_{4-5}^{2}$I1+L1+E1/16/19/5+H1/19/5$^{2}$p$_{3-5}$p$_{4-}$
$_{5}^{3}$I1+L1+E1/16/19/5+H1/19/5$^{2}$c$_{5}$p$_{5-}$
$_{3}$I1+L1+E1/16/19/5+H1/19/5$^{2}$c$_{3}^{2}$C1/19/5I1+L1+E1/16/19/5+H1/19/5$^{2}$p$_{3-5}$p$_{4-}$
$_{5}^{3}$c$_{5}$I1+L1+E1/16/19/5+H1/19/5c$_{5}$I1+L1+E1/16/19/5+H1/19/5p$_{4-5}$I1+L1+E1/16/19/5+H1/19/5p$_{4-}$
$_{5}$c$_{5}$I1+L1+E1/16/19/5+H1/19/5c$_{4}$I1+L1+E1/16/19/5+H1/19/5$^{2}$c$_{3}$C1/19/5I1+L1+E1/16/19/5+H1/19/
5$^{2}$p$_{3-5}$p$_{4-5}^{3}$c$_{5}$I1+L1+E1/16/19/5+H1/19/5c$_{5}$I1+L1+E1/16/19/5+H1/19/5p$_{4-}$
$_{5}$I1+L1+E1/16/19/5+H1/19/5p$_{4-5}$c$_{5}$I1+L1+E1/16/19/5+H1/19/5c$_{4}^{2}$I1+L1+E1/16/19/5+H1/19/5$^{2}$p$_{3-}$
$_{5}$p$_{4-5}^{3}$I1+L1+E1/16/19/5+H1/19/5c$_{5}$I1+L1+E1/16/19/5+H1/19/5c$_{5}$I1+L1+E1/16/19/5+H1/19/5p$_{4-}$
$_{5}$I1+L1+E1/16/19/5+H1/19/5p$_{4-5}$c$_{5}$p$_{4-5}$c$_{4}$p$_{4-5}$I1+L1+E1/16/19/5+H1/19/5p$_{4-}$
$_{5}$c$_{4}$D1/19/5+P1+F1/19+G1/19/5+V1/19.1I1+L1+E1/16/19/5+H1/19/5p$_{4-5}$p$_{4-}$
$_{6}$I1+L1+E1/16/19/5+H1/19/5$^{2}$p$_{4-5}$c$_{4}$p$_{4-5}$I1+L1+E1/16/19/5+H1/19/5p$_{4-}$
$_{5}$I1+L1+E1/16/19/5+H1/19/5c$_{4}$p$_{4-5}$p$_{3-}$
$_{5}$c$_{4}$D1/19/5+P1+F1/19+G1/19/5+V1/19.1I1+L1+E1/16/19/5+H1/19/5p$_{4-5}$p$_{4-}$
$_{6}$I1+L1+E1/16/19/5+H1/19/5$^{2}$p$_{4-5}$c$_{4}$p$_{4-5}$I1+L1+E1/16/19/5+H1/19/5p$_{4-}$
$_{5}$I1+L1+E1/16/19/5+H1/19/5c$_{4}$p$_{4-5}$I1+L1+E1/16/19/5+H1/19/5p$_{4-5}$p$_{5-}$
$_{1}$I1+L1+E1/16/19/5+H1/19/5D1/19/5+P1+F1/19+G1/19/5+V1/19.1p$_{1-5}$p$_{4-}$
$_{5}$I1+L1+E1/16/19/5+H1/19/5p$_{4-5}$c$_{5}$p$_{4-5}$c$_{4}$p$_{4-5}$I1+L1+E1/16/19/5+H1/19/5p$_{4-}$
$_{5}^{2}$I1+L1+E1/16/19/5+H1/19/5$^{2}$p$_{4-5}$I1+L1+E1/16/19/5+H1/19/5$^{2}$c$_{2}$p$_{2-}$
$_{5}^{2}$I1+L1+E1/16/19/5+H1/19/5$^{2}$p$_{3-5}$p$_{4-5}^{2}$c$_{4}$D1/19/5+P1+F1/19+G1/19/5+V1/19.1p$_{1-}$
$_{5}$I1+L1+E1/16/19/5+H1/19/5$^{3}$p$_{4-5}^{4}$I1+L1+E1/16/19/5+H1/19/5c$_{5}$I1+L1+E1/16/19/5+H1/19/5p$_{4-}$
$_{5}$c$_{4}$p$_{4-5}^{2}$I1+L1+E1/16/19/5+H1/19/5p$_{4-5}^{2}$I1+L1+E1/16/19/5+H1/19/5c$_{4}$p$_{4-}$
$_{5}^{2}$I1+L1+E1/16/19/5+H1/19/5c$_{5}$I1+L1+E1/16/19/5+H1/19/5p$_{4-5}$I1+L1+E1/16/19/5+H1/19/5$^{2}$p$_{4-}$
$_{5}$c$_{5}$I1+L1+E1/16/19/5+H1/19/5c$_{4}$p$_{4-5}$c$_{5}$I1+L1+E1/16/19/5+H1/19/5p$_{4-}$
$_{5}^{2}$I1+L1+E1/16/19/5+H1/19/5c$_{5}$I1+L1+E1/16/19/5+H1/19/5p$_{4-5}$c$_{5}$I1+L1+E1/16/19/5+H1/19/5p$_{4-}$
$_{5}^{7}$I1+L1+E1/16/19/5+H1/19/5$^{2}$c$_{5}$I1+L1+E1/16/19/5+H1/19/5p$_{4-5}$I1+L1+E1/16/19/5+H1/19/5p$_{3-5}$p$_{4-}$
$_{5}^{2}$I1+L1+E1/16/19/5+H1/19/5$^{2}$p$_{3-5}$p$_{4-5}^{3}$I1+L1+E1/16/19/5+H1/19/5$^{2}$c$_{5}$p$_{5-}$
$_{3}$I1+L1+E1/16/19/5+H1/19/5$^{2}$c$_{3}^{2}$C1/19/5I1+L1+E1/16/19/5+H1/19/5$^{2}$p$_{3-5}$p$_{4-}$
$_{5}^{3}$c$_{5}$I1+L1+E1/16/19/5+H1/19/5c$_{5}$I1+L1+E1/16/19/5+H1/19/5p$_{4-5}$I1+L1+E1/16/19/5+H1/19/5p$_{4-}$
$_{5}$c$_{5}$I1+L1+E1/16/19/5+H1/19/5c$_{4}$I1+L1+E1/16/19/5+H1/19/5$^{2}$p$_{3-5}$p$_{4-}$
$_{5}^{3}$c$_{5}$I1+L1+E1/16/19/5+H1/19/5c$_{5}$I1+L1+E1/16/19/5+H1/19/5p$_{4-5}$I1+L1+E1/16/19/5+H1/19/5p$_{4-}$
$_{5}$c$_{5}$I1+L1+E1/16/19/5+H1/19/5c$_{4}^{2}$I1+L1+E1/16/19/5+H1/19/5$^{2}$p$_{3-5}$p$_{4-}$
$_{5}^{3}$I1+L1+E1/16/19/5+H1/19/5c$_{5}$I1+L1+E1/16/19/5+H1/19/5c$_{5}$I1+L1+E1/16/19/5+H1/19/5p$_{4-}$
$_{5}$I1+L1+E1/16/19/5+H1/19/5p$_{4-5}$c$_{5}$p$_{4-5}$c$_{4}$p$_{4-5}$I1+L1+E1/16/19/5+H1/19/5p$_{4-}$
$_{5}$c$_{4}$D1/19/5+P1+F1/19+G1/19/5+V1/19.1I1+L1+E1/16/19/5+H1/19/5p$_{4-5}$p$_{4-}$
$_{6}$I1+L1+E1/16/19/5+H1/19/5$^{2}$p$_{4-5}$c$_{4}$p$_{4-5}^{3}$I1+L1+E1/16/19/5+H1/19/5c$_{4}$p$_{4-5}^{2}$p$_{3-}$
$_{5}$c$_{4}$D1/19/5+P1+F1/19+G1/19/5+V1/19.1I1+L1+E1/16/19/5+H1/19/5p$_{4-5}^{2}$p$_{3-}$
$_{5}$c$_{4}$D1/19/5+P1+F1/19+G1/19/5+V1/19.1I1+L1+E1/16/19/5+H1/19/5p$_{4-5}$p$_{4-}$
$_{6}$I1+L1+E1/16/19/5+H1/19/5$^{2}$p$_{4-5}$c$_{4}$p$_{4-5}$I1+L1+E1/16/19/5+H1/19/5p$_{4-}$

$_5$I1+L1+E1/16/19/5+H1/19/5c$_4$p$_{4-5}$I1+L1+E1/16/19/5+H1/19/5p$_{4-5}$p$_{5-}$
$_1$I1+L1+E1/16/19/5+H1/19/5p$_{1-5}$p$_{4-5}$I1+L1+E1/16/19/5+H1/19/5p$_{4-5}$c$_5$p$_{4-5}$c$_4$p$_{4-}$
$_5$I1+L1+E1/16/19/5+H1/19/5c$_4$D1/19/5+P1+F1/19+G1/19/5+V1/19.1I1+L1+E1/16/19/5+H1/19/5p
$_{4-5}$p$_{4-6}$I1+L1+E1/16/19/5+H1/19/5$^2$p$_{4-5}$c$_4$p$_{4-5}$I1+L1+E1/16/19/5+H1/19/5p$_{4-}$
$_5^2$I1+L1+E1/16/19/5+H1/19/5c$_4$p$_{4-5}^2$p$_{3-5}$c$_4$p$_{4-5}$I1+L1+E1/16/19/5+H1/19/5p$_{4-5}^3$c$_4$p$_{4-}$
$_5$I1+L1+E1/16/19/5+H1/19/5$^2$p$_{5-1}$I1+L1+E1/16/19/5+H1/19/5p$_{4-5}$I1+L1+E1/16/19/5+H1/19/5p$_{5-}$
$_1$I1+L1+E1/16/19/5+H1/19/5p$_{4-5}$D1/19/5+P1+F1/19+G1/19/5+V1/19.1p$_{1-5}$p$_{4-}$
$_1$I1+L1+E1/16/19/5+H1/19/5$^2$p$_{4-5}^2$p$_{5-1}$I1+L1+E1/16/19/5+H1/19/5p$_{4-}$
$_5$D1/19/5+P1+F1/19+G1/19/5+V1/19.1p$_{1-5}$p$_{4-1}$I1+L1+E1/16/19/5+H1/19/5$^2$p$_{4-5}$p$_{4-6}$p$_{5-}$
$_1$I1+L1+E1/16/19/5+H1/19/5$^2$p$_{4-5}^2$c$_5$I1+L1+E1/16/19/5+H1/19/5$^2$c$_5$I1+L1+E1/16/19/5+H1/19/5$^2$p$_{4-}$
$_5$c$_5$I1+L1+E1/16/19/5+H1/19/5p$_{4-5}$p$_{4-1}$p$_{3-}$
$_5$I1+L1+E1/16/19/5+H1/19/5$^2$c$_5$I1+L1+E1/16/19/5+H1/19/5$^2$c$_5$I1+L1+E1/16/19/5+H1/19/5c$_4$p$_{4-}$
$_5^2$I1+L1+E1/16/19/5+H1/19/5p$_{5-3}$c$_5$I1+L1+E1/16/19/5+H1/19/5$^2$c$_4$p$_{4-}$
$_5^2$I1+L1+E1/16/19/5+H1/19/5p$_{5-3}$c$_5$I1+L1+E1/16/19/5+H1/19/5$^2$c$_4$p$_{4-}$
$_5^2$c$_5$I1+L1+E1/16/19/5+H1/19/5p$_{4-5}$p$_{4-}$
$_1$I1+L1+E1/16/19/5+H1/19/5c$_5$I1+L1+E1/16/19/5+H1/19/5p$_{4-5}$c$_4$p$_{4-}$
$_5$I1+L1+E1/16/19/5+H1/19/5$^3$p$_{4-5}$c$_4$p$_{4-5}$c$_4$p$_{4-}$
$_5$I1+L1+E1/16/19/5+H1/19/5c$_5$I1+L1+E1/16/19/5+H1/19/5$^3$p$_{4-}$
$_5^2$c$_4$D1/19/5+P1+F1/19+G1/19/5+V1/19.1p$_{4-5}$c$_4$p$_{4-5}$I1+L1+E1/16/19/5+H1/19/5$^3$p$_{4-5}$c$_4$p$_{4-}$
$_5$c$_6$c$_5$I1+L1+E1/16/19/5+H1/19/5p$_{5-1}$I1+L1+E1/16/19/5+H1/19/5p$_{4-}$
$_5$I1+L1+E1/16/19/5+H1/19/5D1/19/5+P1+F1/19+G1/19/5+V1/19.1c$_6$p$_{4-}$
$_5$I1+L1+E1/16/19/5+H1/19/5$^4$p$_{4-5}$I1+L1+E1/16/19/5+H1/19/5c$_4$p$_{4-5}$I1+L1+E1/16/19/5+H1/19/5$^4$p$_{4-}$
$_5$I1+L1+E1/16/19/5+H1/19/5p$_{4-5}$c$_4$p$_{4-5}^2$I1+L1+E1/16/19/5+H1/19/5$^4$p$_{2-5}$p$_{4-5}$c$_4$p$_{4-5}$p$_{2-5}$p$_{4-}$
$_5$I1+L1+E1/16/19/5+H1/19/5$^4$p$_{2-5}$p$_{4-5}$I1+L1+E1/16/19/5+H1/19/5c$_6$p$_{4-5}$c$_4$p$_{4-}$
$_5^2$D1/19/5+P1+F1/19+G1/19/5+V1/19.1c$_6$p$_{4-5}^3$c$_4$p$_{4-5}^4$c$_4$p$_{4-5}^2$c$_4$p$_{4-5}$c$_4$p$_{4-}$
$_5$I1+L1+E1/16/19/5+H1/19/5p$_{4-5}^2$I1+L1+E1/16/19/5+H1/19/5c$_5$I1+L1+E1/16/19/5+H1/19/5$^2$p$_{4-}$
$_5$I1+L1+E1/16/19/5+H1/19/5p$_{3-5}$I1+L1+E1/16/19/5+H1/19/5p$_{4-5}$I1+L1+E1/16/19/5+H1/19/5c$_4$p$_{4-}$
$_5^2$c$_4$p$_{1-5}$c$_4$p$_{4-5}$I1+L1+E1/16/19/5+H1/19/5c$_4$p$_{4-5}^2$c$_4$p$_{1-5}$c$_4$p$_{4-5}$p$_{2-5}$A1I1+L1+E1/16/19/5+H1/19/5p$_{4-5}^2$p$_{1-}$
$_3$I1+L1+E1/16/19/5+H1/19/5p$_{4-5}$I1+L1+E1/16/19/5+H1/19/5$^2$

### cen2

B2c$_2^2$p$_{2-3}$p$_{1-2}$C2p$_{4-2}$c$_1$c$_2^{11}$B2A2c$_4^3$p$_{4-2}$c$_4$D2/20c$_3^2$p$_{3-1}$c$_4$p$_{4-2}$A2c$_4$p$_{4-2}$A2c$_4$p$_{4-2}$p$_{1-2}$A2c$_4^2$p$_{4-2}$A2c$_4$p$_{4-}$
$_2$A2p$_{4-2}$A2c$_4$p$_{4-2}$A2p$_{4-2}$A2c$_4$p$_{4-2}$A2c$_4$p$_{4-2}$A2p$_{4-2}$c$_1^3$A2c$_1$A2p$_{4-2}$A2c$_4^3$p$_{4-2}$A2c$_4$p$_{4-2}$A2c$_4^2$p$_{4-1}$c$_4$p$_{4-2}$A2c$_4^5$p$_{4-}$
$_2$A2c$_4^3$c$_1$p$_{1-2}$A2p$_{4-1}$c$_3^9$p$_{3-1}$c$_4^3$p$_{4-2}$A2c$_4$c$_3$p$_{3-1}$c$_4$c$_3^2$C2c$_3^6$A2c$_4$p$_{4-1}$c$_4$c$_3^2$C2c$_3^2$A2c$_4^2$p$_{4-2}$A2c$_4^2$p$_{4-2}$A2c$_4^9$p$_{4-}$
$_2$A2p$_{4-2}$A2c$_4^2$p$_{4-1}^2$c$_4$p$_{4-1}$c$_4$p$_{4-2}$A2c$_4^2$p$_{4-1}$c$_1$p$_{1-2}$A2c$_4$D2/20c$_3$A2p$_{4-1}$c$_4^3$p$_{4-2}$p$_{4-1}$c$_4^2$p$_{4-2}$A2c$_4$D2/20c$_3$A2p$_{4-}$
$_1$c$_4$p$_{4-2}$A2c$_4$p$_{4-2}$A2p$_{4-1}$c$_4$p$_{4-2}$A2p$_{4-2}$A2p$_{4-2}$A2c$_4^2$p$_{4-2}$A2p$_{4-1}$c$_1$p$_{1-2}$A2c$_4$p$_{4-2}$A2c$_4$p$_{4-1}$c$_4^2$p$_{4-1}$c$_1$p$_{1-2}$A2c$_4$p$_{4-}$
$_1$c$_4^2$p$_{4-1}$c$_1$A2c$_4^4$p$_{4-2}$A2c$_4^4$p$_{4-1}$c$_4^4$p$_{4-1}$c$_4$p$_{4-2}$p$_{1-2}$A2c$_4^2$p$_{4-1}$c$_4^8$p$_{4-1}$c$_4^7$p$_{4-1}$c$_4^4$p$_{4-2}$A2c$_4$p$_{4-2}$A2c$_4$p$_{4-2}$A2c$_4^3$p$_{4-}$
$_2$A2c$_4$p$_{4-2}$A2c$_4$p$_{4-2}$A2c$_4$p$_{4-2}$A2c$_4$p$_{4-2}$A2c$_4$p$_{4-2}$A2p$_{4-1}$c$_4$p$_{4-2}$A2c$_4$p$_{4-2}$A2c$_4^2$p$_{4-2}$c$_1$p$_{1-2}$A2c$_4$p$_{4-2}$A2c$_4$p$_{4-}$
$_2$A2c$_4$p$_{4-2}$A2c$_4$p$_{4-2}$A2c$_4$p$_{4-2}$A2c$_4$p$_{4-2}$A2c$_4$p$_{4-2}$A2p$_{4-1}$c$_4$p$_{4-2}$A2c$_4$p$_{4-2}$A2c$_4$p$_{4-2}$A2c$_4$p$_{4-2}$A2c$_4$p$_{4-2}$A2c$_4$p$_{4-}$
$_2$c$_1^2$A2c$_4^2$p$_{4-1}$c$_4$p$_{4-1}$c$_4^2$p$_{4-1}$c$_4$p$_{4-1}$c$_4$p$_{4-2}$c$_1^5$A2c$_4^4$p$_{4-1}$c$_4$p$_{4-1}$c$_4^2$p$_{4-1}$c$_4^3$p$_{4-1}$c$_4^2$p$_{4-1}^2$c$_1^6$p$_{1-2}$A2c$_4$p$_{4-1}$c$_1$p$_{1-}$
$_2$A2c$_4^6$p$_{4-2}$A2c$_4^2$p$_{4-2}$A2c$_4$p$_{4-1}$c$_4^7$p$_{4-1}$c$_1$p$_{1-2}$A2c$_4$p$_{4-2}$A2c$_4$p$_{4-2}$A2c$_4$p$_{4-2}$A2c$_4$p$_{4-2}$c$_3^2$p$_{3-1}$c$_4$p$_{4-2}$A2c$_4^3$p$_{4-}$
$_2$A2c$_4^2$p$_{4-1}$c$_4$p$_{4-2}$A2c$_4$p$_{4-2}$A2p$_{4-1}$c$_4^5$p$_{4-2}$A2c$_4$p$_{4-2}$A2D2/20c$_3$A2p$_{4-1}$c$_4^4$p$_{4-2}$A2c$_4^2$p$_{4-1}$c$_4$p$_{4-2}$A2c$_4$p$_{4-1}$c$_1^2$p$_{1-}$
$_2$A2c$_4^3$p$_{4-1}$c$_4^2$p$_{4-2}$A2c$_4^2$D2/20c$_3^4$A2c$_4^2$p$_{4-2}$A2c$_4$p$_{4-2}$A2c$_4^6$p$_{4-1}$c$_4$p$_{4-2}$A2c$_4^2$p$_{4-2}$c$_1$p$_{1-2}$A2c$_4^8$p$_{4-1}$c$_1^2$A2c$_1^4$p$_{1-}$
$_2$A2c$_4^2$p$_{4-2}$A2c$_4$p$_{4-2}$A2c$_4^5$p$_{4-2}$A2c$_4^6$p$_{4-2}$A2c$_4^6$p$_{4-2}$A2c$_4$p$_{4-1}$c$_1^3$p$_{1-2}$A2c$_4^4$p$_{4-2}$A2c$_4^4$p$_{4-2}$A2c$_4^4$p$_{4-}$
$_2$A2c$_1$c$_3$c$_1$A2c$_1$p$_{1-2}$c$_1$p$_{1-2}$A2c$_4$p$_{4-2}$c$_1^2$A2c$_1$A2c$_4$p$_{4-2}$A2c$_4^4$p$_{4-2}$A2p$_{4-2}$A2c$_4$p$_{4-2}$A2c$_4$p$_{4-2}$A2p$_{4-2}$A2c$_4$p$_{4-}$
$_2$A2c$_4^2$D2/20c$_3^3$A2p$_{4-2}$A2c$_4$p$_{4-2}$A2c$_4$p$_{4-2}$A2c$_4^3$p$_{4-2}$A2c$_4$p$_{4-2}$A2p$_{4-2}$A2c$_4^5$p$_{4-2}$A2p$_{4-2}$A2c$_4^3$p$_{4-2}$A2p$_{4-2}$A2p$_{4-}$
$_2$A2p$_{4-2}$A2c$_4^2$p$_{4-2}$A2c$_4$p$_{4-2}$A2c$_4^2$p$_{4-2}$A2c$_4^2$p$_{4-2}$A2p$_{4-2}$A2c$_4$p$_{4-2}$c$_1^3$p$_{1-2}$A2p$_{4-2}$c$_1$p$_{1-2}$A2c$_4^3$p$_{4-2}$A2c$_4$p$_{4-2}$A2p$_{4-}$
$_2$c$_4$p$_{4-1}$D2/20c$_3$p$_{3-1}$c$_4^3$p$_{4-1}$D2/20c$_3^3$p$_{3-1}$c$_4^2$p$_{4-2}$c$_1$A2c$_4^2$p$_{4-2}$A2c$_4^4$p$_{4-1}$c$_4^2$p$_{4-2}$A2c$_4^4$p$_{4-2}$p$_{1-2}$A2p$_{4-1}$c$_4^3$p$_{4-2}$p$_{1-}$
$_2$A2p$_{4-1}$c$_4^3$p$_{4-1}$c$_4^2$D2/20c$_4$p$_{4-1}$c$_4^5$p$_{4-1}$c$_4^2$p$_{4-2}$A2c$_4$D2/20c$_4$B2A2c$_4$c$_1^2$A2p$_{4-}$
$_2$A2c$_4$D2/20C2B2A2c$_4^4$D2/20c$_3$A2c$_4$c$_2$B2A2c$_4$D2/20c$_3$A2c$_4^3$p$_{4-1}$p$_{4-2}$A2p$_{4-1}$c$_1$p$_{1-2}$A2c$_4^2$p$_{4-1}$p$_{4-}$

$_{2}$A2$c_4^3p_{4-1}p_{4-2}$A2$c_4$B2A2$c_4^3$D2/20$c_3$A2$c_4c_2^2p_{2-4}c_4^2p_{4-2}$A2$c_4$D2/20C2B2A2$c_4p_{4-1}c_4$D2/20$c_3^2p_{3-}$
$_{4}c_3$A2$p_{4-1}c_1c_3$A2$c_4^3p_{4-1}c_4^3p_{4-1}c_4^3$D2/20$c_3$A2$c_4^2$D2/20$c_3$A2$c_4^3$D2/20$c_3p_{1-3}c_2p_{2-3}$B2$c_1^3$A2$c_4^3$D2/20$c_3p_{1-}$
$_{3}c_2p_{2-3}$B2$c_1^4c_3$A2$c_4$D2/20$c_3$A2$c_4$D2/20$c_3$C2$c_2p_{2-3}$B2A2$c_4^2p_{4-2}$A2$c_4^3$D2/20$c_3p_{1-3}c_2p_{2-3}$B2$c_1^3p_{1-3}c_2p_{2-}$
$_{3}$B2$c_1^4c_3$A2$c_4$D2/20$c_3$A2$c_1c_3$A2$c_4$D2/20C2B2A2$c_4^2p_{4-2}$A2$c_4^3p_{4-1}c_4^3p_{4-2}$A2$p_{4-2}$A2$c_4^2p_{4-2}$A2$c_4p_{4-}$
$_{2}$A2$c_4^7p_{4-2}$A2$c_4$D2/20$c_3^4$A2$c_1p_{1-2}$A2$c_4p_{4-1}p_{4-2}$A2$c_4p_{4-1}c_4p_{4-1}c_4^3p_{4-2}$A2$c_4p_{4-1}c_4^2p_{4-1}c_4^3p_{4-1}c_4^3p_{4-2}$A2$c_4p_{4-}$
$_{1}c_4^3p_{4-1}c_4^3p_{4-1}c_4^3p_{4-2}$A2$c_4p_{4-1}p_{4-2}$A2$c_4^2p_{4-2}$A2$c_4^3p_{4-1}c_4p_{4-1}c_4^3p_{4-1}c_4^2p_{4-2}$A2$c_4^2p_{4-2}$A2$c_4p_{4-1}c_4^2p_{4-2}$A2$c_4p_{4-}$
$_{1}c_4^5$D2/20$c_4$B2A2$c_4p_{1-2}c_2c_4^2$D2/20$p_{3-1}p_{4-2}c_2^3p_{2-4}c_4$B2A2$c_4c_1^2$C2B2A2$c_4^4$D2/20$c_3$A2$c_4^2p_{4-}$
$_{2}$A2$c_4$D2/20C2B2A2$c_4^4$D2/20$c_4c_2p_{2-3}c_1^2p_{3-4}$C2B2A2$c_4^4c_1^2$C2B2A2$p_{4-2}c_4p_{4-}$
$_{2}$A2$c_4$D2/20C2B2A2$c_4^4p_{4-2}$A2$c_4$D2/20$c_3$B2A2$c_4^3$D2/20$c_3$A2$c_4c_2p_{2-}$
$_{3}$B2A2$c_4^2$D2/20$c_3$A2$c_4$B2A2$c_4^2p_{4-2}$A2$c_4$D2/20C2B2A2$c_4p_{4-1}c_4^2$D2/20$c_3$A2$p_{4-1}c_1$A2$c_4^2p_{4-2}$A2$c_4p_{4-}$
$_{1}c_4^3p_{4-1}c_4^4p_{4-1}c_4p_{4-1}c_4^2p_{4-1}^2c_4p_{4-2}$A2$c_4p_{4-1}c_4p_{4-1}c_4p_{4-2}$A2$c_4^3p_{4-1}^3c_1c_3^2p_{3-4}c_3$A2$c_4c_2p_{2-3}$B2A2$c_4p_{4-1}c_4p_{4-}$
$_{2}$A2$c_4^2$D2/20$c_2p_{4-1}^2c_1^3c_3$A2$c_4c_2p_{2-3}$B2A2$c_4p_{4-2}$A2D2/20C2B2$c_1$A2$c_4$D2/20$p_{2-4}c_3$A2$p_{4-1}c_1c_3$A2$c_4^3p_{4-}$
$_{2}$A2$c_4$D2/20C2B2A2$c_4$D2/20$p_{2-4}p_{4-2}$A2$p_{4-1}c_1c_3$A2$c_4^3p_{4-1}p_{4-2}$A2$c_4^3p_{4-2}$A2$c_4p_{4-1}p_{4-}$
$_{2}$A2$c_4$D2/20C2B2A2$p_{4-1}p_{4-2}$A2$c_4^3p_{4-2}$A2$c_4p_{4-1}p_{4-2}$A2$c_4$D2/20C2B2A2$p_{4-}$
$_{1}c_4$D2/20$c_3$A2$c_4^2$D2/20C2B2A2$c_4^4$B2A2$c_4^3$D2/20$c_3$A2$c_4c_2$B2A2$p_{4-1}c_1c_3$A2$c_4^3p_{4-}$
$_{2}$A2$c_4$D2/20C2B2A2$c_4^3$B2A2$c_4^3$D2/20$c_3$A2$c_4c_2^3p_{2-4}$C2B2A2$c_4^4$B2A2$c_4^3$D2/20$c_3$A2$c_4c_2$B2A2$p_{4-}$
$_{1}c_1c_3$A2$c_4^3p_{4-2}$A2$c_4$D2/20C2B2A2$c_4^3$B2A2$c_4^2p_{4-2}$A2$c_4$D2/20C2B2A2$c_4^3$B2A2$c_4^3$D2/20$c_3$A2$c_4c_2p_{2-}$
$_{3}$B2A2$c_4^3$D2/20$c_3$A2$c_4c_2$B2A2$p_{4-1}c_1c_3$A2$c_4^3p_{4-2}$A2$c_4$D2/20C2B2A2$c_4^2$B2A2$c_4^3c_2$B2A2$p_{4-}$
$_{1}c_1c_3$A2$c_4p_{4-2}$A2$c_4$D2/20C2B2A2$c_4p_{4-1}c_4$D2/20$c_3$A2$c_4^3$D2/20C2B2A2$c_4^4c_2^3p_{2-}$
$_{3}$B2A2$c_4^3$D2/20$c_3$A2$c_4c_2$B2A2$p_{4-1}c_1c_3$A2$c_4^3p_{4-2}$A2$c_4c_2$B2A2$p_{4-1}c_1c_3$A2$c_4^3p_{4-}$
$_{2}$A2$c_4$D2/20C2B2A2$c_4^4$B2A2$c_4p_{4-2}$A2$p_{4-1}c_1c_3p_{3-4}c_3$A2$c_4^3p_{4-1}p_{4-2}$A2$c_4$D2/20C2B2A2$c_4p_{4-1}c_1p_{1-}$
$_{3}$B2A2$c_4^3$D2/20$c_3$A2$c_4c_2$B2A2$p_{4-1}c_1$C2B2A2$c_4^2$D2/20$c_3$A2$c_4c_2p_{2-4}c_3$A2$c_4c_2p_{2-4}c_3$A2$c_4c_2$B2A2$p_{4-}$
$_{1}c_1c_3$A2$c_4^3c_2$B2A2$p_{4-1}c_1c_3$A2$c_4^5p_{4-1}c_1c_3$A2$c_4^3p_{4-1}c_1$A2$c_1p_{1-2}$A2$c_4^2p_{4-2}$A2$c_4$D2/20C2B2A2$c_4p_{4-}$
$_{1}c_4$D2/20$c_3^2$A2$c_4$D2/20C2B2A2$c_4p_{4-1}c_1$A2$c_1p_{1-2}$A2$c_4^2p_{4-2}$A2$c_4$D2/20C2B2A2$c_4p_{4-}$
$_{1}c_4$D2/20$c_3$A2$c_4^2$D2/20$c_3$A2$p_{4-1}c_1c_3$A2$c_4^3p_{4-2}$A2$c_4$D2/20C2B2A2$c_4p_{4-1}c_4$D2/20$c_3$A2$c_4^2$D2/20$c_3$A2$p_{4-}$
$_{1}c_1c_3$A2$c_4^3$D2/20C2B2A2$c_4$D2/20C2B2A2$p_{4-2}$A2D2/20C2B2A2$c_4p_{4-1}c_1c_3$A2$c_4^3p_{4-1}p_{4-}$
$_{2}$A2$c_4$D2/20C2B2A2$c_4p_{4-1}c_1$A2$c_1c_3$A2$c_4^3p_{4-1}p_{4-2}$A2$c_4$D2/20C2B2A2$c_4p_{4-1}c_1c_3$A2$c_4c_2^2p_{2-4}c_3$A2$c_4^3p_{4-}$
$_{1}p_{4-2}$A2$c_4$D2/20C2B2A2$c_4^2p_{4-1}c_1c_3$A2$c_4c_2^2c_4p_{4-1}^2c_1$C2B2A2$c_4p_{4-1}c_1p_{3-1}c_4c_2p_{2-}$
$_{3}$B2A2$c_4^2$B2A2$c_4^2c_2$B2A2$p_{4-1}c_1c_3$A2$p_{4-2}$A2$c_4c_2$B2A2$p_{4-1}c_1c_3$A2$c_4^2p_{4-1}p_{4-}$
$_{2}$A2D2/20C2B2A2$c_4$D2/20C2B2A2$p_{4-2}$A2$c_4^3p_{4-1}p_{4-2}$A2D2/20C2B2A2$c_4^3p_{4-1}p_{4-}$
$_{2}$A2$c_4$D2/20C2B2A2$c_4$D2/20C2B2A2$p_{4-2}$A2$c_4^3$D2/20C2B2A2$c_4$D2/20C2B2A2$p_{4-2}$A2$c_4^3p_{4-1}p_{4-}$
$_{2}$A2$c_4$D2/20C2B2A2$c_4$D2/20C2B2A2$p_{4-2}$A2$c_4^2p_{4-1}p_{4-2}$A2$c_4$D2/20C2B2A2$c_4$D2/20C2B2A2$p_{4-}$
$_{2}$A2$c_4^2p_{4-1}c_1p_{3-1}c_4c_2^4p_{2-4}c_3$A2$c_4c_2$B2A2$p_{4-1}c_1c_3$A2$c_4^3p_{4-1}p_{4-}$
$_{2}$A2$c_4^2$D2/20$c_3$A2$c_4$D2/20C2B2A2$c_4$D2/20C2B2A2$p_{4-2}$A2$c_4^3p_{4-1}p_{4-}$
$_{2}$A2$c_4$D2/20C2B2A2$c_4$D2/20C2B2A2$p_{4-2}$A2$c_4^3p_{4-2}$A2$p_{4-1}c_1p_{3-1}p_{4-1}c_1$C2B2A2$c_4^2$D2/20$c_3^2$A2$p_{4-}$
$_{1}c_1c_3$A2$c_4^3p_{4-1}p_{4-2}$A2$c_4^2p_{4-1}c_1p_{3-1}c_4c_2p_{2-3}$B2A2$c_4^4$D2/20$c_3$A2$c_4c_2$B2A2$p_{4-1}c_1^4$A2$p_{4-}$
$_{2}$A2$c_4$D2/20C2B2A2$c_4^2p_{4-1}p_{4-2}$A2$c_4$D2/20C2B2A2$c_4^3p_{4-1}p_{4-2}$A2$c_4c_2^3c_4p_{4-2}$A2$p_{4-1}c_1$C2B2A2$c_4p_{4-}$
$_{2}$A2$p_{4-1}c_1$C2B2A2$c_4^2$D2/20$c_3$C2$c_2^3p_{4-2}$A2$c_4$D2/20$c_3p_{3-1}c_1c_3$A2$c_4c_2c_4p_{4-2}$A2$p_{4-1}c_1$C2B2A2$c_4p_{4-1}c_1p_{3-}$
$_{1}c_1^2$A2$c_4p_{4-1}c_1c_3p_{3-1}c_4p_{4-1}c_1$A2$c_1c_3p_{3-1}c_4p_{4-1}c_1c_3p_{3-1}c_4p_{4-1}c_1$A2$c_1c_3p_{3-1}c_1$A2$c_1c_3p_{3-1}c_4p_{4-1}c_1c_3p_{3-}$
$_{1}c_1^2$A2$c_4p_{4-1}c_1c_3^2p_{3-4}c_3^4p_{3-1}c_1c_3^4$A2$c_4^2p_{4-1}$A2$c_1c_3^2p_{3-1}c_1^2$A2$c_4$D2/20$c_3^2p_{3-4}c_3^3p_{3-1}c_1p_{1-2}$A2$p_{4-1}c_4p_{4-}$
$_{2}$A2$c_1^3p_{1-2}$A2$c_4^2p_{4-1}$A2$c_1c_3^2p_{3-1}c_1^2$A2$c_4$D2/20$c_3^2p_{3-4}c_3^3p_{3-1}c_1p_{1-2}$A2$p_{4-1}c_4p_{4-2}$A2$c_1^3$A2$c_1c_3^2p_{3-}$
$_{4}^{2}$B2A2$p_{4-1}c_4^2p_{4-2}$A2$c_1^2p_{1-2}$A2$c_1^2p_{1-2}$A2$c_1^3$A2$c_1$A2$c_1^2$A2$c_4$D2/20$c_3^2p_{3-4}c_3^3p_{3-1}c_1p_{1-2}$A2$p_{4-1}c_4^2p_{4-2}$A2$p_{4-}$
$_{1}c_1^3$A2$c_1c_3^2p_{3-4}^2$B2A2$p_{4-1}c_4^2p_{4-2}$A2$c_1^2p_{1-2}$A2$c_1^3$A2$c_1^2$A2$c_4$D2/20$c_3^2p_{3-4}c_3^3p_{3-1}c_1p_{1-2}$A2$p_{4-1}c_4^2p_{4-2}$A2$p_{4-}$
$_{1}c_1^2p_{1-2}$A2$c_4$D2/20$c_3p_{3-1}p_{1-2}c_4^2p_{4-2}p_{4-1}p_{1-2}$A2$c_4^2p_{4-2}$A2$p_{4-1}p_{1-2}$A2$c_4$D2/20$c_3p_{3-1}c_1^3p_{1-2}$A2$p_{4-2}$A2$p_{4-1}c_1p_{1-}$
$_{2}$A2$c_4$D2/20$c_3p_{3-1}c_1^3p_{1-2}$A2$p_{4-1}p_{1-2}$A2$c_4^3p_{4-2}$A2$p_{4-1}p_{1-2}$A2$c_4$D2/20$c_3p_{3-1}c_1^3p_{1-2}$A2$p_{4-1}p_{1-2}c_1^2c_3^2$A2$p_{4-1}p_{1-}$
$_{2}$A2$c_4^2$D2/20$c_3p_{3-1}c_1^3c_3^2p_{4-1}p_{1-2}$A2$c_4^2$D2/20$c_3^2$A2$p_{4-2}$A2$c_4^2$D2/20$c_3p_{3-4}c_3^2$A2$p_{4-2}$A2$c_4^2$D2/20$c_3^2$A2$p_{4-}$
$_{2}$A2$c_4^2p_{4-2}$A2$c_4$D2/20$c_3$C2$p_{1-2}$A2$c_4p_{4-2}$A2$p_{4-1}c_1^2p_{1-2}$A2$c_4$D2/20$c_3^2$A2$p_{4-1}c_1c_3^2$A2$p_{4-1}c_1p_{1-2}$A2$p_{4-}$
$_{1}c_1c_3^2$A2$c_4^2p_{4-2}$A2$p_{4-1}c_1c_3^2$A2$c_4^2p_{4-2}$A2$p_{4-1}c_1^2p_{1-2}$A2$p_{4-1}c_1^3c_3^2$A2$p_{4-1}c_1^3c_3^2$A2$p_{4-1}c_1^5p_{4-2}$A2$p_{4-}$
$_{1}c_1^5c_4c_2p_{2-4}$C2B2A2$c_4^3$D2/20C2B2A2$c_4^5$D2/20$c_4$B2A2$c_4c_1p_{1-2}$A2$p_{4-}$
$_{1}c_1^3c_3^2$A2$c_4^3$D2/20$c_4$B2A2$c_4c_1p_{1-2}$A2$p_{4-1}c_1^3c_3^2$A2$p_{4-1}c_1c_3^2$A2$p_{4-1}c_1$A2$c_3$A2$p_{4-1}c_1^3c_3^2$A2$p_{4-1}c_1c_3^2$A2$p_{4-}$
$_{2}$A2$p_{4-1}p_{1-2}$A2$c_4^2p_{4-2}$A2$p_{4-1}c_1^5c_3$A2$p_{4-1}p_{1-2}$A2$c_4^2$D2/20$c_3^2$A2$p_{4-1}p_{1-2}$A2$c_4^5$D2/20$c_3^5p_{3-4}c_3^2$A2$p_{4-1}p_{1-}$

$_{2}A2c_4^{2}p_{4-1}c_4^{3}p_{4-2}c_1^{8}p_{1-2}A2c_4^{6}p_{4-2}A2c_4^{9}p_{4-2}A2c_4^{4}p_{4-1}p_{1-2}A2c_4^{3}p_{4-2}A2c_4D2/20c_3A2c_4^{6}p_{4-2}c_1^{3}p_{1-2}A2c_4^{4}p_{4-1}$
$p_{1-2}A2p_{4-2}A2c_4^{4}p_{4-1}c_1^{5}p_{1-2}A2c_4^{3}p_{4-1}p_{1-2}A2c_4^{5}p_{4-2}A2c_4^{5}p_{4-2}c_1^{3}p_{1-2}A2c_4^{10}p_{4-2}A2D2/20B2A2c_4^{3}p_{4-1}$
$c_4^{2}p_{4-2}A2p_{4-2}A2c_4^{8}p_{4-2}c_1^{2}p_{1-2}A2c_4^{4}p_{4-2}A2c_4^{6}p_{4-2}A2c_4^{2}p_{4-1}p_{1-2}A2c_4^{3}p_{4-2}c_1^{3}p_{1-2}A2c_4^{3}p_{4-2}c_1^{3}p_{1-2}$
$A2c_4^{10}p_{4-2}A2c_4^{3}p_{4-1}c_4^{2}p_{4-2}A2p_{4-2}A2c_4^{5}p_{4-2}c_1A2c_4^{2}p_{4-2}A2c_4^{6}p_{4-2}A2c_4^{2}p_{4-1}^{2}p_{1-2}A2c_4^{12}p_{4-2}A2c_4^{3}p_{4-2}$
$A2c_4p_{4-2}A2c_4^{12}p_{4-2}A2c_4^{13}p_{4-2}A2c_4^{5}D2/20c_3p_{3-4}c_3A2c_4^{2}D2/20c_3A2c_4^{2}D2/20c_3A2c_4^{3}D2/20c_3^{2}p_{3-4}$
$c_3^{2}A2D2/20c_3^{3}A2c_4^{3}p_{4-1}c_4p_{4-2}A2c_4^{3}D2/20c_3^{2}A2p_{4-1}c_4^{3}p_{4-2}A2p_{4-1}c_4^{5}p_{4-1}c_4p_{4-2}A2c_4^{3}D2/20c_3^{2}A2c_4^{4}p_{4-1}$
$c_4^{3}p_{1-2}A2p_{4-1}c_4^{3}p_{1-2}A2c_4^{3}p_{4-1}c_4^{3}p_{4-2}A2c_4^{3}p_{4-2}A2p_{4-1}c_4^{2}p_{4-2}A2p_{4-1}c_4^{8}p_{4-2}A2p_{4-2}A2c_4^{2}p_{4-1}c_4p_{4-2}A2c_4^{4}p_{4-1}$
$c_4p_{4-2}A2c_4^{6}p_{4-1}c_4p_{4-2}A2c_4^{2}p_{4-2}A2c_4^{4}p_{4-1}c_4c_1^{4}A2c_4p_{4-2}A2c_4^{6}p_{4-1}c_4p_{4-2}A2c_4^{4}p_{4-1}c_4^{4}p_{4-2}A2p_{4-1}c_4^{4}p_{4-1}$
$c_4^{5}p_{4-2}c_4^{4}p_{4-2}A2p_{4-1}c_4^{4}p_{4-1}c_4^{5}p_{4-2}D2/20C2A2c_4^{2}p_{4-2}A2c_4^{4}p_{4-2}A2p_{4-1}c_4^{9}D2/20c_3^{3}p_{3-4}$
$c_3^{12}c_4^{12}D2/20c_3^{11}p_{3-1}c_4p_{4-2}A2c_4^{2}p_{4-2}A2p_{4-2}c_4^{2}p_{4-2}c_2^{3}B2^{2}c_1^{6}p_{1-2}A2p_{4-2}c_4p_{4-2}c_2B2c_4^{4}D2/20c_3^{6}p_{3-4}$
$c_3^{3}c_4^{15}p_{4-2}c_3c_4^{6}p_{4-2}c_3c_4^{9}p_{4-2}c_4D2/20c_2^{14}B2c_4^{2}p_{4-2}c_4^{10}D2/20c_3^{10}c_4^{23}D2/20c_3^{4}c_4D2/20c_2^{14}B2c_4^{2}p_{4-2}$
$c_4^{10}D2/20c_3^{9}p_{3-4}c_3^{10}c_4^{23}D2/20c_3^{6}c_4^{21}p_{1-2}c_4^{3}p_{4-2}c_1^{14}p_{1-2}c_4^{5}D2/20c_3^{9}c_1^{5}p_{1-3}c_1^{8}p_{1-2}A2c_4^{4}p_{4-1}c_4p_{4-2}$
$A2c_4^{3}p_{4-1}c_4p_{4-2}A2c_4^{2}p_{4-1}c_4c_1^{4}A2c_4p_{4-2}A2c_4^{4}c_1^{4}A2c_4^{4}p_{4-1}c_4^{2}p_{4-2}A2c_4^{4}p_{4-2}A2p_{4-1}c_4^{3}p_{4-2}A2p_{4-1}c_4^{3}p_{4-2}$
$A2p_{4-1}c_4^{4}p_{4-1}c_4^{5}p_{4-2}D2/20C2A2c_4^{2}p_{4-2}A2c_4^{2}p_{4-2}A2p_{4-2}A2c_4^{2}p_{4-2}A2p_{4-2}A2c_4^{6}D2/20c_3^{3}p_{3-4}$
$c_3^{9}c_4^{12}D2/20c_3^{11}p_{3-1}c_4p_{4-2}A2c_4^{2}p_{4-2}A2p_{4-2}c_4^{2}p_{4-2}c_2^{3}B2c_2^{3}p_{2-4}c_3^{8}A2p_{4-2}c_4p_{4-2}c_2B2c_4^{4}D2/20c_3^{6}p_{3-4}$
$c_3^{3}p_{3-1}c_1^{2}c_3^{9}c_4^{5}p_{4-2}c_3c_4^{6}p_{4-2}c_3c_4^{9}p_{4-2}c_4D2/20c_2^{14}B2c_4^{25}D2/20c_3^{10}c_4^{16}p_{1-2}c_4^{2}D2/20c_2B2p_{1-2}c_4^{9}c_1^{4}p_{1-2}$
$c_4^{5}D2/20c_3^{11}c_1^{4}p_{1-3}c_1^{13}p_{1-3}c_1^{10}p_{1-3}c_1^{8}p_{1-3}c_1^{5}p_{1-2}c_4^{5}D2/20c_2^{2}B2c_4^{4}D2/20c_2^{2}B2c_4^{3}p_{4-2}c_4p_{4-2}c_4^{13}p_{4-2}$
$c_4^{4}p_{4-2}A2c_4^{2}p_{4-2}c_4^{10}p_{4-2}c_2^{2}p_{2-4}c_3^{6}A2p_{4-2}c_4p_{4-2}c_2B2c_4^{6}c_1c_3^{13}A2c_4p_{4-2}A2p_{4-2}c_4p_{4-2}c_2B2c_4^{7}D2/20c_3p_{3-4}$
$c_2^{7}c_1^{2}c_3^{10}c_4^{8}p_{4-2}c_4^{5}p_{4-2}c_4^{4}p_{4-2}c_4^{5}p_{4-2}c_4^{7}c_1^{4}p_{1-2}c_1^{4}p_{1-2}c_4^{3}c_3^{3}p_{3-4}c_2^{7}p_{2-4}c_2^{26}p_{2-3}c_1^{7}B2D2/20c_2^{10}p_{2-3}$
$c_1^{3}c_3^{2}C2c_1^{4}p_{1-3}c_1^{59}c_2^{8}B2c_4^{6}D2/20c_2^{5}B2c_4^{3}p_{4-2}B2c_4p_{4-1}c_3^{6}C2c_2^{2}B2c_4c_2^{3}B2c_4^{2}p_{4-2}c_4c_2^{3}B2c_4^{2}p_{4-2}$
$c_4c_2^{12}p_{2-3}C2c_4D2/20c_3^{6}C2^{2}c_4D2/20p_{3-4}c_4D2/20C2c_4D2/20$

### cen3

$p_{1-4}p_{6-4}^{51}p_{6-8}p_{10-4}p_{6-4}^{6}p_{6-3}p_{6-4}^{4}p_{6-17}p_{16-4}p_{6-4}^{2}c_5^{78}p_{7-4}c_5^{2}p_{7-5}c_6c_{11}^{5}c_{17}^{14}p_{4-12}c_{13}^{22}p_{14-5}c_6^{11}p_{14-5}c_6^{2}p_{8-5}c_6^{2}p_{8-5}$
$c_6^{2}p_{8-5}c_6p_{8-13}c_{14}^{6}p_{1-4}p_{7-13}c_{14}^{2}p_{1-4}c_5^{29}p_{7-10}c_{11}^{109}p_{4-11}c_{12}^{7}p_{1-6}p_{5-6}c_7p_{5-11}p_{1-6}p_{5-11}p_{1-11}c_{12}p_{1-6}p_{5-6}c_7p_{5-6}c_7p_{5-11}$
$c_{12}p_{1-6}p_{5-6}c_7^{6}p_{8-6}c_7p_{8-2}c_3^{6}p_{8-2}c_3^{2}p_{8-6}c_7^{10}p_{8-6}^{2}c_7^{2}p_{8-6}^{2}p_{8-10}p_{15-6}p_{8-6}^{7}p_{8-16}p_{9-6}p_{8-6}^{29}p_{8-4}p_{8-6}^{11}p_{8-4}^{3}p_{8-6}p_{8-4}^{5}$
$p_{8-6}p_{8-4}p_{8-6}^{4}p_{8-4}p_{8-6}p_{8-4}p_{8-6}^{7}C3F3p_{8-12}p_{1-3}F3p_{8-12}p_{1-3}F3p_{8-6}C3F3p_{8-12}p_{1-3}F3p_{8-12}p_{1-3}F3p_{8-6}C3F3p_{8-12}$
$p_{1-3}p_{6-4}F3C3F3H3p_{10-12}A3F3p_{8-3}F3C3F3p_{8-9}p_{11-12}p_{1-3}F3C3F3H3p_{10-12}A3C3p_{6-4}F3C3F3H3p_{10-13}p_{1-4}$

### cen4

$N4p_{16-19}p_{5-14}H4/9p_{3-14}H4/9p_{4-14}H4/9p_{10-14}H4/9p_{3-14}H4/9p_{3-14}H4/9p_{4-14}H4/9p_{4-14}H4/9J4p_{4-14}H4/9p_{4-14}$
$H4/9p_{10-14}H4/9p_{3-14}H4/9p_{3-14}H4/9p_{4-14}H4/9p_{3-14}H4/9c_3^{3}p_{7-10}p_{18-2}G4p_{15-2}p_{7-10}p_{18-11}p_{19-5}c_2^{2}p_{2-7}$
$c_{15}^{5}O4p_{1-10}c_{18}p_{18-5}p_{13-5}^{4}p_{13-4}^{2}p_{13-5}p_{13-4}p_{13-5}p_{13-4}c_{13}p_{13-1}c_6p_{6-1}c_6^{3}p_{6-18}c_1A4c_6^{3}p_{6-11}c_{19}^{4}p_{19-10}R4.1p_{12-10}$
$R4.1p_{12-3}p_{11-1}c_6p_{6-1}c_6^{2}p_{6-18}c_1p_{1-18}c_1A4p_{6-18}c_1A4c_6^{3}p_{6-13}G4p_{15-2}c_7p_{7-2}p_{7-10}p_{18-2}p_{7-2}p_{7-10}p_{18-2}p_{7-2}^{2}$
$c_7^{4}c_{14}^{5}p_{14-10}p_{18-2}p_{7-1}c_6p_{6-1}^{4}p_{6-2}p_{7-5}p_{13-14}c_{14}p_{14-12}c_{12}L4c_{12}L4c_{12}^{2}p_{12-2}c_7p_{7-2}^{2}c_7^{17}p_{7-19}c_{19}^{9}p_{19-14}H4/9p_{16-14}$
$H4/9c_{16}^{2}p_{16-14}H4/9c_{16}p_{16-6}C4c_{17}p_{17-6}C4p_{17-14}H4/9p_{16-1}c_6^{4}c_{13}p_{13-2}p_{7-2}p_{7-18}c_1A4p_{6-10}p_{18-2}c_7^{2}p_{7-2}p_{7-10}$
$p_{18-2}p_{7-2}c_7p_{7-2}c_7p_{7-2}p_{9-2}c_9p_{9-18}c_1^{3}p_{1-14}H4/9p_{16-6}c_{14}^{3}p_{14-6}c_{14}^{2}p_{14-6}c_{14}p_{14-1}p_{6-1}^{4}p_{6-2}c_7^{2}p_{7-2}c_7p_{7-12}c_{12}p_{12-2}p_{7-1}$
$c_6p_{6-2}p_{7-2}p_{7-16}c_{10}^{26}p_{10-19}c_{19}^{14}p_{1-14}H4/9p_{16-14}H4/9p_{16-14}H4/9c_{16}^{6}p_{16-14}H4/9p_{16-14}H4/9p_{16-11}G4p_{10-11}p_{13-14}$
$H4/9p_{16-17}R4.1A4p_{6-14}H4/9p_{16-17}p_{4-11}p_{13-14}H4/9p_{16-17}p_{4-14}H4/9p_{16-17}p_{4-14}H4/9p_{16-17}p_{4-14}H4/9p_{16-18}p_{5-14}$
$H4/9p_{16-17}p_{4-14}H4/9p_{16-18}p_{5-14}H4/9p_{16-17}p_{4-14}H4/9p_{16-18}p_{5-14}H4/9p_{16-17}p_{4-14}H4/9p_{16-17}c_4^{3}R4.1p_{12-10}$
$R4.1c_{12}p_{12-10}R4.1c_{12}^{2}p_{12-2}c_8^{3}p_{8-10}R4.1p_{12-10}R4.1c_{12}^{2}p_{12-10}R4.1c_{12}^{6}p_{12-10}R4.1p_{12-18}c_5^{2}p_{5-10}R4.1p_{12-18}$
$p_{5-10}R4.1p_{12-4}p_{12-18}p_{5-8}R4.1c_{12}^{8}p_{12-10}R4.1c_{12}p_{12-10}R4.1c_{12}^{4}p_{12-10}R4.1c_{12}^{2}p_{12-10}R4.1c_{12}^{2}p_{12-10}$
$R4.1c_{12}^{2}p_{12-10}R4.1c_{12}p_{12-10}R4.1p_{12-10}R4.1c_{12}^{3}p_{12-17}c_3^{16}p_{3-17}p_{19-17}c_{19}^{53}p_{19-12}c_2p_{2-12}c_2^{2}p_{2-18}p_{1-3}c_{17}p_{17-18}$
$p_{1-3}c_{17}p_{17-18}p_{1-3}c_{17}^{2}p_{17-18}p_{1-3}c_{17}p_{17-18}c_1p_{1-3}c_{11}p_{11-18}p_{1-3}c_{17}p_{17-18}p_{1-3}c_{17}^{2}p_{17-18}p_{1-3}c_{17}p_{17-18}p_{1-3}c_{17}^{6}p_{17-18}$
$p_{1-3}c_{17}p_{17-18}p_{1-3}p_{17-10}R4.1c_{12}^{2}p_{12-18}p_{1-3}c_{17}p_{17-18}p_{1-3}p_{17-10}R4.1c_{12}p_{12-18}p_{1-3}c_{17}^{4}p_{17-18}p_{1-3}c_{17}p_{17-18}p_{1-3}p_{17-10}$
$R4.1c_{12}^{2}p_{12-18}p_{1-3}c_{17}p_{17-18}p_{1-3}p_{17-10}R4.1c_{12}p_{12-18}p_{1-3}c_{17}^{8}p_{17-4}c_{12}p_{12-18}p_{1-3}c_{17}p_{17-18}p_{1-3}c_{17}^{4}p_{17-18}p_{1-3}$
$c_{17}p_{17-18}p_{1-3}p_{17-10}R4.1p_{12-3}c_{17}p_{17-18}p_{1-3}c_{17}^{7}p_{17-18}p_{1-3}c_{17}p_{17-18}p_{1-3}p_{17-10}R4.1c_7^{3}p_{7-18}p_{1-3}c_{17}p_{17-18}p_{1-3}p_{17-}$

$_{10}$R4.1p$_{12-3}$c$_{17}$p$_{17-18}$p$_{1-3}$c$_{17}^{4}$p$_{17-18}$c$_{1}^{3}$p$_{1-18}$c$_{1}^{9}$p$_{1-18}$p$_{1-3}$c$_{17}^{15}$p$_{17-18}$c$_{1}^{13}$p$_{1-18}$p$_{1-3}$c$_{17}$p$_{17-18}$p$_{1-3}$p$_{17-3}$c$_{16}^{2}$p$_{16-18}$p$_{1-3}$c$_{17}$p$_{17-18}$p$_{1-3}$c$_{17}^{5}$p$_{17-18}$p$_{1-3}$c$_{17}$p$_{17-18}$c$_{1}^{2}$p$_{1-18}$p$_{1-3}$c$_{17}^{2}$p$_{17-18}$c$_{1}^{2}$p$_{1-18}$p$_{1-3}$c$_{17}^{5}$p$_{17-18}$p$_{1-3}$c$_{17}^{3}$p$_{17-18}$p$_{1-3}$c$_{17}$p$_{17-18}$p$_{1-3}$p$_{17-5}$p$_{16-18}$p$_{1-3}$c$_{17}^{3}$p$_{17-18}$p$_{1-3}$c$_{17}^{2}$p$_{17-18}$p$_{1-3}$c$_{17}^{6}$p$_{17-18}$p$_{1-3}$c$_{17}$p$_{17-18}$p$_{1-3}$c$_{17}^{2}$p$_{17-3}$p$_{11-4}$c$_{12}$p$_{12-18}$p$_{1-3}$c$_{17}$p$_{17-18}$p$_{1-3}$c$_{17}$p$_{17-18}$p$_{1-3}$c$_{17}$p$_{17-18}$p$_{1-3}$c$_{16}^{7}$p$_{16-18}$p$_{1-3}$c$_{17}$p$_{17-18}$p$_{1-3}$c$_{17}$p$_{17-18}$p$_{1-3}$c$_{17}^{4}$p$_{17-18}$p$_{1-3}$c$_{17}$p$_{17-18}$p$_{1-3}$c$_{17}$p$_{17-18}$p$_{1-3}$p$_{17-10}$R4.1p$_{12-3}$c$_{17}^{2}$p$_{17-18}$c$_{1}^{12}$p$_{1-18}$p$_{1-3}$c$_{17}^{5}$p$_{17-18}$p$_{1-3}$c$_{17}^{2}$p$_{17-18}$p$_{1-3}$c$_{17}$p$_{17-18}$p$_{1-3}$c$_{17}$p$_{17-18}$p$_{1-3}$c$_{17}$p$_{17-18}$p$_{1-3}$c$_{17}^{3}$p$_{17-18}$p$_{1-3}$c$_{17}$p$_{17-18}$p$_{1-3}$c$_{10}$p$_{10-18}$p$_{1-3}$c$_{17}$p$_{17-3}$c$_{17}^{2}$p$_{17-18}$p$_{1-15}$c$_{17}^{2}$p$_{17-18}$A4p$_{2-18}$p$_{1-3}$c$_{17}$p$_{17-18}$c$_{1}$p$_{1-18}$p$_{1-3}$c$_{17}$p$_{17-18}$p$_{1-3}$c$_{17}$p$_{17-18}$c$_{1}^{2}$p$_{1-18}$p$_{1-3}$c$_{17}^{4}$p$_{17-18}$p$_{1-3}$c$_{17}$p$_{17-18}$p$_{1-3}$c$_{17}$p$_{17-18}$p$_{1-3}$p$_{17-3}$c$_{17}^{3}$p$_{17-18}$p$_{1-3}$c$_{17}^{2}$p$_{17-18}$p$_{1-3}$c$_{17}^{6}$p$_{17-18}$p$_{1-3}$c$_{17}$p$_{17-18}$p$_{1-3}$c$_{17}^{2}$p$_{17-5}$c$_{2}^{2}$p$_{2-18}$p$_{1-3}$c$_{17}$p$_{17-18}$p$_{1-3}$c$_{17}$p$_{17-18}$p$_{1-3}$c$_{17}$p$_{17-18}$A4p$_{2-18}$p$_{1-3}$p$_{17-3}$c$_{17}^{3}$p$_{17-18}$p$_{1-3}$c$_{17}^{12}$p$_{17-18}$p$_{1-3}$c$_{17}^{3}$p$_{17-18}$p$_{1-18}$p$_{1-3}$c$_{17}$p$_{17-18}$p$_{1-3}$c$_{17}$p$_{17-18}$p$_{1-3}$p$_{17-19}$p$_{5-18}$p$_{1-3}$p$_{17-19}$p$_{5-18}$p$_{1-3}$c$_{17}$p$_{17-18}$p$_{1-3}$c$_{17}$p$_{17-18}$p$_{1-3}$c$_{17}^{2}$p$_{17-18}$p$_{1-18}$p$_{1-11}$p$_{1-5}$p$_{13-5}$c$_{13}$p$_{13-18}$p$_{1-11}$p$_{1-5}$p$_{13-5}$c$_{13}$p$_{13-18}$p$_{1-11}$p$_{1-5}$c$_{13}$p$_{16-12}^{5}$p$_{16-10}$R4.1p$_{12-15}$p$_{1-12}$p$_{16-12}$c$_{16}$p$_{16-12}$p$_{6-10}$R4.1c$_{12}^{4}$p$_{12-10}$R4.1p$_{12-10}$R4.1p$_{12-18}$c$_{1}$p$_{1-18}$p$_{1-3}$p$_{17-19}$p$_{5-18}$p$_{1-3}$c$_{17}$p$_{17-18}$p$_{1-3}$c$_{17}^{2}$p$_{17-18}$p$_{1-18}$p$_{1-11}$p$_{1-5}$c$_{13}$p$_{16-12}^{9}$p$_{16-10}$R4.1c$_{12}^{3}$L4p$_{6-10}$R4.1c$_{12}$L4p$_{6-10}$R4.1c$_{12}^{2}$p$_{12-10}$R4.1p$_{12-10}$R4.1c$_{12}^{2}$p$_{12-18}$p$_{1-3}$c$_{17}$p$_{17-18}$p$_{1-3}$c$_{17}$p$_{17-18}$p$_{1-3}$c$_{17}$p$_{17-18}$p$_{1-18}$c$_{1}$p$_{1-18}$p$_{1-3}$c$_{17}$p$_{17-18}$p$_{1-3}$c$_{17}$p$_{17-18}$p$_{1-3}$c$_{17}$p$_{17-18}$p$_{1-3}$p$_{17-19}$p$_{5-18}$p$_{1-3}$c$_{17}^{2}$p$_{17-18}$p$_{1-4}$S4p$_{5-18}$p$_{1-11}$p$_{1-5}$c$_{13}$p$_{16-11}$p$_{1-5}$c$_{2}^{2}$p$_{2-10}$R4.1p$_{12-10}$R4.1p$_{12-10}$R4.1p$_{12-10}$R4.1p$_{12-10}$R4.1p$_{12-10}$R4.1p$_{12-10}$R4.1p$_{12-10}$R4.1p$_{12-10}$R4.1p$_{12-10}$R4.1p$_{12-10}$R4.1p$_{12-10}$R4.1p$_{12-10}$R4.1p$_{12-10}$R4.1p$_{12-10}$R4.1p$_{12-10}$R4.1p$_{12-18}$p$_{5-10}$R4.1p$_{12-10}$R4.1p$_{12-4}^{2}$p$_{12-10}$R4.1p$_{12-10}$R4.1p$_{12-10}$c$_{4}^{7}$p$_{4-18}$p$_{1-3}$c$_{17}^{2}$p$_{17-18}$p$_{1-3}$c$_{17}^{4}$p$_{17-18}$p$_{1-3}$c$_{17}^{3}$p$_{17-18}$p$_{1-3}$c$_{17}$p$_{17-10}$R4.1p$_{12-10}$R4.1c$_{12}^{3}$p$_{12-18}$p$_{1-3}$c$_{17}$p$_{17-10}$R4.1p$_{12-10}$R4.1c$_{12}^{3}$p$_{12-18}$p$_{1-3}$Q4c$_{4}$p$_{4-10}$R4.1c$_{12}^{2}$p$_{12-18}$p$_{1-3}$c$_{17}$p$_{17-3}$c$_{17}^{2}$p$_{17-18}$p$_{1-3}$c$_{17}$p$_{17-3}$c$_{17}^{6}$p$_{17-18}$p$_{1-3}$c$_{17}$p$_{17-2}$p$_{14-3}$c$_{17}^{2}$p$_{17-18}$c$_{1}^{7}$p$_{1-18}$c$_{1}^{2}$p$_{1-18}$p$_{1-3}$c$_{17}^{2}$p$_{17-18}$p$_{1-3}$c$_{17}$p$_{17-3}$p$_{17-19}$c$_{5}$p$_{5-3}$p$_{17-10}$R4.1c$_{12}$p$_{12-10}$R4.1c$_{12}$p$_{12-10}$R4.1c$_{12}^{2}$p$_{12-6}$p$_{8-18}$p$_{1-3}$c$_{17}^{15}$p$_{17-18}$p$_{1-3}$c$_{17}$p$_{17-9}$p$_{3-4}$c$_{12}$p$_{12-19}$c$_{1}$p$_{1-18}$p$_{1-3}$c$_{17}$p$_{17-18}$p$_{1-3}$c$_{17}$p$_{17-3}$c$_{17}$p$_{17-18}$c$_{1}^{2}$p$_{1-18}$c$_{1}$p$_{1-2}$p$_{7-10}$p$_{18-1}$p$_{6-2}$c$_{7}^{5}$p$_{7-2}$c$_{7}^{2}$p$_{7-10}$p$_{4-5}$p$_{13-4}$p$_{12-2}$p$_{7-4}$p$_{12-2}$p$_{7-4}$p$_{12-2}$c$_{7}^{4}$p$_{7-1}$p$_{6-10}$p$_{18-1}$c$_{6}$p$_{6-2}$c$_{7}^{2}$p$_{7-14}$H4/9c$_{16}^{3}$p$_{16-1}$p$_{6-2}$c$_{7}^{4}$p$_{7-18}$c$_{5}^{8}$p$_{5-11}$c$_{5}^{3}$p$_{5-8}$p$_{2-18}$p$_{1-3}$c$_{17}$p$_{17-18}$p$_{1-3}$c$_{17}^{2}$p$_{17-4}$p$_{1-10}$R4.1c$_{12}^{2}$p$_{12-4}$p$_{1-10}$R4.1c$_{12}$p$_{12-19}$p$_{5-10}$R4.1c$_{12}$p$_{12-17}$p$_{19-4}$p$_{1-10}$R4.1p$_{12-4}$c$_{1}^{7}$p$_{1-4}$c$_{1}$p$_{1-4}$p$_{1-10}$R4.1p$_{12-3}$p$_{11-4}$p$_{1-10}$R4.1c$_{12}^{5}$p$_{12-4}$c$_{1}$p$_{1-4}$p$_{1-10}$R4.1c$_{12}$p$_{12-4}$c$_{12}$p$_{12-14}$H4/9p$_{16-17}$c$_{4}$R4.1c$_{12}$p$_{12-14}$H4/9p$_{16-17}$p$_{4-14}$H4/9p$_{16-17}$c$_{4}$p$_{4-10}$R4.1p$_{12-10}$R4.1c$_{12}$p$_{12-4}$c$_{12}$p$_{12-4}$c$_{12}$p$_{12-14}$H4/9p$_{16-17}$p$_{4-14}$H4/9p$_{16-17}$c$_{4}$p$_{4-10}$R4.1c$_{12}$p$_{12-4}$p$_{1-10}$R4.1c$_{12}$p$_{12-4}$c$_{1}$p$_{1-10}$R4.1c$_{12}^{2}$p$_{12-17}$c$_{4}^{2}$R4.1c$_{1}^{3}$p$_{1-4}$p$_{1-10}$R4.1p$_{12-4}$c$_{1}$p$_{1-4}$p$_{1-10}$R4.1c$_{12}$p$_{12-4}$p$_{1-10}$R4.1p$_{12-13}$p$_{19-10}$R4.1p$_{12-13}$S4p$_{12-10}$R4.1c$_{12}$p$_{12-13}$c$_{15}^{3}$p$_{15-10}$R4.1c$_{12}$p$_{12-10}$R4.1c$_{12}$p$_{12-10}$R4.1c$_{12}^{4}$p$_{12-4}$c$_{1}$p$_{1-4}$p$_{1-9}$p$_{3-4}$c$_{12}$p$_{12-4}$p$_{1-10}$R4.1c$_{12}$p$_{12-4}$p$_{1-4}$c$_{1}$p$_{1-4}$p$_{1-10}$R4.1p$_{12-4}$c$_{1}$p$_{1-4}$c$_{1}$p$_{1-4}$c$_{1}$p$_{1-4}$c$_{1}$p$_{1-15}$p$_{18-4}$p$_{13-5}^{2}$c$_{13}$p$_{13-8}$p$_{2-12}$p$_{2-8}$c$_{2}$p$_{2-18}$c$_{1}^{2}$p$_{1-18}^{4}$p$_{1-5}$c$_{13}^{3}$p$_{13-9}$c$_{5}^{4}$E4p$_{13-11}$E4c$_{13}^{2}$p$_{13-17}$p$_{4-6}$p$_{14-6}$c$_{14}^{7}$p$_{14-12}$c$_{6}^{6}$p$_{6-12}$p$_{6-15}$c$_{1}^{2}$p$_{1-12}$c$_{6}^{8}$p$_{6-7}$c$_{7}$G4c$_{7}$G4c$_{7}$G4c$_{7}$G4c$_{7}$p$_{7-14}$H4/9p$_{16-14}$H4/9p$_{16-14}$H4/9c$_{16}$p$_{16-14}$H4/9p$_{16-14}$H4/9p$_{16-14}$H4/9c$_{16}$p$_{16-14}$H4/9c$_{16}$p$_{16-14}$H4/9p$_{16-19}$p$_{5-11}$G4p$_{10-11}$p$_{13-14}$H4/9p$_{16-17}$R4.1p$_{1-2}$N4H4/9C4p$_{11-14}$H4/9p$_{16-2}$N4H4/9p$_{3-8}$p$_{16-2}$H4/9p$_{3-8}$p$_{16-2}$H4/9p$_{3-10}$p$_{18-2}$H4/9p$_{16-17}$p$_{5-14}$H4/9p$_{16-17}$p$_{4-11}$p$_{13-14}$H4/9p$_{16-17}$p$_{4-14}$H4/9p$_{16-17}$p$_{4-14}$H4/9p$_{16-17}$p$_{4-14}$H4/9p$_{16-17}$p$_{4-14}$H4/9p$_{16-17}$p$_{4-14}$H4/9p$_{16-17}$p$_{4-14}$H4/9p$_{16-17}$p$_{4-14}$H4/9p$_{16-2}$G4p$_{10-14}$H4/9p$_{16-1}$p$_{7-8}$p$_{10-14}$H4/9p$_{16-2}$p$_{7-8}$p$_{10-14}$H4/9p$_{16-1}$p$_{7-8}$p$_{10-14}$p$_{16-2}$p$_{7-8}$p$_{10-14}$p$_{16-2}$p$_{7-8}$p$_{10-14}$H4/9p$_{16-2}$p$_{7-8}$p$_{10-14}$H4/9p$_{16-2}$p$_{7-8}$p$_{10-13}$p$_{18-2}$p$_{7-8}$p$_{10-14}$H4/9p$_{16-2}$p$_{7-8}$p$_{10-13}$p$_{18-2}$p$_{7-8}$p$_{10-14}$H4/9p$_{16-2}$p$_{7-8}$p$_{10-13}$p$_{18-2}$p$_{7-8}$p$_{10-14}$H4/9p$_{16-2}$p$_{7-8}$p$_{10-13}$p$_{18-2}$p$_{7-8}$p$_{10-14}$H4/9p$_{16-2}$p$_{7-8}$p$_{10-13}$p$_{18-2}$p$_{7-8}$p$_{10-14}$H4/9p$_{16-2}$p$_{7-8}$p$_{10-14}$H4/9p$_{16-2}$p$_{7-8}$p$_{10-14}$H4/9p$_{16-2}$p$_{7-8}$p$_{10-14}$H4/9p$_{16-2}$p$_{7-8}$p$_{10-14}$H4/9p$_{16-2}$p$_{7-8}$p$_{10-14}$H4/9p$_{16-2}$p$_{7-8}$p$_{10-14}$H4/9p$_{16-2}$p$_{7-8}$p$_{10-14}$H4/9p$_{16-2}$p$_{7-8}$p$_{10-14}$H4/9p$_{16-2}$p$_{7-8}$p$_{10-14}$H4/9p$_{16-2}$p$_{7-8}$p$_{10-2}$p$_{7-8}$p$_{10-14}$H4/9p$_{16-2}$p$_{7-8}$p$_{10-14}$H4/9p$_{16-2}$p$_{7-8}$p$_{10-14}$H4/9p$_{16-2}$p$_{7-8}$p$_{10-14}$H4/9p$_{16-2}$p$_{7-8}$p$_{10-14}$H4/9p$_{16-2}$p$_{7-8}$p$_{10-15}$p$_{17-2}$p$_{7-8}$p$_{10-15}$p$_{17-2}$p$_{7-8}$p$_{10-14}$H4/9p$_{17-2}$p$_{7-8}$p$_{10-14}$H4/9p$_{16-2}$p$_{7-8}$p$_{10-14}$H4/9p$_{16-2}$p$_{7-8}$p$_{10-14}$H4/9p$_{16-2}$p$_{7-8}$p$_{10-14}$H4/9p$_{16-2}$p$_{7-8}$p$_{10-14}$H4/9p$_{16-2}$p$_{7-8}$p$_{10-14}$H4/9p$_{16-2}$p$_{7-8}$p$_{10-14}$H4/9p$_{16-2}$G4H4/9p$_{16-2}$p$_{7-8}$p$_{10-14}$H4/9p$_{16-2}$p$_{7-8}$p$_{10-14}$H4/9p$_{16-2}$p$_{7-8}$p$_{10-14}$H4/9p$_{16-2}$p$_{7-8}$p$_{10-14}$H4/9p$_{16-2}$p$_{7-8}$p$_{10-14}$H4/9p$_{16-2}$p$_{7-8}$p$_{10-14}$H4/9p$_{16-2}$p$_{7-8}$p$_{10-14}$H4/9p$_{16-2}$p$_{7-8}$p$_{10-14}$H4/9p$_{16-2}$p$_{7-8}$p$_{10-14}$H4/9p$_{16-2}$p$_{7-8}$p$_{10-14}$H4/9p$_{16-2}$p$_{7-8}$p$_{10-14}$H4/9p$_{16-17}$p$_{11-14}$H4/9p$_{16-2}$p$_{7-8}$p$_{10-14}$H4/9p$_{16-2}$p$_{7-8}$p$_{10-14}$H4/9p$_{16-2}$H4/9p$_{10-14}$H4/9p$_{16-2}$p$_{7-8}$p$_{10-14}$H4/9p$_{16-18}$p$_{1-2}$p$_{7-8}$p$_{17-18}$B4G4p$_{6-11}$S4p$_{12-14}$H4/9p$_{16-2}$H4/9p$_{16-19}$p$_{5-9}$p$_{10-13}$p$_{7-14}$H4/9p$_{16-18}$p$_{8-14}^{2}$p$_{5-7}$N4H4/9p$_{16-11}$M4p$_{1-11}$p$_{19-10}$p$_{19-13}$p$_{3-8}$p$_{13-1}$N4H4/9p$_{16-17}$R4.1B4N4H4/9p$_{3-5}$H4/9

### cen5

p$_{1-5}$P5E1/16/19/5P5p$_{5-1}$R1/19/5E1/16/19/5P5p$_{5-1}$p$_{3-5}$P5c$_{5}$E1/16/19/5P5p$_{5-1}$R1/19/5c$_{5}$E1/16/19/5P5p$_{5-1}$R1/19/5E1/16/19/5p$_{2-5}$P5p$_{5-1}$R1/19/5c$_{5}$p$_{1-5}$P5p$_{5-6}$E1/16/19/5P5E1/16/19/5P5p$_{5-1}$R1/19/5E1/16/19/5p$_{3-5}$P5c$_{5}$E1/16/19/5P5p$_{5-1}$R1/19/5E1/16/19/5p$_{2-5}$P5p$_{5-1}$R1/19/5p$_{5-1}$R1/19/5E1/16/19/5P5p$_{5-1}$R1/19/5c$_{5}$E1/16/19/5P5p$_{5-1}$p$_{3-}$

$_5$P5c$_5$E1/16/19/5P5p$_{5-1}$R1/19/5c$_5$E1/16/19/5P5p$_{5-1}$R1/19/5E1/16/19/5p$_{2-5}$P5p$_{5-}$
$_1$R1/19/5E1/16/19/5p$_{2-5}$P5p$_{5-1}$R1/19/5c$_5$E1/16/19/5P5p$_{5-1}$R1/19/5p$_{1-4}$A1/16/19/5R1/19/5p$_{5-3}$p$_{5-}$
$_1$R1/19/5E1/16/19/5p$_{2-5}$P5p$_{5-1}$R1/19/5p$_{5-1}$R1/19/5p$_{5-1}$R1/19/5c$_5$E1/16/19/5P5p$_{5-}$
$_1$R1/19/5c$_5$E1/16/19/5P5p$_{5-1}$R1/19/5p$_{5-1}$R1/19/5c$_5$E1/16/19/5P5p$_{5-1}$R1/19/5p$_{5-}$
$_1$R1/19/5E1/16/19/5p$_{3-5}$P5p$_{5-1}$R1/19/5c$_5$E1/16/19/5P5p$_{5-1}$R1/19/5p$_{5-1}$R1/19/5p$_{5-}$
$_1$R1/19/5c$_5$E1/16/19/5P5p$_{5-1}$R1/19/5p$_{5-1}$R1/19/5c$_5$E1/16/19/5P5p$_{5-1}$R1/19/5E1/16/19/5p$_{2-5}$P5p$_{5-}$
$_1$R1/19/5p$_{2-5}$P5p$_{5-1}$R1/19/5c$_5$E1/16/19/5P5p$_{5-1}$R1/19/5p$_{1-4}$A1/16/19/5R1/19/5p$_{5-3}$p$_{5-}$
$_1$R1/19/5c$_5$E1/16/19/5P5p$_{5-6}$p$_{3-5}$P5p$_{5-1}$R1/19/5p$_{5-1}$R1/19/5c$_5$p$_{5-1}$R1/19/5E1/16/19/5p$_{2-}$
$_5$P5c$_5$E1/16/19/5P5p$_{5-1}$R1/19/5c$_5$E1/16/19/5P5c$_5$E1/16/19/5P5p$_{5-1}$R1/19/5p$_{5-}$
$_1$R1/19/5c$_5$E1/16/19/5P5p$_{5-1}$R1/19/5c$_5$E1/16/19/5P5p$_{5-1}$R1/19/5E1/16/19/5P5p$_{5-}$
$_6$E1/16/19/5P5E1/16/19/5P5p$_{5-1}$R1/19/5E1/16/19/5p$_{3-5}$P5c$_5$E1/16/19/5P5p$_{5-}$
$_1$R1/19/5E1/16/19/5p$_{2-5}$P5p$_{5-1}$R1/19/5p$_{5-1}$R1/19/5E1/16/19/5P5p$_{5-1}$R1/19/5c$_5$E1/16/19/5P5p$_{5-1}$p$_{3-}$
$_5$P5c$_5$E1/16/19/5P5p$_{5-1}$R1/19/5c$_5$E1/16/19/5P5p$_{5-1}$R1/19/5E1/16/19/5p$_{2-5}$P5p$_{5-}$
$_1$R1/19/5E1/16/19/5p$_{2-5}$P5p$_{5-1}$R1/19/5c$_5$E1/16/19/5P5p$_{5-1}$R1/19/5p$_{1-4}$A1/16/19/5R1/19/5p$_{5-}$
$_1$R1/19/5p$_{1-4}$A1/16/19/5R1/19/5p$_{5-3}$p$_{5-1}$R1/19/5E1/16/19/5p$_{2-5}$P5p$_{5-1}$R1/19/5p$_{5-1}$R1/19/5p$_{5-}$
$_1$R1/19/5c$_5$E1/16/19/5P5p$_{5-1}$R1/19/5c$_5$E1/16/19/5P5p$_{5-1}$R1/19/5p$_{5-1}$R1/19/5c$_5$E1/16/19/5P5p$_{5-}$
$_1$R1/19/5p$_{5-1}$R1/19/5E1/16/19/5p$_{3-5}$P5p$_{5-1}$R1/19/5c$_5$E1/16/19/5P5p$_{5-1}$R1/19/5p$_{5-1}$R1/19/5p$_{5-}$
$_1$R1/19/5c$_5$E1/16/19/5P5p$_{5-1}$R1/19/5p$_{5-1}$R1/19/5c$_5$E1/16/19/5P5p$_{5-1}$R1/19/5E1/16/19/5p$_{2-5}$P5p$_{5-}$
$_1$R1/19/5E1/16/19/5p$_{2-5}$P5p$_{5-1}$R1/19/5c$_5$E1/16/19/5P5p$_{5-1}$R1/19/5p$_{1-5}$P5p$_{5-6}$p$_{3-5}$P5p$_{5-1}$R1/19/5p$_{5-}$
$_1$R1/19/5c$_5$p$_{5-1}$R1/19/5E1/16/19/5p$_{2-5}$P5c$_5$E1/16/19/5P5p$_{5-1}$R1/19/5p$_{3-5}$P5p$_{5-}$
$_1$R1/19/5c$_5$E1/16/19/5P5p$_{5-1}$R1/19/5p$_{5-1}$R1/19/5p$_{3-5}$P5c$_5$E1/16/19/5P5p$_{5-1}$R1/19/5p$_{5-1}$R1/19/5c$_5$p$_{5-}$
$_1$R1/19/5E1/16/19/5c$_2$p$_{2-5}$P5c$_5$E1/16/19/5P5p$_{5-1}$R1/19/5p$_{3-5}$P5p$_{5-1}$R1/19/5c$_5$E1/16/19/5P5p$_{5-}$
$_1$R1/19/5c$_5$p$_{5-1}$R1/19/5p$_{3-5}$P5p$_{5-6}$E1/16/19/5P5p$_{5-1}$R1/19/5p$_{5-1}$R1/19/5c$_5$E1/16/19/5P5p$_{5-}$
$_1$R1/19/5p$_{3-5}$P5p$_{5-1}$R1/19/5E1/16/19/5p$_{4-5}$P5c$_5$E1/16/19/5P5c$_5$E1/16/19/5P5c$_5$p$_{5-6}$p$_{5-1}$R1/19/5p$_{5-}$
$_1$R1/19/5p$_{5-1}$R1/19/5p$_{3-5}$P5c$_5$E1/16/19/5P5c$_5$E1/16/19/5P5c$_5$E1/16/19/5P5p$_{5-}$
$_1$R1/19/5c$_5$E1/16/19/5P5p$_{5-1}$R1/19/5c$_5$E1/16/19/5P5p$_{5-1}$R1/19/5c$_5$E1/16/19/5P5p$_{5-}$
$_1$R1/19/5c$_5$E1/16/19/5P5p$_{5-1}$R1/19/5c$_5$E1/16/19/5P5p$_{5-1}$R1/19/5p$_{3-5}$P5p$_{5-1}$R1/19/5p$_{4-5}$P5p$_{5-}$
$_6$R1/19/5p$_{3-5}$P5p$_{5-1}$R1/19/5c$_5$E1/16/19/5P5c$_5$p$_{3-5}$P5p$_{5-1}$R1/19/5c$_5$p$_{3-5}$P5p$_{5-6}$p$_{5-1}$R1/19/5p$_{5-}$
$_1$R1/19/5p$_{5-1}$R1/19/5c$_5$E1/16/19/5P5p$_{5-1}$R1/19/5c$_5$E1/16/19/5P5p$_{5-1}$R1/19/5c$_5$E1/16/19/5P5c$_5$p$_{5-}$
$_1$R1/19/5c$_5$p$_{5-1}$R1/19/5c$_5$E1/16/19/5P5c$_5$E1/16/19/5P5p$_{5-1}$R1/19/5c$_5$E1/16/19/5P5p$_{5-}$
$_1$R1/19/5c$_5$E1/16/19/5P5p$_{5-1}$R1/19/5c$_5$E1/16/19/5P5p$_{5-1}$R1/19/5p$_{5-1}$R1/19/5p$_{5-6}$p$_{3-5}$P5p$_{5-}$
$_1$R1/19/5p$_{5-1}$R1/19/5c$_5$E1/16/19/5P5p$_{5-1}$R1/19/5p$_{5-1}$R1/19/5p$_{5-1}$R1/19/5p$_{5-1}$R1/19/5p$_{5-}$
$_1$R1/19/5c$_5$E1/16/19/5P5p$_{5-1}$R1/19/5p$_{5-1}$R1/19/5p$_{5-1}$R1/19/5p$_{5-1}$R1/19/5p$_{5-}$
$_1$R1/19/5c$_5$E1/16/19/5P5p$_{5-1}$R1/19/5p$_{5-1}$R1/19/5p$_{5-6}$p$_{5-1}$R1/19/5c$_5$E1/16/19/5P5p$_{5-}$
$_1$R1/19/5c$_5$E1/16/19/5P5p$_{5-1}$R1/19/5p$_{5-3}$P5p$_{5-1}$R1/19/5p$_{5-1}$R1/19/5p$_{5-1}$R1/19/5p$_{5-1}$R1/19/5c$_5$p$_{5-}$
$_1$R1/19/5c$_5$E1/16/19/5P5p$_{5-1}$R1/19/5p$_{5-1}$R1/19/5c$_5$p$_{5-1}$R1/19/5c$_5$E1/16/19/5P5p$_{5-1}$R1/19/5p$_{5-}$
$_1$R1/19/5p$_{5-1}$R1/19/5p$_{5-1}$R1/19/5c$_5$E1/16/19/5P5p$_{5-1}$R1/19/5c$_5$E1/16/19/5P5p$_{5-}$
$_1$R1/19/5c$_5$E1/16/19/5P5p$_{5-1}$R1/19/5p$_{5-1}$R1/19/5p$_{5-1}$R1/19/5p$_{5-1}$R1/19/5c$_5$E1/16/19/5P5p$_{5-}$
$_1$R1/19/5c$_5$E1/16/19/5P5p$_{5-1}$R1/19/5c$_5$E1/16/19/5P5p$_{5-1}$R1/19/5p$_{5-1}$R1/19/5p$_{5-}$
$_1$R1/19/5c$_5$E1/16/19/5P5p$_{5-1}$R1/19/5c$_5$E1/16/19/5P5p$_{5-1}$R1/19/5c$_5$E1/16/19/5P5p$_{5-}$
$_1$R1/19/5c$_5$E1/16/19/5P5p$_{5-1}$R1/19/5c$_5$E1/16/19/5P5c$_5$E1/16/19/5P5p$_{5-}$
$_1$R1/19/5c$_5$E1/16/19/5P5p$_{5-1}$R1/19/5c$_5$E1/16/19/5P5p$_{5-1}$R1/19/5c$_5$E1/16/19/5P5p$_{5-1}$R1/19/5p$_{5-}$
$_1$R1/19/5p$_{5-1}$R1/19/5p$_{5-1}$R1/19/5c$_5$E1/16/19/5P5p$_{5-1}$R1/19/5p$_{5-1}$R1/19/5p$_{5-1}$R1/19/5p$_{5-1}$R1/19/5p$_{5-}$
$_1$R1/19/5c$_5$E1/16/19/5P5p$_{5-1}$R1/19/5p$_{5-1}$R1/19/5p$_{5-1}$R1/19/5p$_{5-1}$R1/19/5p$_{5-}$
$_1$R1/19/5c$_5$E1/16/19/5P5p$_{5-1}$R1/19/5c$_5$E1/16/19/5P5p$_{5-1}$R1/19/5c$_5$E1/16/19/5P5p$_{5-1}$R1/19/5p$_{5-}$
$_1$R1/19/5c$_5$E1/16/19/5P5p$_{5-1}$R1/19/5c$_5$E1/16/19/5P5p$_{5-1}$R1/19/5c$_5$E1/16/19/5P5p$_{5-1}$R1/19/5p$_{5-}$
$_1$R1/19/5p$_{5-1}$R1/19/5p$_{5-1}$R1/19/5c$_5$E1/16/19/5P5p$_{5-1}$R1/19/5p$_{5-1}$R1/19/5p$_{5-1}$R1/19/5p$_{5-1}$R1/19/5p$_{5-}$
$_1$R1/19/5c$_5$E1/16/19/5P5p$_{5-1}$R1/19/5p$_{5-1}$R1/19/5p$_{5-1}$R1/19/5p$_{5-1}$R1/19/5c$_5$E1/16/19/5P5p$_{5-}$
$_1$R1/19/5p$_{5-1}$R1/19/5p$_{5-1}$R1/19/5p$_{5-1}$R1/19/5c$_5$E1/16/19/5P5p$_{5-1}$R1/19/5p$_{5-}$
$_1$R1/19/5c$_5$E1/16/19/5P5p$_{5-1}$R1/19/5p$_{5-1}$R1/19/5c$_5$E1/16/19/5P5p$_{5-1}$R1/19/5c$_5$E1/16/19/5P5p$_{5-}$
$_1$R1/19/5p$_{5-1}$R1/19/5p$_{5-1}$R1/19/5p$_{5-1}$R1/19/5p$_{5-1}$R1/19/5c$_5$E1/16/19/5P5p$_{5-1}$R1/19/5p$_{5-1}$R1/19/5p$_{5-}$

$_{1}$R1/19/5p$_{5-1}$R1/19/5p$_{5-1}$R1/19/5c$_{5}$E1/16/19/5P5p$_{5-1}$R1/19/5p$_{5-1}$R1/19/5p$_{5-1}$R1/19/5p$_{5-}$
$_{1}$R1/19/5c$_{5}$E1/16/19/5P5p$_{5-1}$R1/19/5p$_{5-1}$R1/19/5p$_{5-1}$R1/19/5p$_{5-1}$R1/19/5c$_{5}$E1/16/19/5P5p$_{5-}$
$_{1}$R1/19/5p$_{5-1}$R1/19/5c$_{5}$E1/16/19/5P5p$_{5-1}$R1/19/5p$_{5-1}$R1/19/5c$_{5}$E1/16/19/5P5p$_{5-}$
$_{1}$R1/19/5c$_{5}$E1/16/19/5P5p$_{5-1}$R1/19/5p$_{5-1}$R1/19/5p$_{5-1}$R1/19/5c$_{5}$E1/16/19/5P5p$_{5-}$
$_{1}$R1/19/5c$_{5}$E1/16/19/5P5p$_{5-1}$R1/19/5p$_{5-1}$R1/19/5E1/16/19/5p$_{2-5}$P5p$_{5-1}$R1/19/5E1/16/19/5P5p$_{1-}$
$_{5}$P5p$_{5-1}$R1/19/5c$_{5}$E1/16/19/5P5p$_{5-1}$R1/19/5c$_{5}$E1/16/19/5P5p$_{5-1}$R1/19/5p$_{5-1}$R1/19/5p$_{5-1}$R1/19/5p$_{5-}$
$_{1}$R1/19/5c$_{5}$E1/16/19/5P5p$_{5-1}$R1/19/5p$_{5-1}$R1/19/5p$_{5-1}$R1/19/5p$_{5-1}$R1/19/5p$_{5-}$
$_{1}$R1/19/5c$_{5}$E1/16/19/5P5p$_{5-1}$R1/19/5p$_{5-1}$R1/19/5p$_{5-1}$R1/19/5p$_{5-1}$R1/19/5p$_{5-1}$R1/19/5p$_{5-1}$R1/19/5p$_{5-}$
$_{1}$R1/19/5p$_{5-1}$R1/19/5c$_{5}$E1/16/19/5P5p$_{5-1}$R1/19/5p$_{5-1}$R1/19/5p$_{5-1}$R1/19/5p$_{5-1}$R1/19/5p$_{5-}$
$_{1}$R1/19/5c$_{5}$E1/16/19/5P5p$_{5-1}$R1/19/5p$_{5-1}$R1/19/5p$_{5-1}$R1/19/5p$_{5-1}$R1/19/5p$_{5-}$
$_{1}$R1/19/5c$_{5}$E1/16/19/5P5p$_{5-1}$R1/19/5c$_{5}$E1/16/19/5P5p$_{5-1}$R1/19/5c$_{5}$E1/16/19/5P5p$_{5-1}$R1/19/5p$_{5-}$
$_{1}$R1/19/5c$_{5}$E1/16/19/5P5p$_{5-1}$R1/19/5c$_{5}$E1/16/19/5P5p$_{5-1}$R1/19/5c$_{5}$E1/16/19/5P5p$_{5-1}$R1/19/5p$_{5-}$
$_{1}$R1/19/5p$_{5-1}$R1/19/5p$_{5-1}$R1/19/5p$_{5-1}$R1/19/5c$_{5}$E1/16/19/5P5p$_{5-1}$R1/19/5c$_{5}$E1/16/19/5P5p$_{5-}$
$_{1}$R1/19/5c$_{5}$E1/16/19/5P5p$_{5-1}$R1/19/5p$_{5-1}$R1/19/5p$_{5-1}$R1/19/5p$_{5-1}$R1/19/5p$_{5-}$
$_{1}$R1/19/5c$_{5}$E1/16/19/5P5p$_{5-1}$R1/19/5p$_{5-1}$R1/19/5p$_{5-1}$R1/19/5p$_{5-1}$R1/19/5p$_{5-6}$p$_{5-1}$R1/19/5p$_{5-}$
$_{1}$R1/19/5p$_{5-1}$R1/19/5c$_{5}$E1/16/19/5P5p$_{5-1}$R1/19/5p$_{5-1}$R1/19/5p$_{5-1}$R1/19/5p$_{5-1}$R1/19/5p$_{5-1}$R1/19/5p$_{5-}$
$_{1}$R1/19/5p$_{5-1}$R1/19/5p$_{5-1}$R1/19/5c$_{5}$E1/16/19/5P5p$_{5-1}$R1/19/5p$_{5-1}$R1/19/5p$_{5-}$
$_{1}$R1/19/5c$_{5}$E1/16/19/5P5p$_{5-1}$R1/19/5c$_{5}$E1/16/19/5P5p$_{5-1}$R1/19/5c$_{5}$E1/16/19/5P5p$_{5-1}$R1/19/5p$_{5-}$
$_{1}$R1/19/5p$_{5-1}$R1/19/5p$_{5-1}$R1/19/5p$_{5-1}$R1/19/5c$_{5}$E1/16/19/5P5p$_{5-1}$R1/19/5c$_{5}$E1/16/19/5P5p$_{5-}$
$_{1}$R1/19/5c$_{5}$E1/16/19/5P5p$_{5-1}$R1/19/5p$_{5-1}$R1/19/5p$_{5-1}$R1/19/5p$_{5-1}$R1/19/5p$_{5-}$
$_{1}$R1/19/5c$_{5}$E1/16/19/5P5p$_{5-1}$R1/19/5p$_{5-1}$R1/19/5p$_{5-1}$R1/19/5p$_{5-1}$R1/19/5p$_{5-1}$R1/19/5p$_{5-1}$R1/19/5p$_{5-}$
$_{1}$R1/19/5p$_{5-1}$R1/19/5c$_{5}$E1/16/19/5P5p$_{5-1}$R1/19/5p$_{5-1}$R1/19/5p$_{5-1}$R1/19/5p$_{5-}$
$_{1}$R1/19/5c$_{5}$E1/16/19/5P5p$_{5-1}$R1/19/5p$_{5-1}$R1/19/5c$_{5}$E1/16/19/5P5p$_{5-1}$R1/19/5p$_{5-}$
$_{1}$R1/19/5c$_{5}$E1/16/19/5P5p$_{5-1}$R1/19/5c$_{5}$E1/16/19/5P5p$_{5-1}$R1/19/5p$_{5-1}$R1/19/5p$_{5-}$
$_{1}$R1/19/5c$_{5}$E1/16/19/5P5p$_{5-1}$R1/19/5c$_{5}$E1/16/19/5P5p$_{5-1}$R1/19/5p$_{5-1}$R1/19/5p$_{5-}$
$_{1}$R1/19/5c$_{5}$E1/16/19/5P5p$_{5-1}$R1/19/5c$_{5}$E1/16/19/5P5p$_{5-1}$R1/19/5p$_{5-1}$R1/19/5E1/16/19/5p$_{2-5}$P5p$_{5-}$
$_{1}$R1/19/5E1/16/19/5P5p$_{1-5}$P5p$_{5-1}$R1/19/5c$_{5}$E1/16/19/5P5p$_{5-1}$R1/19/5c$_{5}$E1/16/19/5P5p$_{5-1}$R1/19/5p$_{5-}$
$_{1}$R1/19/5p$_{5-1}$R1/19/5p$_{5-1}$R1/19/5c$_{5}$E1/16/19/5P5p$_{5-1}$R1/19/5p$_{5-1}$R1/19/5p$_{5-1}$R1/19/5p$_{5-}$
$_{1}$R1/19/5c$_{5}$p$_{5-1}$R1/19/5c$_{5}$A1/16/19/5R1/19/5c$_{5}$E1/16/19/5P5p$_{1-5}$P5p$_{5-1}$R1/19/5p$_{5-}$
$_{1}$R1/19/5c$_{5}$E1/16/19/5P5p$_{5-1}$R1/19/5c$_{5}$E1/16/19/5P5p$_{5-1}$R1/19/5E1/16/19/5p$_{2-}$
$_{5}$P5c$_{5}$E1/16/19/5P5p$_{5-1}$R1/19/5p$_{5-1}$R1/19/5p$_{5-1}$R1/19/5c$_{5}$E1/16/19/5P5p$_{5-}$
$_{1}$R1/19/5c$_{5}$E1/16/19/5P5p$_{5-1}$R1/19/5E1/16/19/5p$_{2-5}$P5c$_{5}$E1/16/19/5P5p$_{5-1}$R1/19/5p$_{5-1}$R1/19/5p$_{5-}$
$_{1}$R1/19/5c$_{5}$E1/16/19/5P5p$_{5-1}$R1/19/5c$_{5}$E1/16/19/5P5p$_{5-1}$R1/19/5p$_{5-1}$R1/19/5c$_{5}$E1/16/19/5P5p$_{5-}$
$_{1}$R1/19/5p$_{5-1}$R1/19/5E1/16/19/5p$_{3-5}$P5p$_{5-1}$R1/19/5c$_{5}$E1/16/19/5P5p$_{5-1}$R1/19/5p$_{5-1}$R1/19/5p$_{5-}$
$_{1}$R1/19/5c$_{5}$E1/16/19/5P5p$_{5-1}$R1/19/5p$_{5-1}$R1/19/5c$_{5}$E1/16/19/5P5p$_{5-1}$R1/19/5p$_{5-1}$R1/19/5p$_{5-}$
$_{1}$R1/19/5p$_{5-1}$R1/19/5p$_{5-1}$R1/19/5p$_{5-1}$R1/19/5p$_{5-1}$R1/19/5c$_{5}$E1/16/19/5P5p$_{5-1}$R1/19/5p$_{5-1}$R1/19/5p$_{5-}$
$_{1}$R1/19/5p$_{5-1}$R1/19/5p$_{5-1}$R1/19/5p$_{5-1}$R1/19/5p$_{5-1}$R1/19/5c$_{5}$E1/16/19/5P5p$_{5-1}$R1/19/5p$_{5-1}$R1/19/5p$_{5-}$
$_{1}$R1/19/5p$_{5-1}$R1/19/5p$_{5-1}$R1/19/5c$_{5}$E1/16/19/5P5p$_{5-1}$R1/19/5p$_{5-1}$R1/19/5p$_{5-}$
$_{1}$R1/19/5c$_{5}$E1/16/19/5P5p$_{5-1}$R1/19/5p$_{5-1}$R1/19/5p$_{5-1}$R1/19/5c$_{5}$E1/16/19/5P5c$_{5}$p$_{5-}$
$_{1}$R1/19/5c$_{5}$E1/16/19/5P5p$_{5-1}$R1/19/5c$_{5}$p$_{5-1}$R1/19/5c$_{5}$E1/16/19/5P5p$_{5-1}$R1/19/5c$_{5}$E1/16/19/5P5p$_{5-}$
$_{1}$R1/19/5p$_{5-1}$R1/19/5c$_{5}$p$_{5-1}$R1/19/5c$_{5}$E1/16/19/5P5p$_{5-1}$R1/19/5p$_{5-3}$p$_{5-1}$R1/19/5p$_{5-}$
$_{1}$R1/19/5c$_{5}$E1/16/19/5P5p$_{5-1}$R1/19/5c$_{5}$p$_{5-1}$R1/19/5c$_{5}$E1/16/19/5P5p$_{5-1}$R1/19/5p$_{5-}$
$_{1}$R1/19/5c$_{5}$E1/16/19/5P5p$_{5-1}$R1/19/5p$_{5-1}$R1/19/5c$_{5}$E1/16/19/5P5p$_{5-1}$R1/19/5c$_{5}$p$_{5-}$
$_{1}$R1/19/5c$_{5}$E1/16/19/5P5p$_{5-1}$R1/19/5p$_{5-1}$R1/19/5c$_{5}$E1/16/19/5P5p$_{5-1}$R1/19/5p$_{5-}$
$_{1}$R1/19/5c$_{5}$E1/16/19/5P5p$_{5-1}$R1/19/5p$_{5-1}$R1/19/5c$_{5}$E1/16/19/5P5p$_{5-1}$R1/19/5c$_{5}$E1/16/19/5P5p$_{5-}$
$_{1}$R1/19/5p$_{5-1}$R1/19/5c$_{5}$E1/16/19/5P5p$_{5-1}$R1/19/5p$_{5-1}$R1/19/5p$_{5-1}$R1/19/5p$_{5-}$
$_{1}$R1/19/5c$_{5}$E1/16/19/5P5p$_{5-1}$R1/19/5p$_{5-1}$R1/19/5c$_{5}$E1/16/19/5P5p$_{5-1}$R1/19/5c$_{5}$E1/16/19/5P5p$_{5-}$
$_{1}$R1/19/5p$_{5-1}$R1/19/5p$_{5-1}$R1/19/5c$_{5}$E1/16/19/5P5p$_{5-1}$R1/19/5p$_{5-1}$R1/19/5c$_{5}$E1/16/19/5P5p$_{5-}$
$_{1}$R1/19/5p$_{5-1}$R1/19/5c$_{5}$E1/16/19/5P5p$_{5-1}$R1/19/5p$_{5-}$
$_{1}$R1/19/5E1/16/19/5P5c$_{5}$E1/16/19/5P5E1/16/19/5P5c$_{5}$E1/16/19/5P5E1/16/19/5P5p$_{5-}$
$_{1}$R1/19/5E1/16/19/5P5c$_{5}$E1/16/19/5P5p$_{6-1}$R1/19/5c$_{5}$E1/16/19/5P5p$_{5-1}$R1/19/5p$_{5-1}$R1/19/5p$_{5-}$

$_1$R1/19/5$c_5$E1/16/19/5P5$p_{5-1}$R1/19/5$p_{5-1}$R1/19/5$p_{5-1}$R1/19/5$p_{5-1}$R1/19/5$c_5^2$E1/16/19/5P5$p_{5-}$
$_1$R1/19/5$p_{5-1}$R1/19/5$c_5$E1/16/19/5P5$p_{5-1}$R1/19/5$c_5$E1/16/19/5P5$p_{5-1}$R1/19/5$c_5$E1/16/19/5P5$p_{5-}$
$_1$R1/19/5$p_{5-1}$R1/19/5$p_{5-1}$R1/19/5$c_5$E1/16/19/5P5$p_{5-1}$R1/19/5$c_5$E1/16/19/5P5$p_{5-}$
$_1$R1/19/5$c_5$E1/16/19/5P5$p_{5-1}$R1/19/5$p_{5-1}$R1/19/5$p_{5-1}$R1/19/5$c_5$E1/16/19/5P5$p_{5-1}$R1/19/5$p_{5-}$
$_1$R1/19/5$p_{5-1}$R1/19/5$c_5$E1/16/19/5P5$c_5$E1/16/19/5P5$c_5$E1/16/19/5P5A1/16/19/5R1/19/5$p_{5-}$
$_1$R1/19/5$p_{5-6}$$c_5$E1/16/19/5P5$c_5$E1/16/19/5P5$p_{5-1}$R1/19/5$p_{5-1}$R1/19/5$p_{5-1}$R1/19/5$c_5$E1/16/19/5P5$p_{5-}$
$_1$R1/19/5$p_{5-1}$R1/19/5$p_{5-1}$R1/19/5$c_5$$p_{5-1}$R1/19/5$c_5$E1/16/19/5P5$p_{5-1}$R1/19/5$p_{5-1}$R1/19/5$p_{5-}$
$_1$R1/19/5$c_5$E1/16/19/5P5$p_{5-1}$R1/19/5$p_{5-1}$R1/19/5$p_{5-1}$R1/19/5$c_5$E1/16/19/5P5$p_{5-1}$R1/19/5$p_{5-}$
$_1$R1/19/5$c_5$E1/16/19/5P5$p_{5-1}$R1/19/5$p_{5-1}$R1/19/5$c_5$E1/16/19/5P5$p_{5-1}$R1/19/5$p_{5-}$
$_1$R1/19/5$c_5$E1/16/19/5P5$p_{5-1}$R1/19/5$p_{5-1}$R1/19/5$p_{5-1}$R1/19/5$c_5$E1/16/19/5P5$p_{5-1}$R1/19/5$p_{5-}$
$_1$R1/19/5$p_{5-1}$R1/19/5$c_5$E1/16/19/5P5$c_5$E1/16/19/5P5E1/16/19/5P5$p_{5-1}$R1/19/5$p_{5-1}$R1/19/5$p_{5-}$
$_1$R1/19/5$c_5$E1/16/19/5P5$c_5$E1/16/19/5P5$p_{5-1}$R1/19/5$p_{5-1}$R1/19/5$p_{5-1}$R1/19/5$c_5$E1/16/19/5P5$p_{5-}$
$_1$R1/19/5$p_{5-1}$R1/19/5$p_{5-1}$R1/19/5$c_5$E1/16/19/5P5$c_5$E1/16/19/5P5$c_5$E1/16/19/5P5$p_{5-}$
$_1$R1/19/5$c_5$E1/16/19/5P5$c_5$E1/16/19/5P5$c_5$E1/16/19/5P5$p_{5-1}$R1/19/5$p_{5-1}$R1/19/5$p_{5-}$
$_1$R1/19/5$c_5$E1/16/19/5P5$c_5$E1/16/19/5P5$p_{5-1}$R1/19/5$p_{5-1}$R1/19/5$p_{5-}$
$_1$R1/19/5$c_5$E1/16/19/5P5$c_5$E1/16/19/5P5$p_{5-1}$R1/19/5$p_{5-1}$R1/19/5$p_{5-1}$R1/19/5$c_5$E1/16/19/5P5$p_{5-}$
$_1$R1/19/5$p_{5-1}$R1/19/5$p_{5-1}$R1/19/5$c_5$E1/16/19/5P5$p_{5-1}$R1/19/5$p_{5-1}$R1/19/5$c_5$E1/16/19/5P5$p_{5-}$
$_1$R1/19/5$c_5$E1/16/19/5P5$c_5$E1/16/19/5P5$c_5$E1/16/19/5P5$p_{5-1}$R1/19/5$p_{5-1}$R1/19/5$p_{5-}$
$_1$R1/19/5$c_5$E1/16/19/5P5$p_{5-1}$R1/19/5$p_{5-1}$R1/19/5$c_5$E1/16/19/5P5$p_{5-1}$R1/19/5$p_{5-}$
$_1$R1/19/5$c_5$E1/16/19/5P5$p_{5-1}$R1/19/5$p_{5-1}$R1/19/5$c_5$E1/16/19/5P5$c_5$E1/16/19/5P5$p_{5-1}$R1/19/5$p_{5-}$
$_1$R1/19/5E1/16/19/5P5$p_{5-1}$R1/19/5$p_{5-1}$R1/19/5$p_{5-1}$R1/19/5$p_{5-1}$R1/19/5$p_{5-3}$P5$p_{5-1}$R1/19/5$p_{5-}$
$_1$R1/19/5$p_{5-1}$R1/19/5$p_{5-1}$R1/19/5$c_5$E1/16/19/5P5$c_5$E1/16/19/5P5$p_{5-1}$R1/19/5$p_{5-}$
$_1$R1/19/5E1/16/19/5P5$p_{5-1}$R1/19/5$p_{5-1}$R1/19/5$p_{5-1}$R1/19/5$p_{5-1}$R1/19/5$p_{5-3}$P5$p_{5-1}$R1/19/5$p_{5-}$
$_1$R1/19/5$p_{5-1}$R1/19/5$p_{5-1}$R1/19/5$c_5$E1/16/19/5P5$c_5$E1/16/19/5P5$p_{5-1}$R1/19/5$p_{5-1}$R1/19/5$p_{5-}$
$_1$R1/19/5$p_{5-1}$R1/19/5$p_{5-3}$P5$p_{5-1}$R1/19/5$p_{5-1}$R1/19/5$p_{5-1}$R1/19/5$p_{5-1}$R1/19/5$c_5$E1/16/19/5P5$p_{5-}$
$_1$R1/19/5$p_{5-1}$R1/19/5$p_{5-3}$P5$p_{5-1}$R1/19/5$p_{5-3}$P5$p_{5-1}$E1/16/19/5P5$p_{5-1}$R1/19/5$p_{5-}$
$_1$R1/19/5$c_5$E1/16/19/5P5$p_{5-1}$R1/19/5$p_{5-1}$R1/19/5$c_5$E1/16/19/5P5E1/16/19/5$p_{4-}$
$_5$P5$c_5$E1/16/19/5P5$p_{5-1}$R1/19/5$p_{5-1}$R1/19/5$p_{5-1}$R1/19/5$c_5$E1/16/19/5P5$p_{5-1}$R1/19/5$p_{5-1}$R1/19/5$p_{5-}$
$_1$R1/19/5$c_5$E1/16/19/5P5$p_{5-1}$R1/19/5$p_{5-1}$R1/19/5$p_{5-1}$R1/19/5$c_5$E1/16/19/5P5$p_{5-1}$R1/19/5$p_{5-}$
$_1$R1/19/5$p_{5-1}$R1/19/5$c_5$E1/16/19/5P5$p_{5-1}$R1/19/5$p_{5-1}$R1/19/5$p_{5-1}$R1/19/5$c_5$E1/16/19/5P5$p_{5-}$
$_1$R1/19/5$p_{5-1}$R1/19/5$p_{5-1}$R1/19/5$c_5$E1/16/19/5P5$c_5$$p_{5-1}$R1/19/5$c_5$E1/16/19/5P5$p_{5-1}$R1/19/5$p_{5-}$
$_1$R1/19/5$p_{5-1}$R1/19/5$c_5$E1/16/19/5P5$p_{5-1}$R1/19/5$p_{5-1}$R1/19/5$p_{5-1}$R1/19/5$c_5$E1/16/19/5P5$p_{5-}$
$_1$R1/19/5$p_{5-1}$R1/19/5$p_{5-1}$R1/19/5$c_5$E1/16/19/5P5$p_{5-1}$R1/19/5$p_{5-1}$R1/19/5$p_{5-}$
$_1$R1/19/5$c_5$E1/16/19/5P5$p_{5-1}$R1/19/5$p_{5-1}$R1/19/5$p_{5-1}$R1/19/5$c_5$E1/16/19/5P5$p_{5-1}$R1/19/5$p_{5-}$
$_1$R1/19/5$p_{5-1}$R1/19/5$c_5$E1/16/19/5P5$p_{5-1}$R1/19/5$p_{5-1}$R1/19/5$p_{5-1}$R1/19/5$c_5$E1/16/19/5P5$p_{5-}$
$_1$R1/19/5$p_{5-1}$R1/19/5$c_5$E1/16/19/5P5$c_5$E1/16/19/5P5$p_{5-1}$R1/19/5$p_{5-1}$R1/19/5$c_5$E1/16/19/5P5$p_{5-}$
$_1$R1/19/5$c_5$E1/16/19/5P5$p_{5-1}$R1/19/5$p_{5-1}$R1/19/5$c_5$E1/16/19/5P5$p_{5-1}$R1/19/5$p_{5-1}$R1/19/5$p_{5-}$
$_1$R1/19/5$c_5$E1/16/19/5P5$c_5$E1/16/19/5P5$p_{5-1}$R1/19/5$p_{5-1}$R1/19/5$p_{5-1}$R1/19/5$c_5$E1/16/19/5P5$p_{5-}$
$_1$R1/19/5$p_{5-1}$R1/19/5$c_5$E1/16/19/5P5$p_{5-1}$R1/19/5$p_{5-1}$R1/19/5$c_5$E1/16/19/5P5$p_{5-1}$R1/19/5$p_{5-}$
$_1$R1/19/5$c_5$E1/16/19/5P5$p_{5-1}$R1/19/5$p_{5-1}$R1/19/5$c_5$E1/16/19/5P5$p_{5-1}$R1/19/5$p_{5-}$
$_1$R1/19/5$c_5$E1/16/19/5P5$p_{5-1}$R1/19/5$p_{5-1}$R1/19/5$c_5$E1/16/19/5P5$c_5$E1/16/19/5P5$p_{5-1}$R1/19/5$p_{5-}$
$_1$R1/19/5$p_{5-1}$R1/19/5$c_5$E1/16/19/5P5$p_{5-1}$R1/19/5$p_{5-1}$R1/19/5$c_5$E1/16/19/5P5$p_{5-1}$R1/19/5$p_{5-}$
$_1$R1/19/5$p_{5-1}$R1/19/5$c_5$E1/16/19/5P5$p_{5-1}$R1/19/5$p_{5-1}$R1/19/5$c_5$E1/16/19/5P5$p_{5-1}$R1/19/5$p_{5-}$
$_1$R1/19/5$c_5$E1/16/19/5P5$p_{5-1}$R1/19/5$p_{5-1}$R1/19/5$c_5$E1/16/19/5P5$p_{5-1}$R1/19/5$c_5$E1/16/19/5P5$p_{5-}$
$_1$R1/19/5$p_{5-1}$R1/19/5$p_{5-1}$R1/19/5$p_{5-6}$$c_3$$p_{3-5}$P5$p_{5-1}$R1/19/5$p_{5-1}$R1/19/5$p_{5-1}$R1/19/5$c_5$E1/16/19/5P5$p_{5-}$
$_1$R1/19/5$p_{5-1}$R1/19/5$p_{5-1}$R1/19/5$c_5$E1/16/19/5P5$p_{5-1}$R1/19/5$p_{5-1}$R1/19/5$p_{5-}$
$_1$R1/19/5$c_5$E1/16/19/5P5$c_5$E1/16/19/5P5$p_{5-1}$R1/19/5$p_{5-1}$R1/19/5$p_{5-1}$R1/19/5$p_{5-}$
$_1$R1/19/5$c_5$E1/16/19/5P5$p_{5-1}$R1/19/5$p_{5-1}$R1/19/5$p_{5-1}$R1/19/5$p_{5-1}$R1/19/5$p_{5-1}$R1/19/5$p_{5-}$
$_1$R1/19/5$c_5$E1/16/19/5P5$p_{5-1}$R1/19/5$p_{5-1}$R1/19/5$p_{5-3}$P5$p_{5-1}$R1/19/5$p_{5-1}$R1/19/5$p_{5-}$
$_1$R1/19/5$c_5$E1/16/19/5P5$p_{5-1}$R1/19/5$p_{5-1}$R1/19/5$c_5$E1/16/19/5P5$p_{5-1}$R1/19/5$p_{5-1}$R1/19/5$p_{5-}$
$_1$R1/19/5$c_5$E1/16/19/5P5$p_{5-1}$R1/19/5$c_5$E1/16/19/5P5$p_{5-1}$R1/19/5$p_{5-1}$R1/19/5$c_5$E1/16/19/5P5$p_{5-}$

$_1$R1/19/5$p_{5-1}$R1/19/5$c_5$E1/16/19/5P5$p_{5-1}$R1/19/5$p_{5-1}$R1/19/5$p_{5-1}$R1/19/5$c_5$E1/16/19/5P5$p_{5-}$
$_1$R1/19/5$c_5$E1/16/19/5P5$p_{5-1}$R1/19/5$p_{5-1}$R1/19/5$p_{5-1}$R1/19/5$c_5$E1/16/19/5P5$p_{5-1}$R1/19/5$p_{5-}$
$_1$R1/19/5$c_5$E1/16/19/5P5$c_5$E1/16/19/5P5$p_{5-1}$R1/19/5$c_5$E1/16/19/5P5$p_{5-1}$R1/19/5$p_{5-}$
$_1$R1/19/5$c_5$E1/16/19/5P5$p_{5-1}$R1/19/5$p_{5-1}$R1/19/5$p_{5-1}$R1/19/5$c_5$E1/16/19/5P5$p_{5-1}$R1/19/5$p_{5-}$
$_1$R1/19/5$c_5$E1/16/19/5P5$p_{5-1}$$p_{6-1}$R1/19/5$p_{5-1}$R1/19/5$c_5$E1/16/19/5P5$p_{5-1}$R1/19/5$p_{5-1}$R1/19/5$p_{5-}$
$_1$R1/19/5$c_5$E1/16/19/5P5$p_{5-1}$R1/19/5$c_5$E1/16/19/5P5$p_{5-1}$R1/19/5$p_{5-1}$R1/19/5$c_5$E1/16/19/5P5$p_{5-}$
$_1$R1/19/5$p_{5-1}$R1/19/5$p_{5-1}$R1/19/5$c_5$E1/16/19/5P5$p_{5-1}$R1/19/5$c_5$E1/16/19/5P5$p_{5-1}$R1/19/5$p_{5-}$
$_1$R1/19/5$c_5$E1/16/19/5P5$p_{5-1}$R1/19/5$p_{5-1}$R1/19/5$p_{5-1}$R1/19/5$c_5$E1/16/19/5P5$p_{5-1}$R1/19/5$p_{5-}$
$_1$R1/19/5$c_5$E1/16/19/5P5$p_{5-1}$R1/19/5$p_{5-1}$R1/19/5$p_{5-1}$R1/19/5$c_5$E1/16/19/5P5$p_{5-1}$R1/19/5$p_{5-}$
$_1$R1/19/5$p_{5-1}$R1/19/5$c_5$E1/16/19/5P5$p_{5-1}$R1/19/5$c_5$E1/16/19/5P5$p_{5-1}$R1/19/5$c_5$E1/16/19/5P5$p_{5-}$
$_1$R1/19/5$p_{5-1}$R1/19/5$p_{5-1}$R1/19/5$c_5$E1/16/19/5P5$p_{5-1}$R1/19/5$p_{5-1}$R1/19/5E1/16/19/5P5$p_{5-}$
$_1$R1/19/5$p_{5-1}$R1/19/5$p_{5-1}$R1/19/5$c_5$E1/16/19/5P5$p_{5-1}$R1/19/5$p_{5-1}$R1/19/5$p_{5-}$
$_1$R1/19/5$c_5$E1/16/19/5P5$p_{5-1}$R1/19/5$p_{5-1}$R1/19/5$p_{5-1}$R1/19/5$c_5$E1/16/19/5P5$p_{5-1}$R1/19/5$p_{5-}$
$_1$R1/19/5E1/16/19/5P5$p_{5-1}$R1/19/5$p_{5-1}$R1/19/5$p_{5-1}$R1/19/5$c_5$E1/16/19/5P5$p_{5-1}$R1/19/5$p_{5-}$
$_1$R1/19/5$p_{5-1}$R1/19/5$c_5$E1/16/19/5P5$p_{5-1}$R1/19/5$p_{5-1}$R1/19/5$p_{5-1}$R1/19/5$p_{5-1}$R1/19/5$p_{5-}$
$_1$R1/19/5$c_5$E1/16/19/5P5$p_{5-1}$R1/19/5$p_{5-1}$R1/19/5$p_{5-1}$R1/19/5$c_5$E1/16/19/5P5$p_{5-1}$R1/19/5$p_{5-}$
$_1$R1/19/5$p_{5-1}$R1/19/5$c_5$E1/16/19/5P5$p_{5-1}$R1/19/5$p_{5-1}$R1/19/5$p_{5-1}$R1/19/5$c_5$E1/16/19/5P5$p_{5-}$
$_1$R1/19/5$p_{5-1}$R1/19/5E1/16/19/5P5$p_{5-1}$R1/19/5$p_{5-1}$R1/19/5$c_5$E1/16/19/5P5$p_{5-1}$R1/19/5$p_{5-}$
$_1$R1/19/5$p_{5-1}$R1/19/5$c_5$E1/16/19/5P5$p_{5-1}$R1/19/5$p_{5-1}$R1/19/5$p_{5-1}$R1/19/5$c_5$E1/16/19/5P5$p_{5-}$
$_1$R1/19/5$p_{5-1}$R1/19/5$p_{5-1}$R1/19/5$c_5$E1/16/19/5P5$p_{5-1}$R1/19/5$p_{5-1}$R1/19/5$p_{5-}$
$_1$R1/19/5$c_5$E1/16/19/5P5$p_{5-1}$R1/19/5$p_{5-1}$R1/19/5$p_{5-1}$R1/19/5$c_5$E1/16/19/5P5$p_{5-1}$R1/19/5$p_{5-}$
$_1$R1/19/5$p_{5-1}$R1/19/5$c_5$E1/16/19/5P5$p_{5-1}$R1/19/5$p_{5-1}$R1/19/5$c_5$$p_{5-1}$R1/19/5$p_{5-1}$R1/19/5$p_{5-}$
$_1$R1/19/5$c_5$$p_{5-1}$R1/19/5$p_{5-1}$R1/19/5$p_{5-1}$R1/19/5$c_5$$p_{5-1}$R1/19/5$c_5$E1/16/19/5P5$p_{5-1}$R1/19/5$c_5$$p_{5-}$
$_1$R1/19/5$p_{5-1}$R1/19/5E1/16/19/5$p_{2-5}$P5$c_5$E1/16/19/5P5$c_5$E1/16/19/5P5$c_5$E1/16/19/5P5$p_{5-}$
$_1$R1/19/5$p_{5-1}$R1/19/5E1/16/19/5P5$p_{5-1}$R1/19/5$c_5$E1/16/19/5P5$c_5$E1/16/19/5P5$c_5$E1/16/19/5P5$p_{5-}$
$_1$R1/19/5$p_{5-1}$R1/19/5$c_5$E1/16/19/5P5E1/16/19/5P5$p_{5-1}$R1/19/5$c_5$E1/16/19/5P5$p_{5-1}$R1/19/5$p_{5-}$
$_1$R1/19/5$p_{5-1}$R1/19/5$c_5$E1/16/19/5P5$p_{5-1}$R1/19/5$p_{5-1}$R1/19/5$c_5$E1/16/19/5P5$p_{5-1}$R1/19/5$p_{5-}$
$_1$R1/19/5$c_5$$p_{5-1}$R1/19/5$c_5$E1/16/19/5P5$p_{5-1}$R1/19/5$c_5$$p_{5-1}$R1/19/5$c_5$E1/16/19/5P5$p_{5-1}$R1/19/5$c_5$$p_{5-}$
$_1$R1/19/5$p_{5-1}$R1/19/5E1/16/19/5$p_{2-5}$P5$c_5$E1/16/19/5P5$c_5$E1/16/19/5P5$p_{5-1}$R1/19/5$p_{5-}$
$_1$R1/19/5$c_5$E1/16/19/5P5$p_{5-1}$R1/19/5$p_{5-1}$R1/19/5$c_5$$p_{5-1}$R1/19/5$p_{5-1}$R1/19/5$p_{5-}$
$_1$R1/19/5$c_5$E1/16/19/5P5$p_{5-1}$R1/19/5$p_{5-1}$R1/19/5$c_5$E1/16/19/5P5$p_{5-1}$R1/19/5$p_{5-1}$R1/19/5$p_{5-}$
$_1$R1/19/5$c_5$E1/16/19/5P5$c_5$E1/16/19/5P5$p_{5-1}$R1/19/5$p_{5-1}$R1/19/5$c_5$E1/16/19/5P5$p_{5-1}$R1/19/5$p_{5-}$
$_1$R1/19/5$p_{5-1}$R1/19/5$p_{5-1}$R1/19/5$p_{5-1}$R1/19/5$c_5$E1/16/19/5P5$p_{5-1}$R1/19/5$p_{5-1}$R1/19/5$p_{5-1}$R1/19/5$p_{5-}$
$_1$R1/19/5$c_5$E1/16/19/5P5$p_{5-1}$R1/19/5$c_5$E1/16/19/5P5$p_{5-1}$R1/19/5$p_{5-}$
$_6$E1/16/19/5P5$c_5$E1/16/19/5P5$p_{5-1}$R1/19/5$p_{5-1}$R1/19/5$c_5$E1/16/19/5P5$c_5$E1/16/19/5P5$p_{5-}$
$_1$R1/19/5E1/16/19/5P5$c_5$E1/16/19/5P5$p_{5-1}$R1/19/5$c_5$E1/16/19/5P5$p_{5-}$
$_1$R1/19/5E1/16/19/5P5$c_5$E1/16/19/5P5$p_{5-1}$R1/19/5$c_5$E1/16/19/5P5$p_{5-1}$R1/19/5$c_5$E1/16/19/5P5$p_{5-}$
$_1$R1/19/5$p_{5-1}$R1/19/5$c_5$E1/16/19/5P5$p_{5-1}$R1/19/5$p_{5-1}$R1/19/5$c_5$E1/16/19/5P5$p_{5-1}$R1/19/5$p_{5-}$
$_1$R1/19/5$p_{5-1}$R1/19/5$c_5$E1/16/19/5P5$p_{5-1}$R1/19/5$p_{5-1}$R1/19/5$c_5$E1/16/19/5P5$p_{5-1}$R1/19/5$p_{5-}$
$_1$R1/19/5$p_{5-1}$R1/19/5$c_5$E1/16/19/5P5$p_{5-1}$R1/19/5$p_{5-1}$R1/19/5$p_{5-1}$R1/19/5$c_5$E1/16/19/5P5$p_{5-}$
$_1$R1/19/5$p_{5-1}$R1/19/5$c_5$E1/16/19/5P5$p_{5-1}$R1/19/5$p_{5-1}$R1/19/5$p_{5-1}$R1/19/5$c_5$E1/16/19/5P5$p_{5-}$
$_1$R1/19/5$p_{5-1}$R1/19/5$p_{5-1}$R1/19/5$c_5$E1/16/19/5P5$p_{5-1}$R1/19/5$p_{5-1}$R1/19/5$c_5$E1/16/19/5P5$p_{5-}$
$_1$R1/19/5$p_{5-1}$R1/19/5$c_5$$p_{5-1}$R1/19/5$c_5$E1/16/19/5P5$p_{5-1}$R1/19/5$p_{5-1}$R1/19/5$p_{5-}$
$_1$R1/19/5$c_5$E1/16/19/5P5$p_{5-1}$R1/19/5$p_{5-1}$R1/19/5$p_{5-6}$E1/16/19/5P5$c_5$E1/16/19/5P5$p_{5-1}$R1/19/5$p_{5-}$
$_1$R1/19/5$c_5$E1/16/19/5P5$p_{5-1}$R1/19/5E1/16/19/5P5$c_5$E1/16/19/5P5$p_{5-1}$R1/19/5$p_{5-}$
$_1$R1/19/5$c_5$E1/16/19/5P5$p_{5-1}$R1/19/5$p_{5-1}$R1/19/5$p_{5-1}$R1/19/5$c_5$E1/16/19/5P5$p_{5-1}$R1/19/5$p_{5-}$
$_1$R1/19/5$c_5$E1/16/19/5P5$p_{5-1}$R1/19/5$p_{5-1}$R1/19/5$c_5$E1/16/19/5P5$p_{5-1}$R1/19/5$p_{5-}$
$_1$R1/19/5$c_5$E1/16/19/5P5$p_{5-1}$R1/19/5$p_{5-1}$R1/19/5E1/16/19/5P5$c_5$E1/16/19/5P5$p_{5-}$
$_1$R1/19/5E1/16/19/5P5$c_5$E1/16/19/5P5$p_{5-1}$R1/19/5$c_5$E1/16/19/5P5$p_{5-1}$R1/19/5$p_{5-1}$R1/19/5$p_{5-}$
$_6$E1/16/19/5P5$c_5$E1/16/19/5P5$p_{5-1}$R1/19/5$p_{5-1}$R1/19/5E1/16/19/5R1/19/5$c_5$E1/16/19/5P5$p_{5-}$
$_1$R1/19/5$p_{5-1}$R1/19/5E1/16/19/5P5$c_5$E1/16/19/5P5$p_{5-1}$R1/19/5$p_{5-1}$R1/19/5$c_5$E1/16/19/5P5$p_{5-}$

1R1/19/5p5-1R1/19/5p5-1R1/19/5c5E1/16/19/5P5p5-1R1/19/5c5E1/16/19/5P5p5-1R1/19/5p5-
1R1/19/5c5E1/16/19/5P5p5-1R1/19/5p5-1R1/19/5p5-1R1/19/5c5E1/16/19/5P5p5-1R1/19/5p5-
1R1/19/5p5-1R1/19/5E1/16/19/5P5p5-1R1/19/5c5E1/16/19/5P5p5-1R1/19/5p5-
1R1/19/5c5E1/16/19/5P5c5E1/16/19/5P5p5-1R1/19/5p5-1R1/19/5p5-1R1/19/5c5E1/16/19/5P5p5-
1R1/19/5p1-5P5p5-1R1/19/5E1/16/19/5P5c5E1/16/19/5P5p5-1R1/19/5c5E1/16/19/5P5p5-1R1/19/5p5-
1R1/19/5E1/16/19/5P5c5E1/16/19/5P5p5-1R1/19/5E1/16/19/5P5c5E1/16/19/5P5p5-
1R1/19/5c5E1/16/19/5P5c5E1/16/19/5P5p5-1R1/19/5p5-1R1/19/5p5-6c5E1/16/19/5P5p5-1R1/19/5p5-
1R1/19/5E1/16/19/5P5p5-1R1/19/5c5E1/16/19/5P5p5-1R1/19/5p5-1R1/19/5p5-
1R1/19/5c5E1/16/19/5P5p5-1R1/19/5c5E1/16/19/5P5p5-1R1/19/5p5-1R1/19/5p5-
1R1/19/5c5E1/16/19/5P5p5-1R1/19/5c5E1/16/19/5P5p5-1R1/19/5p5-1R1/19/5p5-
1R1/19/5c5E1/16/19/5P5p5-1R1/19/5p5-1R1/19/5c5E1/16/19/5P5p5-1R1/19/5p5-
1R1/19/5c5E1/16/19/5P5p5-1R1/19/5p5-1R1/19/5c5E1/16/19/5P5p5-1R1/19/5p5-
1R1/19/5c5E1/16/19/5P5p5-1R1/19/5p5-1R1/19/5p5-1R1/19/5c5E1/16/19/5P5p5-1R1/19/5p5-
1R1/19/5c5E1/16/19/5P5p5-1R1/19/5c5E1/16/19/5P5p5-1R1/19/5p5-1R1/19/5c5E1/16/19/5P5p5-
1R1/19/5c5E1/16/19/5P5p5-1R1/19/5p5-1R1/19/5c5E1/16/19/5P5p5-1R1/19/5p5-6c5E1/16/19/5P5p5-
1R1/19/5p5-1R1/19/5p5-1R1/19/5c5E1/16/19/5P5p5-1R1/19/5p5-1R1/19/5c5E1/16/19/5p4-5P5p5-
1R1/19/5p5-1R1/19/5p5-6c5E1/16/19/5P5p5-1R1/19/5p5-1R1/19/5p5-1R1/19/5c5E1/16/19/5P5p5-
1R1/19/5p5-1R1/19/5p5-1R1/19/5c5E1/16/19/5P5p5-1R1/19/5p5-1R1/19/5p5-1R1/19/5p5-
1R1/19/5E1/16/19/5P5p1-5P5p5-1R1/19/5p5-1R1/19/5p5-1R1/19/5c5E1/16/19/5P5p5-1R1/19/5p5-
1R1/19/5p5-1R1/19/5c5E1/16/19/5P5p5-1R1/19/5p5-1R1/19/5p5-
1R1/19/5E1/16/19/5P5c5E1/16/19/5P5p5-1R1/19/5p5-1R1/19/5E1/16/19/5P5c5E1/16/19/5P5p5-
1R1/19/5p5-1R1/19/5p5-6E1/16/19/5P5p5-1R1/19/5p5-1R1/19/5p5-1R1/19/5c5E1/16/19/5P5p5-
1R1/19/5p5-1R1/19/5p5-1R1/19/5p5-1R1/19/5c5E1/16/19/5P5E1/16/19/5R1/19/5p5-1R1/19/5p5-
1R1/19/5p5-1R1/19/5c5E1/16/19/5P5p5-1R1/19/5p5-1R1/19/5p5-1R1/19/5p5-
1R1/19/5c5E1/16/19/5P5p5-1R1/19/5p5-1R1/19/5p5-1R1/19/5c5E1/16/19/5P5p5-1R1/19/5p5-
1R1/19/5p5-1R1/19/5p5-1R1/19/5p5-1R1/19/5p5-1R1/19/5p5-1R1/19/5p5-1R1/19/5p5-1R1/19/5p5-
1R1/19/5c5E1/16/19/5P5p5-1R1/19/5p5-1R1/19/5p5-1R1/19/5c5E1/16/19/5P5p5-1R1/19/5p5-
1R1/19/5p5-1R1/19/5p5-1R1/19/5c5E1/16/19/5P5p5-1R1/19/5c5E1/16/19/5P5p5-
1R1/19/5c5E1/16/19/5P5p5-1R1/19/5p5-1R1/19/5p5-1R1/19/5c5E1/16/19/5P5p5-1R1/19/5p5-
1R1/19/5p5-1R1/19/5c5E1/16/19/5P5p5-1R1/19/5c5E1/16/19/5P5p5-1R1/19/5p5-
1R1/19/5c5E1/16/19/5P5p5-1R1/19/5p5-1R1/19/5c5E1/16/19/5P5p5-1R1/19/5p5-1R1/19/5p5-
1R1/19/5c5E1/16/19/5P5p5-1R1/19/5p5-6E1/16/19/5P5p5-1R1/19/5p5-1R1/19/5p5-
1R1/19/5c5E1/16/19/5P5p5-1R1/19/5p5-1R1/19/5p5-1R1/19/5c5E1/16/19/5P5p5-1R1/19/5p5-
1R1/19/5p5-1R1/19/5c5E1/16/19/5P5p5-1R1/19/5p5-1R1/19/5p5-1R1/19/5c5E1/16/19/5P5p5-
1R1/19/5p5-1R1/19/5c5E1/16/19/5P5p5-1R1/19/5c5E1/16/19/5P5p5-1R1/19/5p5-1R1/19/5p5-
1R1/19/5c5E1/16/19/5P5p5-1R1/19/5p5-1R1/19/5p5-1R1/19/5c5E1/16/19/5P5p5-1R1/19/5p5-
1R1/19/5c5E1/16/19/5P5p5-1R1/19/5p5-1R1/19/5c5E1/16/19/5P5p5-1R1/19/5c5E1/16/19/5P5p5-
1R1/19/5p5-1R1/19/5c5E1/16/19/5P5c5p5-1R1/19/5p5-1R1/19/5p5-1R1/19/5p5-1R1/19/5p5-
1R1/19/5p5-1R1/19/5p5-1R1/19/5c5E1/16/19/5P5p5-1R1/19/5p5-1R1/19/5p5-
1R1/19/5c5E1/16/19/5P5p5-1R1/19/5p5-1R1/19/5p5-1R1/19/5p5-1R1/19/5c5E1/16/19/5P5p5-
1R1/19/5c5E1/16/19/5P5p5-1R1/19/5p5-1R1/19/5c5E1/16/19/5P5p5-1R1/19/5p5-1R1/19/5p5-
1R1/19/5c5E1/16/19/5P5p5-1R1/19/5p5-1R1/19/5p5-1R1/19/5c5E1/16/19/5P5p5-
1R1/19/5c5E1/16/19/5P5p5-1R1/19/5p5-1R1/19/5c5E1/16/19/5P5p5-1R1/19/5p5-
1R1/19/5c5E1/16/19/5P5p5-1R1/19/5p5-1R1/19/5p5-1R1/19/5c5E1/16/19/5P5p5-1R1/19/5p5-
6E1/16/19/5P5p5-1R1/19/5p5-1R1/19/5p5-1R1/19/5c5E1/16/19/5P5p5-1R1/19/5p5-1R1/19/5p5-
1R1/19/5c5E1/16/19/5P5p5-1R1/19/5p5-1R1/19/5p5-1R1/19/5c5E1/16/19/5P5p5-1R1/19/5p5-
1R1/19/5p5-1R1/19/5c5E1/16/19/5P5p5-1R1/19/5p5-1R1/19/5c5E1/16/19/5P5p5-
1R1/19/5c5E1/16/19/5P5p5-1R1/19/5p5-1R1/19/5p5-1R1/19/5c5E1/16/19/5P5p5-1R1/19/5p5-
1R1/19/5p5-1R1/19/5c5E1/16/19/5P5p5-1R1/19/5p5-1R1/19/5c5E1/16/19/5P5p5-1R1/19/5p5-
1R1/19/5c5E1/16/19/5P5p5-1R1/19/5c5E1/16/19/5P5p5-1R1/19/5p5-
1R1/19/5c5E1/16/19/5P5c5E1/16/19/5P5p5-1R1/19/5p5-1R1/19/5p5-1R1/19/5c5E1/16/19/5P5p5-

$_1$R1/19/5$p_{5-1}$R1/19/5$c_5$E1/16/19/5P5$p_{5-1}$R1/19/5$p_{5-1}$R1/19/5$p_{5-}$
$_1$R1/19/5$c_5$E1/16/19/5P5$c_5$E1/16/19/5P5$p_{5-1}$R1/19/5$p_{5-1}$R1/19/5$p_{5-1}$R1/19/5E1/16/19/5P5$p_{5-}$
$_1$R1/19/5$c_5$E1/16/19/5P5$p_{5-1}$R1/19/5$p_{5-1}$R1/19/5$p_{5-1}$R1/19/5$c_5$E1/16/19/5P5$c_5$E1/16/19/5P5$p_{5-}$
$_1$R1/19/5$p_{5-1}$R1/19/5$p_{5-1}$R1/19/5$c_5$E1/16/19/5P5$p_{5-1}$R1/19/5$c_5$E1/16/19/5P5$p_{5-}$
$_1$R1/19/5$c_5$E1/16/19/5P5$c_5$E1/16/19/5P5$p_{5-1}$R1/19/5$p_{5-1}$R1/19/5$p_{5-1}$R1/19/5$c_5$E1/16/19/5P5$p_{5-}$
$_1$R1/19/5$p_{5-1}$R1/19/5$p_{5-3}$R1/19/5$p_{5-1}$R1/19/5$c_5$E1/16/19/5P5$c_5$E1/16/19/5P5$p_{5-1}$R1/19/5$p_{5-}$
$_1$R1/19/5$c_5$E1/16/19/5P5$p_{5-1}$R1/19/5$p_{5-1}$R1/19/5$c_5$E1/16/19/5P5$p_{5-1}$R1/19/5$p_{5-1}$R1/19/5$p_{5-}$
$_1$R1/19/5$c_5$E1/16/19/5P5E1/16/19/5R1/19/5$p_{5-1}$R1/19/5$p_{5-1}$R1/19/5$c_5$E1/16/19/5P5$p_{5-1}$R1/19/5$p_{5-}$
$_1$R1/19/5$p_{5-1}$R1/19/5$p_{5-1}$R1/19/5E1/16/19/5P5$c_5$E1/16/19/5P5$p_{5-1}$R1/19/5$p_{5-1}$R1/19/5$p_{5-}$
$_1$R1/19/5$p_{5-1}$R1/19/5$c_5$E1/16/19/5P5$p_{5-1}$R1/19/5$p_{5-1}$R1/19/5$p_{5-}$
$_1$R1/19/5E1/16/19/5P5$c_5^2$E1/16/19/5P5$p_{5-1}$R1/19/5$c_5$E1/16/19/5P5A1/16/19/5R1/19/5$p_{5-}$
$_1$R1/19/5$p_{5-6}p_{3-5}$P5$c_5$E1/16/19/5P5A1/16/19/5R1/19/5$p_{5-1}$R1/19/5$p_{5-}$
$_6$E1/16/19/5P5$c_5$E1/16/19/5P5$p_{5-1}$R1/19/5$p_{5-1}$R1/19/5E1/16/19/5P5$c_5$E1/16/19/5P5$p_{5-1}$R1/19/5$p_{5-}$
$_1$R1/19/5$c_5$E1/16/19/5P5$p_{5-1}$R1/19/5$p_{5-1}$R1/19/5$p_{5-1}$R1/19/5$c_5$E1/16/19/5P5$p_{5-}$
$_1$R1/19/5$c_5$E1/16/19/5P5$p_{5-1}$R1/19/5$p_{5-1}$R1/19/5$p_{5-1}$R1/19/5$c_5$E1/16/19/5P5$p_{5-}$
$_1$R1/19/5$c_5$E1/16/19/5P5$p_{5-1}$R1/19/5$c_5$E1/16/19/5P5$p_{5-1}$R1/19/5$p_{5-1}$R1/19/5$c_5$E1/16/19/5P5$p_{5-}$
$_1$R1/19/5$p_{5-1}$R1/19/5$c_5$E1/16/19/5P5$c_5$E1/16/19/5P5$p_{5-1}$R1/19/5$p_{5-1}$R1/19/5$p_{5-}$
$_1$R1/19/5$c_5$E1/16/19/5P5$p_{5-1}$R1/19/5$c_5$E1/16/19/5P5$p_{5-1}$R1/19/5$p_{5-}$
$_1$R1/19/5$c_5$E1/16/19/5P5$c_5$E1/16/19/5P5$p_{5-1}$R1/19/5$p_{5-1}$R1/19/5$p_{5-1}$R1/19/5$c_5$E1/16/19/5P5$p_{5-}$
$_1$R1/19/5$c_5$E1/16/19/5P5$c_5$E1/16/19/5P5$c_5$E1/16/19/5P5$p_{5-1}$R1/19/5$p_{5-1}$R1/19/5$p_{5-}$
$_1$R1/19/5$c_5$E1/16/19/5P5$c_5$E1/16/19/5P5$p_{5-1}$R1/19/5$p_{5-1}$R1/19/5$p_{5-1}$R1/19/5$c_5$E1/16/19/5P5$p_{5-}$
$_1$R1/19/5$p_{5-1}$R1/19/5$c_5$E1/16/19/5P5$p_{5-1}$R1/19/5$p_{5-1}$R1/19/5$c_5$E1/16/19/5P5$p_{5-1}$R1/19/5$p_{5-}$
$_1$R1/19/5$c_5$E1/16/19/5P5$p_{5-1}$R1/19/5$p_{5-1}$R1/19/5$p_{5-1}$R1/19/5$c_5$E1/16/19/5P5E1/16/19/5R1/19/5$p_{5-}$
$_1$R1/19/5$p_{5-1}$R1/19/5$c_5$E1/16/19/5P5$p_{5-1}$R1/19/5$p_{5-1}$R1/19/5$p_{5-1}$R1/19/5$p_{5-}$
$_1$R1/19/5E1/16/19/5P5$c_5$E1/16/19/5P5$p_{5-1}$R1/19/5$p_{5-1}$R1/19/5$p_{5-1}$R1/19/5$p_{5-1}$R1/19/5$p_{5-}$
$_1$R1/19/5E1/16/19/5P5$c_5$E1/16/19/5P5$p_{5-1}$R1/19/5$p_{5-1}$R1/19/5$p_{5-1}$R1/19/5$p_{5-}$
$_1$R1/19/5$c_5$E1/16/19/5P5$p_{5-1}$R1/19/5$p_{5-1}$R1/19/5$p_{5-1}$R1/19/5E1/16/19/5P5$c_5^2$E1/16/19/5P5$p_{5-}$
$_1$R1/19/5$c_5$E1/16/19/5P5A1/16/19/5R1/19/5$p_{5-1}$R1/19/5$p_{5-6}p_{3-}$
$_5$P5$c_5$E1/16/19/5P5A1/16/19/5R1/19/5$p_{5-1}$R1/19/5$p_{5-6}$E1/16/19/5P5$c_5$E1/16/19/5P5$p_{5-1}$R1/19/5$p_{5-}$
$_1$R1/19/5E1/16/19/5P5$c_5$E1/16/19/5P5$p_{5-1}$R1/19/5$p_{5-1}$R1/19/5$c_5$E1/16/19/5P5$p_{5-1}$R1/19/5$p_{5-}$
$_1$R1/19/5$p_{5-1}$R1/19/5$c_5$E1/16/19/5P5$p_{5-1}$R1/19/5$c_5$E1/16/19/5P5$p_{5-1}$R1/19/5$c_5$E1/16/19/5P5$p_{5-}$
$_1$R1/19/5$p_{5-1}$R1/19/5$c_5$E1/16/19/5P5$p_{5-1}$R1/19/5$c_5$E1/16/19/5P5$p_{5-1}$R1/19/5$c_5$E1/16/19/5P5$p_{5-}$
$_1$R1/19/5$p_{5-1}$R1/19/5$c_5$E1/16/19/5P5$p_{5-1}$R1/19/5$c_5$E1/16/19/5P5$p_{5-1}$R1/19/5$p_{5-}$
$_1$R1/19/5$c_5$E1/16/19/5P5$p_{5-1}$R1/19/5$c_5$E1/16/19/5P5$p_{5-1}$R1/19/5$p_{5-1}$R1/19/5$c_5$E1/16/19/5P5$p_{5-}$
$_1$R1/19/5$p_{5-1}$R1/19/5E1/16/19/5P5$p_{5-1}$R1/19/5$p_{5-1}$R1/19/5$p_{5-1}$R1/19/5$c_5p_{5-1}$R1/19/5$p_{5-}$
$_1$R1/19/5$c_5^2$E1/16/19/5P5$p_{5-1}$R1/19/5$p_{5-1}$R1/19/5E1/16/19/5P5$p_{5-1}$R1/19/5$p_{5-1}$R1/19/5$p_{5-}$
$_1$R1/19/5$c_5$E1/16/19/5P5$p_{5-1}$R1/19/5$c_5$E1/16/19/5P5$p_{5-1}$R1/19/5$p_{5-1}$R1/19/5$c_5$E1/16/19/5P5$p_{5-}$
$_1$R1/19/5$c_5$E1/16/19/5P5$p_{5-1}$R1/19/5$p_{5-1}$R1/19/5$p_{5-1}$R1/19/5$c_5$E1/16/19/5P5$p_{5-1}$R1/19/5$p_{5-}$
$_1$R1/19/5$c_5$E1/16/19/5P5$p_{5-1}$R1/19/5$p_{5-1}$R1/19/5$p_{5-6}p_{5-1}$R1/19/5$p_{5-1}$R1/19/5$p_{5-}$
$_1$R1/19/5E1/16/19/5P5$c_5$E1/16/19/5P5$p_{5-1}$R1/19/5$p_{5-1}$R1/19/5$p_{5-6}$E1/16/19/5P5$p_{5-1}$R1/19/5$p_{5-}$
$_1$R1/19/5E1/16/19/5P5$c_5$E1/16/19/5P5$p_{5-1}$R1/19/5E1/16/19/5P5$c_5$E1/16/19/5P5$p_{5-1}$R1/19/5$p_{5-}$
$_1$R1/19/5$c_5$E1/16/19/5P5$p_{5-1}$R1/19/5$p_{5-1}$R1/19/5$c_5$E1/16/19/5P5$p_{5-1}$R1/19/5$p_{5-}$
$_1$R1/19/5$c_5$E1/16/19/5P5$p_{5-1}$R1/19/5$p_{5-1}$R1/19/5$p_{5-1}$R1/19/5$c_5$E1/16/19/5P5$p_{5-1}$R1/19/5$p_{5-}$
$_1$R1/19/5$p_{5-1}$R1/19/5$c_5$E1/16/19/5P5$p_{5-1}$R1/19/5$p_{5-1}$R1/19/5$c_5$E1/16/19/5P5$p_{5-1}$R1/19/5$p_{5-}$
$_1$R1/19/5$p_{5-1}$R1/19/5$c_5$E1/16/19/5P5$p_{5-1}$R1/19/5$p_{5-1}$R1/19/5$c_5$E1/16/19/5P5$p_{5-1}$R1/19/5$p_{5-}$
$_1$R1/19/5$p_{5-1}$R1/19/5$c_5$E1/16/19/5P5$p_{5-1}$R1/19/5$p_{5-1}$R1/19/5$p_{5-1}$R1/19/5$c_5$E1/16/19/5P5$p_{5-}$
$_1$R1/19/5$p_{5-1}$R1/19/5$c_5p_{5-1}$R1/19/5$c_5$E1/16/19/5P5$p_{5-1}$R1/19/5$p_{5-1}$R1/19/5$c_5$E1/16/19/5P5$p_{5-}$
$_1$R1/19/5$c_5$E1/16/19/5P5$p_{5-1}$R1/19/5$p_{5-1}$R1/19/5$p_{5-1}$R1/19/5$c_5$E1/16/19/5P5$p_{5-1}$R1/19/5$p_{6-}$
$_1$R1/19/5$c_5$E1/16/19/5P5$p_{5-1}$R1/19/5$p_{5-1}$R1/19/5$c_5$E1/16/19/5P5$p_{5-1}$R1/19/5$p_{5-}$
$_1$R1/19/5$c_5$E1/16/19/5P5$p_{5-1}$R1/19/5$p_{5-1}$R1/19/5$c_5$E1/16/19/5P5$p_{5-1}$R1/19/5$p_{5-1}$R1/19/5$p_{5-}$
$_1$R1/19/5$c_5$E1/16/19/5P5$p_{5-1}$R1/19/5$p_{5-1}$R1/19/5$c_5$E1/16/19/5P5$p_{5-1}$R1/19/5$p_{5-1}$R1/19/5$p_{5-}$

${}_{1}R1/19/5E1/16/19/5P5p_{5-1}R1/19/5c_5E1/16/19/5P5p_{5-1}R1/19/5p_{5-1}R1/19/5c_5E1/16/19/5P5p_{5-}$
${}_{1}R1/19/5p_{5-1}R1/19/5p_{5-1}R1/19/5E1/16/19/5P5p_{5-1}R1/19/5c_5E1/16/19/5P5p_{5-1}R1/19/5p_{5-}$
${}_{1}R1/19/5p_{5-1}R1/19/5p_{5-1}R1/19/5E1/16/19/5P5p_{5-1}R1/19/5c_5E1/16/19/5P5p_{5-1}R1/19/5p_{5-}$
${}_{1}R1/19/5c_5E1/16/19/5P5p_{5-1}R1/19/5p_{5-1}R1/19/5p_{5-1}R1/19/5E1/16/19/5P5c_5E1/16/19/5P5p_{5-}$
${}_{1}R1/19/5c_5E1/16/19/5P5p_{5-1}R1/19/5p_{5-1}R1/19/5c_5E1/16/19/5P5p_{5-1}R1/19/5p_{5-}$
${}_{1}R1/19/5c_5E1/16/19/5P5p_{5-1}R1/19/5c_5E1/16/19/5P5p_{5-1}R1/19/5p_{5-1}R1/19/5p_{5-}$
${}_{1}R1/19/5c_5E1/16/19/5P5c_5E1/16/19/5P5p_{5-1}R1/19/5p_{5-1}R1/19/5p_{5-1}R1/19/5c_5E1/16/19/5P5p_{5-}$
${}_{1}R1/19/5p_{5-1}R1/19/5c_5E1/16/19/5P5p_{5-1}R1/19/5p_{5-1}R1/19/5p_{5-1}R1/19/5c_5E1/16/19/5P5p_{5-}$
${}_{1}R1/19/5c_5E1/16/19/5P5p_{5-1}R1/19/5p_{5-1}R1/19/5p_{5-1}R1/19/5c_5E1/16/19/5P5p_{5-}$
${}_{1}R1/19/5c_5E1/16/19/5P5p_{5-1}R1/19/5c_5p_{5-1}R1/19/5p_{5-1}R1/19/5p_{5-1}R1/19/5c_5E1/16/19/5P5p_{5-}$
${}_{1}R1/19/5p_{5-1}R1/19/5p_{5-6}E1/16/19/5P5c_5E1/16/19/5P5p_{5-1}R1/19/5p_{5-1}R1/19/5p_{5-}$
${}_{1}R1/19/5c_5E1/16/19/5P5p_{5-1}R1/19/5p_{5-1}R1/19/5c_5E1/16/19/5P5p_{5-1}R1/19/5p_{5-1}R1/19/5p_{5-}$
${}_{1}R1/19/5c_5E1/16/19/5P5p_{5-1}R1/19/5p_{5-1}R1/19/5c_5E1/16/19/5P5p_{5-1}R1/19/5p_{5-1}R1/19/5p_{5-}$
${}_{1}R1/19/5p_{5-1}R1/19/5E1/16/19/5P5c_5A1/16/19/5R1/19/5p_{5-1}R1/19/5c_5p_{5-1}R1/19/5p_{5-1}R1/19/5p_{5-}$
${}_{1}R1/19/5c_5E1/16/19/5P5p_{5-1}R1/19/5p_{5-1}R1/19/5p_{5-6}E1/16/19/5P5c_5E1/16/19/5P5p_{5-1}R1/19/5p_{5-}$
${}_{1}R1/19/5p_{5-1}R1/19/5c_5E1/16/19/5P5p_{5-1}R1/19/5p_{5-1}R1/19/5p_{5-1}R1/19/5c_5p_{5-1}R1/19/5p_{5-}$
${}_{1}R1/19/5c_5p_{5-1}R1/19/5p_{5-1}R1/19/5c_5p_{5-1}R1/19/5c_5p_{5-1}R1/19/5p_{5-1}R1/19/5c_5p_{5-1}R1/19/5p_{5-}$
${}_{1}R1/19/5c_5p_{5-1}R1/19/5p_{5-1}R1/19/5c_5p_{5-1}R1/19/5p_{5-1}R1/19/5c_5p_{5-1}R1/19/5p_{5-1}R1/19/5c_5p_{5-}$
${}_{1}R1/19/5p_{5-1}R1/19/5c_5p_{5-1}R1/19/5p_{5-1}R1/19/5c_5p_{5-1}R1/19/5p_{5-1}R1/19/5c_5p_{5-1}R1/19/5p_{5-}$
${}_{1}R1/19/5c_5E1/16/19/5p_{5-1}R1/19/5p_{5-1}R1/19/5p_{5-1}R1/19/5p_{5-1}R1/19/5p_{5-}$
${}_{1}R1/19/5E1/16/19/5P5c_5A1/16/19/5R1/19/5p_{5-1}R1/19/5c_5E1/16/19/5P5p_{5-1}R1/19/5p_{5-}$
${}_{1}R1/19/5c_5E1/16/19/5P5p_{5-1}R1/19/5p_{5-1}R1/19/5E1/16/19/5p_{2-5}P5c_5E1/16/19/5P5p_{5-1}R1/19/5p_{5-}$
${}_{1}R1/19/5c_5E1/16/19/5P5p_{5-1}R1/19/5p_{5-1}R1/19/5c_5E1/16/19/5P5p_{5-1}R1/19/5p_{5-}$
${}_{1}R1/19/5c_5E1/16/19/5P5p_{5-1}R1/19/5c_5E1/16/19/5p_{5-1}R1/19/5p_{5-1}R1/19/5p_{5-1}R1/19/5p_{5-1}R1/19/5p_{5-}$
${}_{1}R1/19/5E1/16/19/5P5c_5A1/16/19/5R1/19/5p_{5-1}R1/19/5c_5E1/16/19/5P5p_{5-1}R1/19/5p_{5-}$
${}_{1}R1/19/5c_5E1/16/19/5P5p_{5-1}R1/19/5p_{5-1}R1/19/5c_5E1/16/19/5P5p_{5-1}R1/19/5p_{5-}$
${}_{1}R1/19/5c_5E1/16/19/5P5p_{5-1}R1/19/5p_{5-1}R1/19/5c_5E1/16/19/5P5p_{5-1}R1/19/5p_{5-1}R1/19/5p_{5-}$
${}_{1}R1/19/5c_5E1/16/19/5P5p_{5-1}R1/19/5p_{5-1}R1/19/5c_5E1/16/19/5P5p_{5-1}R1/19/5p_{5-1}R1/19/5p_{5-}$
${}_{1}R1/19/5c_5E1/16/19/5P5p_{5-1}R1/19/5p_{5-1}R1/19/5c_5E1/16/19/5P5p_{5-1}R1/19/5p_{5-1}R1/19/5p_{5-}$
${}_{1}R1/19/5c_5E1/16/19/5P5p_{5-1}R1/19/5p_{5-1}R1/19/5c_5E1/16/19/5P5p_{5-1}R1/19/5p_{5-1}R1/19/5p_{5-}$
${}_{1}R1/19/5c_5E1/16/19/5P5p_{5-1}R1/19/5p_{5-1}R1/19/5p_{5-1}R1/19/5c_5E1/16/19/5P5p_{5-1}R1/19/5p_{5-}$
${}_{1}R1/19/5p_{5-1}R1/19/5c_5E1/16/19/5P5p_{5-1}R1/19/5p_{5-1}R1/19/5p_{5-1}R1/19/5c_5E1/16/19/5P5p_{5-}$
${}_{1}R1/19/5p_{5-1}R1/19/5p_{5-1}R1/19/5c_5E1/16/19/5P5p_{5-1}R1/19/5p_{5-1}R1/19/5p_{5-}$
${}_{1}R1/19/5c_5E1/16/19/5P5p_{5-1}R1/19/5p_{5-1}R1/19/5c_5E1/16/19/5P5p_{5-3}P5p_{5-1}R1/19/5p_{5-}$
${}_{1}R1/19/5A1/16/19/5R1/19/5c_5E1/16/19/5P5p_{5-1}R1/19/5p_{5-1}R1/19/5p_{5-3}P5p_{5-1}R1/19/5p_{5-}$
${}_{1}R1/19/5A1/16/19/5R1/19/5p_{5-1}R1/19/5p_{5-3}P5p_{5-1}R1/19/5p_{5-1}R1/19/5p_{5-}$
${}_{1}R1/19/5c_5E1/16/19/5P5p_{5-1}R1/19/5c_5E1/16/19/5P5p_{5-1}R1/19/5c_5p_{5-1}R1/19/5c_5E1/16/19/5P5p_{5-}$
${}_{6}c_5^{3}p_{5-1}R1/19/5c_5E1/16/19/5P5p_{5-6}c_5p_{5-1}R1/19/5c_5^{2}p_{5-1}R1/19/5c_5E1/16/19/5P5p_{5-}$
${}_{1}R1/19/5c_5^{2}E1/16/19/5P5p_{5-1}R1/19/5c_5p_{5-1}R1/19/5p_{5-1}R1/19/5c_5p_{5-1}p_{6-1}R1/19/5p_{5-1}R1/19/5p_{5-}$
${}_{1}R1/19/5c_5p_{5-1}p_{6-1}R1/19/5p_{5-1}R1/19/5p_{5-1}R1/19/5p_{5-1}R1/19/5c_5p_{5-1}p_{6-1}R1/19/5p_{5-1}R1/19/5p_{5-}$
${}_{1}R1/19/5c_5p_{5-1}p_{6-1}R1/19/5p_{5-1}R1/19/5p_{5-1}R1/19/5c_5E1/16/19/5P5c_5E1/16/19/5P5p_{5-1}R1/19/5p_{5-}$
${}_{1}R1/19/5p_{5-1}R1/19/5c_5E1/16/19/5P5p_{5-1}R1/19/5p_{5-1}R1/19/5c_5p_{5-1}R1/19/5c_5E1/16/19/5P5p_{5-}$
${}_{1}R1/19/5p_{5-1}R1/19/5p_{5-1}R1/19/5c_5E1/16/19/5P5p_{5-1}R1/19/5c_5p_{5-1}R1/19/5c_5E1/16/19/5P5p_{5-6}c_5p_{5-}$
${}_{1}R1/19/5c_5E1/16/19/5P5c_5E1/16/19/5P5p_{5-6}c_5p_{5-1}R1/19/5c_5E1/16/19/5P5c_5E1/16/19/5P5p_{5-}$
${}_{6}c_5E1/16/19/5P5p_{5-6}c_5p_{5-1}R1/19/5c_5E1/16/19/5P5c_5p_{5-1}R1/19/5c_5E1/16/19/5P5c_5E1/16/19/5P5p_{5-}$
${}_{6}c_5p_{5-1}R1/19/5c_5E1/16/19/5P5c_5E1/16/19/5P5p_{5-1}R1/19/5c_5E1/16/19/5P5p_{5-1}R1/19/5c_5p_{5-}$
${}_{1}R1/19/5p_{5-3}P5c_5E1/16/19/5P5c_5E1/16/19/5P5p_{5-6}c_5p_{5-}$
${}_{1}R1/19/5c_5E1/16/19/5P5c_5E1/16/19/5P5p_{5-1}R1/19/5c_5E1/16/19/5P5p_{5-1}R1/19/5c_5^{2}p_{5-}$
${}_{1}R1/19/5c_5p_{5-1}R1/19/5p_{5-1}R1/19/5c_5E1/16/19/5P5p_{5-1}R1/19/5p_{5-1}R1/19/5p_{5-}$
${}_{1}R1/19/5c_5E1/16/19/5P5p_{5-1}R1/19/5c_5p_{5-1}R1/19/5p_{5-1}R1/19/5c_5E1/16/19/5P5p_{5-1}R1/19/5p_{5-}$
${}_{1}R1/19/5p_{5-1}R1/19/5c_5E1/16/19/5P5p_{5-1}R1/19/5p_{5-1}R1/19/5p_{5-1}R1/19/5c_5E1/16/19/5P5p_{5-}$

$_1$R1/19/5p$_{5-1}$R1/19/5c$_5$p$_{5-1}$R1/19/5c$_5$E1/16/19/5P5p$_{5-1}$R1/19/5c$_5$p$_{5-1}$R1/19/5c$_5$E1/16/19/5P5p$_{5-}$
$_1$R1/19/5c$_5$E1/16/19/5P5p$_{5-1}$R1/19/5E1/16/19/5$^2$P5c$_5$E1/16/19/5P5p$_{5-}$
$_1$R1/19/5c$_5$E1/16/19/5P5c$_5$E1/16/19/5P5p$_{5-1}$R1/19/5c$_5$E1/16/19/5P5p$_{5-}$
$_1$R1/19/5E1/16/19/5$^2$P5c$_5$E1/16/19/5P5p$_{5-1}$R1/19/5c$_5$E1/16/19/5P5p$_{5-1}$R1/19/5p$_{5-1}$R1/19/5p$_{5-}$
$_1$R1/19/5c$_5$p$_{5-1}$R1/19/5c$_5$E1/16/19/5P5p$_{5-1}$R1/19/5c$_5$p$_{5-1}$R1/19/5c$_5$E1/16/19/5P5p$_{5-}$
$_1$R1/19/5c$_5$E1/16/19/5P5p$_{5-1}$R1/19/5E1/16/19/5$^2$P5c$_5$E1/16/19/5P5p$_{5-}$
$_1$R1/19/5c$_5$E1/16/19/5P5E1/16/19/5P5p$_{5-}$
$_1$R1/19/5c$_5$E1/16/19/5P5c$_5$E1/16/19/5P5E1/16/19/5P5p$_{5-2}$A1/16/19/5P5p$_{5-}$
$_1$R1/19/5c$_5$E1/16/19/5P5p$_{5-1}$R1/19/5E1/16/19/5$^2$P5c$_5$E1/16/19/5P5p$_{5-}$
$_1$R1/19/5c$_5$E1/16/19/5P5E1/16/19/5P5p$_{5-}$
$_1$R1/19/5c$_5$E1/16/19/5P5c$_5$E1/16/19/5P5E1/16/19/5P5p$_{5-2}$A1/16/19/5P5c$_5$E1/16/19/5P5p$_{5-}$
$_1$R1/19/5p$_{5-1}$R1/19/5c$_5$E1/16/19/5P5p$_{5-}$
$_2$A1/16/19/5R1/19/5c$_5$E1/16/19/5P5c$_5$E1/16/19/5P5c$_5$E1/16/19/5P5p$_{5-}$
$_1$R1/19/5c$_5$E1/16/19/5P5E1/16/19/5P5p$_{5-1}$R1/19/5c$_5$E1/16/19/5P5p$_{5-1}$R1/19/5c$_5$E1/16/19/5P5p$_{5-}$
$_1$R1/19/5c$_5$E1/16/19/5P5E1/16/19/5P5p$_{5-1}$R1/19/5c$_5$E1/16/19/5P5p$_{5-1}$R1/19/5p$_{5-}$
$_1$R1/19/5c$_5$$^2$E1/16/19/5P5p$_{5-1}$R1/19/5c$_5$E1/16/19/5P5c$_5$E1/16/19/5P5p$_{5-}$
$_1$R1/19/5c$_5$E1/16/19/5P5p$_{5-}$
$_1$R1/19/5c$_5$E1/16/19/5P5E1/16/19/5P5c$_5$E1/16/19/5P5c$_5$E1/16/19/5P5c$_5$E1/16/19/5P5p$_{5-}$
$_1$R1/19/5c$_5$E1/16/19/5P5E1/16/19/5P5p$_{5-1}$R1/19/5c$_5$E1/16/19/5P5p$_{5-1}$R1/19/5c$_5$E1/16/19/5P5p$_{5-}$
$_1$R1/19/5c$_5$E1/16/19/5P5E1/16/19/5P5p$_{5-1}$R1/19/5c$_5$E1/16/19/5P5p$_{5-1}$R1/19/5p$_{5-}$
$_1$R1/19/5c$_5$$^2$E1/16/19/5P5p$_{5-1}$R1/19/5c$_5$E1/16/19/5P5c$_5$E1/16/19/5P5p$_{5-}$
$_1$R1/19/5c$_5$E1/16/19/5P5p$_{5-1}$R1/19/5c$_5$E1/16/19/5P5E1/16/19/5P5c$_5$E1/16/19/5P5p$_{5-}$
$_1$R1/19/5c$_5$E1/16/19/5P5E1/16/19/5p$_{5-}$
$_1$R1/19/5c$_5$E1/16/19/5P5c$_5$E1/16/19/5P5c$_5$E1/16/19/5P5E1/16/19/5P5p$_{5-1}$R1/19/5p$_{5-}$
$_1$R1/19/5c$_5$E1/16/19/5P5E1/16/19/5P5p$_{1-5}$P5p$_{5-1}$R1/19/5E1/16/19/5P5c$_5$E1/16/19/5P5p$_{5-}$
$_1$R1/19/5c$_5$E1/16/19/5P5c$_5$E1/16/19/5P5c$_5$E1/16/19/5P5p$_{5-1}$R1/19/5p$_{1-5}$P5c$_5$$^2$E1/16/19/5P5c$_5$p$_{5-}$
$_1$R1/19/5c$_5$E1/16/19/5P5E1/16/19/5R1/19/5c$_5$E1/16/19/5P5p$_{5-1}$R1/19/5c$_5$E1/16/19/5P5c$_5$p$_{1-}$
$_5$P5p$_{5-1}$R1/19/5c$_5$E1/16/19/5P5p$_{5-1}$R1/19/5c$_5$$^2$p$_{1-5}$P5p$_{5-1}$R1/19/5c$_5$E1/16/19/5P5E1/16/19/5P5p$_{5-}$
$_1$R1/19/5c$_5$E1/16/19/5P5p$_{5-1}$R1/19/5p$_{5-1}$R1/19/5c$_5$E1/16/19/5P5c$_5$E1/16/19/5P5p$_{5-1}$R1/19/5c$_5$p$_{5-}$
$_1$R1/19/5c$_5$E1/16/19/5P5p$_{5-1}$R1/19/5c$_5$E1/16/19/5P5p$_{5-1}$R1/19/5c$_5$E1/16/19/5P5p$_{5-}$
$_1$R1/19/5c$_5$E1/16/19/5P5p$_{5-1}$R1/19/5c$_5$E1/16/19/5P5p$_{5-}$
$_1$R1/19/5c$_5$E1/16/19/5P5c$_5$E1/16/19/5P5c$_5$E1/16/19/5P5p$_{5-1}$R1/19/5c$_5$E1/16/19/5P5p$_{5-}$
$_1$R1/19/5c$_5$E1/16/19/5P5c$_5$E1/16/19/5P5p$_{5-}$
$_1$R1/19/5c$_5$E1/16/19/5P5c$_5$E1/16/19/5P5c$_5$E1/16/19/5P5p$_{5-}$
$_1$R1/19/5c$_5$$^2$E1/16/19/5P5c$_5$E1/16/19/5P5c$_5$E1/16/19/5P5p$_{5-}$
$_1$R1/19/5c$_5$E1/16/19/5P5c$_5$E1/16/19/5P5c$_5$E1/16/19/5P5p$_{5-1}$R1/19/5c$_5$E1/16/19/5P5p$_{5-}$
$_1$R1/19/5c$_5$E1/16/19/5P5c$_5$E1/16/19/5P5c$_5$E1/16/19/5P5p$_{5-}$
$_1$R1/19/5c$_5$E1/16/19/5P5c$_5$E1/16/19/5P5c$_5$E1/16/19/5P5c$_5$E1/16/19/5P5p$_{5-}$
$_1$R1/19/5c$_5$E1/16/19/5P5p$_{5-6}$p$_{3-}$
$_5$P5c$_5$E1/16/19/5P5c$_5$E1/16/19/5P5c$_5$E1/16/19/5P5c$_5$E1/16/19/5P5p$_{5-1}$R1/19/5p$_{5-}$
$_1$R1/19/5c$_5$E1/16/19/5P5p$_{5-1}$R1/19/5p$_{5-3}$P5p$_{5-1}$R1/19/5p$_{5-1}$R1/19/5c$_5$E1/16/19/5P5p$_{5-1}$R1/19/5p$_{5-}$
$_3$P5p$_{5-6}$p$_{5-1}$R1/19/5p$_{5-1}$R1/19/5c$_5$E1/16/19/5P5p$_{5-1}$R1/19/5c$_5$E1/16/19/5P5p$_{5-}$
$_1$R1/19/5c$_5$E1/16/19/5P5p$_{5-1}$R1/19/5c$_5$E1/16/19/5P5p$_{5-1}$R1/19/5c$_5$E1/16/19/5P5p$_{5-}$
$_1$R1/19/5c$_5$E1/16/19/5P5p$_{5-1}$R1/19/5c$_5$E1/16/19/5P5p$_{5-1}$R1/19/5p$_{5-1}$R1/19/5p$_{5-6}$p$_{3-}$
$_5$P5c$_5$E1/16/19/5P5p$_{5-1}$R1/19/5p$_{5-1}$R1/19/5c$_5$E1/16/19/5P5p$_{5-1}$R1/19/5c$_5$E1/16/19/5P5p$_{5-}$
$_1$R1/19/5c$_5$E1/16/19/5P5p$_{5-1}$R1/19/5c$_5$E1/16/19/5P5p$_{5-1}$R1/19/5c$_5$E1/16/19/5P5p$_{5-1}$R1/19/5p$_{5-}$
$_1$R1/19/5c$_5$E1/16/19/5P5p$_{5-1}$R1/19/5c$_5$E1/16/19/5P5p$_{5-1}$R1/19/5c$_5$E1/16/19/5P5p$_{5-}$
$_1$R1/19/5c$_5$E1/16/19/5P5p$_{5-1}$R1/19/5c$_5$E1/16/19/5P5p$_{5-1}$R1/19/5c$_5$E1/16/19/5P5p$_{5-1}$R1/19/5p$_{5-}$
$_1$R1/19/5c$_5$E1/16/19/5P5p$_{5-1}$R1/19/5p$_{5-1}$R1/19/5c$_5$E1/16/19/5P5p$_{5-1}$R1/19/5p$_{5-}$
$_1$A1/16/19/5R1/19/5c$_5$E1/16/19/5P5p$_{5-1}$R1/19/5p$_{5-1}$R1/19/5c$_5$E1/16/19/5P5p$_{5-}$
$_1$R1/19/5E1/16/19/5P5p$_{5-1}$R1/19/5c$_5$E1/16/19/5P5p$_{5-1}$R1/19/5E1/16/19/5P5c$_5$E1/16/19/5P5p$_{5-}$

$_{1}$R1/19/5E1/16/19/5P5p$_{5-1}$R1/19/5c$_{5}$E1/16/19/5P5p$_{5-1}$R1/19/5p$_{5-1}$R1/19/5c$_{5}$E1/16/19/5P5p$_{5-1}$R1/19/5E1/16/19/5P5c$_{5}$E1/16/19/5P5p$_{5-1}$R1/19/5E1/16/19/5P5p$_{5-1}$R1/19/5c$_{5}$E1/16/19/5P5p$_{5-1}$R1/19/5p$_{5-1}$R1/19/5c$_{5}$E1/16/19/5P5p$_{5-1}$R1/19/5E1/16/19/5P5p$_{5-1}$R1/19/5c$_{5}$E1/16/19/5P5p$_{5-1}$R1/19/5p$_{5-1}$R1/19/5c$_{5}$E1/16/19/5P5p$_{5-1}$R1/19/5E1/16/19/5P5p$_{5-1}$R1/19/5c$_{5}$E1/16/19/5P5E1/16/19/5$^{2}$P5p$_{5-1}$R1/19/5p$_{5-1}$R1/19/5c$_{5}$E1/16/19/5P5E1/16/19/5$^{2}$P5p$_{5-1}$R1/19/5c$_{5}$E1/16/19/5P5p$_{5-1}$R1/19/5E1/16/19/5P5E1/16/19/5R1/19/5c$_{5}$E1/16/19/5P5p$_{5-1}$R1/19/5E1/16/19/5P5c$_{5}$E1/16/19/5P5p$_{5-1}$R1/19/5E1/16/19/5P5p$_{5-1}$R1/19/5E1/16/19/5P5p$_{5-1}$R1/19/5c$_{5}$A1/16/19/5R1/19/5E1/16/19/5P5E1/16/19/5p$_{4-1}$R1/19/5E1/16/19/5P5p$_{5-2}$E1/16/19/5P5E1/16/19/5P5E1/16/19/5P5A1/16/19/5R1/19/5E1/16/19/5P5E1/16/19/5p$_{4-1}$R1/19/5E1/16/19/5P5p$_{5-2}$E1/16/19/5P5E1/16/19/5P5E1/16/19/5P5A1/16/19/5R1/19/5c$_{5}$E1/16/19/5P5p$_{5-3}$A1/16/19/5R1/19/5p$_{5-1}$R1/19/5p$_{5-1}$R1/19/5c$_{5}$E1/16/19/5P5A1/16/19/5R1/19/5p$_{5-1}$R1/19/5p$_{5-1}$R1/19/5p$_{5-1}$R1/19/5p$_{5-1}$R1/19/5p$_{5-1}$R1/19/5c$_{5}$p$_{5-3}$A1/16/19/5R1/19/5p$_{5-1}$R1/19/5p$_{5-6}$C5$^{2}$E1/16/19/5C5E1/16/19/5P5A1/16/19/5p$_{3-5}$P5E1/16/19/5P5E1/16/19/5p$_{2-3}$C5E1/16/19/5C5E1/16/19/5P5$^{2}$B5E1/16/19/5p$_{1-3}$

### cen6

D6F6K6p$_{11-12}$c$_{16}^{7}$c$_{17}^{21}$Q6c$_{1}^{16}$p$_{1-9}$p$_{5-11}$c$_{14}^{3}$p$_{14-11}$c$_{15}^{2}$p$_{15-11}$c$_{15}$p$_{15-11}$p$_{14-11}^{2}$c$_{14}^{6}$p$_{14-9}$p$_{5-11}$c$_{14}^{4}$p$_{14-11}$p$_{13-11}$p$_{14-11}^{2}$c$_{14}^{8}$p$_{14-11}^{3}$c$_{14}^{44}$p$_{14-12}$p$_{17-12}$c$_{17}$p$_{17-12}$c$_{17}^{15}$p$_{17-12}$p$_{16-12}$c$_{16}^{2}$p$_{16-12}^{2}$c$_{16}$p$_{16-12}^{2}$c$_{16}$p$_{16-12}$c$_{16}^{11}$p$_{16-14}$c$_{18}^{6}$p$_{18-12}$c$_{16}^{20}$p$_{16-12}$c$_{17}$p$_{17-12}$c$_{17}$p$_{17-12}$c$_{17}$p$_{17-12}$c$_{16}$p$_{16-12}^{2}$p$_{17-12}$c$_{16}^{3}$p$_{16-12}$c$_{16}$p$_{16-12}$p$_{17-12}^{2}$p$_{15-12}$c$_{16}^{3}$p$_{16-12}^{2}$p$_{17-12}$c$_{16}^{2}$p$_{16-12}$c$_{16}^{3}$p$_{16-12}$c$_{17}^{3}$p$_{17-12}$c$_{16}$p$_{16-12}$c$_{16}^{3}$p$_{16-17}$p$_{11-12}$p$_{16-12}$c$_{16}^{5}$p$_{16-12}$c$_{16}^{12}$p$_{16-12}$c$_{16}^{2}$p$_{16-12}$c$_{16}^{12}$p$_{16-14}$p$_{18-12}$p$_{16-12}$c$_{17}$p$_{17-12}$p$_{16-12}$c$_{17}^{2}$p$_{17-12}$p$_{16-12}$p$_{17-12}$c$_{17}$p$_{17-12}$c$_{17}$p$_{17-12}$p$_{16-12}$c$_{16}$p$_{16-12}^{3}$c$_{16}^{2}$p$_{16-12}^{4}$p$_{17-12}$p$_{16-12}^{3}$c$_{16}$p$_{16-12}^{3}$c$_{16}^{2}$p$_{16-12}$p$_{16-7}$p$_{15-12}$c$_{16}$p$_{16-12}^{2}$p$_{17-12}$c$_{16}$p$_{16-12}^{2}$p$_{17-12}^{2}$p$_{16-12}$c$_{17}$p$_{17-12}^{2}$p$_{16-12}$c$_{17}$p$_{17-12}$p$_{16-12}$c$_{17}$p$_{17-12}$c$_{16}^{2}$p$_{16-12}^{2}$p$_{17-12}$p$_{16-13}$c$_{17}^{3}$p$_{17-12}$p$_{16-12}$c$_{16}$p$_{16-12}^{2}$c$_{16}^{3}$p$_{16-12}^{2}$c$_{16}^{2}$p$_{16-12}^{2}$c$_{16}$p$_{16-12}$c$_{16}^{2}$p$_{16-12}$c$_{16}^{2}$p$_{16-12}$c$_{16}^{2}$p$_{16-12}$c$_{16}^{2}$p$_{16-12}$p$_{17-12}$p$_{16-12}^{2}$c$_{16}^{4}$p$_{16-12}^{3}$c$_{16}^{2}$p$_{16-12}^{2}$c$_{16}$p$_{16-12}$c$_{16}$p$_{16-12}$c$_{16}^{2}$p$_{16-12}$c$_{16}$p$_{16-12}^{2}$c$_{16}$p$_{16-12}^{2}$c$_{16}$p$_{16-12}$c$_{16}$p$_{16-12}$c$_{16}$p$_{16-12}^{2}$c$_{16}$p$_{16-12}$c$_{16}^{2}$p$_{16-12}$c$_{16}$p$_{16-12}$c$_{16}^{22}$p$_{16-12}^{2}$p$_{17-12}^{2}$p$_{16-12}$p$_{17-12}^{2}$p$_{16-12}^{4}$c$_{17}$p$_{17-12}$p$_{16-12}$p$_{17-12}^{2}$c$_{17}^{2}$p$_{17-12}$p$_{16-12}$p$_{17-12}$c$_{17}$p$_{17-12}$p$_{16-12}$c$_{16}$p$_{16-12}^{3}$c$_{16}^{2}$p$_{16-12}^{6}$p$_{17-12}$p$_{16-12}^{3}$c$_{16}$p$_{16-12}^{3}$c$_{16}^{4}$p$_{16-12}$c$_{17}$p$_{17-12}$c$_{16}^{2}$p$_{16-12}^{2}$p$_{16-13}$c$_{17}^{3}$p$_{17-12}$p$_{16-12}$c$_{16}$p$_{16-12}$c$_{16}^{2}$p$_{16-12}$c$_{16}^{2}$p$_{16-12}^{2}$c$_{16}^{2}$p$_{16-12}$c$_{16}^{3}$p$_{16-12}^{3}$c$_{16}$p$_{16-12}^{2}$c$_{16}$p$_{16-12}$c$_{16}$p$_{16-12}$c$_{16}^{2}$p$_{16-12}$c$_{16}$p$_{16-12}^{2}$c$_{16}$p$_{16-12}$c$_{16}$p$_{16-12}^{3}$c$_{16}$p$_{16-12}$c$_{16}$p$_{16-12}$c$_{16}$p$_{16-12}$c$_{16}$p$_{16-12}$c$_{16}$p$_{16-12}$c$_{16}$p$_{16-12}$c$_{16}^{6}$p$_{16-12}^{2}$c$_{16}$p$_{16-12}$c$_{16}$p$_{16-12}$c$_{17}$p$_{17-12}$c$_{17}^{7}$p$_{17-12}$p$_{16-12}^{2}$c$_{16}$p$_{16-12}^{4}$c$_{16}^{3}$p$_{16-12}$c$_{16}^{4}$p$_{16-12}$c$_{16}^{4}$p$_{16-12}$c$_{16}$p$_{16-12}$c$_{16}^{19}$p$_{16-12}$c$_{16}$p$_{16-12}$c$_{16}^{4}$p$_{16-12}$c$_{16}^{2}$p$_{16-12}$c$_{16}^{5}$p$_{16-12}$c$_{16}^{6}$p$_{16-12}^{2}$c$_{16}$p$_{16-12}$c$_{17}^{5}$p$_{17-12}$c$_{16}$p$_{16-12}$c$_{16}^{23}$p$_{16-12}$c$_{16}$p$_{16-12}$c$_{16}^{4}$p$_{16-12}$c$_{16}$p$_{16-12}$c$_{16}^{2}$p$_{16-12}$c$_{16}$p$_{16-12}$c$_{16}^{8}$p$_{16-12}$c$_{16}$p$_{16-12}$c$_{16}^{4}$p$_{16-12}$c$_{16}^{2}$p$_{16-12}$c$_{16}^{5}$p$_{16-12}$c$_{16}^{6}$p$_{16-12}$c$_{16}$p$_{16-12}$c$_{16}^{5}$p$_{16-12}$c$_{16}$p$_{16-12}$c$_{16}^{24}$p$_{16-12}$c$_{16}$p$_{16-12}$c$_{16}^{4}$p$_{16-12}$c$_{16}$p$_{16-12}$c$_{16}^{3}$p$_{16-12}$c$_{16}$p$_{16-12}$c$_{16}^{2}$p$_{16-12}$c$_{16}$p$_{16-12}$c$_{16}$p$_{16-12}$c$_{16}^{4}$p$_{16-14}$c$_{18}^{2}$p$_{18-12}$c$_{16}$p$_{16-12}$c$_{16}^{2}$p$_{16-12}^{2}$c$_{16}^{3}$p$_{16-12}$c$_{16}^{23}$p$_{16-12}$c$_{16}^{6}$p$_{16-12}$c$_{16}$p$_{16-12}$c$_{16}^{6}$p$_{16-12}$c$_{16}$p$_{16-12}$c$_{17}^{3}$p$_{17-12}$c$_{16}^{3}$p$_{16-12}$p$_{17-12}$c$_{16}$p$_{16-12}$c$_{16}$p$_{16-12}$c$_{17}^{3}$p$_{17-12}$c$_{16}$p$_{16-12}$c$_{16}$p$_{16-14}$c$_{18}$p$_{18-12}$p$_{16-12}^{3}$c$_{16}^{4}$p$_{16-12}$c$_{16}$p$_{16-12}$c$_{16}^{6}$p$_{16-12}$c$_{16}^{3}$p$_{16-12}$c$_{16}^{3}$p$_{16-12}^{2}$p$_{16-17}$c$_{15}^{15}$p$_{15-12}$c$_{16}$p$_{16-12}$c$_{16}$p$_{16-10}$L6c$_{16}^{2}$p$_{16-12}$c$_{16}^{5}$p$_{16-12}$c$_{16}$p$_{16-12}$c$_{16}^{2}$p$_{16-10}$L6c$_{16}^{8}$p$_{16-12}^{4}$c$_{16}^{4}$p$_{16-12}^{2}$c$_{16}^{5}$p$_{16-12}^{4}$c$_{16}$p$_{16-12}$c$_{16}^{3}$p$_{16-12}^{4}$c$_{16}$p$_{16-12}$c$_{16}^{2}$p$_{16-12}$c$_{16}$p$_{16-12}$c$_{17}$p$_{17-15}$

### cen7

A7p$_{2-3}$E7A7p$_{3-5}$E7$^{2}$F7A7p$_{5-6}$p$_{2-3}$c$_{3}$C7c$_{5}$p$_{5-3}$C7p$_{5-6}$p$_{2-3}$p$_{3-6}$A7C7F7p$_{2-3}$C7A7C7p$_{5-6}$p$_{2-3}$c$_{3}$C7c$_{3}^{2}$C7c$_{3}^{6}$C7c$_{3}^{3}$C7c$_{3}$C7c$_{3}$C7p$_{3-1}$c$_{3}^{3}$C7c$_{3}^{2}$C7c$_{3}^{2}$C7$^{2}$A7C7p$_{5-6}$p$_{2-3}$c$_{3}$C7c$_{3}$C7c$_{3}$C7c$_{3}$C7c$_{3}$p$_{3-6}$p$_{2-3}$c$_{3}$C7c$_{3}^{2}$C7c$_{3}^{3}$C7c$_{3}$C7c$_{3}^{3}$C7c$_{3}$C7c$_{3}^{3}$C7c$_{3}^{3}$C7c$_{3}$C7c$_{3}^{14}$C7c$_{6}^{3}$p$_{6-3}$c$_{6}^{3}$p$_{6-3}$c$_{6}$p$_{6-4}$c$_{1}^{64}$A7c$_{4}^{59}$c$_{6}^{4}$p$_{6-3}$c$_{6}^{3}$p$_{6-3}$c$_{6}^{10}$p$_{6-3}$c$_{6}^{6}$p$_{6-3}$c$_{6}^{9}$p$_{6-3}$p$_{6-1}$c$_{4}^{4}$c$_{6}^{8}$p$_{6-3}$p$_{6-1}$c$_{4}^{4}$c$_{6}^{3}$p$_{6-3}$c$_{6}^{5}$p$_{6-3}$c$_{6}^{6}$p$_{6-3}$c$_{6}^{29}$p$_{6-3}$c$_{6}^{4}$p$_{6-3}$c$_{6}^{2}$p$_{6-3}$c$_{6}^{18}$p$_{6-3}$c$_{6}^{4}$F7c$_{3}^{181}$p$_{3-1}$c$_{4}^{2}$p$_{4-1}$c$_{4}^{58}$p$_{4-1}$c$_{4}^{2}$p$_{4-1}$c$_{4}^{40}$p$_{4-2}$c$_{5}^{9}$p$_{5-1}$c$_{6}^{2}$p$_{6-1}$c$_{4}^{2}$p$_{4-1}$c$_{4}^{7}$p$_{4-1}$c$_{4}^{25}$p$_{4-1}$c$_{6}^{5}$p$_{6-1}$c$_{6}^{4}$p$_{6-1}$c$_{6}^{25}$p$_{6-1}$c$_{4}$p$_{4-1}$c$_{4}^{42}$p$_{4-1}$c$_{4}^{28}$p$_{4-1}$c$_{6}^{8}$p$_{6-1}$c$_{6}^{103}$p$_{6-1}$c$_{4}^{4}$p$_{4-1}$c$_{4}^{36}$p$_{4-2}$c$_{5}^{4}$p$_{5-2}$c$_{5}^{6}$p$_{5-2}$c$_{5}^{4}$p$_{5-2}$c$_{5}^{7}$p$_{5-2}$c$_{5}^{3}$p$_{5-2}$c$_{5}^{9}$p$_{5-2}$c$_{5}^{6}$p$_{5-2}$c$_{5}^{8}$p$_{5-2}$c$_{5}^{271}$p$_{5-3}$c$_{2}^{6}$p$_{2-3}$c$_{5}^{5}$p$_{5-3}$c$_{2}^{4}$p$_{2-3}$c$_{5}^{8}$p$_{5-3}$c$_{2}^{4}$p$_{2-3}$c$_{5}^{5}$p$_{5-3}$c$_{2}^{4}$p$_{2-3}$c$_{5}^{157}$E7c$_{2}$p$_{2-4}$p$_{3-5}$c$_{2}^{2}$p$_{2-5}$c$_{2}^{4}$p$_{2-5}$c$_{2}^{4}$p$_{2-5}$c$_{2}^{4}$p$_{2-5}$c$_{2}^{4}$p$_{2-5}$c$_{2}^{4}$p$_{2-5}$c$_{2}^{6}$p$_{2-5}$c$_{2}^{6}$p$_{2-5}$c$_{2}^{15}$p$_{2-5}$c$_{2}^{3}$p$_{2-5}$p$_{2-4}$p$_{3-5}$c$_{2}^{4}$p$_{2-5}$c$_{2}^{3}$p$_{2-5}$p$_{2-4}$p$_{3-5}$c$_{2}^{14}$p$_{2-5}$c$_{2}^{5}$p$_{2-5}$c$_{2}^{11}$p$_{2-5}$c$_{2}^{11}$p$_{2-5}$c$_{2}^{4}$p$_{2-5}$p$_{2-4}$p$_{3-5}$c$_{2}^{18}$p$_{2-5}$c$_{2}^{11}$p$_{2-5}$c$_{2}^{11}$p$_{2-5}$c$_{2}^{26}$c$_{4}^{4}$p$_{4-1}$c$_{4}$p$_{4-1}$c$_{4}^{17}$c$_{2}^{4}$p$_{2-3}$c$_{2}^{8}$p$_{2-3}$c$_{2}^{11}$p$_{2-3}$c$_{2}^{23}$p$_{2-3}$c$_{2}^{63}$c$_{4}^{27}$p$_{4-1}$c$_{4}^{24}$p$_{4-2}$c$_{5}^{18}$p$_{5-2}$c$_{5}^{6}$p$_{5-2}$c$_{5}^{5}$p$_{5-2}$c$_{5}$p$_{5-}$

${}_2c_5^{43}c_3^9p_{3-4}c_3c_5^5p_{5-2}c_5^3p_{5-2}c_5^{10}p_{5-2}c_5^{18}p_{5-2}c_5^{14}p_{5-2}c_5^{39}c_3c_5^5p_{5-2}c_5^8p_{5-2}c_5^2p_{5-2}c_5^4p_{5-2}c_5^5p_{5-2}c_5^2p_{5-2}c_5^2p_{5-}$
${}_2c_5^2p_{5-2}c_5^5p_{5-2}c_5^{20}p_{5-1}c_4^{14}p_{4-2}c_5^7p_{5-2}c_5^{23}p_{5-2}c_5^{20}p_{5-2}c_5^{82}p_{5-1}c_1^4A7c_1^{244}A7c_4^{242}p_{4-1}c_4^{117}p_{4-1}c_1^{153}p_{1-5}c_2^{57}p_{2-}$
${}_4c_2^5p_{2-3}c_2p_{2-4}c_2p_{2-4}c_2p_{2-4}^2c_2p_{2-3}c_2p_{2-4}c_2p_{2-3}p_{2-4}c_2p_{2-4}c_2p_{3-4}c_2^5p_{2-6}B7c_4^2p_{4-6}c_2^2p_{2-6}c_2p_{2-4}c_2^2p_{2-4}c_2^2p_{2-4}c_2p_{2-}$
${}_4c_2p_{3-4}c_3^3p_{3-4}c_2p_{2-4}p_{2-3}c_3p_{3-4}B7c_2p_{2-4}c_2^2p_{3-1}c_3^2p_{3-4}p_{2-4}c_2^3p_{2-4}c_2p_{3-4}p_{2-4}c_2^3p_{2-4}c_2p_{2-3}p_{5-1}D7c_2^2p_{2-4}^2c_2B7p_{5-}$
${}_2p_{2-4}c_2^2p_{2-4}^3c_2^3p_{2-4}^2c_2p_{2-3}c_5p_{2-4}p_{3-4}c_2^2B7p_{2-4}c_2p_{2-4}^2c_2^3p_{2-4}c_3^4p_{3-4}p_{2-4}c_2^4p_{2-3}B7c_5p_{2-3}c_2^3p_{2-4}p_{2-3}c_6^6p_{6-4}p_{2-}$
${}_3p_{6-2}c_5^2c_2^2p_{2-4}c_2p_{2-4}^2c_2p_{2-5}c_2p_{2-3}c_2p_{2-4}B7c_5^3c_2p_{2-4}c_2p_{2-4}c_2^2p_{2-4}c_2p_{2-4}p_{2-5}c_2^2D7c_2p_{2-4}c_2^2p_{2-4}c_2^2p_{2-3}p_{3-}$
${}_1C7c_5p_{3-1}C7p_{5-2}c_2p_{2-3}p_{3-4}F7p_{2-3}p_{5-1}C7$

### cen8

$c_4^{17}D8c_5^2p_{5-1}c_1^{10}p_{1-4}p_{5-3}c_5p_{5-1}c_1^{16}p_{1-7}p_{1-3}L8c_8c_1p_{1-3}L8p_{8-3}L8c_8p_{1-3}L8c_8p_{1-7}p_{1-3}L8c_8p_{1-7}p_{1-3}L8c_8p_{1-}$
${}_3L8c_8p_{1-7}p_{1-3}L8c_8p_{1-7}p_{1-3}L8p_{8-3}L8c_8p_{1-7}p_{1-3}L8p_{8-3}L8c_8p_{1-3}L8p_{8-3}L8c_8^7p_{1-3}L8c_8p_{8-3}L8c_8p_{1-3}L8c_8p_{1-}$
${}_3L8c_8p_{1-7}^2p_{1-3}L8c_8p_{1-7}p_{1-3}L8p_{8-3}L8c_8p_{8-3}L8c_8p_{1-3}L8c_8p_{1-3}L8c_8p_{1-3}L8p_{8-3}L8c_8^3p_{1-7}^2p_{1-3}L8c_8p_{1-3}L8c_8^3p_{1-}$
${}_3L8c_8^4p_{1-3}L8p_{8-3}L8p_{8-3}L8p_{8-3}L8p_{8-3}L8p_{8-3}L8p_{8-3}L8p_{8-3}L8p_{8-3}L8p_{8-3}L8p_{8-1}p_{2-3}L8p_{8-3}L8p_{8-3}L8p_{8-1}p_{2-}$
${}_3L8p_{8-3}L8p_{8-3}L8p_{8-3}L8p_{8-3}L8p_{8-3}L8p_{8-3}L8p_{8-3}L8p_{8-3}L8p_{8-3}L8p_{8-3}L8c_8p_{8-3}L8p_{8-3}L8p_{8-3}L8p_{8-3}L8p_{8-3}L8p_{8-}$
${}_3L8p_{8-3}L8c_8^3p_{10-3}p_{5-7}p_{1-7}^2p_{1-3}L8c_8p_{1-3}L8p_{8-3}L8c_8p_{1-3}L8p_{8-3}L8c_8^2p_{1-3}L8c_8^2p_{1-3}L8c_8^2p_{1-3}L8c_8^2p_{1-3}L8p_{8-}$
${}_3L8c_8^2p_{1-3}L8p_{8-3}L8p_{8-3}L8p_{8-3}L8p_{8-3}L8p_{8-3}L8c_8p_{8-3}L8c_8p_{8-3}L8p_{8-3}L8c_8p_{8-3}L8p_{8-3}L8c_8^2p_{1-3}L8c_8p_{1-3}L8p_{8-}$
${}_3L8c_8p_{1-3}L8c_8p_{1-3}L8c_8p_{1-3}L8c_8^2p_{1-3}L8c_8p_{1-3}L8c_8^3p_{1-3}L8c_8p_{1-3}L8c_8p_{1-3}L8c_8p_{1-3}L8c_8p_{1-3}L8c_8p_{1-3}L8c_8p_{1-}$
${}_3L8p_{8-3}L8c_8p_{1-3}L8c_8^2p_{1-3}L8c_8p_{1-3}L8c_8p_{1-3}L8c_8p_{1-3}L8c_8p_{1-3}L8p_{8-3}L8c_8p_{1-3}L8c_8p_{1-3}L8p_{8-3}L8c_8p_{1-3}L8c_8p_{1-}$
${}_3L8c_8p_{1-3}L8p_{8-3}L8c_8p_{1-3}L8c_8p_{1-3}L8p_{8-3}L8p_{8-3}L8c_8p_{1-3}L8p_{8-3}L8p_{8-3}L8c_8p_{1-3}L8c_8p_{1-3}L8p_{8-3}L8p_{8-3}L8p_{8-}$
${}_3L8p_{8-3}L8p_{8-3}L8p_{8-3}L8p_{8-3}L8p_{8-3}L8p_{8-3}L8p_{8-3}L8p_{8-3}L8p_{8-3}L8p_{8-3}L8p_{8-3}L8p_{8-3}L8p_{8-3}L8c_8p_{1-3}L8c_8p_{1-7}p_{1-}$
${}_3L8c_8^2p_{1-3}L8c_8p_{1-3}L8c_8p_{1-3}L8p_{8-3}L8c_8p_{1-3}L8c_8p_{1-3}L8c_8p_{1-3}L8p_{8-3}L8p_{8-3}L8p_{8-3}L8c_8^2p_{1-7}^2p_{1-3}p_{5-7}p_{1-7}p_{1-}$
${}_3L8p_{8-3}L8c_8p_{1-3}L8p_{8-3}L8p_{8-3}L8p_{8-3}L8p_{8-3}L8c_8p_{1-3}L8p_{8-3}L8p_{8-3}L8c_8p_{1-3}L8p_{8-3}L8p_{8-3}L8p_{8-3}L8p_{8-3}L8p_{8-}$
${}_3L8p_{8-3}L8p_{8-3}L8p_{8-3}L8p_{8-3}L8p_{8-3}L8p_{8-3}L8p_{8-3}L8c_8p_{1-3}L8p_{8-3}L8p_{8-3}L8c_8p_{1-3}L8p_{8-3}L8p_{8-3}L8p_{8-3}L8p_{8-}$
${}_3L8p_{8-3}L8p_{8-3}L8c_8p_{1-3}L8p_{8-3}L8p_{8-3}L8p_{8-3}L8p_{8-3}L8p_{8-3}L8c_8p_{1-3}L8p_{8-3}L8p_{8-3}L8p_{8-3}L8p_{8-3}L8c_8p_{1-3}L8c_8p_{1-}$
${}_7p_{1-3}L8c_8p_{1-7}p_{1-3}L8c_8p_{2-7}p_{1-3}L8p_{8-3}L8p_{8-3}L8c_8p_{1-7}p_{1-3}L8p_{8-3}L8p_{8-3}L8c_8p_{1-3}L8p_{8-3}L8p_{8-3}L8c_8p_{1-3}L8c_8p_{1-}$
${}_3L8p_{8-3}L8c_8p_{1-3}L8p_{8-3}L8c_8p_{1-3}L8p_{8-3}L8c_8p_{1-3}L8c_8p_{1-3}L8c_8p_{1-3}L8c_8p_{1-3}L8c_8p_{1-3}L8c_8p_{1-3}L8c_8p_{1-3}L8c_8p_{1-}$
${}_3L8c_8p_{1-3}L8c_8p_{1-3}L8c_8p_{1-3}L8c_8p_{1-3}L8c_8p_{1-3}L8c_8p_{1-3}L8c_8p_{1-3}L8c_8p_{1-3}L8p_{8-3}L8c_8p_{1-3}L8p_{8-3}L8p_{8-3}L8c_8p_{1-}$
${}_3L8c_8p_{1-3}G8p_{1-3}L8c_8p_{1-3}L8c_8p_{1-3}L8c_8p_{1-3}L8c_8p_{1-3}G8p_{1-3}L8c_8p_{1-3}L8p_{8-3}L8p_{8-3}L8c_8p_{1-3}L8p_{8-3}L8c_8p_{1-}$
${}_3L8p_{8-3}L8c_8p_{1-3}L8p_{8-3}L8c_8p_{1-3}L8p_{8-3}L8p_{8-3}L8p_{8-3}L8c_8p_{1-3}L8c_8p_{1-3}L8c_8p_{1-3}L8c_8p_{1-3}L8p_{8-3}L8c_8p_{1-3}L8p_{8-}$
${}_3L8c_8p_{1-3}L8c_8p_{1-3}L8c_8p_{1-3}L8c_8p_{1-3}L8c_8p_{1-3}L8c_8p_{1-3}L8p_{8-3}p_{9-7}p_{1-3}L8c_8p_{1-3}L8c_8p_{1-3}L8c_8p_{1-3}L8c_8p_{1-}$
${}_3L8c_8p_{1-3}L8c_8p_{1-3}L8c_8p_{1-3}L8c_8p_{1-3}L8c_8p_{1-3}L8c_8p_{1-3}L8c_8p_{1-3}L8c_8p_{1-3}L8c_8p_{1-3}L8c_8p_{1-3}L8c_8p_{1-3}L8p_{8-}$
${}_3L8c_8p_{1-3}L8p_{8-3}L8p_{8-3}L8c_8p_{1-3}L8p_{8-3}L8p_{8-3}L8c_8p_{1-3}L8p_{8-3}L8p_{8-3}L8c_8p_{1-3}L8c_8p_{1-3}L8c_8p_{1-3}L8p_{8-3}L8c_8p_{1-}$
${}_3L8c_8p_{1-3}L8c_8p_{1-3}L8c_8p_{1-3}L8c_8p_{1-3}L8p_{8-3}L8p_{8-3}L8p_{8-3}L8p_{8-3}L8p_{8-3}L8c_8p_{1-3}L8c_8p_{1-3}L8c_8p_{1-3}L8c_8p_{1-}$
${}_3L8p_{8-3}L8c_8p_{1-3}L8c_8p_{1-3}L8c_8p_{1-3}L8c_8p_{1-3}L8p_{8-3}L8p_{8-3}L8c_8p_{1-3}L8c_8p_{1-3}L8p_{8-3}L8p_{8-3}L8c_8p_{1-3}L8c_8p_{1-}$
${}_3L8p_{8-3}L8c_8p_{1-3}L8p_{8-3}L8p_{8-3}L8c_8p_{1-3}L8c_8p_{1-7}^3p_{1-3}L8p_{8-3}L8p_{8-3}L8p_{8-3}L8p_{8-3}L8c_8p_{1-3}L8c_8p_{1-7}^3p_{1-3}L8p_{8-}$
${}_3L8p_{8-3}L8c_8p_{1-3}L8c_8p_{1-3}L8c_8p_{1-3}L8c_8p_{1-7}p_{1-3}L8c_8p_{1-3}L8c_8p_{1-3}L8c_8p_{1-3}L8p_{8-3}L8c_8p_{1-3}L8c_8p_{1-3}L8p_{8-}$
${}_3L8p_{8-3}L8p_{8-3}L8p_{8-3}L8c_8p_{1-3}L8c_8p_{1-3}L8p_{8-3}L8c_8p_{1-3}L8c_8p_{1-3}L8p_{8-3}L8c_8p_{1-3}L8c_8p_{1-3}L8p_{8-3}L8c_8p_{1-}$
${}_3L8c_8p_{1-3}L8c_8p_{1-3}L8p_{8-3}L8c_8p_{1-3}L8c_8p_{1-3}L8p_{8-3}L8c_8p_{1-3}L8c_8p_{1-3}L8c_8p_{1-3}L8p_{8-3}L8c_8p_{1-3}L8c_8p_{1-3}L8p_{8-}$
${}_3L8p_{8-3}L8c_8p_{1-3}L8c_8p_{1-3}L8c_8p_{1-7}c_1^2p_{1-2}G8p_{1-7}^6c_1^2p_{1-7}^2c_1p_{1-7}^8p_{1-3}p_{5-7}c_1p_{1-7}c_1p_{1-7}^9p_{1-3}p_{5-7}p_{1-7}^6p_{1-3}p_{5-7}p_{1-}$
${}_7^3p_{1-3}p_{5-7}p_{1-7}^2c_1p_{1-7}c_1^2p_{1-7}c_1p_{1-7}c_1^2p_{1-7}c_1p_{1-7}c_1p_{1-7}p_{1-3}p_{5-7}p_{1-7}c_1p_{1-7}c_1p_{1-7}c_1p_{1-7}c_1^2p_{1-7}c_1p_{1-7}c_1p_{1-7}p_{1-3}p_{5-}$
${}_7c_1p_{1-7}c_1p_{1-7}^3p_{1-3}p_{5-7}p_{1-7}^2c_1p_{1-7}c_1p_{1-7}c_1p_{1-7}c_1^2p_{1-7}c_1p_{1-7}c_1p_{1-7}p_{1-3}p_{5-7}c_1p_{1-7}^3p_{1-3}p_{5-7}p_{1-7}c_1p_{1-7}c_1^2p_{1-7}^2c_1p_{1-}$
${}_7^{98}c_1p_{1-7}c_1p_{1-7}^2p_{1-3}L8p_{8-3}L8c_8p_{1-7}c_1p_{1-7}^3p_{1-3}L8p_{8-3}L8c_8p_{1-7}^2c_1p_{1-7}^2c_1p_{1-7}^3p_{1-3}L8p_{8-3}L8c_8p_{1-7}^2c_1p_{1-7}c_1p_{1-}$
${}_7^3p_{1-3}L8c_8p_{1-7}^4p_{1-3}L8c_8p_{1-7}^3p_{1-3}L8c_8p_{1-7}^{25}p_{1-3}p_{5-7}p_{1-3}p_{5-7}p_{1-7}^4p_{1-3}L8c_8p_{1-3}L8p_{8-3}L8p_{8-3}L8c_8p_{1-7}^2p_{1-}$
${}_3L8c_8p_{1-7}^6p_{1-3}L8c_8p_{1-3}L8p_{8-3}L8c_8p_{1-7}^3p_{1-3}L8p_{8-3}L8c_8p_{1-7}^4p_{1-3}L8c_8p_{1-7}p_{1-3}L8c_8p_{1-3}L8p_{8-3}L8p_{8-3}L8p_{8-}$
${}_3L8c_8p_{1-3}L8p_{8-3}L8p_{8-3}L8c_8p_{1-3}L8p_{8-3}L8p_{8-3}L8c_8p_{1-7}p_{1-3}L8p_{8-3}L8p_{8-3}L8p_{8-3}L8p_{8-3}L8c_8p_{1-7}p_{1-3}L8c_8c_1p_{1-}$
${}_3L8c_8c_1p_{1-7}p_{1-3}L8p_{8-3}L8c_8p_{1-3}L8p_{8-3}L8p_{8-3}L8c_8p_{1-7}p_{1-3}L8p_{8-3}L8p_{8-3}L8c_8p_{1-7}p_{1-3}L8c_8c_1p_{1-7}p_{1-3}L8p_{8-}$
${}_3L8p_{8-3}L8c_8p_{1-7}p_{1-3}L8p_{8-3}L8p_{8-3}L8c_8p_{1-7}p_{1-3}L8c_8c_1p_{1-3}L8c_8c_1p_{1-7}c_1p_{1-7}p_{1-3}L8p_{8-3}L8p_{8-3}L8c_8p_{1-7}p_{1-}$
${}_3L8p_{8-3}L8p_{8-3}L8c_8c_1p_{1-7}p_{1-3}L8p_{8-3}L8p_{8-3}L8c_8p_{1-7}p_{1-3}L8p_{8-3}L8c_8p_{1-7}c_1p_{1-7}^2c_1p_{1-3}L8c_8c_1p_{1-3}L8c_8p_{1-7}c_1p_{1-}$
${}_7p_{1-3}L8p_{8-3}L8p_{8-3}L8c_8p_{1-3}L8p_{8-3}L8p_{8-3}L8c_8p_{1-7}p_{1-3}L8p_{8-3}L8c_8p_{1-7}p_{1-3}L8p_{8-3}L8p_{8-3}L8c_8p_{1-7}p_{1-3}L8p_{8-}$
${}_3L8p_{8-3}L8p_{8-3}L8c_8p_{1-7}p_{1-3}L8p_{8-3}L8c_8p_{1-7}p_{1-3}L8p_{8-3}L8c_8p_{1-7}p_{1-3}L8p_{8-3}L8c_8p_{1-7}p_{1-3}L8p_{8-3}L8p_{8-3}L8c_8p_{8-}$

$_{3}L8c_8c_1p_{1-3}L8c_8c_1^{2}p_{1-7}p_{1-3}L8p_{8-3}L8c_8c_1p_{1-3}L8c_8c_1^{2}p_{1-3}L8c_8p_{1-3}L8c_8p_{1-7}p_{1-3}L8c_8p_{1-7}p_{1-3}L8p_{8-3}L8p_{8-}$
$_{3}L8p_{8-3}L8c_8c_1p_{1-3}L8c_8c_1^{2}p_{1-7}^{2}p_{1-3}L8p_{8-3}L8c_8c_1^{2}p_{1-7}^{2}p_{1-3}L8c_8p_{1-7}p_{1-3}L8p_{8-3}L8c_8p_{1-7}p_{1-3}L8c_8p_{1-7}p_{1-}$
$_{3}L8c_8p_{1-7}p_{1-3}L8p_{8-3}L8p_{8-3}L8p_{8-3}L8c_8c_1p_{1-7}c_1p_{1-7}p_{1-3}L8c_8p_{1-3}L8c_8c_1p_{1-3}L8c_8c_1p_{1-7}c_1^{2}p_{1-3}L8c_8p_{1-3}L8c_8p_{1-}$
$_{7}p_{1-3}L8c_8p_{1-7}p_{1-3}L8p_{8-3}L8p_{8-3}L8p_{8-3}L8c_8c_1p_{1-3}L8c_8c_1^{2}p_{1-7}^{2}p_{1-3}L8p_{8-3}L8c_8c_1^{2}p_{1-7}^{2}p_{1-3}L8c_8p_{1-7}p_{1-3}L8p_{8-}$
$_{3}L8c_8p_{1-7}p_{1-3}L8c_8p_{1-7}p_{1-3}L8c_8p_{1-7}p_{1-3}L8p_{8-3}L8p_{8-3}L8p_{8-3}L8c_8c_1p_{1-7}c_1p_{1-7}p_{1-3}L8c_8p_{1-7}p_{1-3}L8c_8p_{1-7}p_{1-}$
$_{3}L8c_8p_{1-7}p_{1-3}L8c_8p_{1-7}p_{1-3}L8p_{8-3}L8p_{8-3}L8p_{8-3}L8c_8c_1^{2}p_{1-7}^{2}p_{1-3}L8c_8p_{1-7}p_{1-3}L8c_8p_{1-7}p_{1-3}L8c_8p_{1-7}p_{1-3}L8p_{8-}$
$_{3}L8p_{8-3}L8p_{8-3}L8c_8c_1p_{1-7}c_1p_{1-7}p_{1-3}L8c_8p_{1-7}p_{1-3}L8c_8p_{1-7}p_{1-3}L8c_8p_{1-7}p_{1-3}L8c_8p_{1-7}p_{1-3}L8p_{8-3}L8p_{8-3}L8p_{8-}$
$_{3}L8c_8c_1^{2}p_{1-7}^{2}p_{1-3}L8c_8p_{1-7}p_{1-3}L8c_8p_{1-7}p_{1-3}L8c_8p_{1-7}p_{1-3}L8p_{8-3}L8c_8p_{1-7}p_{1-3}L8c_8p_{1-7}p_{1-3}L8c_8p_{1-7}p_{1-3}L8p_{8-}$
$_{3}L8p_{8-3}L8p_{8-3}L8c_8c_1p_{1-3}L8c_8c_1p_{1-7}c_1p_{1-7}c_1p_{1-7}c_1p_{1-7}c_1p_{1-3}L8c_8c_1p_{1-3}L8c_8p_{8-3}L8c_8c_1p_{1-3}L8c_8c_1p_{1-3}L8p_{8-}$
$_{3}L8p_{8-3}L8p_{8-3}L8p_{8-3}L8c_8c_1p_{1-7}p_{1-3}L8c_8p_{1-3}L8p_{8-3}L8c_8p_{1-3}L8p_{8-5}p_{1-3}L8c_8c_1p_{1-7}^{2}p_{1-3}L8p_{8-3}L8p_{8-3}L8p_{8-}$
$_{3}L8p_{8-3}L8p_{8-3}L8c_8p_{8-3}L8p_{8-3}L8p_{8-3}L8p_{8-3}L8p_{8-3}L8p_{8-3}L8c_8p_{8-3}L8p_{8-3}L8p_{8-3}L8c_8p_{8-3}L8p_{8-3}L8p_{8-3}L8p_{8-}$
$_{3}L8c_8p_{8-3}L8p_{8-3}L8p_{8-3}L8c_8^{4}p_{8-3}L8c_8^{4}p_{1-3}L8c_8^{9}p_{8-1}c_3^{18}p_{3-4}c_9^{48}B8$

### cen9

$A4/9p_{1-2}p_{4-1}^{2}p_{4-5}p_{7-2}p_{4-6}c_6p_{7-1}p_{4-5}G9{+}M9L9{+}B9{+}Y9p_{4-2}^{2}E9p_{7-3}E9p_{7-1}p_{7-2}E9p_{7-4}p_{7-2}^{2}c_4p_{4-5}p_{7-2}F9p_{7-5}p_{7-}$
$_{1}c_3p_{6-1}F9c_7p_{7-2}F9c_7p_{7-2}F9p_{7-1}p_{1-2}F9p_{7-2}p_{4-2}F9p_{7-2}p_{7-1}p_{7-2}F9p_{7-5}p_{7-2}p_{7-3}E9p_{7-4}c_6p_{6-2}p_{4-5}p_{7-2}F9p_{7-}$
$_{3}E9c_7p_{7-4}c_6p_{6-2}F9c_7p_{7-4}p_{7-2}c_7p_{7-5}p_{7-2}F9p_{7-2}E9p_{7-2}p_{7-3}p_{7-2}p_{7-5}F9p_{7-5}p_{7-2}^{2}c_4^{3}D9p_{7-5}p_{7-1}G9{+}M9p_{5-2}p_{7-4}p_{6-}$
$_{2}F9p_{7-5}p_{7-1}c_7p_{7-5}p_{7-2}F9p_{7-2}E9p_{7-2}p_{7-3}p_{7-2}p_{7-5}F9p_{7-5}p_{7-2}^{2}c_4^{3}D9p_{7-5}p_{7-1}G9{+}M9p_{5-2}p_{7-4}p_{6-2}F9p_{7-5}p_{7-1}p_{7-3}p_{1-}$
$_{4}c_6p_{7-2}F9p_{7-3}G9{+}M9L9{+}B9{+}Y9p_{7-1}c_3p_{7-1}F9p_{7-4}c_6p_{6-2}p_{7-3}p_{5-3}p_{7-5}p_{5-3}c_7p_{7-3}c_7p_{7-2}F9p_{7-3}p_{7-5}p_{4-2}F9p_{7-}$
$_{3}c_7p_{7-2}F9p_{7-3}p_{7-5}p_{4-2}F9p_{7-3}p_{7-5}p_{4-2}F9c_7p_{7-2}F9p_{7-3}p_{7-5}p_{4-6}p_{3-5}p_{7-2}L9{+}B9{+}Y9F9p_{7-5}c_7p_{7-2}F9p_{7-3}c_7p_{7-}$
$_{2}F9p_{7-3}c_7p_{7-2}F9p_{7-1}p_{4-2}F9p_{7-2}p_{4-2}F9p_{7-3}c_7p_{7-2}F9p_{7-2}F9p_{7-3}^{2}c_7p_{7-2}F9p_{7-3}c_7p_{7-2}F9p_{7-3}c_7p_{7-2}F9p_{7-3}c_7p_{7-}$
$_{2}F9p_{7-2}F9p_{7-3}c_7p_{7-1}F9p_{7-2}F9p_{7-2}F9p_{7-2}F9p_{7-3}c_7p_{7-5}F9p_{7-3}c_7p_{7-2}F9p_{7-3}c_7p_{7-2}F9p_{7-3}c_7p_{7-2}F9p_{7-2}F9c_7p_{7-}$
$_{2}F9p_{7-3}c_7p_{7-2}F9p_{7-3}c_7p_{7-2}F9p_{7-3}c_7p_{7-1}F9p_{7-2}F9p_{7-3}c_7p_{7-2}F9C9c_7p_{7-2}F9p_{7-3}c_7p_{7-2}F9c_7p_{7-2}F9p_{7-3}c_7p_{7-}$
$_{2}F9p_{7-3}c_7p_{7-2}F9p_{7-2}F9p_{7-3}c_7p_{7-2}F9p_{7-3}c_7p_{7-2}F9p_{7-3}c_7p_{7-2}F9p_{7-3}c_7p_{7-2}F9p_{7-3}c_7p_{7-2}F9c_7p_{7-2}F9p_{7-3}c_7p_{7-}$
$_{2}F9p_{7-2}F9p_{7-3}c_7p_{7-2}F9c_7p_{7-3}c_7p_{7-2}F9c_7p_{7-2}F9p_{7-3}p_{5-6}p_{1-2}F9c_7p_{7-2}F9p_{7-3}c_7p_{7-2}F9c_7p_{7-2}F9p_{7-3}p_{5-2}F9c_7p_{7-}$
$_{2}F9p_{7-3}c_7p_{7-2}F9p_{7-3}c_7p_{7-2}F9c_7p_{7-3}p_{7-2}F9p_{7-3}p_{5-2}F9c_7p_{7-2}F9p_{7-3}c_7p_{7-2}F9c_7p_{7-2}F9p_{7-3}p_{5-2}F9c_7p_{7-2}F9p_{7-}$
$_{3}c_7p_{7-2}F9p_{7-3}c_7p_{7-2}F9c_7p_{7-2}F9p_{7-3}c_7p_{7-2}F9c_7p_{7-2}F9p_{7-3}^{2}c_7p_{7-2}F9p_{7-3}p_{5-2}F9c_7p_{7-2}F9p_{7-3}p_{5-2}F9c_7p_{7-2}F9p_{7-}$
$_{2}F9p_{7-3}c_7p_{7-2}F9c_7p_{7-2}F9p_{7-3}c_7p_{7-2}F9p_{7-3}c_7p_{7-2}F9p_{7-3}c_7p_{7-2}F9p_{7-3}c_7p_{7-2}F9c_7p_{7-1}F9p_{7-2}F9p_{7-3}c_7p_{7-2}F9p_{7-}$
$_{3}c_7p_{7-2}F9c_7p_{7-2}F9p_{7-3}c_7p_{7-2}F9c_7p_{7-2}F9p_{7-3}^{2}c_7p_{7-2}F9p_{7-3}p_{5-2}F9c_7p_{7-2}F9p_{7-3}p_{5-2}F9c_7p_{7-2}F9p_{7-3}p_{5-2}F9c_7p_{7-}$
$_{2}F9p_{7-2}F9p_{7-3}c_7p_{7-2}F9c_7p_{7-2}F9p_{7-3}c_7p_{7-2}F9p_{7-3}c_7p_{7-2}F9p_{7-3}c_7p_{7-2}F9p_{7-3}c_7p_{7-2}F9c_7p_{7-1}F9p_{7-2}F9p_{7-3}c_7p_{7-}$
$_{2}F9p_{7-2}F9p_{7-3}c_7p_{7-2}F9p_{7-3}c_7p_{7-2}F9p_{7-3}c_7p_{7-2}F9p_{7-3}c_7p_{7-2}F9p_{7-3}c_7p_{7-2}F9p_{7-3}c_7p_{7-2}F9p_{7-3}c_7p_{7-2}F9p_{7-3}c_7p_{7-}$
$_{1}F9p_{7-3}p_{7-2}F9p_{7-3}c_7p_{7-2}F9p_{7-3}p_{7-2}F9p_{7-3}c_7p_{7-2}F9p_{7-3}c_7p_{7-2}F9p_{7-3}p_{7-2}F9p_{7-3}c_7^{2}p_{7-2}F9p_{7-3}c_7p_{7-2}F9p_{7-}$
$_{3}c_7p_{7-2}F9p_{7-2}F9p_{7-3}p_{7-2}F9c_7p_{7-2}F9p_{7-2}F9p_{7-3}p_{7-2}F9c_7p_{7-2}F9p_{7-3}p_{7-2}F9p_{7-3}c_7p_{7-2}F9c_7p_{7-2}F9p_{7-3}p_{7-}$
$_{2}F9c_7p_{7-2}F9p_{7-2}F9p_{7-3}p_{7-2}F9c_7p_{7-2}F9p_{7-3}p_{7-2}F9p_{7-3}c_7p_{7-2}F9c_7p_{7-2}F9p_{7-3}c_7p_{7-2}F9c_7p_{7-2}p_{6-3}c_7p_{7-2}F9c_7p_{7-}$
$_{2}p_{6-3}c_7p_{7-3}p_{7-1}F9p_{7-3}c_7p_{7-2}F9p_{7-2}F9p_{7-3}c_7p_{7-2}F9c_7p_{7-2}p_{6-2}F9c_7p_{7-2}p_{6-2}F9c_7p_{7-2}F9p_{7-2}F9p_{7-3}p_{7-2}F9c_7p_{7-}$
$_{2}F9p_{7-2}F9p_{7-3}p_{7-2}F9c_7p_{7-2}F9p_{7-3}p_{7-2}F9c_7p_{7-2}F9p_{7-3}p_{7-2}F9p_{7-3}c_7p_{7-2}F9c_7p_{7-2}F9p_{7-3}p_{7-2}F9c_7p_{7-2}F9p_{7-3}p_{7-}$
$_{2}F9p_{7-3}c_7p_{7-2}F9c_7p_{7-2}F9p_{7-3}c_7p_{7-2}F9c_7p_{7-2}F9p_{7-3}c_7p_{7-2}F9c_7p_{7-2}p_{6-3}c_7p_{7-2}F9c_7p_{7-2}p_{6-3}c_7p_{7-3}p_{7-1}F9p_{7-}$
$_{3}c_7p_{7-2}F9p_{7-2}F9p_{7-3}c_7p_{7-2}F9c_7p_{7-2}p_{6-2}F9c_7p_{7-2}p_{6-2}F9c_7p_{7-2}F9p_{7-3}c_7p_{7-2}F9p_{7-5}F9p_{7-2}F9c_7p_{7-3}c_7p_{7-2}F9p_{7-}$
$_{2}F9p_{7-3}p_{7-2}F9p_{7-3}c_7p_{7-2}F9p_{7-3}c_7p_{7-2}F9p_{7-3}c_7p_{7-2}F9c_7p_{7-2}F9p_{7-2}F9p_{7-3}G9{+}M9p_{5-1}F9p_{7-3}p_{7-2}F9p_{7-3}c_7p_{7-}$
$_{3}c_7p_{7-2}F9p_{7-3}c_7p_{7-2}F9p_{7-3}p_{7-2}F9p_{7-3}c_7p_{7-3}c_7p_{7-2}F9p_{7-3}p_{7-2}F9p_{7-3}c_7p_{7-3}c_7p_{7-2}F9p_{7-3}c_7p_{7-2}F9p_{7-3}p_{7-1}F9p_{7-}$
$_{3}c_7p_{7-3}c_7p_{7-2}F9p_{7-2}F9p_{7-3}p_{7-1}F9p_{7-3}c_7p_{7-3}c_7p_{7-2}F9p_{7-2}F9p_{7-3}c_7p_{7-2}F9p_{7-2}F9p_{7-3}p_{7-2}F9p_{7-3}^{3}c_7p_{7-2}F9p_{7-}$
$_{2}F9p_{7-3}p_{7-2}F9p_{7-3}c_7p_{7-1}F9p_{7-3}p_{7-2}F9p_{7-3}c_7p_{7-2}F9c_7p_{7-2}F9p_{7-2}F9p_{7-3}^{3}c_7p_{7-2}F9p_{7-2}F9p_{7-3}p_{7-2}F9p_{7-3}c_7p_{7-}$
$_{2}F9p_{7-3}p_{7-1}F9p_{7-3}c_7p_{7-2}F9c_7p_{7-2}F9p_{7-2}F9p_{7-3}c_7p_{7-2}F9p_{7-3}p_{7-1}F9p_{7-3}c_7p_{7-2}F9c_7p_{7-2}F9p_{7-2}F9p_{7-3}c_7p_{7-}$
$_{2}F9p_{7-3}c_7p_{7-2}F9c_7p_{7-2}F9p_{7-3}c_7p_{7-2}F9p_{7-2}F9c_7p_{7-2}F9p_{7-3}c_7p_{7-2}F9c_7p_{7-2}F9p_{7-3}c_7p_{7-2}F9c_7p_{7-2}F9p_{7-3}c_7p_{7-}$
$_{2}F9c_7p_{7-2}F9p_{7-2}F9c_7p_{7-2}F9p_{7-3}p_{5-2}F9c_7p_{7-2}F9p_{7-3}c_7p_{7-2}F9c_7p_{7-2}p_{7-3}p_{5-2}F9c_7p_{7-2}F9p_{7-2}F9p_{7-3}c_7p_{7-2}F9p_{7-}$
$_{2}F9c_7p_{7-2}F9p_{7-3}c_7p_{7-2}F9c_7p_{7-2}F9p_{7-3}c_7p_{7-2}F9c_7p_{7-2}F9c_7p_{7-2}p_{7-3}p_{5-2}F9c_7p_{7-2}F9p_{7-2}F9p_{7-3}c_7p_{7-2}F9p_{7-}$
$_{2}F9c_7p_{7-2}F9p_{7-3}c_7^{2}p_{7-2}F9p_{7-3}c_7p_{7-2}F9c_7p_{7-2}F9p_{7-3}c_7p_{7-2}F9c_7p_{7-2}F9p_{7-2}F9c_7p_{7-2}F9p_{7-3}p_{5-2}F9c_7p_{7-2}F9p_{7-}$
$_{3}c_7p_{7-2}F9c_7p_{7-2}p_{7-3}p_{5-2}F9c_7p_{7-2}F9p_{7-3}p_{5-2}F9c_7p_{7-2}F9p_{7-3}p_{5-2}F9c_7p_{7-2}F9p_{7-2}F9p_{7-3}c_7p_{7-2}F9p_{7-3}c_7p_{7-}$
$_{2}F9p_{7-3}c_7p_{7-2}F9c_7p_{7-2}F9p_{7-3}c_7p_{7-2}F9p_{7-3}p_{5-2}F9c_7p_{7-2}F9p_{7-3}p_{5-2}F9c_7p_{7-2}F9p_{7-2}F9p_{7-3}c_7p_{7-2}F9p_{7-3}p_{5-}$
$_{2}F9p_{7-3}p_{5-2}F9c_7p_{7-2}F9p_{7-2}F9p_{7-3}c_7p_{7-2}F9p_{7-3}c_7p_{7-2}F9p_{7-3}c_7p_{7-2}F9c_7p_{7-2}F9p_{7-2}F9p_{7-3}c_7p_{7-2}F9p_{7-3}c_7p_{7-}$

2F9p7-3c7p7-2F9p7-3c7p7-2F9p7-3c7p7-2F9c7p7-2F9p7-3c7p7-2F9p7-3^2c7p7-2F9p7-3c7p7-2F9p7-3p7-1F9p7-
3c7p7-3p7-2F9p7-3p7-2F9p7-3p7-2F9p7-3c7p7-2F9p7-3p7-2F9p7-3c7p7-3p7-2F9p7-3p7-2F9p7-3p7-2F9p7-3c7p7-
2F9p7-3p7-2F9p7-3c7p7-2F9c7p7-2F9p7-3c7p7-2F9p7-2F9c7p7-2F9p7-3c7p7-2F9p7-2F9p7-3c7p7-2F9c7p7-2F9p7-
3c7p7-2F9p7-2F9p7-3c7p7-2F9c7p7-2F9p7-3p5-2F9c7p7-2F9p7-3p5-2F9c7p7-2F9p7-3p5-2F9c7p7-2F9p7-3p5-
2F9c7p7-2F9p7-2F9p7-3c7p7-2F9p7-3c7p7-2F9p7-3c7p7-2F9p7-3p7-2F9c7p7-2F9p7-3c7p7-2F9p7-3p5-2F9c7p7-
2F9p7-3p5-2F9c7p7-2F9p7-2F9p7-3c7p7-2F9p7-3c7p7-2F9p7-3c7p7-2F9c7p7-2F9p7-3c7p7-2F9p7-3c7p7-2F9p7-
3c7p7-2F9p7-3c7p7-2F9c7p7-2F9p7-3c7p7-2F9p7-3c7p7-2F9p7-3c7p7-2F9p7-3c7p7-2F9p7-3c7p7-2F9p7-3c7p7-
2F9p7-2F9p7-3c7p7-2F9p7-3c7p7-2F9p7-3c7p7-2F9c7p7-2F9p7-3c7p7-2F9p7-3c7p7-2F9p7-3c7p7-2F9p7-3c7p7-
2F9p7-3c7p7-2F9p7-2F9p7-3c7p7-2F9p7-3c7p7-2F9p7-3c7p7-2F9c7p7-2F9p7-3c7p7-2F9p7-3c7p7-2F9p7-3c7p7-
2F9p7-3c7p7-2F9p7-2F9p7-3c7^2p7-2F9p7-3c7^2p7-2F9p7-3c7p7-2F9p7-3c7p7-2F9p7-3c7p7-2F9p7-2F9p7-3p7-
2F9p7-3c7p7-2F9p7-3p7-2F9p7-3c7p7-2F9c7p7-1F9p7-2F9p7-3c7p7-2F9p7-3c7p7-2F9p7-3p7-2F9p7-3c7^2p7-2F9p7-
3c7p7-2F9p7-3c7p7-2F9p7-2F9p7-3p7-2F9p7-3c7p7-2F9p7-3c7p7-2F9c7^2p7-2F9p7-2F9p7-3p7-2F9p7-3c7p7-2F9p7-
3c7p7-2F9c7p7-2F9p7-3c7p7-2F9c7p7-2F9p7-2F9p7-3c7p7-2F9c7p7-2p6-3c7p7-2F9p7-3p7-2F9p7-2F9p7-3c7p7-
2F9c7p7-2p6-3c7p7-2F9p7-3p7-2F9p7-3c7p7-2F9c7p7-2F9p7-3c7p7-2F9c7p7-2p6-2p7-3c7p7-2F9c7p7-2F9c7p7-
2F9c7p7-2F9p7-2F9c7p7-2F9p7-3p7-2F9c7p7-1F9G9+M9L9+B9+Y9F9p7-3c7p7-2F9c7p7-2F9p7-3c7p7-2F9p7-
2F9c7p7-2F9p7-3c7p7-3c7p7-2F9c7p7-2F9c7p7-2F9p7-2F9c7p7-2F9p7-3p7-2F9c7p7-
1F9G9+M9L9+B9+Y9F9p7-3c7p7-2F9c7p7-2F9p7-3c7p7-2F9p7-2F9c7p7-2F9p7-3c7p7-2F9p7-3^2c7p7-2F9p7-
3c7p7-2F9p7-2F9c7p7-2F9p7-3c7p7-2F9p7-2F9p7-3p7-2F9p7-3c7p7-2F9p7-3c7p7-2F9p7-2F9p7-3p7-2F9p7-3c7p7-
2F9p7-3c7p7-2F9p7-3c7p7-2F9c7p7-1F9p7-3c7p7-2F9p7-3c7p7-2F9p7-3c7p7-2F9p7-2F9p7-3c7p7-2F9p7-3c7p7-
2F9p7-3c7p7-2F9p7-3c7p7-2F9p7-2F9p7-2C9c7p7-2F9p7-3c7p7-2F9p7-2F9p7-2c3F9p7-3c7p7-2F9p7-3c7p7-2F9p7-
3c7p7-2F9p7-3c7p7-2F9p7-3c7p7-2F9p7-3c7p7-2F9p7-3c7p7-2F9p7-3c7p7-3c7p7-2F9p7-3c7p7-2F9p7-3c7p7-2F9p7-
3c7p7-2F9p7-3c7p7-2F9p7-3c7p7-2F9p7-3c7p7-2F9p7-3c7p7-2F9p7-3c7p7-2F9p7-3c7p7-3c7p7-2F9p7-3c7p7-2F9p7-
3c7p7-2F9p7-3c7p7-2F9p7-3c7p7-2F9p7-3c7p7-2F9p7-3c7p7-2F9p7-3c7p7-2F9p7-3c7p7-2F9p7-3c7p7-2F9p7-
3^2c7^2p7-2F9c7p7-2F9p7-3c7p7-2F9p7-3c7p7-2F9p7-3c7p7-2F9p7-3^2c7p7-2F9p7-3c7p7-2F9p7-3c7p7-2F9p7-3^2c7p7-
2F9p7-3c7p7-2F9p7-3c7p7-2F9p7-3^2c7p7-2F9p7-3c7p7-3c7p7-2F9p7-3c7p7-2F9p7-3c7p7-2F9p7-3c7p7-2F9p7-3c7p7-
3c7p7-2F9p7-3c7p7-3c7p7-2F9p7-3c7p7-2F9p7-3c7p7-3c7p7-3c7p7-2F9p7-3c7p7-3c7p7-3c7p7-2F9p7-3c7p7-2F9p7-
3^2c7p7-2F9p7-3c7p7-2F9p7-3c7p7-2F9p7-3c7p7-2F9p7-3c7p7-3c7p7-2F9p7-3c7p7-3c7p7-2F9p7-3c7p7-2F9p7-3c7p7-
2F9p7-3c7p7-3c7p7-3c7p7-2F9p7-3^2c7p7-2F9p7-3^2c7p7-2F9p7-3c7^2p7-3^2c7p7-2F9p7-3^2c7p7-2F9p7-3c7p7-2F9p7-
3c7p7-3^2c7p7-2F9p7-3c7p7-3^2c7p7-2F9p7-3c7p7-2F9p7-3c7p7-2F9p7-3c7p7-2F9p7-3c7p7-2F9p7-3c7p7-2F9p7-3c7p7-
2F9p7-3c7p7-3c7p7-2F9p7-3c7p7-2F9p7-3c7p7-2F9p7-3c7p7-2F9p7-3c7p7-2F9p7-3c7p7-2F9c7p7-2F9p7-2F9p7-
3c7p7-2F9p7-2p4-2F9p7-2c4p4-2F9p7-2p5-2F9p7-2c4p4-2F9c7p7-2F9p7-3c7p7-2F9p7-3c7p7-2F9p7-2F9c7p7-
2F9p7-3c7p7-2F9c7p7-2F9p7-2F9c7p7-2F9p7-3c7p7-2F9p7-2p4-2F9p7-3p5-2F9p7-3c7p7-2F9c7^2p7-2F9p7-2F9p7-
3c7p7-3c7p7-1c3p1-2F9p7-4c1p1-2F9p7-3c7p7-2F9p7-2F9c7^2p7-2F9c7p7-2F9p7-2p4-2F9p7-3^3c7p7-2F9p7-3p5-
2F9p7-3c7p7-2F9c7^2p7-2F9p7-4L9+B9+Y9F9p7-4L9+B9+Y9F9c7p7-2F9p7-2F9p7-4p1-4p1-2F9c7p7-2F9p7-
2F9p7-3c7p7-2F9p7-3c7p7-2F9p7-4c1p1-2F9p7-4c1p1-2F9p7-1F9c7p7-2F9p7-4c1p1-2F9p7-1F9c7p7-2F9p7-4c1p1-
2F9p7-4c1p1-2F9p7-3c7p7-2F9p7-3c7p7-2F9c7p7-1F9p7-2F9p7-3p7-4c1p1-2F9c7p7-1F9p7-2F9c7p7-1F9p7-2F9p7-
3c7p7-2F9p7-2F9p7-2F9p7-3c7p7-2F9p7-4c1p1-2F9p7-4p1-2p5-1F9p7-2F9c7p7-2F9p7-1F9c7p7-2F9p7-4c1p1-
2F9p7-1F9c7p7-2F9p7-4c1p1-2F9p7-4c1p1-2F9p7-3c7p7-2F9p7-3c7p7-2F9c7p7-1F9p7-2F9p7-3p7-4c1p1-2F9c7p7-
1F9p7-2F9c7p7-1F9p7-2F9p7-3c7p7-2F9p7-2F9p7-2F9p7-3c7p7-2F9p7-3c7p7-2F9p7-4p1-2p5-1F9p7-2F9c7p7-
1F9p7-2F9p7-3c7p7-2F9p7-2F9p7-2F9p7-4c1p1-2F9p7-4c1p1-2F9p7-4c1p1-2F9p7-4c1p1-2F9p7-4c1p1-2F9p7-
3c7p7-2F9p7-2F9c7p7-1F9p7-3c7p7-1F9p7-2F9p7-3c7p7-2F9p7-3c7p7-2F9p7-2p4-2F9p7-2F9c7p7-2F9p7-3c7p7-
2F9p7-2F9c7p7-1F9p7-3c7p7-2F9p7-2F9p7-3c7p7-2F9p7-3c7p7-2F9p7-2p4-2F9p7-2F9p7-3c7p7-2F9p7-3c7p7-
2F9p7-2p4-2F9p7-2F9c7p7-2F9p7-2F9p7-3c7p7-2F9p7-2F9p7-3c7p7-2F9c7p7-1F9p7-2F9p7-3c7p7-1F9p7-3c7p7-
2F9p7-5F9p7-2F9p7-3c7p7-2F9p7-2F9p7-5F9p7-2F9c7p7-2F9p7-3c7p7-2F9p7-2F9c7p7-1F9p7-3c7p7-2F9p7-
2F9p7-3c7p7-2F9p7-3c7p7-2F9p7-2p4-2F9p7-2F9p7-3c7p7-2F9p7-3c7p7-2F9p7-2p4-2F9p7-2F9c7p7-2F9p7-2F9p7-
3c7p7-2F9p7-2F9p7-3c7p7-2F9p7-3c7p7-2F9p7-2F9p7-3c7p7-2F9c7p7-1F9p7-2F9p7-3c7p7-1F9p7-3c7p7-2F9p7-
5F9p7-2F9p7-3c7p7-2F9p7-2F9p7-5F9p7-2F9p7-3c7p7-2F9p7-2F9p7-5F9p7-2F9p7-3c7p7-2F9p7-2F9p7-5F9p7-
2F9p7-3c7p7-2F9p7-2F9c7p7-2F9p7-2F9p7-3p7-1F9c7p7-3c7^2p7-2p5-2F9c7p7-2F9c7p7-2F9p7-3p7-1c3^2F9p7-3p7-
1c3p3-5p7-3^2p7-2F9p7-3c7p7-2F9c3^2F9c3F9c3C9c7p7-2F9c3F9c3F9p7-3c7p7-2F9c3F9c3C9c7p7-
2F9c3F9c3F9p7-3c7p7-2F9c3F9p7-3c7p7-2F9c3F9p7-3c7p7-2F9c3F9p7-3c7p7-2F9p7-2E9c7p7-2F9p7-2C9c7p7-
2F9p7-2F9c7p7-2F9p7-2F9p7-3p7-2F9p7-2F9p7-3c7p7-2F9p7-2F9c7p7-2F9p7-2F9p7-3p7-2F9p7-2F9p7-3c7p7-

2F9p7-3c7p7-2F9p7-2F9p7-3p7-2F9p7-2F9p7-3c7p7-2F9p7-2F9p7-3c7p7-2F9p7-1F9p7-3p7-2F9p7-2F9p7-3c7p7-
2F9p7-2F9p7-3c7p7-2F9p7-3p7-2F9p7-3c7p7-2F9p7-3c7p7-2F9p7-3c7p7-2F9p7-2F9p7-2p6-2F9p7-2F9p7-3c7p7-
2F9p7-2C9c7p7-2F9p7-3c7p7-2F9p7-2C9c7p7-2F9p7-3p7-2F9p7-3c7p7-2F9p7-2F9p7-2C9p7-2F9p7-3c7p7-2F9p7-
3p7-2F9p7-3c7p7-2F9p7-3c7p7-2F9c7p7-2F9p7-2C9p7-2F9c7p7-2F9p7-2F9c7p7-2F9c7p7-2F9p7-2F9p7-2C9p7-
2F9p7-2C9c7p7-2F9p7-2F9p7-2C9c7p7-2F9p7-3c7p7-2F9p7-3c7p7-2F9p7-3c7p7-2F9p7-3c7p7-2F9p7-2F9p7-
2C9p7-2F9p7-3c7p7-2F9p7-1F9p7-3c7p7-2F9p7-3p7-2C9c7p7-2F9p7-2C9p7-4p1-2F9p7-3p5-2F9p7-3c7p7-2F9p7-
3c7p7-2F9c3F9p7-3c7p7-2F9p7-3c7p7-2F9G9+M9p5-2F9p7-3p7-2F9p7-1F9p7-3c7p7-2F9p7-
2F9G9+M9c3F9c7p7-3p7-2F9p7-2F9p7-2F9p7-2F9p7-3c7p7-1F9p7-1C9c7^2p7-2F9p7-2E9p7-3c7p7-2F9p7-3c7p7-
2F9p7-2E9p7-3c7p7-2F9p7-2c4c7p7-2F9c7p7-2F9p7-4L9+B9+Y9F9p7-2E9F9p7-3c7p7-2F9p7-3c7p7-2F9c7p7-
2F9p7-2F9c7p7-2F9c7p7-2F9p7-2F9p7-3^2c7p7-2F9p7-3c7p7-2F9c7p7-2F9D9p1-2F9c7p7-2F9D9p1-2F9c7p7-
2F9p7-3c7p7-2F9c7p7-2F9D9p1-2F9c7p7-2p6-2F9c7p7-2F9D9p1-2F9c7p7-2p6-2F9c7p7-2F9D9p1-2F9c7p7-
2F9D9p1-2F9c7p7-2F9D9p1-2F9c7p7-2F9p7-3c7p7-2F9c7p7-2F9D9p1-2F9c7p7-2p6-2F9c7p7-2F9D9p1-
2F9c7p7-2p6-2F9c7p7-2F9D9p1-2F9c7p7-3c7p7-2F9c3F9p7-3c7p7-2F9p7-3c7p7-1c3C9p7-1c3F9p7-3p7-
1c3F9c7p7-3p7-2F9p7-3p7-2F9p7-3c7p7-2F9p7-3c7p7-2F9p7-3p7-1c3F9p7-3p7-5p7-2F9c7c3F9p7-3p5-2F9p7-5c7p7-
2p5-2F9c7p7-2F9p7-5p7-1C9p7-2F9c7^2p7-2F9p7-3c7D9^2F9p7-3p5-2F9p7-5p7-2p7-4p6-2F9c7p7-2F9p7-3p5-2F9p7-
5c7p7-2F9p7-4L9+B9+Y9F9p7-3p5-2F9p7-2F9p7-3c7p7-2F9p7-3c7p7-2F9c7p7-2F9G9+M9C9c7p7-2F9p7-
2F9c7p7-2F9p7-2F9p7-3c7p7-1F9p7-2F9p7-3c7p7-2F9p7-3p7-4p1-2F9p7-3p5-2F9p7-3c7p7-2F9p7-3c7p7-3^2p7-
1F9c7p7-2F9p7-3c7p7-2F9p7-3c7p7-2F9c7p7-1c3F9c7p7-1c3F9p7-4p1-2F9c7p7-1c3F9p7-4p1-2F9c7p7-1c3F9p7-
4p1-2F9p7-5L9+B9+Y9F9p7-3c7p7-1F9p7-3c7p7-1F9c7p7-2F9p7-3c7p7-1F9c7p7-2F9p7-3c7p7-2F9p7-3c7p7-
2F9c3F9G9+M9p2-3c7p7-1F9p7-3c7p7-1F9c7p7-2F9p7-3c7p7-1F9c7p7-2F9p7-3c7p7-2F9p7-3c7p7-
2F9c3F9G9+M9p2-3c7p7-2F9c3F9G9+M9p2-3c7p7-2F9p7-3c7p7-1c3C9p7-1p3-1c3C9p7-1c3F9p7-2F9p7-
2F9p7-3c7p7-2F9p7-3c7p7-2F9p7-3c7p7-2F9p7-2F9p7-3c7p7-2F9p7-3p7-2F9p7-3c7p7-2F9p7-3p7-1F9p7-3c7p7-
3^3p7-2F9p7-3^2c7p7-2F9p7-3p7-1c3F9p7-3c7p7-2F9p7-1C9p7-3^2c7p7-2F9p7-3p7-1c3F9p7-3c7p7-2F9p7-2F9p7-
1c3F9p7-3p7-2p4-6c3F9p7-4p7-1p7-2F9c7p7-2F9p7-1c3F9p7-5c7D9p4-5p7-1c3c7D9p4-5c7p7-2c7p4-5c7p7-2F9p7-
5p7-2c3F9p7-3E9L9+B9+Y9F9p7-2F9p7-2p2-3c7p7-1p3-1F9c7p7-2F9p7-2^2p5-3p7-1p3-6D9F9p7-1c3C9p7-
3D9p1-3p7-2c5p5-2F9p7-5p7-2p7-1p3-5p7-5p7-3E9p7-3c7p7-2p7-5c7p7-5p7-2E9p7-3c7p7-5^2p7-3^2c7p7-5p7-2E9p7-3p7-
2F9p7-2p5-2F9c7p7-1F9p7-2p5-2F9p7-4p6-2F9c7p7-2F9c7p7-2p6-2F9c7p7-2F9c7p7-2F9c7p7-4c6p6-2F9p7-4p6-
2F9c7^2p7-2F9p7-4p6-2F9c7p7-2F9p7-4p6-2F9c7p7-2F9p7-3c7p7-2F9p7-5p7-2^2c5E9p7-2F9p7-5p7-3E9p7-1c7D9p4-
2F9p7-2F9p7-5p7-3p5-2F9c7p7-3p7-5^2p7-1p7-5c7p3-5p7-1p3-5p7-3p7-2p5-2F9p7-5p7-3E9p7-2F9p7-5^2c7p7-5p7-3E9p7-
3c7p7-4p7-2E9p7-3c7p7-5c7p7-5p7-3E9p7-3c7p7-5p7-3p5-3c7p7-5p7-2F9c7p7-2F9c7p7-2F9p7-4c7p7-2F9c7p7-
2F9c7p7-1F9p7-4c7p7-2F9c7p7-2F9c7^2p7-2F9c7^2p7-2F9c7p7-2F9p7-2c1p1-2F9c7p7-2F9p7-5p7-2F9c7p7-
2F9c7p7-1F9p7-5p7-2F9c7p7-1F9c7p7-2F9c7p7-2F9c7p7-2F9c7p7-2F9c7p7-2F9c7p7-2F9c7p7-2F9c7p7-
2F9c7p7-2F9G9+M9p5-2F9c7p7-1F9p7-2F9c7p7-2F9p7-2F9c7p7-2F9c7p7-2F9p7-2F9c7p7-2F9G9+M9p5-
2F9c7p7-5p7-2F9c7p7-2F9c7p7-1F9c7p7-2F9G9+M9p5-2F9c7p7-2F9G9+M9p5-2F9G9+M9p5-6p5-
1F9G9+M9p5-7p6-1F9G9+M9p5-2F9c7p7-1F9G9+M9p5-2F9c7p7-2F9G9+M9E9c3F9c7p7-2F9c7p7-
2F9c7p7-2F9c7p7-2F9c7p7-2F9c7p7-2F9p7-4p6-2F9c7p7-2F9c7p7-2F9c7p7-2F9c7p7-2F9c7p7-2F9c7p7-
2F9c7p7-2F9c7p7-2F9c7p7-2F9p7-2F9c7p7-2F9c7p7-2F9c7p7-2F9c7p7-2F9c7p7-2F9c7p7-2F9c7p7-2F9c7p7-
2F9c7^2p7-2F9c7p7-2F9c7^2p7-2F9c7p7-2F9c7^2p7-2F9c7^2p7-2F9c7p7-2F9c7p7-2F9c7p7-4p6-2F9c7p7-2F9c7p7-
2F9c7p7-2F9c7p7-2F9p7-2F9p7-4p6-2F9c7p7-2F9c7p7-2F9p7-4p6-2F9p7-4p6-2F9c7p7-2F9c7p7-2F9c7p7-
2F9c7p7-2F9c7p7-2F9c7p7-2F9c7p7-2F9c7p7-2F9c7p7-2F9c7p7-2F9c7p7-2F9c7p7-2F9c7p7-2F9c7p7-2F9p7-
4p7-2F9c7p7-2F9G9+M9p5-2F9G9+M9E9c7p7-2F9G9+M9c7p7-2F9G9+M9E9c7p7-2F9G9+M9E9c7p7-
2F9G9+M9E9c7p7-2F9G9+M9E9p7-2F9c7p7-2F9c7p7-2F9c7p7-2F9c7p7-2F9c7p7-2F9p7-5c7p7-2F9p7-5p7-
2F9c7p7-2F9c7p7-2F9c7p7-2F9c7^2p7-2F9c7p7-2F9c7p7-2F9c7p7-2F9c7p7-2F9c7p7-2F9c7p7-2F9p7-5p7-2F9p7-
4p6-2F9p7-1p3-5p7-5p7-2p7-3p7-5p7-2p7-3p7-4p7-5p7-3p7-5p7-2p5-2F9p7-5^5p7-4

### cen10

p3-5p2-5p1-2p4-5p2-3E10A10G10p4-5p2-3E10p1-7p2-3E10G10B10E10G10p1-2E10c1p1-2E10c1p1-2p5-
6c1p1-2E10c1p1-2E10c1p1-2E10c1p1-2E10p1-6p1-2E10p1-2E10c1p1-2E10c1p1-2E10c1A10p2-7p1-5G10c8p8-
2E10c1p1-2E10c1p1-2E10c1p1-2E10p1-6c1p1-2E10p1-7c1p1-2E10c1p1-2E10p1-6c1p1-2E10c1p1-2E10c1p1-
2E10c1p1-2E10p1-6c1p1-2E10c1p1-2E10c1p1-2E10c1p1-2E10p1-6c1p1-2E10p1-6c1p1-2E10c1p1-2E10c1p1-

${}_{2}E10c_{1}p_{1-2}E10c_{1}p_{1-2}E10c_{1}p_{1-2}E10p_{1-5}E10p_{1-3}B10E10p_{1-2}c_{5}E10^{2}p_{1-2}B10E10p_{1-5}c_{7}p_{7-2}E10p_{2-}$
${}_{5}C10B10p_{5-6}p_{8-2}E10c_{1}p_{1-2}E10c_{1}p_{1-2}E10c_{1}p_{1-5}E10p_{1-2}B10E10p_{1-5}p_{7-2}E10c_{1}p_{1-5}E10p_{1-2}B10E10p_{1-}$
${}_{5}c_{7}p_{7-2}c_{5}E10C10B10p_{5-2}E10c_{1}p_{1-2}c_{5}E10p_{5-2}E10p_{1-3}B10p_{5-2}E10p_{1-3}B10E10c_{1}p_{1-5}p_{1-2}E10p_{5-2}^{2}E10p_{1-}$
${}_{5}E10p_{1-3}p_{7-3}E10^{2}p_{5-2}E10p_{5-3}E10p_{5-2}p_{5-3}p_{5-2}p_{5-3}E10p_{5-2}p_{5-3}E10p_{5-3}E10p_{5-2}p_{5-3}p_{5-2}p_{5-3}E10p_{5-2}p_{5-}$
${}_{3}E10^{2}p_{5-3}E10p_{5-2}E10^{2}p_{5-3}E10p_{5-3}^{2}E10p_{5-3}E10p_{5-2}p_{5-3}E10p_{5-2}p_{5-3}E10^{2}p_{5-3}E10^{2}p_{5-2}^{2}p_{5-7}E10c_{1}p_{1-2}p_{5-}$
${}_{2}p_{5-7}E10c_{1}p_{1-2}E10c_{1}p_{1-2}p_{5-1}E10p_{1-3}p_{6-3}E10p_{5-3}^{2}E10p_{5-3}E10p_{5-2}p_{5-3}E10p_{5-2}c_{5}p_{5-2}c_{5}p_{5-2}c_{5}p_{5-2}c_{5}p_{5-2}c_{5}p_{5-}$
${}_{2}p_{5-3}E10p_{5-2}c_{5}p_{5-2}c_{5}p_{5-2}c_{5}p_{5-2}c_{5}^{2}p_{5-2}c_{5}p_{5-2}c_{5}p_{5-2}c_{5}p_{5-2}p_{5-6}A10p_{8-3}p_{5-2}p_{5-3}p_{5-2}^{2}p_{5-3}p_{5-2}^{2}c_{5}p_{5-2}^{2}c_{5}p_{5-2}c_{5}p_{5-}$
${}_{2}c_{5}^{2}p_{5-2}c_{5}^{4}p_{5-2}c_{5}p_{5-2}c_{5}p_{5-2}c_{5}p_{5-2}^{3}c_{5}p_{5-2}c_{5}p_{5-2}c_{5}p_{5-2}^{2}c_{5}p_{5-2}c_{5}p_{5-2}c_{5}p_{5-2}c_{5}p_{5-2}c_{5}p_{5-2}c_{5}p_{5-2}c_{5}p_{5-2}c_{5}p_{5-}$
${}_{2}c_{5}p_{5-2}c_{5}p_{5-2}c_{5}p_{5-2}c_{5}p_{5-2}c_{5}p_{5-2}c_{5}p_{5-2}^{2}c_{5}p_{5-2}^{2}c_{5}p_{5-2}^{2}c_{5}p_{5-2}^{2}c_{5}p_{5-2}c_{5}p_{5-2}^{2}c_{5}p_{5-2}c_{5}p_{5-2}c_{5}p_{5-2}c_{5}p_{5-2}c_{5}p_{5-2}^{2}c_{5}p_{5-}$
${}_{2}c_{5}p_{5-2}c_{5}p_{5-2}c_{5}p_{5-2}c_{5}p_{5-2}c_{5}p_{5-2}c_{5}p_{5-2}^{2}c_{5}p_{5-2}c_{5}p_{5-2}c_{5}p_{5-2}c_{5}^{2}p_{5-2}c_{5}^{5}p_{5-2}c_{5}^{2}p_{5-2}c_{5}^{2}p_{5-2}c_{5}^{2}p_{5-2}c_{5}^{2}p_{5-2}c_{5}^{2}p_{5-}$
${}_{2}c_{5}^{2}p_{5-2}c_{5}^{2}p_{5-2}c_{5}^{2}p_{5-2}c_{5}p_{5-2}c_{5}p_{5-2}c_{5}p_{5-2}c_{5}^{2}p_{5-2}c_{5}^{2}p_{5-2}c_{5}p_{5-2}c_{5}C10c_{6}^{3}p_{6-2}c_{5}^{2}p_{5-2}c_{5}p_{5-2}c_{5}C10c_{6}p_{6-2}c_{5}p_{5-}$
${}_{2}c_{5}^{2}p_{5-2}^{2}c_{5}p_{5-2}c_{5}^{2}p_{5-2}c_{5}p_{5-2}c_{5}C10c_{6}p_{6-2}c_{5}p_{5-2}c_{5}^{2}p_{5-2}^{2}c_{5}p_{5-2}c_{5}p_{5-2}c_{5}^{2}p_{5-2}c_{5}^{2}p_{5-2}^{2}c_{5}p_{5-2}c_{5}^{2}p_{5-2}c_{5}^{2}p_{5-2}^{2}c_{5}p_{5-}$
${}_{2}c_{5}^{2}p_{5-2}^{2}c_{5}p_{5-2}c_{5}^{2}p_{5-2}c_{5}^{2}p_{5-2}^{2}c_{5}p_{5-2}c_{5}p_{5-2}^{2}c_{5}p_{5-2}c_{5}p_{5-2}c_{5}p_{5-2}^{2}c_{5}p_{5-2}c_{5}E10p_{5-2}c_{5}p_{5-2}c_{5}p_{5-2}c_{5}E10p_{5-}$
${}_{2}c_{5}E10p_{5-2}c_{5}p_{5-2}c_{5}p_{5-2}c_{5}p_{5-2}c_{5}p_{5-2}^{2}c_{5}p_{5-2}c_{5}p_{5-2}^{2}c_{5}p_{5-2}^{4}c_{5}p_{5-2}^{2}c_{5}p_{5-2}^{2}c_{5}p_{5-2}c_{5}p_{5-2}c_{5}p_{5-2}c_{5}p_{5-2}^{3}c_{5}p_{5-2}^{3}c_{5}p_{5-}$
${}_{2}^{2}c_{5}p_{5-2}c_{5}p_{5-2}c_{5}p_{5-2}c_{5}p_{5-2}c_{5}p_{5-2}c_{5}p_{5-2}c_{5}p_{5-2}c_{5}^{3}p_{5-2}p_{5-3}p_{5-2}^{2}c_{5}p_{5-2}^{2}c_{5}p_{5-2}c_{5}p_{5-2}c_{5}p_{5-2}c_{5}p_{5-2}c_{5}p_{5-2}c_{5}p_{5-2}c_{5}p_{5-}$
${}_{2}c_{5}^{3}p_{5-2}p_{5-3}p_{5-2}p_{5-3}p_{5-2}c_{5}p_{5-2}c_{5}p_{5-3}p_{5-2}p_{5-3}c_{6}p_{6-2}c_{5}p_{5-2}c_{5}p_{5-2}c_{5}p_{5-2}c_{5}p_{5-2}c_{5}p_{5-2}c_{5}p_{5-2}c_{5}^{2}p_{5-2}c_{5}^{2}p_{5-2}c_{5}^{2}p_{5-}$
${}_{2}c_{5}p_{5-3}p_{5-2}p_{5-3}c_{6}p_{6-2}c_{5}p_{5-2}c_{5}p_{5-2}c_{5}p_{5-2}c_{5}p_{5-2}c_{5}^{2}p_{5-2}c_{5}p_{5-2}c_{5}^{3}p_{5-2}c_{5}p_{5-2}c_{5}p_{5-2}c_{5}p_{5-1}p_{4-2}c_{5}p_{5-2}c_{5}p_{5-2}c_{5}p_{5-2}c_{5}p_{5-}$
${}_{1}p_{4-2}c_{5}^{2}p_{5-2}c_{5}p_{5-2}c_{5}p_{5-2}c_{5}^{2}p_{5-2}c_{5}p_{5-2}c_{5}p_{5-2}c_{5}p_{5-2}c_{5}p_{5-2}c_{5}p_{5-2}c_{5}p_{5-2}^{2}c_{5}p_{5-2}c_{5}p_{5-2}c_{5}p_{5-2}c_{5}^{2}p_{5-2}c_{5}p_{5-2}c_{5}^{2}p_{5-}$
${}_{2}c_{5}p_{5-2}c_{5}p_{5-2}c_{5}p_{5-2}c_{5}p_{5-2}c_{5}^{2}p_{5-2}c_{5}p_{5-2}c_{5}p_{5-2}c_{5}p_{5-2}c_{5}p_{5-2}c_{5}p_{5-2}c_{5}p_{5-2}c_{5}^{3}p_{5-2}c_{5}p_{5-2}^{2}c_{5}p_{5-2}^{3}c_{5}^{4}p_{5-2}c_{5}p_{5-2}^{3}c_{5}^{5}p_{5-}$
${}_{2}c_{5}p_{5-2}c_{5}p_{5-2}c_{5}p_{5-2}^{2}c_{5}p_{5-2}^{3}c_{5}^{4}p_{5-2}c_{5}^{3}p_{5-2}c_{5}p_{5-2}c_{5}p_{5-2}^{2}c_{5}p_{5-2}^{2}c_{5}p_{5-2}^{3}c_{5}^{7}p_{5-2}c_{5}p_{5-2}p_{5-1}p_{4-2}p_{5-2}^{2}c_{5}^{8}p_{5-2}c_{5}p_{5-}$
${}_{2}c_{5}p_{5-2}c_{5}p_{5-2}^{2}c_{5}p_{5-2}^{2}c_{5}p_{5-2}^{2}c_{5}p_{5-2}c_{5}p_{5-2}^{2}c_{5}p_{5-2}c_{5}p_{5-2}^{2}c_{5}p_{5-2}c_{5}p_{5-2}^{2}c_{5}p_{5-2}^{2}c_{5}p_{5-2}^{2}c_{5}p_{5-2}^{2}c_{5}p_{5-2}^{2}c_{5}p_{5-2}c_{5}p_{5-}$
${}_{2}c_{5}^{3}p_{5-2}^{3}c_{5}p_{5-2}c_{5}p_{5-2}^{2}c_{5}^{2}E10B10p_{5-2}c_{5}^{2}E10B10c_{5}E10B10p_{5-2}c_{5}^{2}p_{5-2}p_{5-3}c_{5}^{2}p_{5-2}c_{5}p_{5-2}c_{5}p_{5-2}^{2}c_{5}p_{5-2}c_{5}p_{5-}$
${}_{2}c_{5}p_{5-2}^{2}c_{5}p_{5-2}^{2}c_{5}p_{5-2}^{2}c_{5}p_{5-2}c_{5}p_{5-2}c_{5}p_{5-2}c_{5}p_{5-2}c_{5}p_{5-2}^{2}c_{5}p_{5-2}^{2}c_{5}p_{5-2}^{2}c_{5}^{2}p_{5-2}^{2}c_{5}p_{5-2}^{2}c_{5}^{2}p_{5-2}c_{5}^{2}p_{5-2}^{3}c_{5}p_{5-2}c_{5}p_{5-}$
${}_{2}^{2}c_{5}^{2}p_{5-2}c_{5}^{2}p_{5-2}c_{5}p_{5-2}c_{5}p_{5-2}c_{5}^{2}p_{5-2}c_{5}p_{5-2}c_{5}p_{5-2}p_{5-3}c_{5}p_{5-2}p_{5-3}c_{6}p_{6-2}c_{5}p_{5-2}c_{5}E10c_{5}E10p_{3-5}c_{5}E10p_{3-}$
${}_{5}^{2}c_{5}E10p_{3-5}c_{5}E10p_{3-5}c_{5}E10p_{3-5}c_{5}E10p_{3-5}E10c_{5}E10p_{3-5}E10^{3}c_{5}E10p_{3-5}c_{5}E10p_{3-5}c_{5}E10p_{3-5}c_{5}E10p_{3-}$
${}_{5}E10c_{5}E10p_{3-5}E10p_{5-1}p_{4-2}p_{5-2}c_{5}^{2}p_{5-2}c_{5}E10c_{5}E10p_{5-2}c_{5}p_{5-7}c_{5}p_{5-2}c_{5}E10c_{5}p_{5-2}c_{5}p_{5-2}c_{5}p_{5-7}c_{5}p_{5-2}c_{5}p_{5-}$
${}_{7}c_{5}p_{5-2}^{2}c_{5}p_{5-2}c_{5}p_{5-2}c_{5}E10c_{5}p_{5-2}c_{5}p_{5-2}^{2}c_{5}p_{5-2}^{2}c_{5}p_{5-2}c_{5}E10c_{5}E10p_{5-2}c_{5}E10p_{5-2}c_{5}E10c_{5}^{2}E10p_{5-}$
${}_{2}c_{5}E10c_{5}p_{5-2}c_{5}E10p_{5-2}c_{5}^{2}E10p_{5-6}p_{1-2}p_{5-2}c_{5}E10p_{5-2}c_{5}p_{5-2}c_{5}p_{5-7}p_{5-2}c_{5}p_{5-2}c_{5}^{2}p_{5-7}c_{5}p_{5-7}c_{5}p_{5-7}c_{5}^{2}E10c_{5}^{2}p_{5-}$
${}_{2}c_{5}p_{5-2}E10p_{5-2}c_{5}E10c_{5}E10c_{5}E10c_{5}p_{5-2}c_{5}E10B10E10p_{5-2}c_{5}E10c_{5}p_{5-1}c_{5}E10p_{5-1}c_{5}p_{5-1}p_{1-2}p_{5-1}E10c_{1}p_{1-}$
${}_{2}c_{5}p_{5-1}E10c_{1}p_{1-2}c_{5}p_{5-2}c_{5}p_{5-2}c_{5}p_{5-7}c_{5}p_{5-7}c_{5}p_{5-2}c_{5}E10c_{5}E10c_{5}E10c_{5}p_{5-2}c_{5}p_{5-2}E10p_{5-2}c_{5}E10c_{5}E10c_{1}p_{1-}$
${}_{2}c_{5}E10^{2}c_{1}p_{1-2}E10c_{1}p_{1-2}c_{5}p_{5-2}E10c_{1}p_{1-5}p_{5-2}E10c_{1}p_{1-2}c_{5}p_{5-2}E10c_{1}p_{1-5}p_{5-2}c_{5}^{2}p_{5-2}E10p_{5-6}p_{1-2}c_{5}p_{5-}$
${}_{2}c_{5}E10p_{5-2}c_{5}E10c_{5}p_{5-2}c_{5}E10^{2}c_{1}^{2}p_{1-5}E10c_{1}^{2}p_{1-2}c_{5}p_{5-2}c_{5}E10p_{5-2}c_{5}E10p_{5-2}c_{5}E10p_{5-2}p_{5-7}p_{4-5}c_{2}p_{2-}$
${}_{5}^{2}c_{5}E10p_{3-5}p_{5-2}c_{5}E10p_{3-5}p_{2-5}p_{5-2}c_{5}E10p_{3-5}p_{5-2}c_{5}E10c_{3}c_{5}E10^{2}C10p_{7-2}c_{5}E10p_{5-2}c_{5}p_{5-2}c_{5}p_{5-2}c_{5}p_{5-}$
${}_{2}c_{5}E10p_{2-5}p_{5-2}c_{5}E10p_{2-5}c_{5}E10p_{3-5}p_{5-2}^{2}c_{5}E10p_{2-5}c_{5}E10p_{3-5}p_{5-2}c_{5}E10p_{3-5}p_{5-2}c_{5}^{2}E10p_{5-}$
${}_{2}c_{5}E10c_{3}c_{5}^{2}E10p_{5-2}c_{5}p_{5-2}^{2}c_{5}p_{5-2}^{3}c_{5}p_{5-2}^{3}c_{5}E10p_{5-2}c_{5}E10c_{5}E10c_{5}^{2}E10p_{5-2}c_{5}E10p_{5-2}c_{5}E10p_{5-2}c_{5}p_{5-}$
${}_{2}^{3}c_{5}E10p_{5-2}c_{5}^{2}E10p_{5-2}^{3}c_{5}E10p_{5-2}^{8}c_{5}E10p_{5-2}c_{5}^{2}p_{5-2}c_{5}E10p_{5-7}E10c_{3}p_{5-6}c_{3}C10p_{1-5}c_{5}E10c_{3}c_{5}p_{5-2}c_{5}p_{5-}$
${}_{2}E10c_{1}p_{1-2}c_{5}E10p_{3-5}p_{5-1}E10p_{1-2}c_{5}p_{5-2}E10c_{1}p_{1-2}c_{5}E10p_{3-5}p_{5-1}E10c_{1}p_{1-2}c_{5}p_{5-2}E10p_{1-5}c_{5}E10p_{3-5}p_{5-}$
${}_{2}c_{5}E10p_{5-2}c_{5}E10p_{3-5}p_{5-2}c_{5}E10p_{5-2}c_{5}E10p_{5-2}c_{5}E10p_{5-2}^{2}c_{5}E10c_{5}^{2}E10p_{5-2}c_{5}E10c_{3}c_{5}p_{5-2}c_{5}E10p_{5-}$
${}_{2}^{5}c_{5}E10p_{5-2}c_{5}E10c_{5}E10c_{5}E10c_{5}^{3}E10c_{5}^{2}E10c_{5}p_{5-2}^{3}c_{5}p_{5-2}^{2}p_{5-3}p_{5-}$
${}_{2}c_{5}E10c_{5}^{2}E10^{2}c_{5}^{2}E10c_{5}^{2}E10c_{5}E10c_{5}^{2}p_{5-2}c_{5}p_{5-2}^{2}c_{5}E10p_{5-2}c_{5}E10c_{5}E10c_{5}E10c_{5}^{3}E10c_{5}^{2}E10c_{5}p_{5-}$
${}_{2}^{3}c_{5}p_{5-2}^{2}p_{5-3}p_{5-2}c_{5}E10c_{5}^{2}E10^{2}c_{5}E10c_{5}^{2}E10c_{5}E10c_{5}^{2}p_{5-2}c_{5}E10p_{5-2}c_{5}E10c_{3}c_{5}p_{5-2}c_{5}E10p_{5-2}^{5}c_{5}E10p_{5-}$
${}_{2}c_{5}E10c_{5}E10c_{5}E10c_{5}^{2}p_{5-2}c_{5}p_{5-2}c_{5}E10c_{5}^{2}p_{5-2}^{2}c_{5}p_{5-2}c_{5}E10c_{5}^{3}E10c_{5}^{2}p_{5-}$
${}_{2}c_{5}E10c_{5}^{2}E10^{2}c_{5}^{2}E10c_{5}^{3}E10c_{5}E10c_{5}^{3}E10c_{5}^{3}E10c_{5}^{3}E10c_{5}^{2}E10c_{5}^{2}p_{5-}$
${}_{2}c_{5}E10c_{5}E10c_{5}E10c_{5}E10c_{5}E10c_{5}E10c_{5}E10c_{5}E10c_{5}E10c_{5}E10p_{5-2}c_{5}p_{5-2}^{2}E10c_{1}^{2}p_{1-2}E10c_{3}^{2}p_{3-5}p_{5-}$
${}_{2}E10c_{1}p_{1-2}c_{5}E10c_{3}^{2}E10c_{1}p_{1-2}p_{5-2}c_{5}p_{5-2}E10p_{1-5}p_{5-2}c_{5}^{2}E10c_{1}p_{1-2}c_{5}p_{5-2}c_{5}p_{5-2}E10p_{1-5}c_{5}^{2}E10c_{5}p_{5-}$
${}_{2}c_{5}E10c_{5}E10p_{3-5}c_{5}E10c_{5}^{2}E10c_{5}E10c_{5}E10c_{5}E10c_{5}E10c_{5}E10c_{5}E10p_{5-2}c_{5}p_{5-6}p_{8-2}E10c_{1}^{2}p_{1-2}E10c_{1}^{2}p_{1-}$
${}_{2}p_{5-2}E10c_{1}p_{1-2}E10A10p_{6-8}E10c_{1}^{2}p_{1-2}E10c_{1}p_{1-2}p_{5-2}p_{5-6}c_{1}p_{1-2}c_{5}E10p_{5-2}c_{5}E10p_{5-2}E10c_{1}^{2}p_{1-5}p_{5-}$
${}_{2}E10c_{1}^{2}p_{1-2}E10c_{1}p_{1-5}c_{5}E10B10c_{5}^{2}E10c_{1}p_{1-5}c_{1}p_{1-2}E10c_{1}p_{1-5}p_{5-6}p_{8-2}E10c_{1}p_{1-2}E10c_{1}p_{1-2}E10c_{1}^{3}p_{1-}$
${}_{2}E10c_{1}p_{1-5}c_{5}p_{5-2}E10c_{1}p_{1-2}E10c_{1}^{2}p_{1-2}E10c_{1}p_{1-5}c_{5}p_{5-2}E10c_{1}^{2}p_{1-2}E10p_{1-5}p_{5-6}p_{8-2}E10c_{1}p_{1-2}E10c_{1}^{2}p_{1-}$
${}_{2}E10c_{1}p_{1-5}p_{5-6}p_{8-2}E10c_{1}p_{1-2}E10c_{1}p_{1-5}c_{1}p_{1-5}p_{5-6}p_{8-2}E10c_{1}p_{1-2}E10c_{1}^{3}p_{1-5}p_{5-2}E10c_{1}p_{1-2}E10c_{1}p_{1-}$
${}_{2}E10c_{1}p_{1-2}E10p_{1-2}p_{5-1}p_{4-5}c_{5}E10^{2}p_{1-5}c_{5}E10c_{5}E10p_{2-5}c_{5}E10c_{5}E10^{2}c_{1}p_{1-2}c_{5}E10p_{5-2}c_{5}E10c_{5}p_{5-}$

$_{2}E10c_1^2p_{1-2}p_{5-2}E10c_1p_{1-2}E10c_1^2p_{1-2}p_{5-2}c_5p_{5-7}E10c_1p_{1-5}p_{5-2}^2p_{5-7}E10A10p_{7-5}p_{5-6}c_1p_{1-2}c_5p_{5-2}E10c_1p_{1-2}p_{5-2}$
$c_5E10p_{5-2}E10c_1p_{1-5}p_{5-2}c_5E10c_3p_{5-3}p_{1-2}c_5E10c_1p_{1-2}p_{5-2}c_5p_{5-2}E10c_1p_{1-2}c_5p_{5-2}c_5^2E10p_{5-2}E10p_{1-5}c_5p_{5-2}^2$
$E10c_1p_{1-2}p_{5-2}^2E10c_1p_{1-2}E10c_1p_{1-2}E10p_{1-5}c_5E10p_{5-2}^2E10c_1A10E10c_1p_{1-2}E10c_1^2p_{1-2}E10c_1^3p_{1-2}$
$E10c_1p_{1-2}E10p_{1-3}c_5p_{5-2}E10c_1^2p_{1-2}E10c_1p_{1-2}E10p_{1-3}c_5p_{5-2}E10c_1p_{1-5}c_5E10^2c_1^2p_{1-5}E10p_{1-5}c_3^2p_{3-5}$
$B10E10p_{7-2}p_{5-2}p_{5-8}B10c_5E10c_1p_{1-2}c_5E10c_1p_{1-2}p_{5-2}^2c_5E10^2c_1p_{1-5}E10c_1p_{1-5}E10c_1^3p_{1-6}B10E10c_7p_{7-3}$

### cen11

$p_{1-4}c_1^{61}c_2^{71}p_{2-3}c_4^{41}D11c_1^{75}p_{1-2}c_5^{92}p_{5-2}c_4^{193}p_{4-1}c_3^{142}p_{3-1}A11c_3^{136}C11c_2^6p_{2-3}c_2^{129}p_{2-5}c_2^{179}p_{2-3}c_5^{150}p_{5-1}$
$A11c_3^{88}C11c_2^3p_{2-3}c_2^{240}B11c_5p_{5-1}c_3c_5^{48}p_{5-1}c_3^{24}p_{3-1}c_3^{170}p_{3-1}c_4^{26}p_{4-2}c_5p_{5-1}c_3c_5^{151}p_{5-1}c_3^{18}c_4^{89}p_{4-1}c_3^{22}p_{3-1}$
$c_3^{84}p_{3-1}c_3^{22}p_{3-1}c_3^{86}c_5^{25}p_{5-2}c_5^{157}p_{5-2}c_5^{26}p_{5-2}c_5^{54}p_{5-2}c_5^4p_{5-2}c_5^8p_{5-2}c_5^4p_{5-2}^2c_5^{18}p_{5-1}c_1^{48}p_{1-2}c_4^5p_{4-2}c_4^{330}p_{4-2}$
$c_4^{18}p_{4-2}c_5^3p_{5-2}c_5p_{5-1}c_3c_5p_{5-1}c_3^{291}p_{3-5}c_2^{56}p_{2-5}c_2^{86}p_{2-4}c_1^{27}p_{1-4}c_1^{37}p_{1-4}c_1^{237}p_{1-3}c_4^{79}p_{4-1}c_4^{11}p_{4-1}c_4^4p_{4-1}c_4^{23}$

### cen12

$F12H12D12p_{6-1}C12E12p_{3-4}p_{6-8}p_{4-1}^2p_{4-8}^2p_{3-4}p_{6-7}p_{4-1}p_{3-8}p_{3-7}p_{4-1}p_{3-4}p_{6-8}p_{4-8}^2p_{3-4}p_{6-8}D12p_{6-8}p_{4-8}D12p_{6-8}$
$p_{3-8}p_{4-8}p_{6-8}p_{4-1}D12p_{7-8}D12p_{6-8}D12p_{6-1}D12p_{6-1}D12p_{6-1}D12F12D12p_{6-4}^3p_{6-3}p_{6-4}p_{6-8}p_{2-4}p_{6-4}c_5^2p_{6-1}p_{6-8}$
$p_{4-8}D12p_{6-8}p_{3-8}p_{4-8}p_{6-8}p_{4-1}D12p_{7-8}D12p_{6-8}D12p_{6-1}D12p_{6-1}D12p_{6-1}D12F12D12p_{6-4}^3p_{6-3}p_{6-4}p_{6-8}p_{2-4}p_{6-4}$
$c_5^2p_{6-4}c_5p_{6-7}c_8p_{5-1}D12p_{6-1}D12p_{6-7}A12D12p_{6-8}p_{7-1}D12p_{6-1}p_{6-8}D12p_{6-1}D12p_{6-1}p_{7-8}p_{4-1}p_{3-4}F12D12p_{6-1}$
$D12p_{6-7}c_8B12F12p_{2-4}c_5p_{2-5}p_{6-4}^2c_6p_{1-4}p_{6-1}c_2^4p_{6-4}c_5c_6p_{1-4}p_{7-4}p_{6-4}c_5^3F12c_7^2p_{2-3}p_{8-3}^2p_{8-4}F12c_7^2p_{2-3}p_{8-3}^2$
$p_{8-4}p_{6-1}C12p_{2-4}c_5^3p_{6-3}c_4p_{8-1}c_2p_{6-1}c_2D12p_{6-4}F12c_7p_{2-4}p_{6-7}p_{6-4}F12c_7p_{2-4}p_{6-7}D12p_{6-4}F12c_7p_{2-4}p_{6-7}$
$D12p_{6-4}F12c_7p_{1-4}p_{6-4}F12c_7A12D12F12c_7p_{2-4}p_{6-7}p_{6-3}p_{6-4}F12c_7p_{2-4}p_{6-7}p_{6-4}F12c_7p_{2-4}p_{6-7}p_{6-4}^3p_{6-8}p_{2-5}p_{2-4}$
$p_{6-2}p_{4-8}c_1c_5^4p_{6-4}p_{6-1}c_2^3p_{6-1}c_2^3F12p_{6-3}c_4^3p_{8-1}p_{6-3}c_2^2p_{6-1}^2c_2p_{6-1}c_2p_{6-1}c_2^3F12p_{6-3}c_4^4p_{8-1}p_{6-3}c_2^2p_{6-3}c_4^3p_{2-3}$
$c_4^2p_{2-3}c_4^2p_{2-3}c_4^2p_{2-4}c_5p_{6-4}p_{2-7}p_{7-4}c_5c_6p_{1-4}F12c_7p_{2-4}p_{6-7}p_{6-4}^3F12c_7p_{2-4}p_{7-4}p_{6-4}^9F12c_7^2p_{5-7}c_2p_{6-3}c_4p_{2-7}$
$c_8^2p_{2-3}c_4^3p_{2-3}c_4^2p_{2-3}c_4^3p_{2-3}c_4^2p_{2-4}c_5c_6p_{2-4}c_5F12c_7p_{2-4}p_{6-7}p_{6-4}F12c_7p_{2-4}p_{6-4}F12c_7p_{2-4}p_{6-4}^2F12p_{6-4}^3$
$F12p_{6-4}^7F12c_7^3c_5p_{2-7}c_8p_{2-4}c_5^2p_{6-4}c_5p_{6-4}F12c_7^3p_{5-7}p_{2-4}c_5F12c_7^3p_{5-7}p_{2-4}c_5p_{2-7}c_8p_{2-5}p_{2-4}p_{2-7}c_8p_{2-4}c_5^2p_{6-4}$
$c_5p_{6-4}F12c_7^3p_{5-7}p_{2-7}^2c_8p_{2-8}c_1^2p_{2-7}p_{2-5}c_6^2p_{1-7}c_8^3p_{2-5}p_{4-7}c_8^7p_{2-7}c_8^2p_{2-8}c_1p_{3-7}c_8p_{2-7}^2c_8p_{2-7}c_8p_{2-7}^2c_8p_{2-7}c_8p_{2-7}$
$c_8p_{2-7}c_8p_{2-7}c_8^2p_{2-7}c_8^3c_2^3p_{4-1}c_2p_{4-2}c_3p_{4-7}c_2^6p_{4-7}p_{2-7}c_8^{10}p_{2-7}c_8^{16}p_{2-4}c_5^2p_{2-8}c_1p_{2-8}c_1p_{2-7}p_{2-8}c_1p_{2-7}^2c_8p_{2-8}c_1p_{2-8}$
$c_1p_{2-7}^2c_8p_{2-7}c_8p_{2-8}c_1^2p_{2-7}c_8^3p_{2-8}c_1^2c_2^7p_{4-7}p_{2-7}c_8^5B12p_{6-3}c_4^3c_2^2p_{6-7}c_8p_{2-3}c_2p_{6-7}c_8p_{2-3}p_{2-7}c_8p_{2-7}c_8p_{2-8}c_1^2p_{2-7}$
$c_2^5p_{4-7}c_2^2p_{4-7}p_{2-7}c_8^4p_{2-7}c_2^3p_{6-7}c_8p_{2-3}p_{2-7}c_8p_{2-3}p_{2-7}c_8p_{2-8}c_1^2p_{2-7}^2c_8p_{2-7}c_8^2c_2^5p_{4-7}p_{2-3}c_4^2c_2^2p_{6-7}c_8^{14}p_{2-5}p_{4-7}$
$c_8p_{2-8}c_1^2p_{2-7}^2p_{4-7}c_8^4p_{2-5}p_{4-7}c_8p_{4-7}p_{6-7}c_8^2p_{6-8}c_1^3p_{2-6}p_{2-7}p_{2-8}c_1p_{2-7}p_{2-8}c_1^2p_{2-7}^2c_8p_{2-7}c_8p_{2-5}G12c_8B12c_3p_{4-7}$
$p_{2-7}c_8p_{2-7}c_8^2p_{2-7}c_8^2p_{2-8}c_1^2p_{2-7}p_{2-8}c_1^3p_{2-7}^2c_8p_{2-7}c_8B12c_3p_{4-7}p_{2-7}c_8^2c_2^3p_{4-7}p_{2-7}c_8^3p_{2-7}^2p_{6-7}c_8p_{2-7}^2p_{2-3}$
$c_4^6c_2^2p_{6-3}c_4^2c_2p_{6-3}c_4^4c_2^2p_{6-7}c_8p_{2-3}c_2^8p_{4-7}c_8^2p_{2-5}p_{4-6}p_{2-6}c_7p_{5-7}p_{2-5}p_{4-7}c_8p_{6-1}c_2p_{6-7}c_8p_{2-5}c_6D12p_{1-7}c_8^2p_{2-7}$
$c_8^2p_{2-5}c_6^2p_{4-5}c_6^2p_{4-7}c_8^3p_{6-7}c_8p_{6-7}c_8p_{6-7}c_8p_{6-7}c_8p_{6-4}c_5^3p_{1-4}c_5^5c_6p_{2-4}c_5^6c_6p_{2-7}c_8^2p_{6-1}c_2p_{6-1}c_2p_{6-1}^2c_2^2p_{6-1}$
$c_2^3p_{6-1}^2c_2p_{6-1}c_2p_{6-1}c_2p_{6-1}c_2p_{6-1}c_2p_{6-1}c_2p_{6-1}^3c_2p_{6-1}^3c_2p_{6-1}c_2p_{6-3}p_{6-1}^2c_2^6p_{6-1}c_2^2p_{6-1}c_2^3p_{6-1}c_2p_{6-1}c_2p_{6-1}c_2p_{6-1}$
$c_2p_{6-1}c_2p_{6-1}c_2p_{6-1}c_2^2p_{6-1}c_2^2p_{6-1}c_2^2F12c_7^2p_{3-4}A12c_2^7p_{6-1}c_2p_{6-1}c_2^6p_{6-1}c_2p_{6-1}c_2p_{6-1}c_2^4p_{6-1}c_2^{11}p_{6-1}^2c_2p_{6-1}$
$c_2p_{6-1}c_2p_{6-1}c_2^2p_{6-1}c_2^2p_{6-1}c_2^9p_{6-1}c_2p_{6-1}c_2p_{6-1}c_2^2p_{6-4}c_5A12p_{6-1}c_2^2p_{6-1}c_2p_{6-1}c_2p_{6-1}c_2p_{6-8}p_{4-1}p_{6-1}c_2^2p_{6-1}c_2p_{6-1}$
$c_2p_{6-1}c_2^3p_{6-1}c_2^{14}p_{6-1}c_2^2p_{6-1}c_2p_{6-1}c_2p_{6-1}c_2p_{6-1}c_2^2p_{6-1}c_2p_{6-1}c_2^2p_{6-1}c_2p_{6-1}c_2^3p_{6-1}c_2p_{6-4}c_5^3D12c_5p_{4-1}p_{6-1}c_2p_{6-1}^2$
$c_2p_{6-1}c_2p_{6-1}c_2p_{6-1}c_2^2p_{6-2}p_{7-4}c_5^3p_{7-1}c_2p_{6-1}c_2p_{6-1}c_2^2p_{6-1}c_2p_{6-1}c_2p_{6-1}c_2^2p_{6-1}c_2p_{6-1}c_2p_{6-1}c_2p_{6-1}c_2^2p_{6-7}c_8^2p_{2-7}$
$c_8^2p_{2-3}c_4^9p_{2-7}c_8c_2^2F12c_7^{14}E12p_{4-7}c_8p_{4-7}p_{6-7}c_8^2p_{2-7}c_8p_{2-8}c_1^3p_{2-7}^2c_8p_{2-5}p_{7-8}c_1^2p_{2-7}p_{2-8}c_1^3p_{2-7}^2c_8p_{2-5}p_{7-8}$
$c_1^2p_{2-7}^2c_8p_{2-8}c_1^2p_{2-7}^2c_8^3p_{2-7}c_8^2c_2^5p_{6-7}c_8p_{2-3}^2c_4^3c_2^3p_{6-1}c_2^6p_{6-1}c_2^5p_{6-1}c_2^5p_{6-1}c_2^5p_{6-7}c_8^9p_{2-5}p_{4-7}c_8p_{4-7}p_{6-7}$
$c_8^2p_{2-7}c_8p_{2-8}c_1p_{2-7}p_{2-5}c_6p_{7-8}c_1p_{2-7}c_8p_{2-8}c_1^2p_{2-7}p_{2-8}c_1p_{2-7}^2c_8p_{2-8}c_1^3p_{2-7}c_8p_{2-7}c_8p_{2-7}c_8^3p_{2-7}c_8^6p_{2-7}c_8p_{2-8}$
$c_1^3p_{2-7}^2c_8p_{2-7}^2c_8p_{2-7}c_8p_{2-7}c_8^2p_{2-6}c_7^2p_{1-7}c_8c_2^6p_{4-7}c_2^6p_{4-7}p_{2-3}c_4^3c_2^2p_{6-3}c_4^4p_{2-3}c_4^4c_2^2p_{6-3}c_4^2p_{2-7}c_8^{13}p_{2-5}p_{4-7}$
$c_8p_{4-7}c_8^2p_{6-7}c_8^3p_{2-5}p_{4-7}c_8p_{4-7}c_8^2p_{6-1}c_2p_{6-1}c_2p_{6-7}p_{2-7}p_{4-7}c_8^2p_{6-1}c_2p_{6-1}c_2p_{6-7}c_8p_{2-5}c_6p_{4-1}c_2^{13}F12c_7p_{5-7}p_{2-5}p_{4-7}$
$c_8p_{4-7}c_8^2c_6p_{4-5}c_6^2p_{4-7}c_8p_{4-7}c_8^2c_6D12c_5A12p_{6-7}c_8p_{2-5}D12c_5^2p_{1-5}p_{4-1}c_2c_8p_{2-7}c_8p_{2-8}c_1E12p_{4-1}c_2^2p_{6-8}p_{5-7}$
$c_8p_{6-8}p_{5-7}c_8p_{6-7}c_8^3p_{6-8}p_{5-7}c_8p_{6-7}c_8^8F12c_7^2c_5^4p_{1-4}c_5^5p_{1-4}c_5^4c_6p_{2-4}c_5^3c_6c_2p_{6-1}c_2^3p_{6-1}c_2p_{6-1}c_2p_{6-1}c_2p_{6-1}c_2p_{6-1}$
$c_2p_{6-1}c_2p_{6-4}c_5A12c_2p_{6-1}c_2p_{6-1}c_2^2p_{6-1}c_2p_{6-1}c_2p_{6-4}A12c_2p_{6-1}c_2p_{6-1}c_2^2p_{6-1}c_2p_{6-1}c_2p_{6-1}c_2p_{6-1}c_2p_{6-1}c_2p_{6-4}$
$c_5A12c_2p_{6-1}c_2p_{6-1}c_2^4p_{6-1}c_2^3p_{6-1}c_2^2p_{6-3}c_4p_{8-1}p_{6-1}^2c_2^6p_{6-1}c_2^3p_{6-1}c_2p_{6-3}c_4^3p_{8-1}p_{6-1}^2c_2^7p_{6-1}c_2^4p_{6-1}c_2^3p_{6-1}c_2^3p_{6-1}$
$c_2p_{6-1}c_2p_{6-1}c_2^4p_{6-1}c_2^3p_{6-1}c_2^3p_{6-1}c_2p_{6-3}c_4^6p_{8-1}p_{6-1}c_2^5p_{6-3}c_4^2p_{8-1}F12c_7^2p_{3-4}A12c_2^6p_{6-3}c_4p_{8-1}p_{6-1}c_2p_{6-3}c_4p_{8-1}$
$p_{6-1}p_{6-3}c_4p_{8-1}p_{6-3}c_4^2p_{8-1}p_{6-1}p_{6-3}c_4p_{8-1}p_{6-3}c_4p_{8-1}p_{6-1}p_{6-3}c_4^2p_{8-1}p_{6-1}c_2^6p_{6-1}c_2^3p_{6-3}c_4^2p_{8-1}p_{6-1}p_{6-3}c_4p_{8-1}p_{6-}$

$_{3}c_{4}p_{8-1}p_{6-1}p_{6-3}c_{4}p_{8-1}p_{6-3}c_{4}^{2}p_{8-1}p_{6-3}c_{4}p_{8-1}p_{6-1}c_{2}^{7}p_{6-3}c_{4}^{2}p_{8-1}p_{6-3}c_{4}^{2}p_{8-1}p_{6-1}p_{6-3}c_{4}p_{8-1}p_{6-3}c_{4}p_{8-1}p_{6-1}p_{6-3}c_{4}^{2}p_{8-1}$
$p_{6-1}c_{2}^{6}p_{6-1}c_{2}^{3}p_{6-3}c_{4}^{2}p_{8-1}p_{6-1}p_{6-3}c_{4}p_{8-1}p_{6-3}c_{4}p_{8-1}p_{6-1}p_{6-3}c_{4}p_{8-1}p_{6-3}c_{4}^{2}p_{8-1}p_{6-3}c_{4}p_{8-1}p_{6-1}c_{2}^{7}p_{6-3}c_{4}^{2}p_{8-1}p_{6-1}$
$c_{2}^{8}p_{6-3}c_{4}^{2}p_{8-1}p_{6-1}p_{6-3}c_{4}p_{8-1}\mathrm{F12}c_{7}\mathrm{C12}p_{8-1}p_{6-1}p_{6-3}c_{4}p_{8-1}p_{6-1}c_{2}p_{6-3}c_{4}p_{8-1}p_{6-1}c_{2}p_{6-3}c_{4}p_{8-1}p_{6-1}p_{6-3}c_{4}p_{8-1}p_{6-1}$
$c_{2}^{8}\mathrm{F12}c_{2}p_{6-1}c_{2}p_{6-3}c_{4}p_{8-1}p_{6-4}c_{5}p_{1-3}p_{8-1}p_{6-1}c_{2}p_{6-4}c_{5}p_{1-3}p_{8-1}p_{6-1}c_{2}^{4}\mathrm{F12}c_{2}p_{6-3}c_{4}p_{8-1}p_{6-4}c_{5}p_{1-3}p_{8-1}p_{6-1}c_{2}p_{6-4}$
$c_{5}p_{1-3}p_{8-1}p_{6-4}\mathrm{A12}c_{2}p_{6-1}c_{2}^{2}p_{6-1}c_{2}^{2}p_{6-1}c_{2}^{2}p_{6-1}c_{2}^{2}p_{6-4}c_{5}^{17}p_{8-1}c_{2}^{10}p_{4-6}c_{7}^{9}\mathrm{F12}c_{7}^{6}p_{6-7}c_{8}^{23}\mathrm{D12}p_{1-7}c_{8}^{6}\mathrm{D12}p_{1-7}$
$c_{8}^{2}p_{4-7}c_{8}^{12}p_{4-7}c_{8}^{97}\mathrm{D12}p_{1-7}\mathrm{D12}p_{1-7}c_{8}^{2}\mathrm{D12}p_{1-7}\mathrm{D12}p_{1-7}c_{8}^{4}\mathrm{D12}p_{1-4}c_{5}\mathrm{G12}c_{8}^{2}\mathrm{D12}p_{1-7}c_{8}^{2}\mathrm{D12A12}c_{2}^{10}p_{4-7}$
$c_{8}^{47}\mathrm{D12}p_{1-7}c_{8}^{2}\mathrm{D12}p_{1-7}c_{8}^{2}\mathrm{D12}p_{1-7}\mathrm{D12}p_{1-7}c_{8}^{4}\mathrm{D12}p_{1-4}c_{5}\mathrm{G12}c_{8}^{2}\mathrm{D12}p_{1-7}c_{8}^{2}\mathrm{D12A12}c_{2}^{10}c_{4}^{5}p_{3-4}p_{6-4}^{2}p_{6-8}$
$\mathrm{C12}c_{4}^{2}p_{3-4}p_{6-3}c_{4}p_{3-4}p_{6-3}c_{4}p_{3-4}p_{6-3}c_{4}p_{3-4}p_{6-1}c_{2}p_{4-8}c_{1}^{5}\mathrm{C12}c_{4}^{2}p_{3-4}p_{6-4}^{2}p_{6-8}\mathrm{C12}c_{4}^{2}p_{3-4}p_{6-3}c_{4}p_{3-4}p_{6-4}^{2}p_{6-8}$
$\mathrm{C12}c_{4}\mathrm{D12}p_{6-4}p_{2-5}c_{6}p_{3-8}c_{2}p_{8-1}\mathrm{C12}p_{8-1}\mathrm{D12}p_{6-1}p_{3-8}c_{1}^{3}p_{2-4}p_{6-3}\mathrm{H12}c_{4}p_{2-4}c_{5}p_{6-7}p_{2-4}p_{6-3}p_{8-1}c_{2}p_{6-3}\mathrm{H12}p_{2-6}p_{8-1}$
$p_{6-3}c_{4}\mathrm{H12}p_{2-4}p_{6-4}c_{5}^{2}p_{6-4}c_{5}p_{6-4}p_{6-8}p_{5-2}p_{8-6}\mathrm{D12}p_{6-4}^{2}c_{5}p_{6-7}p_{1-4}p_{6-7}p_{2-4}\mathrm{F12}p_{6-7}p_{2-4}\mathrm{F12}p_{6-4}^{2}p_{6-8}p_{2-4}p_{6-8}p_{2-4}$
$p_{6-8}c_{1}p_{2-4}p_{6-4}p_{6-8}c_{1}p_{2-4}p_{6-4}c_{5}p_{6-8}p_{2-8}\mathrm{B12D12}c_{5}p_{6-8}p_{2-4}\mathrm{H12}p_{2-3}c_{4}p_{8-1}c_{4}^{2}p_{8-1}\mathrm{D12}c_{5}^{2}p_{6-8}p_{2-4}\mathrm{H12}p_{2-8}p_{2-4}$
$c_{5}p_{6-8}p_{2-4}\mathrm{H12}p_{2-4}p_{6-3}p_{8-1}\mathrm{D12}c_{5}p_{6-8}p_{2-4}p_{6-1}\mathrm{D12}c_{5}p_{6-4}p_{7-4}p_{6-4}^{4}p_{6-3}c_{4}p_{8-4}p_{6-4}^{2}p_{6-3}c_{4}p_{8-4}p_{6-4}p_{6-1}p_{8-1}\mathrm{D12}^{2}p_{6-7}$
$\mathrm{D12G12D12}p_{6-7}\mathrm{A12}p_{3-4}\mathrm{H12}p_{3-4}\mathrm{G12}$

### cen13

$p_{2-11}p_{8-9}c_{11}\mathrm{G13/21.0}c_{1}p_{8-11}c_{1}p_{8-10}\mathrm{G13/21.0}c_{1}p_{8-11}c_{1}^{3}c_{8}^{4}p_{1-7}c_{8}p_{1-7}c_{8}p_{1-7}^{2}c_{8}p_{1-7}c_{8}p_{1-7}p_{1-5}p_{11-7}p_{1-7}^{4}c_{8}^{3}p_{1-7}$
$c_{8}p_{1-7}^{2}c_{8}p_{1-7}c_{8}p_{1-7}c_{8}p_{1-7}^{3}c_{8}p_{1-7}c_{8}p_{1-7}^{2}c_{8}^{4}p_{1-7}p_{1-2}c_{3}^{2}p_{9-7}p_{1-7}^{3}c_{8}^{3}p_{1-7}c_{8}p_{1-7}c_{8}p_{1-7}c_{8}p_{1-7}c_{8}^{2}p_{1-7}c_{8}p_{1-7}c_{8}p_{1-7}$
$c_{8}p_{1-7}^{4}c_{8}p_{1-7}c_{8}p_{1-7}c_{8}p_{1-7}^{3}c_{8}^{2}p_{1-7}c_{8}^{5}p_{1-7}^{2}c_{8}^{2}p_{1-7}^{2}p_{1-10}\mathrm{G13/21.0}p_{1-7}^{2}c_{8}c_{1}c_{8}p_{1-7}^{3}c_{8}p_{1-7}c_{8}p_{1-7}c_{8}p_{1-7}^{3}c_{8}p_{1-7}$
$c_{8}p_{1-7}^{2}c_{8}p_{1-7}c_{8}p_{1-7}^{3}c_{8}^{2}p_{1-7}^{3}c_{8}p_{1-7}^{2}c_{8}p_{1-7}c_{8}p_{1-7}^{3}c_{8}p_{1-7}c_{8}^{2}p_{1-7}^{5}c_{1}c_{8}^{2}p_{1-7}^{3}c_{8}p_{1-7}^{2}c_{8}^{2}p_{1-7}c_{8}p_{1-7}c_{8}p_{1-7}^{2}c_{8}p_{1-7}^{2}$
$c_{8}p_{1-7}^{5}c_{8}^{2}p_{1-7}c_{8}^{2}p_{1-7}^{3}c_{8}p_{1-7}^{2}c_{8}p_{1-7}c_{1}c_{8}p_{1-7}^{2}c_{8}p_{1-7}c_{1}c_{8}p_{1-7}c_{8}p_{1-7}c_{8}p_{1-7}^{3}c_{8}p_{1-7}c_{8}^{2}p_{1-7}^{2}c_{8}p_{1-7}c_{8}p_{1-7}^{2}c_{8}p_{1-7}$
$c_{8}p_{1-7}^{2}c_{8}p_{1-7}^{2}c_{8}p_{1-7}c_{8}p_{1-7}^{3}c_{8}p_{1-7}c_{8}p_{1-7}^{2}p_{1-3}p_{4-7}p_{1-7}^{5}c_{1}c_{8}^{3}p_{1-7}^{2}c_{8}p_{1-7}^{3}c_{8}p_{1-7}^{3}c_{1}^{5}p_{8-11}c_{1}p_{8-11}^{2}c_{1}^{2}c_{8}^{3}p_{1-7}$
$c_{8}^{2}c_{1}^{3}c_{8}p_{1-7}^{2}c_{8}^{2}p_{1-7}c_{8}^{2}p_{1-7}^{2}c_{8}p_{1-7}c_{8}p_{1-4}c_{5}p_{11-7}c_{8}p_{1-7}c_{8}p_{1-7}c_{8}p_{1-7}c_{8}p_{1-7}^{2}c_{8}^{2}p_{1-7}c_{8}p_{1-7}c_{1}^{9}c_{8}^{4}p_{1-7}^{2}c_{8}^{2}p_{1-7}c_{8}^{2}p_{1-7}$
$c_{8}p_{1-7}^{5}c_{8}^{2}p_{1-7}c_{8}^{2}p_{1-7}c_{8}^{2}p_{1-7}c_{8}p_{1-7}^{7}c_{8}p_{1-7}c_{8}^{2}p_{1-7}^{7}c_{8}^{2}p_{1-7}c_{8}^{2}p_{1-7}^{3}c_{8}p_{1-7}^{3}c_{8}p_{1-7}c_{1}c_{8}p_{1-7}^{2}c_{8}p_{1-7}c_{1}c_{8}p_{1-7}c_{8}p_{1-7}$
$c_{8}p_{1-7}^{3}c_{8}p_{1-7}c_{8}p_{1-7}^{3}c_{8}p_{1-7}^{2}c_{8}p_{1-7}c_{8}p_{1-7}^{2}c_{8}p_{1-7}^{4}c_{8}p_{1-7}c_{8}p_{1-7}^{2}c_{8}p_{1-7}^{2}c_{8}p_{1-7}c_{8}p_{1-7}^{3}c_{8}p_{1-7}c_{8}p_{1-7}^{8}c_{1}c_{8}^{3}p_{1-7}^{2}c_{8}p_{1-7}^{3}$
$c_{8}p_{1-7}^{3}c_{1}^{5}p_{8-11}c_{1}p_{8-11}^{2}c_{1}^{2}c_{8}^{3}p_{1-7}c_{8}^{2}c_{1}^{3}c_{8}p_{1-7}^{2}c_{8}^{2}p_{1-7}c_{8}^{2}p_{1-7}^{2}c_{8}p_{1-7}c_{8}p_{1-7}c_{8}^{2}p_{1-7}c_{8}p_{1-7}c_{8}p_{1-7}c_{8}p_{1-7}^{2}c_{8}^{2}p_{1-7}$
$c_{8}p_{1-7}^{2}c_{8}^{2}p_{1-7}^{2}c_{8}^{2}p_{1-7}c_{8}p_{1-7}c_{1}^{7}c_{8}^{3}p_{1-7}^{2}c_{8}^{2}p_{1-7}c_{8}^{2}p_{1-7}c_{8}p_{1-7}^{5}c_{8}^{2}p_{1-7}c_{8}^{2}p_{1-7}c_{8}^{2}p_{1-7}c_{8}p_{1-7}^{7}c_{8}p_{1-7}c_{8}^{2}p_{1-7}^{4}p_{1-8}p_{3-7}$
$p_{1-7}^{4}p_{1-5}\mathrm{G13/21.0}c_{8}p_{1-7}^{2}p_{1-5}\mathrm{G13/21.0}c_{8}p_{1-7}^{2}c_{8}p_{1-7}^{2}c_{8}^{2}p_{1-2}c_{3}p_{9-7}c_{8}p_{1-7}c_{8}^{2}p_{1-7}c_{8}p_{1-7}c_{8}^{4}p_{1-7}c_{8}^{2}p_{1-7}c_{8}p_{1-7}$
$c_{8}p_{1-7}c_{8}p_{1-7}c_{8}p_{1-7}^{2}c_{8}p_{1-7}^{2}c_{8}p_{1-4}\mathrm{G13/21.0}c_{8}p_{1-7}^{2}p_{1-4}\mathrm{G13/21.0}c_{8}p_{1-7}^{3}c_{8}p_{1-7}^{2}c_{8}p_{1-7}^{9}c_{8}p_{1-7}c_{8}p_{1-7}^{2}c_{8}^{2}p_{1-7}^{3}$
$c_{8}p_{1-7}c_{8}p_{1-7}^{3}c_{8}^{6}p_{1-7}c_{8}p_{1-7}c_{8}p_{1-7}^{5}c_{8}^{2}p_{1-7}^{2}c_{8}p_{1-7}^{2}c_{8}^{3}p_{1-7}^{2}p_{1-6}c_{7}p_{5-7}c_{8}p_{1-7}^{3}c_{8}p_{1-7}^{2}c_{8}p_{1-7}c_{8}p_{1-7}^{2}c_{8}p_{1-7}p_{1-3}p_{4-7}$
$p_{1-7}^{2}c_{8}p_{1-7}^{2}c_{8}p_{1-7}^{2}c_{8}^{2}p_{1-7}^{2}c_{8}p_{1-7}c_{8}^{2}p_{1-7}c_{8}p_{1-7}c_{8}p_{1-7}^{2}c_{8}p_{1-7}^{7}c_{8}^{2}p_{1-7}c_{8}^{2}p_{1-7}^{2}c_{8}^{4}p_{1-7}^{3}c_{8}p_{1-7}^{9}c_{8}^{2}p_{1-7}c_{8}^{2}p_{1-7}c_{8}^{4}p_{1-7}^{2}$
$c_{8}^{3}p_{1-7}^{2}c_{8}^{2}p_{1-7}c_{8}p_{1-7}^{3}c_{8}p_{1-7}c_{8}p_{1-7}c_{8}p_{1-7}c_{8}^{2}p_{1-7}^{3}c_{8}p_{1-7}c_{8}^{2}p_{1-7}c_{8}p_{1-7}^{2}c_{8}p_{1-7}^{2}c_{8}^{2}p_{1-7}^{6}c_{8}p_{1-7}c_{1}c_{8}p_{1-6}c_{7}p_{11-7}p_{1-7}^{3}$
$p_{1-6}p_{11-7}p_{1-7}^{3}c_{8}p_{1-7}^{3}c_{8}p_{1-7}^{2}c_{8}p_{1-7}c_{8}p_{1-7}c_{8}p_{1-7}c_{8}p_{1-7}c_{8}p_{1-7}c_{8}p_{1-7}c_{8}p_{1-7}c_{8}p_{1-7}^{2}c_{8}p_{1-7}^{6}c_{8}^{2}p_{1-7}c_{8}p_{1-7}^{2}c_{8}^{4}p_{1-7}^{2}$
$c_{8}p_{1-7}^{5}c_{8}^{2}p_{1-7}c_{8}^{2}p_{1-7}^{4}c_{8}^{2}p_{1-7}c_{8}^{2}p_{1-7}c_{8}^{4}p_{1-7}^{2}c_{8}p_{1-7}^{5}c_{8}^{2}p_{1-7}^{2}c_{8}^{2}p_{1-7}c_{8}^{2}c_{1}p_{8-11}c_{1}^{2}c_{8}^{4}p_{1-7}c_{8}^{2}p_{1-7}^{6}c_{8}^{2}p_{1-7}c_{8}^{6}p_{1-7}^{2}$
$c_{8}^{4}p_{1-7}^{2}c_{8}^{4}p_{1-7}^{2}c_{8}^{3}p_{1-7}c_{8}p_{1-7}^{2}c_{8}p_{1-7}c_{8}p_{1-7}^{3}c_{8}p_{1-7}^{3}c_{8}^{2}p_{1-7}c_{8}p_{1-7}^{6}c_{8}^{4}p_{1-7}^{5}c_{8}p_{1-7}^{2}c_{8}p_{1-7}^{3}c_{8}p_{1-7}c_{8}p_{1-7}^{2}c_{8}p_{1-7}^{4}$
$c_{8}^{2}p_{1-7}^{6}c_{8}p_{1-7}^{2}c_{8}p_{1-7}^{7}c_{8}p_{1-7}c_{8}p_{1-7}^{5}p_{1-6}p_{1-7}^{2}p_{1-6}p_{1-7}^{4}p_{1-6}p_{5-7}p_{1-7}^{7}p_{1-6}p_{1-7}^{3}p_{1-3}c_{4}p_{4-7}c_{8}p_{1-7}c_{8}p_{1-7}c_{8}^{4}p_{1-7}c_{8}^{4}p_{1-7}^{4}$
$c_{8}^{3}p_{1-7}^{3}c_{8}^{3}p_{1-7}c_{8}^{2}p_{1-7}^{3}c_{8}^{3}p_{1-7}c_{8}^{6}p_{1-7}c_{8}^{2}p_{1-7}c_{8}^{2}p_{1-7}^{3}c_{8}^{3}p_{1-7}c_{8}^{5}p_{1-7}c_{8}p_{1-7}c_{8}^{5}p_{1-7}c_{8}^{4}p_{1-7}c_{8}p_{1-7}c_{8}p_{1-7}c_{8}p_{1-7}$
$c_{8}^{2}p_{1-7}c_{8}p_{1-7}c_{8}^{6}p_{1-7}c_{8}p_{1-7}c_{8}p_{1-7}c_{8}^{2}p_{1-7}c_{8}^{2}p_{1-5}p_{11-7}c_{8}p_{1-7}c_{8}^{2}p_{1-7}c_{8}p_{1-7}c_{8}p_{1-7}c_{8}p_{1-7}c_{8}^{2}p_{1-7}c_{8}^{4}p_{1-7}c_{8}^{4}p_{1-7}c_{8}p_{1-7}$
$c_{8}^{2}p_{1-7}c_{8}^{4}p_{1-7}^{2}c_{8}p_{1-7}c_{8}p_{1-7}^{6}c_{8}p_{1-7}^{3}c_{8}p_{1-7}^{4}c_{8}p_{1-7}^{6}c_{8}p_{1-7}c_{8}p_{1-7}^{2}c_{8}p_{1-7}^{3}c_{8}p_{1-7}c_{8}p_{1-7}^{5}c_{8}p_{1-7}c_{8}p_{1-7}c_{8}^{2}p_{1-7}c_{8}p_{1-7}^{2}$
$c_{8}p_{1-7}c_{8}^{2}p_{1-7}^{3}c_{8}p_{1-7}^{2}c_{8}p_{1-7}^{2}c_{8}^{12}p_{1-6}p_{11-7}c_{8}^{3}p_{1-6}p_{11-7}c_{8}^{8}p_{1-7}^{3}c_{8}p_{1-7}^{2}c_{8}^{2}p_{1-7}^{2}c_{8}^{2}p_{1-7}c_{8}^{6}p_{1-7}c_{8}p_{1-7}c_{8}p_{1-7}p_{1-4}c_{5}$

### cen14

$p_{5-7}p_{1-2}\mathrm{G14/22}p_{1-2}p_{4-5}p_{7-8}\mathrm{E14/22}p_{1-5}p_{7-5}c_{6}\mathrm{G14/22}c_{2}p_{4-5}\mathrm{G14/22}p_{2-7}p_{1-5}c_{6}^{2}p_{7-5}c_{6}\mathrm{G14/22}p_{1-5}c_{6}^{2}p_{7-5}$
$c_{6}\mathrm{G14/22}c_{2}p_{4-5}\mathrm{G14/22}p_{1-5}c_{6}p_{7-1}p_{4-5}\mathrm{G14/22}p_{1-2}c_{3}^{39}p_{8-2}c_{3}^{13}p_{8-2}c_{3}^{25}p_{8-4}c_{5}^{406}\mathrm{D14/22}c_{5}^{21}p_{4-5}c_{6}^{8}p_{4-5}$
$c_{6}^{2}\mathrm{D14/22}c_{5}^{67}p_{4-5}c_{6}^{4}p_{4-5}c_{6}^{5}p_{4-5}c_{6}^{3}p_{4-5}c_{6}^{19}p_{4-5}c_{6}^{2}p_{4-5}c_{6}^{4}p_{4-5}c_{6}^{4}p_{4-5}c_{6}^{4}p_{4-5}c_{6}^{4}p_{4-5}c_{6}^{6}p_{4-5}$
$c_{6}\mathrm{D14/22}c_{5}^{347}\mathrm{D14/22}c_{5}^{62}\mathrm{D14/22}c_{5}^{4}p_{4-5}c_{6}^{6}p_{4-5}c_{6}^{2}\mathrm{D14/22}c_{5}^{36}p_{4-5}c_{6}^{4}p_{4-5}c_{6}^{6}p_{4-5}c_{6}^{19}p_{4-5}c_{6}^{3}p_{4-5}c_{6}^{4}p_{4-5}$
$c_{6}^{3}p_{4-5}c_{6}^{3}p_{4-5}c_{6}^{4}p_{4-5}c_{6}^{4}p_{4-5}c_{6}^{4}p_{4-5}c_{6}^{4}p_{4-5}c_{6}^{3}p_{4-5}c_{6}^{4}p_{4-5}c_{6}^{3}p_{4-5}c_{6}^{3}p_{4-5}c_{6}^{3}\mathrm{D14/22}c_{5}^{7}p_{4-5}c_{6}^{97}p_{7-5}$
$c_{6}^{409}\mathrm{G14/22}p_{1-7}p_{2-5}\mathrm{G14/22}p_{1-7}c_{8}\mathrm{G14/22B14/22}p_{2-7}p_{1-4}p_{2-7}c_{8}^{18}p_{2-5}\mathrm{G14/22}c_{8}p_{1-5}\mathrm{G14/22}p_{1-7}c_{8}p_{2-5}$
$\mathrm{G14/22}p_{1-2}c_{3}p_{4-5}p_{7-5}c_{6}\mathrm{G14/22}c_{8}^{4}p_{2-5}\mathrm{G14/22}c_{8}p_{1-5}\mathrm{G14/22}p_{1-7}c_{8}p_{2-5}\mathrm{G14/22}p_{1-2}c_{3}p_{4-5}p_{7-5}c_{6}p_{7-}$

$_5$c$_6^9$G14/22p$_{1-2}$G14/22p$_{1-5}$G14/22p$_{1-7}$c$_8^2$p$_{2-5}$G14/22p$_{1-5}$G14/22p$_{1-5}$G14/22p$_{2-3}$p$_{3-5}$c$_6^{71}$G14/22p$_{2-7}$p$_{2-5}$
G14/22p$_{1-7}$c$_8$G14/22B14/22p$_{2-7}$p$_{1-4}$p$_{2-7}$c$_8^5$p$_{2-5}$G14/22c$_8$p$_{1-5}$G14/22p$_{1-7}$c$_8$p$_{2-5}$G14/22p$_{1-2}$c$_3$p$_{4-5}$p$_{7-5}$
c$_6$p$_{7-5}$c$_6^5$G14/22p$_{1-2}$G14/22p$_{1-5}$G14/22p$_{1-7}$c$_8^2$p$_{2-5}$G14/22c$_8$p$_{1-5}$G14/22p$_{1-7}$c$_8$p$_{2-5}$G14/22p$_{1-2}$c$_3$p$_{4-5}$
G14/22c$_8^3$p$_{1-5}$G14/22p$_{2-7}$c$_8^3$p$_{4-5}$c$_6^2$G14/22p$_{1-5}$G14/22p$_{1-5}$G14/22p$_{1-5}$G14/22p$_{1-7}$c$_8$c$_1$p$_{4-2}$c$_3$E14/22

### cen15

p$_{9-10}$H15D15+L15c$_1$p$_{9-10}$H15D15+L15p$_{1-4}$c$_5$p$_{1-4}$c$_5$p$_{1-4}$p$_{1-3}$p$_{5-11}$p$_{3-4}$p$_{1-3}$c$_5$p$_{1-3}$c$_4$p$_{5-10}$p$_{1-3}$p$_{5-3}$p$_{1-4}$c$_5$p$_{1-3}$c$_4$p$_{5-10}$
p$_{1-3}$p$_{5-3}$p$_{1-3}$c$_4$p$_{1-3}$c$_4$p$_{5-3}$p$_{1-3}$c$_4$p$_{1-4}$c$_5$p$_{1-4}$p$_{1-3}$c$_4$p$_{1-4}$c$_5$p$_{1-4}^2$c$_5$p$_{1-4}$c$_5$p$_{1-4}$c$_5$p$_{1-3}$c$_4$p$_{1-3}$p$_{5-3}$p$_{1-3}$c$_4$p$_{1-4}$c$_5$p$_{1-4}$c$_5$p$_{1-4}$
c$_5$p$_{1-4}$p$_{1-3}$c$_4$p$_{1-3}$c$_4$p$_{1-4}$c$_5$p$_{1-4}$c$_5$p$_{1-4}$c$_5$p$_{1-4}^2$c$_5$p$_{1-4}$c$_5$p$_{1-4}$c$_5$p$_{1-4}$c$_5$p$_{1-4}$c$_5$p$_{1-4}$c$_5$p$_{1-4}$c$_5$p$_{1-4}$c$_5$p$_{1-4}$c$_5$p$_{1-4}$c$_5$p$_{1-4}$c$_5$p$_{1-4}$
c$_5$p$_{1-4}$c$_5$p$_{1-4}$c$_5$p$_{1-4}$c$_5$p$_{1-4}$c$_5$p$_{1-4}$c$_5$p$_{1-4}$c$_5$p$_{1-4}$c$_5^2$p$_{1-4}$c$_5$p$_{1-4}$c$_5$p$_{1-4}$c$_5$p$_{1-4}$c$_5$p$_{1-4}$c$_5^2$p$_{1-4}$c$_5$p$_{1-4}$c$_5$p$_{1-4}$c$_5$p$_{1-4}$c$_5$p$_{1-4}$
c$_5$p$_{1-4}$c$_5$p$_{1-4}$c$_5^2$p$_{1-4}$c$_5$p$_{1-8}$p$_{10-4}$p$_{1-4}$c$_5$p$_{1-4}$c$_5^2$p$_{1-4}$c$_5$p$_{1-4}$c$_5$p$_{1-4}$c$_5$p$_{1-4}$c$_5^2$p$_{1-4}$c$_5$p$_{1-4}$c$_5$p$_{1-4}$c$_5$p$_{1-4}$c$_5^2$p$_{1-4}^2$c$_5$p$_{1-4}$c$_5$p$_{1-4}$
c$_5$p$_{1-4}$c$_5$p$_{1-4}$c$_5$p$_{1-4}$c$_5$p$_{1-4}$c$_5$p$_{1-4}$c$_5$p$_{1-4}$c$_5$p$_{1-4}$c$_5$p$_{1-4}$c$_5$p$_{1-4}$c$_5$p$_{1-4}$c$_5$p$_{1-4}$c$_5$p$_{1-4}$c$_5$p$_{1-4}$c$_5$p$_{1-4}$c$_5$p$_{1-4}$c$_5$p$_{1-4}$c$_5$p$_{1-4}$c$_5^3$p$_{1-4}$
c$_5$p$_{1-4}$c$_5$p$_{1-4}$c$_5$p$_{1-4}$c$_5$p$_{1-4}$c$_5$p$_{1-4}$c$_5$p$_{1-4}$c$_5$p$_{1-4}$c$_5$p$_{1-7}$c$_8$D15+L15c$_5$p$_{1-6}$c$_7$p$_{9-4}$A15p$_{5-1}$c$_5$p$_{1-4}$A15p$_{5-1}$c$_5$p$_{1-4}$
A15p$_{5-6}$c$_7$p$_{9-4}$p$_{1-4}$c$_5$p$_{1-4}$c$_5$p$_{1-6}$c$_7$p$_{9-4}$p$_{1-4}$A15p$_{5-6}$c$_7$p$_{9-4}$c$_5$p$_{1-4}$c$_5$p$_{1-6}$p$_{9-4}$c$_5$p$_{1-4}$c$_5$p$_{1-6}$p$_{9-4}$c$_5$p$_{1-6}$p$_{9-4}$c$_5$p$_{1-6}$p$_{9-4}$
c$_5$p$_{1-4}$c$_5^2$p$_{1-4}$c$_5^2$p$_{1-4}$c$_5$p$_{1-6}$p$_{9-4}$c$_5$p$_{1-4}$c$_5$p$_{1-6}$c$_7$p$_{9-4}$A15c$_5$p$_{1-4}$A15c$_5$p$_{1-4}$A15p$_{5-6}$c$_7$p$_{9-4}$p$_{1-4}$c$_5$p$_{1-4}$c$_5$p$_{1-6}$c$_7$p$_{9-4}$p$_{1-4}$
A15p$_{5-6}$c$_7$p$_{9-4}$c$_5$p$_{1-6}$p$_{9-4}$c$_5$p$_{1-6}$c$_7$p$_{9-4}$c$_5$p$_{1-4}$c$_5$p$_{1-6}$p$_{9-4}$c$_5$p$_{1-4}$c$_5$p$_{1-6}$p$_{9-4}$c$_5$p$_{1-6}$p$_{9-4}$c$_5$p$_{1-6}$p$_{9-4}$c$_5$p$_{1-4}$c$_5^2$p$_{1-4}$c$_5^2$p$_{1-4}$
c$_5$p$_{1-6}$p$_{9-4}$c$_5$p$_{1-4}$c$_5$p$_{1-6}$c$_7$p$_{9-4}$A15c$_5$p$_{1-4}$A15c$_5$p$_{1-4}$A15p$_{5-6}$c$_7$p$_{9-4}$p$_{1-4}$c$_5$p$_{1-4}$c$_5$p$_{1-6}$c$_7$p$_{9-4}$p$_{1-4}$A15p$_{5-6}$c$_7$p$_{9-4}$c$_5$p$_{1-6}$
p$_{9-4}$c$_5$p$_{1-4}$c$_5$p$_{1-6}$p$_{9-4}$c$_5$p$_{1-4}$c$_5$p$_{1-4}$c$_5$p$_{1-4}$c$_5$p$_{1-4}^2$c$_5$p$_{1-4}$c$_5$p$_{1-4}$c$_5$p$_{1-4}$c$_5$p$_{1-4}$c$_5$p$_{1-4}$c$_5$p$_{1-4}$c$_5$p$_{1-4}$c$_5$p$_{1-4}$c$_5$p$_{1-4}$c$_5$p$_{1-4}$
c$_5$p$_{1-4}$c$_5$p$_{1-4}$c$_5$p$_{1-4}$c$_5$p$_{1-4}$c$_5$p$_{1-4}$c$_5$p$_{1-4}$c$_5$p$_{1-4}$c$_5$p$_{1-4}$c$_5$p$_{1-4}$c$_5$p$_{1-4}$c$_5$p$_{1-4}$c$_5$p$_{1-4}$c$_5$p$_{1-4}$c$_5$p$_{1-4}$c$_5$p$_{1-4}$c$_5$p$_{1-4}$c$_5$p$_{1-4}$c$_5$p$_{1-4}$
c$_5$p$_{1-4}$c$_5$p$_{1-4}$c$_5$p$_{1-4}$c$_5$p$_{1-4}$c$_5$p$_{1-4}$c$_5$p$_{1-4}$c$_5$p$_{1-4}$c$_5$p$_{1-4}$c$_5$p$_{1-4}$c$_5$p$_{1-4}$c$_5$p$_{1-4}$c$_5$p$_{1-4}$c$_5$p$_{1-4}$c$_5$p$_{1-4}$c$_5^2$p$_{1-4}$c$_5$p$_{1-4}$c$_5$p$_{1-4}$c$_5$p$_{1-4}$
c$_5$p$_{1-4}$c$_5$p$_{1-4}$c$_5$p$_{1-4}$c$_5$p$_{1-4}$c$_5$p$_{1-4}$c$_5$p$_{1-4}$c$_5$p$_{1-4}$c$_5$p$_{1-4}$c$_5$p$_{1-4}$c$_5$p$_{1-4}$c$_5$p$_{1-4}$c$_5$p$_{1-4}$c$_5$p$_{1-4}$c$_5$p$_{1-4}$c$_5$p$_{1-4}$c$_5$p$_{1-4}$c$_5$p$_{1-4}$c$_5$p$_{1-4}$
c$_5$p$_{1-4}$c$_5$p$_{1-4}$c$_5$p$_{1-10}$J15p$_{10-4}$p$_{1-4}$c$_5$p$_{1-4}$c$_5$p$_{1-4}$c$_5$A15c$_2$D15+L15p$_{1-4}$c$_5^2$p$_{1-4}$c$_5$p$_{1-4}$c$_5$p$_{1-4}$c$_5$p$_{1-4}$c$_5$p$_{1-4}$c$_5$p$_{1-4}$
c$_5$p$_{1-4}$c$_5$p$_{1-4}$c$_5$p$_{1-4}$c$_5$p$_{1-4}$c$_5$p$_{1-4}$c$_5$p$_{1-4}$c$_5$p$_{1-4}$c$_5$p$_{1-4}$c$_5$p$_{1-4}$c$_5$p$_{1-4}$c$_5$p$_{1-4}$c$_5$p$_{1-4}$c$_5$p$_{1-4}$c$_5$p$_{1-4}$c$_5$p$_{1-4}$c$_5$p$_{1-4}$c$_5$p$_{1-4}$c$_5$p$_{1-4}$
c$_5$p$_{1-4}$c$_5$p$_{1-4}^2$c$_5$p$_{1-4}$c$_5$p$_{1-4}$c$_5$p$_{1-4}$c$_5$p$_{1-4}$c$_5$p$_{1-4}$c$_5$p$_{1-4}$c$_5$p$_{1-4}$c$_5$p$_{1-4}^2$c$_5$p$_{1-4}$c$_5$p$_{1-4}$c$_5$p$_{1-4}^2$c$_5$p$_{1-4}$c$_5$p$_{1-4}$c$_5$p$_{1-4}$c$_5$p$_{1-4}$
c$_5$p$_{1-4}$c$_5$p$_{1-4}^2$c$_5$p$_{1-4}$c$_5$p$_{1-4}^2$c$_5$p$_{1-4}$c$_5$p$_{1-4}$c$_5$p$_{1-4}^2$c$_5$p$_{1-4}$c$_5$p$_{1-4}$c$_5$p$_{1-4}^2$c$_5$p$_{1-4}$c$_5$p$_{1-4}$c$_5$p$_{1-6}$c$_7$p$_{9-4}$p$_{1-4}$c$_5$p$_{1-4}$c$_5$p$_{1-4}^2$
c$_5$p$_{1-4}$c$_5$p$_{1-4}$c$_5$p$_{1-6}$c$_7$p$_{9-4}$A15p$_{5-6}$c$_7$p$_{9-4}$p$_{1-4}$c$_5^2$p$_{1-4}$c$_5^4$p$_{1-4}$c$_5^2$p$_{1-4}$c$_5$p$_{1-4}$c$_5$p$_{1-4}$c$_5$p$_{1-4}$c$_5^4$p$_{1-4}$c$_5^2$p$_{1-6}$p$_{9-4}$p$_{1-6}$
c$_7^2$p$_{9-4}$p$_{1-4}$c$_5^7$p$_{1-6}$p$_{9-4}$p$_{1-6}$p$_{9-4}$p$_{1-6}$c$_7^2$p$_{9-4}$p$_{1-4}$c$_5^7$p$_{1-6}$p$_{9-4}$p$_{1-6}$p$_{9-4}$p$_{1-4}$c$_5^7$p$_{1-6}$p$_{9-4}$p$_{1-4}$c$_5$p$_{1-3}$c$_4$c$_5^3$p$_{1-9}$
D15+L15p$_{1-4}$c$_5^2$p$_{1-9}$D15+L15p$_{1-4}$c$_5^2$p$_{6-4}$p$_{1-9}$D15+L15p$_{1-7}$p$_{9-10}$p$_{1-3}$p$_{1-4}$p$_{9-4}$p$_{1-4}$p$_{9-3}$p$_{1-4}$p$_{9-10}$p$_{1-4}^2$p$_{9-4}^2$p$_{1-4}$
p$_{9-3}$p$_{1-4}$p$_{1-2}$p$_{1-3}$B15K15p$_{2-4}$p$_{2-3}$p$_{2-4}$J15p$_{2-3}$D15+L15p$_{7-2}$

### cen16

c$_2^2$p$_{2-6}$p$_{8-9}$p$_{4-5}$p$_{7-8}$A1/16/19/5p$_{10-6}$p$_{8-1}$p$_{10-5}$p$_{7-10}$p$_{2-7}$p$_{9-10}$c$_2$p$_{2-6}$c$_8$p$_{8-3}$p$_{7-10}$p$_{2-4}$p$_{8-5}$p$_{7-5}$c$_7^2$p$_{8-6}$p$_{8-9}$
B16E1/16/19/5p$_{7-5}^3$p$_{7-2}$c$_5^9$p$_{5-9}$p$_{1-8}$p$_{1-9}$c$_2^6$p$_{2-9}$p$_{1-8}$p$_{1-9}$c$_2^9$p$_{2-5}$p$_{4-5}$c$_7^9$p$_{7-5}$c$_4^2$p$_{4-5}$c$_4^{10}$p$_{4-5}$c$_4$p$_{4-9}$p$_{2-9}$p$_{2-5}$
c$_4^{19}$c$_6^{45}$p$_{6-9}$c$_4^{23}$p$_{4-6}$A1/16/19/5c$_{10}^5$c$_4^{75}$D1/16/19/5p$_{7-4}$c$_7$p$_{7-4}$c$_7^{22}$p$_{7-4}$c$_7^{22}$p$_{7-3}$c$_6^9$p$_{6-2}$c$_5^{11}$p$_{5-2}$c$_5^{11}$p$_{5-3}$c$_6^5$p$_{6-3}$
c$_8^5$p$_{8-5}$c$_4^{10}$p$_{4-5}$c$_4^{98}$c$_6^5$p$_{6-3}$c$_6^6$p$_{6-3}$c$_8^7$p$_{8-3}$c$_8^2$p$_{8-3}$c$_8$p$_{8-3}$c$_8$p$_{8-3}$c$_8$p$_{8-3}$c$_8^2$p$_{8-3}$c$_8^{13}$p$_{8-3}$c$_8^2$p$_{8-3}$c$_8$p$_{8-3}$c$_8$p$_{8-3}$c$_8$p$_{8-3}$
c$_8^2$p$_{8-3}$c$_8^2$p$_{8-3}$c$_8$p$_{8-3}$c$_8$H16/19A1/16/19/5c$_{10}$p$_{10-3}$c$_6^6$p$_{6-3}$c$_8^7$p$_{8-3}$c$_8^2$p$_{8-3}$c$_8$p$_{8-3}$c$_8$p$_{8-3}$H16/19c$_{10}$p$_{10-3}$c$_8^2$p$_{8-3}$
c$_8^{13}$p$_{8-3}$c$_8^2$p$_{8-3}$c$_8$p$_{8-3}$c$_8$p$_{8-3}$c$_8^2$p$_{8-3}$c$_8^2$p$_{8-3}$c$_8$p$_{8-3}$c$_8$H16/19A1/16/19/5c$_{10}$p$_{10-3}$c$_8$H16/19A1/16/19/5c$_{10}$p$_{10-3}$
c$_8^4$p$_{8-9}$c$_2^{21}$p$_{2-3}$c$_8$p$_{8-3}$c$_8$p$_{8-3}$c$_8$p$_{8-3}$c$_8^3$p$_{8-3}$c$_8$p$_{8-3}$c$_8$p$_{8-3}$c$_8$p$_{8-3}$c$_8$p$_{8-3}$c$_8$p$_{8-3}$c$_8$p$_{8-3}$c$_8$p$_{8-3}$c$_8^2$p$_{8-3}$
c$_8$H16/19A1/16/19/5c$_{10}$p$_{10-3}$c$_8$H16/19A1/16/19/5c$_{10}^3$p$_{10-3}$c$_8^2$p$_{8-9}$c$_2$p$_{2-9}$c$_2^{36}$p$_{2-3}$c$_6^6$p$_{6-3}$c$_6^7$p$_{6-3}$c$_6^2$p$_{6-3}$
c$_6^8$p$_{6-3}$c$_6^5$c$_8^2$p$_{8-3}$c$_6$p$_{8-2}$c$_5$E1/16/19/5c$_8^9$p$_{8-3}$c$_6^3$p$_{6-4}$c$_7^3$p$_{7-3}$c$_6$p$_{6-4}$c$_7^3$p$_{7-3}$c$_6$p$_{6-4}$c$_7^4$p$_{7-3}$c$_6$p$_{6-3}$c$_6$p$_{6-4}$c$_7^6$p$_{7-5}$
c$_4^{14}$c$_6^{41}$p$_{6-9}$c$_2^{52}$p$_{2-3}$c$_6^3$p$_{6-3}$c$_6$p$_{6-3}^2$c$_6^8$p$_{6-3}^2$c$_6^{13}$p$_{6-8}$p$_{5-6}$I16c$_2^2$p$_{2-6}$c$_9^6$p$_{9-5}$p$_{4-6}$p$_{9-10}$c$_5^6$p$_{5-3}$c$_6^{20}$F16/19I16c$_2^2$p$_{2-6}$
I16c$_2$p$_{2-6}$I16p$_{2-6}$I16c$_2^7$p$_{2-6}$I16c$_2^2$p$_{2-6}$I16c$_2$p$_{2-6}$I16c$_2$p$_{2-6}$I16c$_2^3$p$_{2-6}$I16c$_2^2$p$_{2-8}$p$_{1-6}$I16c$_2^3$p$_{2-6}$I16p$_{2-6}$I16p$_{2-6}$
I16p$_{2-6}$I16c$_2^2$p$_{2-6}$I16c$_2^2$p$_{2-6}$I16p$_{2-3}$c$_6^2$F16/19I16c$_2$p$_{2-6}$I16p$_{2-6}$I16p$_{2-3}$c$_6^3$p$_{6-9}$p$_{4-6}$I16p$_{2-3}$c$_6^4$F16/19I16p$_{2-6}$
I16p$_{2-6}$I16p$_{2-6}$I16c$_2$p$_{2-6}$I16p$_{2-4}$c$_7^2$I16p$_{2-6}$I16c$_2$p$_{2-6}$I16p$_{2-6}$I16p$_{2-8}$p$_{1-8}^2$p$_{1-6}$I16p$_{2-8}$c$_1$p$_{1-6}$I16p$_{2-8}$c$_1$p$_{1-6}$I16p$_{2-8}$
c$_1$p$_{1-6}$I16c$_2^2$p$_{2-6}$I16p$_{2-8}$c$_1$p$_{1-6}$I16c$_2$p$_{2-6}$I16c$_2$p$_{2-6}$I16p$_{2-6}$I16p$_{2-8}$p$_{1-8}^2$c$_1$p$_{1-6}$I16c$_2$p$_{2-6}$I16p$_{2-6}$I16c$_2$p$_{2-6}$I16p$_{2-6}$
I16c$_2$p$_{2-6}$I16p$_{2-6}$I16c$_2$p$_{2-6}$I16p$_{2-8}$p$_{1-8}$p$_{1-6}$I16p$_{2-8}$p$_{1-6}$I16p$_{2-8}$c$_1$p$_{1-6}$I16c$_2$p$_{2-6}$I16p$_{2-8}$c$_1^2$p$_{1-6}$I16p$_{2-8}$c$_1$p$_{1-6}$
I16p$_{2-8}$c$_1$p$_{1-6}$I16p$_{2-6}$I16p$_{2-8}$c$_1$p$_{1-9}$c$_2^4$p$_{2-9}$c$_2^9$p$_{2-6}$c$_9^{73}$p$_{9-6}$I16c$_2^2$p$_{2-6}$I16c$_2^4$p$_{2-6}$I16p$_{2-3}$c$_6$p$_{6-10}$I16c$_2^3$p$_{2-6}$I16p$_{2-6}$
I16c$_2^5$p$_{2-6}$I16p$_{2-3}$c$_6^2$F16/19I16c$_2$p$_{2-6}$I16c$_2$p$_{2-6}$I16c$_2^2$p$_{2-6}$I16c$_2^2$p$_{2-6}$I16c$_2$p$_{2-6}$I16c$_2^3$p$_{2-6}$I16p$_{2-}$

$_{3}$c$_{6}$F16/19I16c$_{2}$p$_{2-3}$

### cen17

c1$_{30}^{8}$p$_{30-31}$c1$_{19}^{28}$p$_{19-28}$c1$_{22}^{65}$p$_{22-19}$c1$_{21}^{12}$p$_{21-24}$c1$_{18}^{2}$p$_{18-21}$c1$_{29}$p$_{29-21}$c1$_{29}$p$_{29-21}$c1$_{29}$p$_{29-21}$c1$_{29}$p$_{29-21}$c1$_{29}^{48}$p$_{29-30}$
c1$_{18}^{21}$p$_{18-21}$p$_{12-13}$c$_{15}$p$_{15-13}$P17p$_{2-13}$p$_{15-9}$p$_{11-9}^{2}$c$_{11}^{38}$p$_{11-4}$c$_{8}^{11}$p$_{8-10}$c$_{13}^{7}$p$_{13-16}$c$_{2}^{9}$p$_{2-4}$c$_{8}$p$_{8-4}$c$_{8}^{2}$p$_{8-13}$c$_{15}$p$_{16-14}$
c$_{16}^{3}$p$_{16-13}$p$_{15-9}$p$_{11-13}$c$_{15}$p$_{15-13}$c$_{15}$p$_{15-4}$p$_{8-14}$p$_{16-7}$p$_{11-14}$c$_{16}^{7}$p$_{16-4}$p$_{8-10}$c$_{13}$M17c$_{15}$p$_{15-13}$c$_{15}^{2}$p$_{15-4}$p$_{8-14}$c$_{16}$p$_{16-13}$
c$_{15}$p$_{15-13}$p$_{16-2}$p$_{8-13}$c$_{16}^{7}$p$_{16-14}$c$_{16}^{3}$p$_{16-7}$c$_{9}$p$_{9-4}$c$_{8}$p$_{8-9}$p$_{11-4}$c$_{8}^{2}$p$_{8-9}$p$_{11-4}$c$_{8}^{8}$p$_{8-14}$c$_{16}^{18}$c$_{14}^{13}$AP17+N17c$_{16}^{2}$p$_{16-14}$
c$_{16}^{2}$p$_{16-7}$c$_{9}$p$_{9-4}$c$_{8}$p$_{8-9}$p$_{11-4}$c$_{8}^{6}$p$_{8-14}$c$_{16}^{18}$c$_{14}^{7}$AP17+N17c$_{16}$p$_{16-13}$c$_{16}$p$_{16-13}$c$_{16}^{3}$p$_{16-13}$c$_{15}^{2}$p$_{15-9}$p$_{11-13}$p$_{15-9}$p$_{11-13}$
c$_{15}^{27}$p$_{15-10}$c$_{10}^{4}$J17c$_{10}^{2}$J17c$_{10}^{2}$J17c$_{10}^{41}$p$_{10-14}$c$_{16}^{24}$p$_{16-14}$c$_{16}^{59}$p$_{16-14}$c$_{16}^{103}$P17c$_{2}^{24}$p$_{2-15}$c$_{1}^{32}$p$_{1-5}$c$_{6}^{55}$p$_{6-14}$c$_{7}$p$_{7-14}$
c$_{7}^{2}$p$_{7-14}$c$_{7}^{5}$p$_{7-15}$p$_{1-14}$c$_{7}$p$_{7-14}$c$_{7}$p$_{7-15}$p$_{1-14}$c$_{7}^{3}$p$_{7-14}$c$_{7}^{66}$p$_{7-16}$c$_{16}$P17c$_{16}^{3}$P17c$_{16}^{20}$p$_{16-13}$c$_{16}^{19}$p$_{16-14}$c$_{7}^{17}$p$_{7-8}$
c$_{12}^{3}$p$_{12-13}$c$_{15}^{21}$p$_{15-16}$c$_{16}^{21}$p$_{16-14}$c$_{1}^{13}$p$_{1-14}$c$_{7}^{19}$p$_{7-13}$c$_{15}^{52}$O17c$_{1}^{73}$p$_{1-14}$c$_{16}^{9}$c$_{1}$p$_{1-15}$c$_{1}^{19}$p$_{1-15}$c$_{1}^{17}$p$_{1-14}$c$_{16}^{30}$p$_{1-15}^{3}$p$_{1-7}$
p$_{11-7}^{10}$c$_{11}^{7}$p$_{11-13}$c$_{15}$p$_{15-13}$c$_{15}^{2}$p$_{15-13}$p$_{15-3}$c$_{9}$p$_{9-14}$P17c$_{2}^{2}$p$_{2-13}$p$_{15-13}$c$_{15}$p$_{15-13}$c$_{15}$p$_{15-13}^{2}$c$_{15}$P17c$_{2}$p$_{2-13}$c$_{15}$O17p$_{1-14}$
p$_{16-4}$p$_{11-15}$c$_{15}$p$_{15-13}$O17p$_{15-13}$O17$^{2}$p$_{15-13}^{2}$O17$^{2}$p$_{15-13}$O17$^{2}$p$_{15-13}^{2}$c$_{15}$P17$^{2}$p$_{2-14}$p$_{10-15}$AP17+N17p$_{16-13}$
O17AP17+N17c$_{16}^{4}$p$_{16-13}$c$_{14}^{5}$p$_{15-13}^{2}$O17c$_{14}$p$_{14-15}$p$_{15-13}$p$_{15-1}$p$_{3-13}$p$_{15-13}^{2}$c$_{15}$p$_{16-13}$P17p$_{2-13}$p$_{16-14}$P17p$_{2-13}$
p$_{15-16}$p$_{2-13}$p$_{15-16}$p$_{2-16}^{4}$p$_{2-13}$P17p$_{2-13}$P17p$_{2-16}^{23}$p$_{2-13}$P17p$_{2-16}$p$_{2-13}$P17p$_{2-16}^{21}$p$_{2-13}$P17p$_{2-16}^{2}$p$_{2-14}$P17p$_{2-16}^{3}$
c$_{2}$p$_{2-7}$p$_{3-4}$p$_{12-16}$p$_{2-16}^{2}$c$_{3}$p$_{3-16}$p$_{2-16}^{6}$p$_{2-13}$P17p$_{2-16}^{2}$p$_{2-14}$P17p$_{2-16}^{3}$c$_{2}$p$_{2-7}$p$_{3-4}$p$_{12-16}$p$_{2-16}^{2}$c$_{3}$p$_{3-16}$p$_{2-16}$p$_{2-13}$
c$_{15}$p$_{15-1}$p$_{7-13}$P17p$_{2-16}^{4}$p$_{2-14}$P17p$_{2-16}$p$_{2-15}$p$_{14-16}$p$_{2-16}^{8}$p$_{3-1}$c$_{3}$p$_{3-16}$p$_{2-14}$c$_{16}$p$_{16-13}$c$_{15}^{3}$c$_{16}$p$_{16-14}^{2}$c$_{16}$P17p$_{7-15}$
O17p$_{1-6}$p$_{13-14}$p$_{16-14}$AP17+N17p$_{16-13}$p$_{15-13}^{2}$p$_{15-9}$p$_{11-13}$p$_{15-10}$p$_{2-13}$p$_{15-13}^{2}$p$_{15-5}$M17p$_{16-13}$p$_{15-13}^{3}$p$_{15-7}$p$_{3-4}$p$_{8-13}$
p$_{15-16}$p$_{2-13}$p$_{15-7}$p$_{3-4}$p$_{8-13}$p$_{15-16}$c$_{2}$p$_{2-13}$p$_{15-13}$c$_{15}$p$_{15-16}$p$_{2-16}^{23}$p$_{2-13}$p$_{15-16}$p$_{2-16}^{6}$c$_{2}^{2}$p$_{2-16}^{3}$c$_{2}$p$_{2-16}$p$_{2-6}$p$_{14-16}$p$_{2-16}^{4}$
c$_{2}$p$_{2-13}$p$_{16-13}^{2}$O17c$_{15}^{2}$O17c$_{1}^{3}$p$_{1-14}$p$_{16-13}$c$_{16}$AP17+N17p$_{16-13}$p$_{16-14}$P17p$_{16-14}$P17c$_{16}^{3}$p$_{16-13}$p$_{15-13}$p$_{16-14}$
P17p$_{2-10}$p$_{12-14}$p$_{16-13}$c$_{16}$p$_{16-13}$c$_{15}^{2}$p$_{15-13}^{4}$p$_{15-10}$p$_{12-13}$c$_{16}$p$_{16-13}$p$_{15-6}$c$_{8}$p$_{8-13}$c$_{15}$p$_{15-13}$p$_{15-16}$c$_{2}$p$_{2-13}$p$_{15-13}^{2}$p$_{15-12}$
p$_{2-5}$p$_{8-12}$p$_{15-16}^{2}$p$_{2-5}$p$_{8-13}$p$_{15-16}^{2}$p$_{2-5}$p$_{8-13}$p$_{15-5}$p$_{8-13}$P17p$_{15-3}$

### cen18

I18L18G18.1p$_{4-5}$p$_{7-9}$L18p$_{7-9}$p$_{12-2}$G18.1p$_{4-5}$p$_{7-9}$p$_{12-2}$G18.1p$_{5-9}$p$_{1-2}$G18.1p$_{4-9}$p$_{1-2}$G18.1p$_{4-7}$I18p$_{12-2}$
G18.1p$_{5-9}$p$_{1-2}$G18.1p$_{4-9}$p$_{12-2}$G18.1p$_{4-5}$G18.1I18B18G18.1p$_{5-9}$p$_{12-2}$G18.1p$_{4-5}$p$_{7-9}$p$_{12-2}$G18.1p$_{4-9}$p$_{12-2}$
G18.1p$_{4-9}$p$_{12-2}$G18.1p$_{4-10}$I18p$_{12-2}$G18.1p$_{4-7}$p$_{4-6}$G18.0p$_{8-10}$E18p$_{1-2}$c$_{7}$p$_{7-2}$c$_{7}^{2}$p$_{7-3}$c$_{8}^{47}$p$_{8-2}$c$_{7}^{3}$G18.1c$_{4}^{13}$p$_{4-10}$
c$_{9}^{3}$p$_{9-6}$G18.0c$_{8}^{2}$p$_{8-3}$c$_{8}^{3}$p$_{8-3}$c$_{8}^{6}$J18.1c$_{11}^{3}$p$_{11-7}$J18.1p$_{11-7}$J18.1c$_{11}$p$_{11-9}$p$_{12-2}$p$_{7-9}$p$_{12-2}$G18.1p$_{4-9}$p$_{12-9}^{2}$p$_{12-2}$
G18.1p$_{4-9}$p$_{12-2}$G18.1p$_{4-9}$p$_{12-9}^{2}$p$_{12-2}$G18.1p$_{4-9}$p$_{12-2}$G18.1p$_{4-9}$p$_{12-2}$G18.1p$_{4-9}$p$_{12-2}$G18.1p$_{4-9}$p$_{12-2}$p$_{7-9}$p$_{12-2}$
G18.1p$_{4-9}$p$_{12-2}$p$_{7-9}$I18B18G18.1p$_{4-9}$p$_{12-2}$G18.1J18.1A18p$_{12-2}$p$_{7-9}$p$_{12-2}$G18.1p$_{4-9}$p$_{12-2}$p$_{7-9}$B18G18.1p$_{4-9}$
p$_{12-1}$p$_{1-2}$p$_{7-9}$p$_{12-2}$G18.1p$_{4-9}$p$_{12-1}$p$_{12-2}$G18.1p$_{4-9}$p$_{12-9}$p$_{12-2}$G18.1p$_{4-9}$p$_{12-2}$p$_{7-9}$p$_{12-2}$p$_{7-9}$p$_{12-2}$G18.1p$_{4-9}$p$_{12-2}$p$_{7-9}$
p$_{12-2}$G18.1p$_{4-9}$p$_{12-2}$G18.1p$_{4-9}$p$_{12-2}$G18.1p$_{4-9}$p$_{12-2}$G18.1p$_{4-9}$p$_{12-2}$G18.1p$_{4-9}$p$_{12-2}$p$_{7-9}$p$_{12-2}$G18.1p$_{4-9}$p$_{12-2}$
G18.1p$_{4-9}$p$_{12-2}$G18.1p$_{4-9}$p$_{12-2}$G18.1p$_{4-9}$p$_{12-2}$p$_{7-9}$p$_{12-2}$G18.1p$_{4-9}$p$_{12-2}$G18.1p$_{4-9}$p$_{12-2}$G18.1p$_{4-9}$p$_{12-2}$
G18.1p$_{4-9}$p$_{12-2}$G18.1p$_{4-9}$p$_{12-2}$G18.1p$_{4-9}$p$_{12-2}$G18.1p$_{4-9}$p$_{12-2}$p$_{7-9}$p$_{12-2}$G18.1p$_{4-9}$p$_{12-2}$G18.1p$_{4-9}$p$_{12-2}$p$_{7-9}$
p$_{12-2}$G18.1p$_{4-9}$p$_{12-2}$G18.1p$_{4-9}$p$_{12-2}$G18.1p$_{4-9}$p$_{12-2}$p$_{7-9}$p$_{12-2}$p$_{7-9}$p$_{12-2}$G18.1p$_{4-9}$p$_{12-2}$G18.1p$_{4-9}$p$_{12-2}$p$_{7-9}$p$_{12-2}$
p$_{7-9}$p$_{12-2}$G18.1p$_{4-9}$p$_{12-2}$G18.1p$_{4-9}$p$_{12-2}$G18.1p$_{4-9}$p$_{12-2}$G18.1p$_{4-9}$p$_{12-2}$G18.1p$_{4-9}$p$_{12-2}$G18.1p$_{4-7}$I18p$_{12-2}$p$_{7-9}$
p$_{12-2}$G18.1p$_{4-9}$p$_{12-2}$G18.1p$_{4-9}$p$_{12-2}$p$_{7-9}$p$_{12-2}$G18.1p$_{4-9}$p$_{12-2}$G18.1p$_{4-9}$p$_{12-2}$p$_{7-9}$p$_{12-2}$p$_{7-9}$p$_{12-2}$p$_{7-9}$p$_{12-2}$
G18.1p$_{4-9}$p$_{12-2}$G18.1p$_{4-9}$p$_{12-2}$G18.1p$_{4-9}$p$_{12-2}$G18.1p$_{4-9}$p$_{12-2}$G18.1p$_{12-2}$p$_{7-9}$p$_{12-2}$G18.1p$_{4-9}$p$_{12-2}$p$_{7-9}$
L18p$_{8-9}$p$_{12-9}$p$_{12-2}$p$_{7-9}$p$_{12-2}$p$_{7-9}$L18p$_{8-9}$L18p$_{8-9}$p$_{12-2}$p$_{7-9}$L18p$_{8-9}$p$_{12-9}$I18p$_{12-2}$p$_{7-9}$p$_{12-2}$p$_{7-9}$L18H18/20p$_{12-9}$
I18p$_{12-2}$p$_{7-9}$L18H18/20p$_{12-9}$I18p$_{12-2}$p$_{7-9}$L18p$_{8-9}$p$_{12-9}$I18p$_{12-2}$p$_{7-9}$p$_{12-2}$G18.1p$_{4-6}$G18.0p$_{8-9}$p$_{12-2}$G18.1p$_{4-9}$
p$_{12-9}^{2}$p$_{12-2}$G18.1p$_{4-9}$p$_{12-2}$G18.1p$_{4-9}$p$_{12-2}$p$_{7-9}$p$_{12-2}$p$_{7-9}$L18p$_{8-9}$p$_{12-2}$p$_{7-9}$p$_{8-9}$p$_{12-1}$I18$^{2}$p$_{12-2}$G18.1p$_{4-5}$p$_{7-9}$p$_{12-2}$
p$_{7-9}$L18p$_{8-9}$p$_{12-9}^{2}$p$_{12-2}$p$_{7-9}$L18p$_{8-9}$p$_{12-2}$p$_{7-9}$p$_{8-9}$p$_{12-2}$G18.1p$_{4-7}$L18c$_{8}^{6}$p$_{8-11}$c$_{12}^{2}$p$_{12-2}$G18.1p$_{4-2}$G18.1c$_{4}^{2}$p$_{4-2}$
c$_{9}^{2}$p$_{9-2}$c$_{9}$p$_{9-7}$B18c$_{9}$p$_{9-7}$B18p$_{9-2}$G18.1c$_{12}$p$_{12-2}$G18.1c$_{12}$p$_{12-2}$G18.1p$_{4-2}$G18.1c$_{12}$p$_{12-2}$G18.1c$_{12}$p$_{12-2}$
G18.1c$_{12}$p$_{12-2}$G18.1p$_{4-2}$G18.1c$_{12}$p$_{12-2}$G18.1p$_{4-7}$p$_{4-2}$G18.1p$_{4-2}$G18.1c$_{4}^{9}$p$_{4-9}$p$_{8-9}$c$_{12}^{2}$p$_{12-9}$c$_{2}^{12}$p$_{2-7}$
J18c$_{12}$p$_{12-7}$J18p$_{12-3}$c$_{8}^{3}$c$_{10}$p$_{10-7}$J18c$_{12}^{2}$p$_{12-7}$J18c$_{12}^{2}$p$_{12-7}$J18c$_{12}^{3}$c$_{10}^{2}$p$_{10-2}$c$_{9}^{2}$p$_{9-2}$G18.1p$_{4-7}$
c$_{2}$B18G18.1c$_{4}$p$_{4-2}$c$_{9}^{3}$p$_{9-2}$c$_{9}$p$_{9-2}$c$_{9}^{2}$p$_{9-2}$c$_{9}$p$_{9-2}$G18.1p$_{4-2}$G18.1p$_{4-2}$G18.1p$_{4-2}$c$_{9}$p$_{9-7}$B18p$_{9-7}$B18c$_{9}^{2}$p$_{9-2}$
G18.1p$_{4-2}$G18.1c$_{12}$p$_{12-2}$G18.1c$_{12}$p$_{12-2}$G18.1p$_{4-2}$G18.1c$_{12}$p$_{12-2}$G18.1p$_{4-2}$G18.1p$_{12-2}$G18.1p$_{4-2}$
G18.1p$_{4-2}$G18.1p$_{4-2}$G18.1c$_{12}$p$_{12-2}$G18.1c$_{12}$p$_{12-2}$G18.1p$_{4-2}$G18.1c$_{4}$p$_{4-2}$G18.1p$_{4-2}$G18.1p$_{4-}$

2G18.1c4^2p4-2G18.1p4-2p9-6c3^2G18.1p4-2p9-12c3p9-12c3p9-12c3p9-2G18.1p4-2p9-2c7^3p7-9p12-2c7p7-9p12-2p9-
2G18.1p4-2G18.1p4-2p9-2G18.1p4-2G18.1p4-2c9p9-2G18.1c4p4-2c7^2G18.1J18c12p12-7J18p12-2G18.1p4-
2p7-2G18.1c4p4-2c7p7-2p7-9p4-2p7-9p4-2G18.1p4-11c10p10-2G18.1p4-11c10p10-2G18.1p4-7B18c9p9-11c10p10-
7B18c9p9-7B18c9^2p9-2G18.1p4-2G18.1c4^2p4-7c4p4-2G18.1c4p4-2G18.1c4^2p4-7c4p4-2G18.1p4-7c4^2p4-
2G18.1p4-2G18.1c4^2p4-11c10^3p10-2p9-2G18.1c4^2p4-9p12-2p9-2G18.1c4^23p4-6c3^24p3-9p4-6G18.0c8^6B18p9-
7B18I18p12-2G18.1c4p4-2G18.1p4-2G18.1p4-2G18.1p4-2G18.1p4-2G18.1c4p4-2G18.1c4p4-
6c3^2G18.1c12p12-2G18.1p4-2G18.1c4p4-2G18.1c12p12-2G18.1p4-2G18.1c4p4-2G18.1c12p12-2G18.1p4-
2G18.1c12^2p12-2G18.1c12p12-2G18.1p4-2G18.1c12p12-2G18.1p4-2G18.1c12G18.1p4-2G18.1p4-8p7-
2G18.1c4p4-2G18.1c12p12-2G18.1p4-2G18.1p4-2G18.1c12p12-2G18.1p4-2G18.1p4-2G18.1p4-
2G18.1c12p12-2G18.1p4-2G18.1p4-2G18.1p4-2G18.1c12p12-2G18.1p4-2G18.1c12p12-2G18.1c4p4-
2G18.1c4p4-5H18/20c7p7-2G18.1p4-6c3G18.0p8-5H18/20c7^2c3p3-6c3p3-6G18.0c8^4p8-9p12-2G18.1c4^2p4-
6c3^3G18.1p4-2G18.1p4-2G18.1p4-2G18.1p4-2G18.1p4-2c7p7-2p7-11c10p10-2G18.1p4-2c7p7-2c7p7-
2G18.1c4^2p4-2G18.1p4-11c10p10-11c10p10-2G18.1p4-11c10p10-7B18c9p9-7B18c9p9-2G18.1p4-2G18.1p4-
2G18.1c4p4-2G18.1c4p4-7p12-2G18.1c12^2p12-2G18.1c12^2p12-2G18.1c12p12-2G18.1p4-2G18.1p4-
2G18.1c12^3p12-2G18.1c12p12-2G18.1p4-2G18.1c12^3p12-2G18.1p4-2G18.1c12^3p12-2G18.1c12p12-2G18.1p4-
2G18.1c12^3p12-2G18.1p4-2c9p9-7B18p9-2G18.1c4p4-2G18.1p4-2G18.1p4-6c3p7-2G18.1c4p4-2G18.1p12-
1p9-2G18.1p4-2G18.1c4p4-2G18.1p4-2G18.1p4-2G18.1c4^2p4-2c7p7-2G18.1p4-2G18.1p4-2G18.1c4^2p4-
2G18.1p4-2G18.1c4^2p4-2p7-2^2G18.1p4-2G18.1p4-9p12-2G18.1p4-6p8-2p7-9p12-2p7-2G18.1p4-9p12-2p7-
2G18.1p4-9p12-2p7-2G18.1p4-9p12-2p7-2c7p7-9p12-2p7-2G18.1p4-9p12-2c7p7-2c7p7-2p5-2c7p7-2p9-2G18.1p4-
9p12-2G18.1p4-2G18.1p4-2G18.1p4-7c4^2p4-2G18.1c4^2p4-7p4-2G18.1c4p4-2c7^2p7-2G18.1c4p4-
2c7G18.1J18c12p12-2p7-2G18.1p4-11p10-2G18.1p4-2G18.1p4-2G18.1p4-2G18.1p4-11c10^2p10-11p10-
2G18.1p4-11c10^3p10-11^2c10^4p10-2G18.1p4-11c10^2p10-11p10-2G18.1p4-11c10^3p10-11^2c10^4p10-2G18.1p4-11c10p10-
2G18.1p4-11p10-2c9p9-7B18p9-2G18.1p4-2G18.1p4-2G18.1p4-2G18.1c12p12-7p12-2G18.1c12^4p12-
2G18.1c12p12-2G18.1p4-2G18.1p12-2G18.1c4p4-2G18.1c12p12-2G18.1J18.1c3^2c9p9-2^2c7p7-2G18.1c4p8-
2G18.1p4-2G18.1p4-2G18.1c12p12-2G18.1p4-2G18.1c4p4-2G18.1p4-2G18.1p12-2G18.1p4-2G18.1p4-
2G18.1p12-2G18.1p4-2G18.1p4-2G18.1c4p4-2G18.1p4-2G18.1p4-2G18.1c4^2p4-2G18.1p4-2G18.1p4-
2G18.1c4^2p4-2p7-2G18.1p4-2G18.1p4-2G18.1c4^2p4-2c7^3p7-2c7p7-9p12-2p7-2G18.1p4-2G18.1p4-9p12-2p7-
2G18.1p4-9p12-2p7-2G18.1p4-9c12p12-9p12-2p7-2G18.1p4-9p12-1p7-2c7^3p7-2c7^2p7-2p9-2p7-9p12-2p7-2G18.1c4p4-
2p7-9p12-9c12p12-2G18.1c4p4-2G18.1p4-2G18.1c4p4-2G18.1c4p4-2G18.1p4-2G18.1c4^14p4-7c9^7p9-
2G18.1p4-2G18.1c4^6p4-2G18.1p4-2G18.1p4-2G18.1p4-2G18.1p4-2c7^2p7-2p9-2p7-2G18.1p4-2c7p7-
2G18.1p4-2G18.1p4-11c10^2p10-11c10^3p10-2G18.1p4-11c10p10-2G18.1p4-11p10-2G18.1p4-11c10p10-2G18.1p4-
10p5-2G18.1J18.1p11-2G18.1p4-10p5-2G18.1c4^2p4-2G18.1c12p12-2G18.1p4-2G18.1c12p12-2G18.1c4p4-
2G18.1p4-2G18.1p12-2G18.1p4-2G18.1p4-2G18.1p4-2G18.1p12-2G18.1p4-2G18.1p4-2G18.1p12-
2G18.1p4-2G18.1p4-2G18.1c4p4-2G18.1c12p12-2G18.1p4-2G18.1p4-2G18.1c4p4-2G18.1c12p12-
2G18.1p4-2G18.1p4-2G18.1c4p4-2c7p7-2G18.1c4p4-2G18.1c4^2p4-2c7p7-2G18.1p4-2G18.1c4p4-2c7p7-
2G18.1p4-2G18.1p4-2G18.1c4^7p4-6G18.0c8^3p8-9p12-2c7^3p7-9p12-2p7-2G18.1c4^8p4-2G18.1c4p4-9p12-2c7^2p7-
2p9-2G18.1p4-2G18.1c4p4-2p7-2^2G18.1p4-2p9-2G18.1p4-9p12-2G18.1c4^3p4-7c4p4-2G18.1p4-7c4^2p4-7p4-
2G18.1c4^3p4-2c7^17p7-2G18.1c4^30p4-6c3^8G18.1c4^58p4-2G18.1c4^7p4-2G18.1p4-2G18.1c4^2p4-2G18.1p4-
2G18.1c4^9p4-2G18.1c4^8p4-7p9-2G18.1p4-2G18.1c4p4-2G18.1c4p4-2G18.1p4-2G18.1p4-2G18.1p4-
2G18.1c4p4-2G18.1c4^2p4-2G18.1c4^5p4-2G18.1c4^6p4-2G18.1c4^4p4-2G18.1c4p4-2G18.1c4^9p4-
2G18.1c4^17p4-2G18.1c4^2p4-2G18.1c4^30p4-2G18.1p4-2G18.1c4p4-7c9^2p9-2G18.1c4p4-2G18.1p4-2G18.1p4-
2G18.1p4-2G18.1p4-7p9-2G18.1c4^8p4-2G18.1p4-2G18.1c4p4-2G18.1c4^2p4-2G18.1c4^2p4-2G18.1p4-
2G18.1c4^3p4-2G18.1c4p4-2G18.1c4^2p4-2G18.1c4^30J18.1c11^6p11-2G18.1p4-2G18.1J18.1c11^5p11-
2G18.1p4-2G18.1p4-2G18.1J18.1c11p11-2G18.1p4-7p4-2G18.1p4-2G18.1c4p4-2G18.1p4-2G18.1p4-
2G18.1J18.1c11^2p11-2G18.1p4-7p4-2G18.1p4-2G18.1c4^3p4-7p4-2G18.1c4^2p4-2G18.1c4p4-2G18.1p4-
2G18.1c4^10p4-6c3^5G18.1p4-6c3G18.1c4p4-6c3^2G18.1c4^5p4-2G18.1c4^3p4-2G18.1c4^30J18.1c11^5p11-
2G18.1p4-2G18.1J18.1c11^5p11-2G18.1p4-2G18.1p4-2G18.1J18.1c11p11-2G18.1p4-2G18.1J18.1c11p11-
2G18.1J18.1c11p11-2G18.1J18.1c11p11-2G18.1c4^12J18.1c11^5p11-2G18.1p4-2G18.1J18.1c11^5p11-
2G18.1p4-2G18.1p4-2G18.1J18.1c11p11-2G18.1p4-2G18.1J18.1c11p11-2G18.1J18.1c11p11-
2G18.1J18.1c11p11-2G18.1p4-7p4-2G18.1c4^2p4-2G18.1p4-2G18.1p4-2G18.1J18.1c11^2p11-2G18.1p4-
7c4p4-2G18.1c4^3p4-7p4-2G18.1c4^4p4-2G18.1p4-2G18.1c4^7p4-6c3^6G18.1p4-6c3G18.1c4p4-6c3G18.1c4p4-

$_{7}p_{4-2}$G18.1$p_{4-7}$J18$c_{5}p_{5-6}c_{3}$G18.1$p_{4-2}$G18.1$p_{4-2}$G18.1$p_{4-2}$G18.1$p_{4-2}$G18.1$c_{4}p_{4-2}$G18.1$p_{4-2}$G18.1$p_{4-2}$
G18.1$p_{4-2}$G18.1$p_{4-7}$J18$p_{5-2}$G18.1$p_{4-7}$J18$p_{5-2}$G18.1$p_{4-7}p_{4-2}$G18.1$c_{4}p_{4-2}$G18.1$p_{4-2}$G18.1$c_{4}^{3}p_{4-2}$
G18.1$p_{4-2}$G18.1$p_{4-2}$G18.1$p_{4-2}$G18.1$c_{4}^{2}p_{4-2}$G18.1$p_{4-2}$G18.1$p_{4-7}$J18$p_{5-2}$G18.1$p_{4-7}$J18$p_{5-2}$G18.1$p_{4-7}p_{4-2}$
G18.1$p_{4-2}$G18.1$p_{4-2}$G18.1$c_{4}p_{4-2}$G18.1$c_{4}^{4}p_{4-2}$G18.1$p_{4-2}$G18.1$p_{4-7}c_{4}p_{4-2}$G18.1$c_{4}p_{4-2}$G18.1$p_{4-2}$
G18.1$c_{4}p_{4-2}$G18.1$c_{4}^{3}p_{4-2}$G18.1$c_{4}p_{4-2}$G18.1$p_{4-2}$G18.1$c_{4}^{5}p_{4-7}p_{4-2}$G18.1$c_{4}^{2}p_{4-7}c_{4}p_{4-7}c_{4}^{3}p_{4-2}$G18.1$c_{4}p_{4-2}$
G18.1$c_{4}^{3}p_{4-2}$G18.1$c_{4}p_{4-2}$G18.1$c_{4}^{4}p_{4-2}$G18.1$c_{4}^{2}p_{4-7}p_{4-2}$G18.1$c_{4}^{6}p_{4-2}$G18.1$c_{4}p_{4-2}$G18.1$c_{4}p_{4-2}$
G18.1$c_{4}p_{4-2}$G18.1$c_{4}p_{4-2}$G18.1$c_{4}p_{4-2}$G18.1$c_{4}^{33}p_{4-2}$G18.1$p_{4-7}p_{4-2}$G18.1$p_{4-7}c_{4}^{2}p_{4-2}$G18.1$p_{4-2}$
G18.1$c_{4}p_{4-2}$G18.1$p_{4-2}$G18.1$c_{4}^{2}p_{4-2}$G18.1$p_{4-2}$G18.1$p_{4-2}$G18.1$c_{4}^{7}p_{4-2}$G18.1$p_{4-7}p_{4-2}$G18.1$p_{4-2}$
G18.1$p_{4-2}$G18.1$p_{4-2}$G18.1$p_{4-2}$G18.1$c_{4}p_{4-2}$G18.1$p_{4-2}$G18.1$p_{4-2}$G18.1$c_{4}p_{4-2}$G18.1$p_{4-2}$G18.1$c_{4}p_{4-2}$
G18.1$c_{4}p_{4-2}$G18.1$p_{4-9}p_{12-2}$G18.1$p_{4-2}$G18.1$c_{4}p_{4-2}$G18.1$p_{4-2}$G18.1$p_{4-2}$G18.1$p_{4-2}$G18.1$c_{4}p_{4-2}$G18.1$p_{4-2}$
G18.1$p_{4-2}$G18.1$p_{4-2}$G18.1$c_{4}^{2}p_{4-2}$G18.1$c_{4}p_{4-2}$G18.1$p_{4-9}p_{12-2}$G18.1$p_{4-2}$G18.1$c_{4}p_{4-2}$G18.1$p_{4-9}p_{12-2}$
G18.1$p_{4-2}$G18.1$c_{4}p_{4-2}$G18.1$c_{4}p_{4-7}p_{4-2}$G18.1$c_{4}p_{4-7}p_{4-2}$G18.1$p_{4-2}$G18.1$p_{4-2}$G18.1$p_{4-2}$G18.1$p_{4-2}$
G18.1$p_{4-2}$G18.1$p_{4-2}$G18.1$p_{4-2}$G18.1$p_{4-1}p_{12-2}$G18.1$p_{4-1}p_{12-2}$G18.1$p_{4-1}p_{12-1}p_{12-2}$G18.1$p_{4-1}p_{12-2}$
G18.1$c_{4}p_{4-2}$G18.1$c_{4}^{2}p_{4-7}^{2}p_{10-2}p_{7-2}c_{7}p_{7-2}$G18.1$p_{10-2}p_{7-2}c_{7}^{2}$G18.1$c_{4}^{5}p_{4-7}c_{9}p_{9-2}$G18.1$p_{4-2}$G18.1$c_{4}p_{4-2}$
G18.1$p_{4-2}$G18.1$c_{4}^{2}p_{4-2}$G18.1$p_{4-2}$G18.1$p_{4-2}$G18.1$c_{4}^{8}p_{4-2}$G18.1$p_{4-2}p_{7-2}^{3}$G18.1$c_{4}^{2}p_{4-7}c_{4}^{2}p_{4-2}p_{7-2}^{2}$
$c_{7}^{3}$G18.1$c_{4}^{5}p_{4-2}$G18.1$c_{4}p_{4-2}$G18.1$c_{4}^{2}p_{4-2}$G18.1$c_{4}^{2}p_{4-6}c_{3}$G18.1$p_{4-6}c_{3}$G18.1$p_{4-10}p_{5-6}c_{3}$G18.1$p_{4-7}p_{4-2}$
G18.1$p_{4-7}p_{4-2}$G18.1$c_{4}p_{4-2}$G18.1$p_{4-6}c_{3}$G18.1$p_{4-2}$G18.1$c_{4}^{6}p_{4-6}$G18.0$c_{8}p_{8-2}$G18.1$c_{4}p_{4-6}$G18.0$p_{8-2}$
G18.1$p_{4-2}$G18.1$p_{4-10}p_{5-6}c_{3}$G18.1$p_{4-6}c_{3}$G18.1$p_{4-6}c_{3}$G18.1$p_{4-7}p_{4-2}$G18.1$c_{4}^{2}p_{4-6}$G18.0$c_{8}p_{8-6}$
G18.0$c_{8}^{2}p_{8-2}$G18.1$c_{4}p_{4-2}$G18.1$p_{4-7}c_{4}p_{4-2}$G18.1$p_{4-2}$G18.1$p_{4-7}c_{4}^{2}p_{4-2}$G18.1$p_{4-7}c_{4}^{4}p_{4-2}$G18.1$p_{4-7}p_{4-2}$
G18.1$p_{4-2}$G18.1$c_{4}^{17}p_{4-2}$G18.1$p_{4-2}$G18.1$c_{4}^{4}p_{4-2}c_{7}^{3}p_{7-2}c_{7}^{2}p_{7-2}p_{7-10}c_{9}^{2}p_{9-2}p_{7-10}c_{9}^{2}p_{9-2}p_{7-10}c_{9}^{2}p_{9-2}p_{7-10}$
$c_{9}^{2}p_{9-2}p_{7-10}p_{9-2}p_{7-10}c_{9}p_{9-10}p_{9-11}$J18.1$c_{1}p_{1-2}p_{7-2}^{3}p_{7-10}p_{12-2}p_{7-2}$G18.1$p_{10-2}p_{7-2}^{9}c_{7}p_{7-4}$A18$p_{5-11}$J18.1$c_{1}^{2}p_{1-2}$
$p_{7-8}p_{10-11}$J18.1$p_{1-2}p_{7-2}^{7}p_{7-3}p_{8-2}p_{7-11}p_{10-11}p_{10-2}p_{7-2}p_{7-11}$J18$p_{9-2}p_{7-2}^{3}p_{7-11}p_{10-2}p_{7-11}p_{10-3}c_{8}^{2}c_{10}p_{10-2}$
$c_{7}^{2}$G18.1$p_{10-2}p_{7-2}c_{7}^{2}p_{7-2}p_{7-1}p_{6-11}p_{10-2}p_{7-1}p_{6-11}p_{10-2}p_{7-2}^{2}c_{7}^{2}p_{7-2}^{4}c_{7}^{2}p_{7-3}p_{8-2}p_{7-2}^{2}c_{7}^{2}p_{7-2}c_{7}p_{7-10}p_{5-6}$G18.0$p_{10-2}$
$c_{7}$G18.1$p_{10-2}$G18.1$p_{10-7}p_{10-2}p_{7-2}c_{7}^{2}$G18.1$p_{10-2}$G18.1$p_{10-7}p_{10-2}p_{7-2}c_{7}^{2}$G18.1$p_{10-2}p_{7-2}^{9}p_{7-11}p_{10-2}$
$c_{7}$G18.1$p_{10-2}p_{7-2}^{2}p_{7-11}p_{10-2}c_{7}$G18.1$p_{10-2}p_{7-2}^{5}p_{7-11}p_{10-11}p_{10-2}p_{7-2}p_{7-11}$J18$p_{9-2}p_{7-2}^{3}p_{7-11}p_{10-2}p_{7-2}p_{7-1}p_{7-11}p_{10-2}$
$p_{7-2}p_{7-11}$J18$p_{9-11}$J18.1$p_{1-2}$G18.1J18.1$p_{1-2}p_{7-2}^{2}c_{7}^{5}p_{7-2}c_{7}p_{7-2}$G18.1$p_{4-7}p_{10-2}p_{7-2}c_{7}^{2}p_{7-2}^{4}p_{7-1}p_{6-2}c_{7}p_{7-2}p_{7-1}$
$p_{6-11}p_{10-2}c_{7}p_{7-2}c_{7}^{2}$G18.1$p_{10-2}p_{7-2}^{4}c_{7}^{2}p_{7-2}^{4}c_{7}^{2}p_{7-2}c_{7}p_{7-10}p_{5-6}$G18.0J18I18$p_{12-2}c_{7}$G18.1$p_{10-2}c_{7}$G18.1$p_{10-2}$
$p_{7-2}^{3}c_{7}$G18.1$p_{10-2}p_{7-2}^{3}c_{7}$G18.1$p_{10-2}c_{7}$G18.1$p_{10-2}p_{7-2}^{14}p_{7-11}$B18$p_{7-2}^{45}p_{7-11}p_{8-2}p_{7-2}^{12}p_{7-8}p_{11-2}p_{7-2}^{2}p_{7-9}p_{12-2}$
$p_{7-2}^{7}c_{7}^{6}p_{7-2}^{72}p_{7-11}p_{8-2}p_{7-2}^{4}p_{7-9}p_{12-2}p_{7-2}^{13}p_{7-11}p_{10-2}p_{7-9}p_{12-2}p_{7-2}^{6}c_{7}^{5}$G18.1$c_{10}p_{10-7}c_{10}^{3}p_{10-2}p_{7-2}$G18.1$p_{4-11}$
J18.1$c_{1}^{2}p_{1-2}p_{7-2}$G18.1$p_{4-10}p_{9-10}p_{9-11}$J18.1$p_{1-2}c_{7}p_{7-10}p_{1-2}$G18.1J18.1$p_{1-2}c_{7}p_{7-10}p_{1-2}$G18.1J18.1$p_{1-2}p_{7-3}$
$p_{10-2}$G18.1$p_{4-10}p_{9-11}$J18.1$p_{1-9}$J18.1$c_{1}p_{1-2}p_{7-10}p_{9-2}$G18.1$p_{4-10}p_{1-2}$G18.1J18.1$p_{1-2}p_{7-3}$J18$p_{1-2}p_{7-3}$J18$p_{1-2}$
$p_{5-11}p_{10-11}$J18.1$p_{1-2}$G18.1J18.1$p_{1-2}p_{7-2}^{4}p_{7-11}p_{10-2}p_{7-1}p_{8-2}p_{7-2}^{2}p_{7-11}p_{10-2}p_{7-2}^{10}p_{7-11}$J18$p_{9-11}$J18.1$p_{1-2}$
G18.1J18.1$p_{1-2}p_{7-2}^{3}p_{7-11}p_{10-2}p_{7-2}^{8}p_{7-3}p_{8-2}p_{7-2}$G18.1J18.1$p_{1-3}p_{8-2}p_{7-2}^{3}p_{7-3}p_{8-2}p_{7-2}^{2}$G18.1J18.1$p_{1-3}p_{8-2}$
$p_{7-2}^{17}p_{7-11}p_{10-2}p_{7-10}p_{9-2}p_{7-2}^{2}p_{7-10}p_{10-2}$G18.1J18.1$p_{1-2}p_{7-2}^{2}p_{7-8}p_{1-2}p_{7-2}p_{7-11}p_{10-2}c_{5}^{2}p_{5-2}c_{7}^{2}$G18.1$c_{9}p_{9-7}$
$c_{9}^{2}p_{9-2}p_{7-2}^{3}p_{7-11}p_{10-2}p_{7-2}^{10}c_{7}p_{7-10}c_{9}p_{9-2}p_{7-10}p_{9-11}$J18.1$p_{1-10}c_{9}p_{9-2}p_{7-10}p_{9-11}$J18.1$p_{1-2}p_{7-2}^{6}p_{7-9}p_{12-2}p_{7-2}^{5}p_{7-9}$
$p_{12-2}p_{7-2}^{4}p_{7-10}p_{9-10}$J18.1$p_{1-2}p_{7-2}^{4}p_{7-11}p_{10-2}p_{7-2}^{17}p_{7-11}p_{8-11}^{2}p_{10-2}p_{7-11}p_{8-11}p_{10-2}p_{7-2}^{5}p_{7-11}c_{10}p_{10-2}$G18.1$c_{4}p_{4-7}$
$p_{10-2}p_{7-2}^{2}c_{7}$G18.1$p_{10-2}p_{7-2}^{2}c_{7}$G18.1$p_{10-2}p_{7-2}c_{7}$G18.1$p_{10-2}p_{7-2}^{3}c_{7}$G18.0$p_{10-2}p_{7-2}^{2}c_{7}$G18.1$p_{10-2}p_{7-2}^{4}$
$c_{7}$G18.1J18$p_{9-2}p_{7-2}^{4}p_{7-9}p_{12-2}p_{7-2}^{4}p_{7-9}p_{12-2}p_{7-2}^{5}p_{7-9}p_{12-2}p_{7-2}^{17}p_{7-10}$I18$p_{12-2}p_{7-2}^{15}p_{7-9}p_{12-2}p_{7-2}^{8}p_{7-9}p_{12-2}p_{7-2}$
$p_{7-8}p_{11-2}p_{7-2}^{8}p_{7-9}p_{12-2}p_{7-2}^{33}p_{7-9}p_{12-2}p_{7-2}^{61}p_{7-11}p_{10-2}p_{7-8}p_{10-2}p_{7-2}^{10}p_{7-11}p_{10-2}p_{7-2}^{8}p_{7-9}c_{12}$L18

### cen19

AD1/16/19/5+E1/19/5$c_{1}^{2}$AD1/16/19/5+E1/19/5$^{2}c_{2}c_{1}^{5}$AD1/16/19/5+E1/19/5$c_{2}^{3}$AD1/16/19/5+E1/19
/5$c_{1}^{4}c_{2}^{5}$AD1/16/19/5+E1/19/5$c_{1}^{4}c_{2}^{2}$G1/19/5+F19+R19$c_{1}$AD1/16/19/5+E1/19/5$c_{1}^{3}c_{2}c_{1}^{3}$AD1/16/19/
5+E1/19/5$c_{1}$AD1/16/19/5+E1/19/5$c_{1}^{4}$AD1/16/19/5+E1/19/5$c_{1}^{5}$AD1/16/19/5+E1/19/5$^{2}c_{1}^{4}$AD1/16/1
9/5+E1/19/5$^{2}c_{2}c_{1}^{2}$AD1/16/19/5+E1/19/5$c_{2}^{5}c_{1}^{2}$AD1/16/19/5+E1/19/5$c_{1}$AD1/16/19/5+E1/19/5$c_{2}c_{1}^{2}$
AD1/16/19/5+E1/19/5$c_{2}^{5}c_{1}^{2}$AD1/16/19/5+E1/19/5$c_{1}$AD1/16/19/5+E1/19/5$c_{2}^{6}$AD1/16/19/5+E1/19/
5$c_{1}^{2}$AD1/16/19/5+E1/19/5$c_{2}^{4}c_{1}$AD1/16/19/5+E1/19/5$c_{2}c_{1}^{3}$AD1/16/19/5+E1/19/5$c_{1}$AD1/16/19/5+E
1/19/5$c_{2}c_{1}c_{2}$AD1/16/19/5+E1/19/5$c_{2}^{19}$AD1/16/19/5+E1/19/5$c_{2}^{9}c_{1}$AD1/16/19/5+E1/19/5$c_{2}^{2}$G1/19/5
+F19+R19$c_{2}^{7}$G1/19/5+F19+R19$c_{2}^{4}c_{1}c_{2}^{4}c_{1}c_{2}^{5}$G1/19/5+F19+R19$c_{2}^{3}$G1/19/5+F19+R19$c_{2}^{2}$G1/19/5
+F19+R19$c_{2}^{7}$G1/19/5+F19+R19$c_{2}^{6}$G1/19/5+F19+R19$c_{2}^{16}$G1/19/5+F19+R19$c_{2}^{6}$G1/19/5+F19+R1

9$c_{2}^{2}$G1/19/5+F19+R19$c_{2}$G1/19/5+F19+R19$c_{2}^{15}$G1/19/5+F19+R19$c_{2}^{3}$G1/19/5+F19+R19$c_{2}^{13}$G1/1
9/5+F19+R19$c_{2}^{4}$G1/19/5+F19+R19$c_{2}^{3}$G1/19/5+F19+R19$c_{2}^{6}$G1/19/5+F19+R19$c_{2}^{5}$AD1/16/19/5+
E1/19/$5^{2}$$c_{2}^{7}$AD1/16/19/5+E1/19/5$c_{2}c_{1}^{7}c_{2}^{4}c_{1}^{5}$AD1/16/19/5+E1/19/5$c_{2}^{4}c_{1}^{5}$AD1/16/19/5+E1/19/5$c_{2}^{4}c$
$_{1}^{4}$AD1/16/19/5+E1/19/5$c_{1}^{5}$AD1/16/19/5+E1/19/5$c_{2}^{10}$G1/19/5+F19+R19$c_{2}^{7}c_{1}^{5}$AD1/16/19/5+E1/19/
5G1/19/5+F19+R19$c_{2}^{7}c_{1}^{8}$AD1/16/19/5+E1/19/5$c_{1}^{5}$AD1/16/19/5+E1/19/5$c_{2}^{16}c_{1}^{8}$AD1/16/19/5+E1/
19/5$c_{1}^{5}$AD1/16/19/5+E1/19/5$c_{2}^{9}c_{1}^{2}$AD1/16/19/5+E1/19/5$c_{2}^{20}c_{1}^{10}$AD1/16/19/5+E1/19/5$c_{1}^{7}$AD1/16/
19/5+E1/19/5G1/19/5+F19+R19$c_{2}^{7}c_{1}^{3}$AD1/16/19/5+E1/19/5$c_{2}^{5}$G1/19/5+F19+R19$c_{2}^{9}c_{1}^{8}$AD1/16/1
9/5+E1/19/5$c_{1}^{5}$AD1/16/19/5+E1/19/5$c_{2}^{18}$AD1/16/19/5+E1/19/5$c_{2}^{17}c_{1}^{10}$AD1/16/19/5+E1/19/5$c_{1}^{7}$A
D1/16/19/5+E1/19/5$c_{1}^{3}$AD1/16/19/5+E1/19/5$c_{2}^{25}c_{1}^{10}$AD1/16/19/5+E1/19/5$c_{1}^{7}$AD1/16/19/5+E1/19
/5$c_{1}^{3}$AD1/16/19/5+E1/19/5$c_{2}^{2}c_{1}^{3}$AD1/16/19/5+E1/19/5$c_{2}^{25}c_{1}^{10}$AD1/16/19/5+E1/19/5$c_{1}^{7}$AD1/16/19
/5+E1/19/5$c_{1}^{3}$AD1/16/19/5+E1/19/5$c_{2}^{14}c_{1}^{3}$AD1/16/19/5+E1/19/5$c_{2}^{4}c_{1}^{6}$AD1/16/19/5+E1/19/5$c_{2}^{13}$A
D1/16/19/5+E1/19/5$c_{2}$AD1/16/19/5+E1/19/5$c_{1}c_{2}^{5}$G1/19/5+F19+R19$c_{2}^{13}c_{1}$AD1/16/19/5+E1/19/5$c$
$_{1}^{2}$AD1/16/19/5+E1/19/5$c_{1}^{4}$AD1/16/19/5+E1/19/5$c_{1}^{8}$AD1/16/19/5+E1/19/5$c_{2}^{14}c_{1}^{6}c_{2}^{11}$AD1/16/19/5+
E1/19/5$c_{2}^{39}c_{1}^{5}$AD1/16/19/5+E1/19/$5^{2}$$c_{2}^{9}$AD1/16/19/5+E1/19/5$c_{2}^{7}c_{1}^{11}$AD1/16/19/5+E1/19/$5^{2}$$c_{2}^{7}c_{1}^{34}$
AD1/16/19/5+E1/19/5$c_{1}^{20}$AD1/16/19/5+E1/19/5$c_{1}^{7}$AD1/16/19/5+E1/19/5$c_{2}^{5}c_{1}^{5}c_{2}^{6}$AD1/16/19/5+E1
/19/5$c_{1}^{4}$AD1/16/19/5+E1/19/5$c_{2}^{7}$AD1/16/19/5+E1/19/5$c_{2}^{7}c_{1}^{5}$AD1/16/19/5+E1/19/5$c_{2}^{4}c_{1}^{2}$AD1/16/1
9/5+E1/19/5$c_{1}^{5}$AD1/16/19/5+E1/19/5$c_{1}^{5}$AD1/16/19/5+E1/19/5$c_{2}^{4}c_{1}^{5}$AD1/16/19/5+E1/19/$5^{2}$$c_{1}^{11}$AD
1/16/19/5+E1/19/5$c_{1}^{7}$AD1/16/19/5+E1/19/5$c_{1}^{6}$AD1/16/19/5+E1/19/5$c_{2}^{7}c_{1}^{5}$AD1/16/19/5+E1/19/5$c$
$_{1}^{5}$AD1/16/19/5+E1/19/5$c_{1}^{5}$AD1/16/19/5+E1/19/5$c_{1}^{6}c_{2}^{9}c_{1}$AD1/16/19/5+E1/19/5$c_{2}^{7}c_{1}^{6}$AD1/16/19/5+
E1/19/5$c_{1}^{6}c_{2}^{4}c_{1}^{4}c_{2}^{4}c_{1}^{11}c_{2}^{4}c_{1}^{6}$AD1/16/19/5+E1/19/5$c_{1}^{2}c_{2}^{30}c_{1}^{9}c_{2}^{39}$G1/19/5+F19+R19$c_{2}^{11}$AD1/16/19/
5+E1/19/5$c_{2}^{4}c_{1}^{10}$AD1/16/19/5+E1/19/5$c_{2}^{13}$AD1/16/19/5+E1/19/5$c_{2}^{39}c_{1}^{22}$AD1/16/19/5+E1/19/5$c_{1}$A
D1/16/19/5+E1/19/5$c_{1}^{22}$AD1/16/19/5+E1/19/5$c_{1}$AD1/16/19/5+E1/19/5$c_{1}^{5}$AD1/16/19/5+E1/19/5$c_{1}$
AD1/16/19/5+E1/19/5$c_{1}^{19}$AD1/16/19/5+E1/19/5$c_{1}$AD1/16/19/5+E1/19/5$c_{1}^{7}$AD1/16/19/5+E1/19/5$c$
$_{1}$AD1/16/19/5+E1/19/5$c_{1}^{7}$AD1/16/19/5+E1/19/5$c_{1}$AD1/16/19/5+E1/19/5$c_{1}^{37}$AD1/16/19/5+E1/19/5
$c_{1}$AD1/16/19/5+E1/19/5$c_{1}^{32}$AD1/16/19/5+E1/19/5$c_{1}$AD1/16/19/5+E1/19/5$c_{1}^{52}$AD1/16/19/5+E1/19
/5$c_{1}$AD1/16/19/5+E1/19/5$c_{1}^{16}$AD1/16/19/5+E1/19/5$c_{1}$AD1/16/19/5+E1/19/5$c_{1}^{808}$AD1/16/19/5+E1/
19/5$c_{1}^{477}$AD1/16/19/5+E1/19/5$c_{1}^{15}$AD1/16/19/5+E1/19/5$c_{1}^{44}$AD1/16/19/5+E1/19/5$c_{1}^{25}$AD1/16/19/
5+E1/19/5$c_{1}^{65}$AD1/16/19/5+E1/19/5$c_{1}^{2}$AD1/16/19/5+E1/19/5$c_{1}^{19}$AD1/16/19/5+E1/19/5$c_{1}^{22}$AD1/1
6/19/5+E1/19/5$c_{1}^{14}$AD1/16/19/5+E1/19/5$c_{1}^{144}$AD1/16/19/5+E1/19/5$c_{1}^{133}$AD1/16/19/5+E1/19/5$c_{1}^{15}$
AD1/16/19/5+E1/19/5$c_{1}^{45}$AD1/16/19/5+E1/19/5$c_{1}^{25}$AD1/16/19/5+E1/19/5$c_{1}^{65}$AD1/16/19/5+E1/19/
5$c_{1}^{2}$AD1/16/19/5+E1/19/5$c_{1}^{25}$AD1/16/19/5+E1/19/5$c_{1}^{28}$AD1/16/19/5+E1/19/5$c_{1}^{14}$AD1/16/19/5+E
1/19/5$c_{1}^{2}$AD1/16/19/5+E1/19/5$c_{1}^{56}$AD1/16/19/5+E1/19/5$c_{1}^{7}$AD1/16/19/5+E1/19/5$c_{1}^{30}$AD1/16/19/
5+E1/19/5$c_{1}^{54}$AD1/16/19/5+E1/19/5$c_{1}^{9}$AD1/16/19/5+E1/19/5$c_{1}^{31}$AD1/16/19/5+E1/19/5$c_{1}^{11}$AD1/1
6/19/5+E1/19/5$c_{1}^{2}$AD1/16/19/5+E1/19/5$c_{1}^{20}$AD1/16/19/5+E1/19/5$c_{1}^{15}$AD1/16/19/5+E1/19/5$c_{1}^{17}$A
D1/16/19/5+E1/19/5$c_{1}^{27}$AD1/16/19/5+E1/19/5$c_{1}^{43}$AD1/16/19/5+E1/19/5$c_{1}^{2}$AD1/16/19/5+E1/19/5$c$
$_{1}^{28}$AD1/16/19/5+E1/19/5$c_{1}^{75}$AD1/16/19/5+E1/19/5$c_{1}^{12}$AD1/16/19/5+E1/19/5$c_{1}^{133}$AD1/16/19/5+E1/
19/5$c_{1}^{20}$AD1/16/19/5+E1/19/5$c_{1}^{38}$AD1/16/19/5+E1/19/5$c_{1}^{12}$AD1/16/19/5+E1/19/5$c_{1}^{6}$AD1/16/19/5
+E1/19/5$c_{1}^{5}$AD1/16/19/5+E1/19/5$c_{1}^{6}$AD1/16/19/5+E1/19/5$c_{1}^{5}$AD1/16/19/5+E1/19/5$c_{1}^{10}$AD1/16/1
9/5+E1/19/5$c_{1}^{34}$AD1/16/19/5+E1/19/5$c_{1}^{5}$AD1/16/19/5+E1/19/5$c_{1}^{4}$AD1/16/19/5+E1/19/5$c_{1}^{5}$AD1/1
6/19/5+E1/19/5$c_{1}^{4}$AD1/16/19/5+E1/19/5$c_{1}^{11}$AD1/16/19/5+E1/19/5$c_{1}^{13}$AD1/16/19/5+E1/19/5$c_{1}^{10}$A
D1/16/19/5+E1/19/5$c_{1}^{2}$AD1/16/19/5+E1/19/5$c_{1}^{8}$AD1/16/19/5+E1/19/5$c_{1}$AD1/16/19/5+E1/19/5$c_{1}^{12}$
$^{9}$AD1/16/19/5+E1/19/5$c_{1}^{4}$AD1/16/19/5+E1/19/5$c_{1}^{238}$AD1/16/19/5+E1/19/5$c_{1}^{14}c_{2}^{4}c_{1}^{10}$AD1/16/19/5
+E1/19/5$c_{1}^{78}$AD1/16/19/5+E1/19/5$c_{1}^{25}$AD1/16/19/5+E1/19/5$c_{1}^{38}$AD1/16/19/5+E1/19/5$c_{1}^{3}$AD1/16/
19/5+E1/19/5$c_{1}^{172}c_{2}^{24}c_{1}^{500}$AD1/16/19/5+E1/19/5$c_{1}^{289}$AD1/16/19/5+E1/19/5$c_{1}^{6}$AD1/16/19/5+E1/19
/5$c_{1}^{753}$AD1/16/19/5+E1/19/5$c_{2}^{66}c_{1}^{112}$AD1/16/19/5+E1/19/5$c_{1}^{8}$AD1/16/19/5+E1/19/5$c_{1}^{7}$AD1/16/19/
5+E1/19/5$c_{1}^{8}$AD1/16/19/5+E1/19/5$c_{1}^{35}$AD1/16/19/5+E1/19/5$c_{1}^{53}$AD1/16/19/5+E1/19/5$c_{1}^{8}$AD1/16/
19/5+E1/19/5$c_{1}^{13}$AD1/16/19/5+E1/19/5$c_{1}^{138}$AD1/16/19/5+E1/19/5$c_{1}^{61}$AD1/16/19/5+E1/19/5$c_{2}^{104}c$
$_{1}^{8}c_{2}^{35}c_{1}^{6}$AD1/16/19/5+E1/19/5$c_{1}^{17}$AD1/16/19/5+E1/19/5$c_{1}^{8}$AD1/16/19/5+E1/19/5$c_{1}^{10}$AD1/16/19/5
+E1/19/5$c_{1}^{3}$AD1/16/19/5+E1/19/5$c_{1}^{10}$AD1/16/19/5+E1/19/5$c_{1}^{4}$AD1/16/19/5+E1/19/5$c_{1}^{9}$AD1/16/1
9/5+E1/19/5$c_{1}^{9}$AD1/16/19/5+E1/19/5$c_{1}^{25}$AD1/16/19/5+E1/19/5$c_{1}^{15}$AD1/16/19/5+E1/19/5$c_{1}c_{2}^{5}c_{1}^{8}$A
D1/16/19/5+E1/19/5$c_{1}^{33}$AD1/16/19/5+E1/19/5$c_{1}^{2}$AD1/16/19/5+E1/19/5$c_{1}^{13}$AD1/16/19/5+E1/19/5$c$

$_1^{23}$AD1/16/19/5+E1/19/5$c_1^{6}$AD1/16/19/5+E1/19/5$c_1^{63}$AD1/16/19/5+E1/19/5$c_1^{10}$AD1/16/19/5+E1/1
9/5$c_1^{6}$AD1/16/19/5+E1/19/5$c_1^{6}$AD1/16/19/5+E1/19/5$c_1^{7}$AD1/16/19/5+E1/19/5$c_1^{4}$AD1/16/19/5+E1
/19/5$c_1^{55}$AD1/16/19/5+E1/19/5$c_1^{8}$AD1/16/19/5+E1/19/5$^{2}c_1^{18}$AD1/16/19/5+E1/19/5$c_1^{11}$AD1/16/19/
5+E1/19/5$c_1^{4}$AD1/16/19/5+E1/19/5$c_1^{8}$AD1/16/19/5+E1/19/5$c_1^{16}$AD1/16/19/5+E1/19/5$c_1^{8}$AD1/16/
19/5+E1/19/5$c_1^{10}$AD1/16/19/5+E1/19/5$c_1^{3}$AD1/16/19/5+E1/19/5$c_1^{8}$AD1/16/19/5+E1/19/5$c_1^{20}$AD1/
16/19/5+E1/19/5$c_1^{2}$AD1/16/19/5+E1/19/5$c_1$AD1/16/19/5+E1/19/5$c_1^{13}$AD1/16/19/5+E1/19/5$c_1^{8}$AD
1/16/19/5+E1/19/5$c_1^{15}$AD1/16/19/5+E1/19/5$c_1^{2}$AD1/16/19/5+E1/19/5$c_1^{2}$AD1/16/19/5+E1/19/5$c_1^{6}$
$c_2^{3}$G1/19/5+F19+R19$c_2^{12}$AD1/16/19/5+E1/19/5$c_1^{22}c_2^{9}$G1/19/5+F19+R19$c_2^{13}c_1^{6}c_2^{11}$G1/19/5+F19+
R19$c_2^{5}c_1^{7}$AD1/16/19/5+E1/19/5$^{2}c_1^{5}$AD1/16/19/5+E1/19/5$c_1$AD1/16/19/5+E1/19/5$^{2}c_1$AD1/16/19/5
+E1/19/5$c_1^{2}$AD1/16/19/5+E1/19/5$c_1c_2c_1^{13}$AD1/16/19/5+E1/19/5$c_1^{4}$AD1/16/19/5+E1/19/5$c_1^{7}$AD1/1
6/19/5+E1/19/5$c_2^{19}c_1^{5}$AD1/16/19/5+E1/19/5$c_1^{4}$AD1/16/19/5+E1/19/5$c_1^{3}$AD1/16/19/5+E1/19/5$c_1^{5}$
AD1/16/19/5+E1/19/5$c_1^{4}$AD1/16/19/5+E1/19/5$c_1^{5}$AD1/16/19/5+E1/19/5$c_1^{4}$AD1/16/19/5+E1/19/5c
$_1^{2}$AD1/16/19/5+E1/19/5$c_1^{5}$AD1/16/19/5+E1/19/5$c_2^{5}c_1^{4}$AD1/16/19/5+E1/19/5$c_2^{5}c_1^{4}$AD1/16/19/5+E
1/19/5$^{2}c_1^{6}$AD1/16/19/5+E1/19/5$c_1^{10}$AD1/16/19/5+E1/19/5$c_2^{8}$G1/19/5+F19+R19$c_2^{18}c_1^{6}$AD1/16/19/
5+E1/19/5$c_2^{12}c_1^{8}$AD1/16/19/5+E1/19/5$c_2^{5}$AD1/16/19/5+E1/19/5$c_1^{6}$AD1/16/19/5+E1/19/5$c_2^{5}c_1^{5}$AD
1/16/19/5+E1/19/5$c_1^{5}$AD1/16/19/5+E1/19/5$c_1$AD1/16/19/5+E1/19/5$^{2}c_1^{4}$AD1/16/19/5+E1/19/5$c_1^{7}$A
D1/16/19/5+E1/19/5$c_1^{4}$AD1/16/19/5+E1/19/5$c_1^{14}$AD1/16/19/5+E1/19/5G1/19/5+F19+R19$c_2^{3}$G1/1
9/5+F19+R19$c_2c_1^{13}$AD1/16/19/5+E1/19/5$c_2^{4}$G1/19/5+F19+R19$c_2^{12}$G1/19/5+F19+R19$c_2^{6}c_1^{3}$AD1/
16/19/5+E1/19/5$c_1c_2^{4}$G1/19/5+F19+R19$c_2^{17}$G1/19/5+F19+R19$c_2^{8}c_1c_2c_1^{2}$AD1/16/19/5+E1/19/5$c_1^{5}$
$c_2^{5}$AD1/16/19/5+E1/19/5$c_1c_2^{4}c_1^{12}$AD1/16/19/5+E1/19/5$c_1^{3}$AD1/16/19/5+E1/19/5$c_1c_2^{6}c_1^{7}$AD1/16/1
9/5+E1/19/5$c_1c_2^{7}c_1^{19}$AD1/16/19/5+E1/19/5$c_1^{13}$AD1/16/19/5+E1/19/5$^{2}c_1^{10}$AD1/16/19/5+E1/19/5$c_1^{1}$
$^{2}$AD1/16/19/5+E1/19/5$c_1^{3}$AD1/16/19/5+E1/19/5$c_2^{3}$AD1/16/19/5+E1/19/5$c_1^{3}$AD1/16/19/5+E1/19/5
$c_2^{3}c_1^{2}$AD1/16/19/5+E1/19/5$c_1^{9}$AD1/16/19/5+E1/19/5$c_2^{4}c_1^{2}$AD1/16/19/5+E1/19/5$c_1^{3}$AD1/16/19/5+
E1/19/5$c_1^{3}$AD1/16/19/5+E1/19/5$c_1^{2}$AD1/16/19/5+E1/19/5$^{2}c_1^{8}$AD1/16/19/5+E1/19/5$c_1^{2}$AD1/16/19/
5+E1/19/5$^{2}c_1^{8}$AD1/16/19/5+E1/19/5$c_1^{7}$AD1/16/19/5+E1/19/5$c_2^{2}c_1^{5}$AD1/16/19/5+E1/19/5$c_1^{2}$AD1/1
6/19/5+E1/19/5$c_2^{2}c_1^{2}$AD1/16/19/5+E1/19/5$c_1^{9}$AD1/16/19/5+E1/19/5$c_1^{2}$AD1/16/19/5+E1/19/5$c_1^{9}$A
D1/16/19/5+E1/19/5$c_1$AD1/16/19/5+E1/19/5$c_1^{2}$AD1/16/19/5+E1/19/5$c_2^{8}c_1^{8}$AD1/16/19/5+E1/19/5
$c_1^{2}$AD1/16/19/5+E1/19/5$c_1^{8}$AD1/16/19/5+E1/19/5$c_2^{9}c_1^{5}$AD1/16/19/5+E1/19/5$c_1^{8}$AD1/16/19/5+E1/
19/5$c_1^{5}c_2c_1^{9}$AD1/16/19/5+E1/19/5$c_1^{15}$AD1/16/19/5+E1/19/5$c_1^{14}c_2^{9}c_1^{24}$AD1/16/19/5+E1/19/5$c_2^{6}$G1
/19/5+F19+R19$c_2^{16}$G1/19/5+F19+R19$c_2^{13}$G1/19/5+F19+R19$c_2^{8}c_1^{2}c_2^{8}$AD1/16/19/5+E1/19/5$c_2^{6}c_1^{7}$
AD1/16/19/5+E1/19/5$c_2^{6}c_1^{6}c_2^{10}c_1^{10}$AD1/16/19/5+E1/19/5$c_1^{10}$AD1/16/19/5+E1/19/5$c_1^{2}$AD1/16/19/
5+E1/19/5$c_1^{13}$AD1/16/19/5+E1/19/5$c_1$AD1/16/19/5+E1/19/5$c_1^{3}$AD1/16/19/5+E1/19/5$c_1^{4}$AD1/16/1
9/5+E1/19/5$c_2c_1^{5}$AD1/16/19/5+E1/19/5$c_2^{5}c_1c_2^{6}$AD1/16/19/5+E1/19/5$c_1^{7}$AD1/16/19/5+E1/19/5$^{2}c_1^{4}$
AD1/16/19/5+E1/19/5$c_2^{26}c_1^{10}c_2^{8}$G1/19/5+F19+R19$c_2^{5}c_1$AD1/16/19/5+E1/19/5$c_1^{7}c_2^{8}c_1^{7}$AD1/16/19/
5+E1/19/5$c_1^{15}$AD1/16/19/5+E1/19/5$c_1^{7}$AD1/16/19/5+E1/19/5$c_1^{49}$AD1/16/19/5+E1/19/5$c_1$AD1/16/
19/5+E1/19/5$^{3}c_1$AD1/16/19/5+E1/19/5$c_1^{3}$AD1/16/19/5+E1/19/5$c_1^{2}$AD1/16/19/5+E1/19/5$c_1$AD1/1
6/19/5+E1/19/5$^{2}c_1^{8}$AD1/16/19/5+E1/19/5$c_2^{12}$G1/19/5+F19+R19$c_2^{7}$G1/19/5+F19+R19$c_2^{3}$AD1/16/
19/5+E1/19/5$^{3}c_1^{3}$AD1/16/19/5+E1/19/5$c_1$AD1/16/19/5+E1/19/5$c_1^{16}$AD1/16/19/5+E1/19/5$c_1$AD1/1
6/19/5+E1/19/5$^{2}c_1^{2}$AD1/16/19/5+E1/19/5$^{2}c_1^{3}$AD1/16/19/5+E1/19/5$c_1$AD1/16/19/5+E1/19/5$^{2}c_1$AD
1/16/19/5+E1/19/5$^{2}c_1^{9}c_2^{12}c_1$AD1/16/19/5+E1/19/5$c_2^{3}c_1^{2}$AD1/16/19/5+E1/19/5$c_1$AD1/16/19/5+E1/
19/5$c_2^{4}c_1$AD1/16/19/5+E1/19/5$^{2}c_1$AD1/16/19/5+E1/19/5$^{2}c_1^{5}$AD1/16/19/5+E1/19/5$c_2^{6}$G1/19/5+F1
9+R19$c_2^{3}$AD1/16/19/5+E1/19/5$c_1^{2}$AD1/16/19/5+E1/19/5$^{2}c_1^{7}$AD1/16/19/5+E1/19/5$c_1^{8}$AD1/16/19/5
+E1/19/5$c_1^{10}$AD1/16/19/5+E1/19/5$c_1^{10}$AD1/16/19/5+E1/19/5$c_1^{2}$AD1/16/19/5+E1/19/5$c_1^{16}$AD1/16/
19/5+E1/19/5$c_1^{3}$AD1/16/19/5+E1/19/5$c_1$AD1/16/19/5+E1/19/5$^{4}c_1$AD1/16/19/5+E1/19/5$^{3}c_1^{4}$AD1/1
6/19/5+E1/19/5$c_1$AD1/16/19/5+E1/19/5$^{2}c_1$AD1/16/19/5+E1/19/5$^{2}c_1^{5}$AD1/16/19/5+E1/19/5$c_1c_2^{5}c_1^{1}$
$^{1}$AD1/16/19/5+E1/19/5$^{2}c_1$AD1/16/19/5+E1/19/5$^{2}c_1^{9}c_2^{12}c_1^{10}$AD1/16/19/5+E1/19/5$c_2^{9}$G1/19/5+F19
+R19$c_2^{9}$G1/19/5+F19+R19$c_2^{15}$G1/19/5+F19+R19$c_2^{10}c_1^{11}$AD1/16/19/5+E1/19/5$c_2^{5}c_1^{3}$AD1/16/19/5
+E1/19/5$c_2^{38}c_1^{17}$AD1/16/19/5+E1/19/5$c_2^{5}c_1^{5}$AD1/16/19/5+E1/19/5$c_1^{7}$AD1/16/19/5+E1/19/5$c_2^{17}c_1^{8}$
AD1/16/19/5+E1/19/5$c_1^{6}$AD1/16/19/5+E1/19/5$c_1^{6}$AD1/16/19/5+E1/19/5$c_1^{19}c_2^{52}c_1^{2}$AD1/16/19/5+E
1/19/5$^{2}c_1^{3}$AD1/16/19/5+E1/19/5$c_1$AD1/16/19/5+E1/19/5$c_2c_1^{6}$AD1/16/19/5+E1/19/5$^{2}c_1^{6}$AD1/16/19
/5+E1/19/5$c_1^{3}$AD1/16/19/5+E1/19/5$c_2c_1$AD1/16/19/5+E1/19/5$c_2^{5}c_1c_2^{3}c_1$AD1/16/19/5+E1/19/5$c_2^{5}$c

$_1$AD1/16/19/5+E1/19/5$c_2^{23}$G1/19/5+F19+R19$c_2^{2}$G1/19/5+F19+R19$c_2^{2}$AD1/16/19/5+E1/19/5$^{2}c_1^{6}$A
D1/16/19/5+E1/19/5$c_1^{7}$AD1/16/19/5+E1/19/5$c_2^{4}$G1/19/5+F19+R19AD1/16/19/5+E1/19/5$c_1^{2}$AD1/
16/19/5+E1/19/5$c_2^{4}$G1/19/5+F19+R19AD1/16/19/5+E1/19/5$c_1^{3}$AD1/16/19/5+E1/19/5$c_1$AD1/16/1
9/5+E1/19/5$c_1$AD1/16/19/5+E1/19/5$c_1$AD1/16/19/5+E1/19/5$c_1$AD1/16/19/5+E1/19/5$c_1$AD1/16/19
/5+E1/19/5$c_1$AD1/16/19/5+E1/19/5$c_1$AD1/16/19/5+E1/19/5$c_1^{2}$AD1/16/19/5+E1/19/5$c_1$AD1/16/19/
5+E1/19/5$^{2}c_1$AD1/16/19/5+E1/19/5$c_1$AD1/16/19/5+E1/19/5$c_2c_1$AD1/16/19/5+E1/19/5$^{2}c_1$AD1/16/1
9/5+E1/19/5$c_1$AD1/16/19/5+E1/19/5$c_1^{2}$AD1/16/19/5+E1/19/5$c_2c_1^{2}$AD1/16/19/5+E1/19/5$c_1$AD1/16
/19/5+E1/19/5$c_1^{2}$AD1/16/19/5+E1/19/5$c_1$AD1/16/19/5+E1/19/5$c_1^{2}$AD1/16/19/5+E1/19/5$c_1$AD1/16
/19/5+E1/19/5$^{2}c_1$AD1/16/19/5+E1/19/5$c_1^{2}$AD1/16/19/5+E1/19/5$^{2}c_1$AD1/16/19/5+E1/19/5$c_1$AD1/1
6/19/5+E1/19/5$^{2}c_1$AD1/16/19/5+E1/19/5$c_1$AD1/16/19/5+E1/19/5$c_1^{2}$AD1/16/19/5+E1/19/5$c_1$AD1/1
6/19/5+E1/19/5$^{2}c_1$AD1/16/19/5+E1/19/5$c_1$AD1/16/19/5+E1/19/5$c_1^{2}$AD1/16/19/5+E1/19/5$c_1$AD1/1
6/19/5+E1/19/5$c_1^{2}$AD1/16/19/5+E1/19/5$c_1$AD1/16/19/5+E1/19/5$^{2}c_1$AD1/16/19/5+E1/19/5$c_1$AD1/1
6/19/5+E1/19/5$c_1^{2}$AD1/16/19/5+E1/19/5$c_1$AD1/16/19/5+E1/19/5$c_1^{2}$AD1/16/19/5+E1/19/5$c_1$AD1/1
6/19/5+E1/19/5$c_1^{2}$AD1/16/19/5+E1/19/5$c_1$AD1/16/19/5+E1/19/5$c_1^{2}$AD1/16/19/5+E1/19/5$c_1$AD1/1
6/19/5+E1/19/5$c_1^{2}$AD1/16/19/5+E1/19/5$c_1$AD1/16/19/5+E1/19/5$c_1^{2}$AD1/16/19/5+E1/19/5$c_1$AD1/1
6/19/5+E1/19/5$c_1^{2}$AD1/16/19/5+E1/19/5$c_1$AD1/16/19/5+E1/19/5$c_1^{2}$AD1/16/19/5+E1/19/5$c_1$AD1/1
6/19/5+E1/19/5$^{2}c_1^{2}$AD1/16/19/5+E1/19/5$c_1$AD1/16/19/5+E1/19/5$c_1^{2}$AD1/16/19/5+E1/19/5$c_1$AD1/
16/19/5+E1/19/5$c_1^{2}$AD1/16/19/5+E1/19/5$c_1$AD1/16/19/5+E1/19/5$c_1^{2}$AD1/16/19/5+E1/19/5$c_1$AD1/
16/19/5+E1/19/5$c_1^{2}$AD1/16/19/5+E1/19/5$c_1$AD1/16/19/5+E1/19/5$^{2}c_1$AD1/16/19/5+E1/19/5$c_1$AD1/
16/19/5+E1/19/5$^{2}c_1$AD1/16/19/5+E1/19/5$c_1$AD1/16/19/5+E1/19/5$^{2}c_1$AD1/16/19/5+E1/19/5$^{2}c_1$AD
1/16/19/5+E1/19/5$c_1$AD1/16/19/5+E1/19/5$^{2}c_1$AD1/16/19/5+E1/19/5$^{2}c_1$AD1/16/19/5+E1/19/5$c_2c_1^{2}$
AD1/16/19/5+E1/19/5$c_1$AD1/16/19/5+E1/19/5$c_1^{2}$AD1/16/19/5+E1/19/5$c_1$AD1/16/19/5+E1/19/5$c_1$
AD1/16/19/5+E1/19/5$c_2^{3}c_1$AD1/16/19/5+E1/19/5$c_1$AD1/16/19/5+E1/19/5$c_1^{3}$AD1/16/19/5+E1/19/5
$c_1$AD1/16/19/5+E1/19/5$^{2}c_1$AD1/16/19/5+E1/19/5$c_1$AD1/16/19/5+E1/19/5$c_1^{10}$AD1/16/19/5+E1/19/
5$c_1$AD1/16/19/5+E1/19/5$c_1^{6}$AD1/16/19/5+E1/19/5$c_1^{2}$AD1/16/19/5+E1/19/5$c_1^{3}$AD1/16/19/5+E1/1
9/5$c_1^{6}$AD1/16/19/5+E1/19/5$c_2c_1^{3}$AD1/16/19/5+E1/19/5$c_1^{5}$AD1/16/19/5+E1/19/5$c_1$AD1/16/19/5+E
1/19/5$^{2}c_1^{3}$AD1/16/19/5+E1/19/5$c_1^{3}$AD1/16/19/5+E1/19/5$c_1^{2}$AD1/16/19/5+E1/19/5$c_1^{2}$AD1/16/19/5
+E1/19/5$c_1^{3}$AD1/16/19/5+E1/19/5$c_1^{28}$AD1/16/19/5+E1/19/5$c_1^{14}$AD1/16/19/5+E1/19/5$c_1^{9}$AD1/16/1
9/5+E1/19/5$c_1^{9}$AD1/16/19/5+E1/19/5$c_1^{7}$AD1/16/19/5+E1/19/5$c_1^{15}$AD1/16/19/5+E1/19/5$c_1^{16}$AD1/1
6/19/5+E1/19/5$c_1^{11}$AD1/16/19/5+E1/19/5$c_1^{12}$AD1/16/19/5+E1/19/5$c_1^{6}$AD1/16/19/5+E1/19/5$c_1^{7}$AD
1/16/19/5+E1/19/5$c_1^{12}$AD1/16/19/5+E1/19/5$c_1^{2}$AD1/16/19/5+E1/19/5$c_1^{7}$AD1/16/19/5+E1/19/5$c_1^{11}$
AD1/16/19/5+E1/19/5$c_1^{6}$AD1/16/19/5+E1/19/5$c_1^{13}c_2^{4}c_1^{6}$AD1/16/19/5+E1/19/5$c_1^{5}$AD1/16/19/5+E1
/19/5$^{2}c_1^{8}$AD1/16/19/5+E1/19/5$c_1^{4}$AD1/16/19/5+E1/19/5$c_1^{3}$AD1/16/19/5+E1/19/5$c_1^{4}$AD1/16/19/5+
E1/19/5$c_2^{3}c_1$AD1/16/19/5+E1/19/5$c_1^{2}$AD1/16/19/5+E1/19/5$^{2}c_1^{4}$AD1/16/19/5+E1/19/5$c_2^{11}c_1$AD1/1
6/19/5+E1/19/5$c_1^{5}$AD1/16/19/5+E1/19/5$c_2^{6}c_1$AD1/16/19/5+E1/19/5$c_1^{9}$AD1/16/19/5+E1/19/5$c_2^{4}c_1^{2}$
AD1/16/19/5+E1/19/5$c_1^{9}$AD1/16/19/5+E1/19/5$c_1^{2}$AD1/16/19/5+E1/19/5$c_1^{4}$AD1/16/19/5+E1/19/5$c$
$_1^{2}c_2^{3}c_1^{7}$AD1/16/19/5+E1/19/5$c_1^{2}$AD1/16/19/5+E1/19/5$c_1^{3}$AD1/16/19/5+E1/19/5$c_1^{2}c_2^{5}c_1^{3}$AD1/16/1
9/5+E1/19/5$c_1^{2}c_2^{2}c_1^{2}$AD1/16/19/5+E1/19/5$c_1^{7}$AD1/16/19/5+E1/19/5$c_1^{14}$AD1/16/19/5+E1/19/5$c_1^{5}$A
D1/16/19/5+E1/19/5$c_1^{11}$AD1/16/19/5+E1/19/5$c_1^{5}$AD1/16/19/5+E1/19/5$c_1^{12}$AD1/16/19/5+E1/19/5$c$
$_1^{3}$AD1/16/19/5+E1/19/5$c_1^{4}$AD1/16/19/5+E1/19/5$c_1^{2}$AD1/16/19/5+E1/19/5$c_1^{3}$AD1/16/19/5+E1/19/
5$c_1^{11}$AD1/16/19/5+E1/19/5$c_1^{11}$AD1/16/19/5+E1/19/5$c_1^{3}$AD1/16/19/5+E1/19/5$c_1^{8}$AD1/16/19/5+E1/
19/5$c_1^{2}$AD1/16/19/5+E1/19/5$c_1^{4}$AD1/16/19/5+E1/19/5$c_1^{2}$AD1/16/19/5+E1/19/5$c_1$AD1/16/19/5+E
1/19/5$c_1^{4}$AD1/16/19/5+E1/19/5$c_1^{4}$AD1/16/19/5+E1/19/5$c_1$AD1/16/19/5+E1/19/5$c_1$AD1/16/19/5+E
1/19/5$c_1^{4}$AD1/16/19/5+E1/19/5$c_1^{4}$AD1/16/19/5+E1/19/5$c_1$AD1/16/19/5+E1/19/5$c_1^{2}$AD1/16/19/5+
E1/19/5$c_1^{2}c_2^{9}c_1^{2}$AD1/16/19/5+E1/19/5$c_1^{3}$AD1/16/19/5+E1/19/5$c_1^{4}$AD1/16/19/5+E1/19/5$c_1$AD1/16
/19/5+E1/19/5$c_1^{8}$AD1/16/19/5+E1/19/5$c_1^{34}$AD1/16/19/5+E1/19/5$c_1^{6}$AD1/16/19/5+E1/19/5$c_1^{9}$AD1/
16/19/5+E1/19/5$c_1^{6}$AD1/16/19/5+E1/19/5$c_1^{9}$AD1/16/19/5+E1/19/5$c_1^{37}c_2$G1/19/5+F19+R19$c_2^{10}c_1^{9}$
AD1/16/19/5+E1/19/5$c_1c_2^{2}c_1^{7}$AD1/16/19/5+E1/19/5$c_1^{5}$AD1/16/19/5+E1/19/5$c_1^{10}$AD1/16/19/5+E1/
19/5$c_1^{3}$AD1/16/19/5+E1/19/5$c_1^{6}$AD1/16/19/5+E1/19/5$^{2}c_1^{5}c_2^{2}c_1^{2}$AD1/16/19/5+E1/19/5$^{3}c_1^{5}$AD1/16/
19/5+E1/19/5$c_2^{2}c_1$AD1/16/19/5+E1/19/5$c_1$AD1/16/19/5+E1/19/5$c_2^{2}c_1$AD1/16/19/5+E1/19/5$c_2^{3}c_1^{3}$
AD1/16/19/5+E1/19/5$c_2^{6}$AD1/16/19/5+E1/19/5$c_2^{18}c_1$AD1/16/19/5+E1/19/5$c_2^{9}c_1$AD1/16/19/5+E1/

19/5$c_2^{6}c_1^{2}$AD1/16/19/5+E1/19/5$c_2^{4}$AD1/16/19/5+E1/19/5$c_2^{4}$G1/19/5+F19+R19$c_2^{8}c_1$AD1/16/19/5+E1/19/5$c_2^{8}$G1/19/5+F19+R19$c_1^{6}$AD1/16/19/5+E1/19/5$c_2^{23}c_1$AD1/16/19/5+E1/19/5$c_2^{49}c_1^{7}c_2^{9}c_1^{9}$AD1/16/19/5+E1/19/5$^{4}c_1^{4}$AD1/16/19/5+E1/19/5

### cen20

$p_{12-13}p_{12-14}p_{1-2}$D20$p_{6-10}$L20A20$p_{7-13}p_{15-7}c_{14}^{4}p_{14-3}p_{16-7}c_{14}^{2}p_{14-3}p_{16-7}c_{14}^{2}c_{12}^{12}p_{14-11}^{2}p_{14-5}p_{9-12}$K20.1$p_{14-16}p_{7-15}p_{14-15}$F2/20$c_{15}^{2}$O20$c_{12}c_{14}^{19}p_{14-9}c_2^{3}p_{2-9}c_2^{7}p_{2-14}p_{7-10}c_{16}p_{16-12}$N20$c_7^{15}p_{7-5}p_{6-16}p_{7-11}p_{14-5}p_{7-11}$N20$c_7^{6}c_9^{12}p_{9-6}p_{9-10}c_7^{8}p_{7-10}c_{13}p_{13-11}c_{14}^{2}c_{12}^{4}p_{14-11}^{3}p_{14-12}$K20.1$c_{14}p_{14-11}p_{14-5}c_5^{9}p_{5-10}$K20$c_{12}^{36}p_{14-10}c_{12}^{9}p_{12-3}c_{16}^{10}p_{16-13}p_{12-15}c_{11}p_{11-13}c_{12}^{2}p_{12-13}c_{12}p_{12-13}c_{12}p_{12-13}c_{12}p_{12-13}c_{12}^{5}p_{12-13}c_{12}p_{12-13}c_{12}p_{12-13}c_{12}^{7}p_{12-13}c_{12}p_{12-13}c_{12}p_{12-13}c_{12}^{4}p_{12-13}c_{12}p_{12-13}c_{12}^{2}p_{12-13}c_{12}^{8}p_{12-13}c_{12}^{10}p_{12-5}c_{16}$P20$c_3^{3}p_{3-16}c_3^{2}p_{3-14}c_{14}$N20$c_{14}$N20$c_{12}^{6}p_{12-3}c_{10}^{4}p_{10-6}c_{15}^{5}p_{15-6}c_{15}^{5}p_{15-6}c_{15}^{7}p_{15-3}p_{10-14}c_{14}^{14}p_{14-16}c_3^{9}p_{3-6}c_{15}^{6}p_{15-6}c_{15}p_{15-6}c_{15}^{2}p_{1-6}c_{15}^{7}p_{15-16}c_3^{8}p_{3-6}c_{15}^{6}p_{15-6}c_{15}^{2}p_{1-6}c_{15}^{12}p_{15-16}c_3p_{3-16}^{2}c_3p_{3-16}c_2^{44}p_{2-5}c_{16}^{23}p_{16-10}c_{13}p_{13-10}c_{13}^{3}$K20$p_{12-1}c_3^{8}p_{3-13}c_{12}^{2}p_{12-13}c_{12}^{4}p_{12-6}c_9^{5}p_{9-13}p_{12-1}p_{3-13}p_{12-1}p_{3-1}c_3^{8}p_{3-13}c_{12}^{6}p_{12-13}c_{12}^{34}$L20$c_{15}^{16}p_{15-12}c_{15}^{12}p_{15-11}c_{14}^{18}p_{14-16}c_3p_{3-16}c_3^{7}p_{3-16}c_3p_{3-14}c_{11}^{6}p_{11-13}c_{12}^{2}p_{12-5}p_{7-13}c_{12}^{4}p_{12-13}c_{12}^{2}p_{12-13}c_{12}^{2}p_{12-5}c_7^{2}p_{7-11}c_{14}^{7}c_{12}^{3}p_{12-13}c_{12}p_{12-13}c_{12}p_{12-13}c_{12}p_{12-5}c_7^{8}p_{7-16}c_3p_{3-13}c_{12}^{2}p_{12-13}c_{12}^{2}p_{12-13}c_{12}^{3}p_{12-13}c_{12}^{7}p_{12-7}c_2^{13}p_{2-13}c_{12}^{3}p_{12-13}c_{12}p_{12-13}c_{12}^{14}c_{14}p_{14-11}c_{14}p_{14-11}c_{14}^{29}c_{12}^{4}p_{12-13}c_{12}^{7}p_{12-13}c_{12}^{5}p_{12-13}c_{12}^{5}p_{12-13}c_{12}^{3}p_{12-13}c_{12}^{22}p_{12-5}p_{7-11}c_{14}p_{14-5}p_{7-11}c_{14}^{20}p_{14-7}c_{14}p_{14-6}p_{8-11}p_{14-6}p_{8-11}c_{14}^{3}p_{14-15}$F2/20$p_{8-11}c_{14}^{10}p_{14-7}c_{14}p_{14-6}p_{8-11}p_{14-6}p_{8-11}c_{14}^{3}p_{14-15}$F2/20$p_{8-11}c_{14}^{5}p_{14-6}p_{11-6}p_{11-15}$F2/20$c_{13}p_{13-5}p_{7-11}p_{14-6}c_8^{4}p_{8-14}c_{12}^{2}p_{12-14}c_{12}^{2}p_{12-14}c_{12}^{3}$L20

### cen21

G13/21.1$p_{1-3}p_{1-2}p_{9-7}p_{1-2}p_{9-3}p_{8-2}c_3^{13}p_{4-2}^{2}p_{4-6}c_7^{23}p_{11-6}c_7^{5}$G13/21$c_1^{5}$I13/21$p_{4-6}c_7^{2}p_{5-7}c_8p_{10-7}c_8p_{10-7}c_8p_{10-7}c_8^{2}p_{10-7}c_8p_{10-7}c_8^{3}p_{10-7}c_8^{3}p_{10-7}c_8^{3}p_{10-7}c_8^{2}p_{10-7}c_8^{3}p_{10-7}c_8^{3}p_{10-7}c_8^{7}p_{1-3}c_4^{33}p_{1-5}c_6p_{7-11}c_1^{6}p_{8-11}c_1^{2}p_{8-4}c_5^{39}$

### cen22

$p_{5-7}p_{1-2}$D14/22$c_1^{64}p_{3-8}c_1p_{3-1}c_2^{41}$D14/22$c_5^{13}p_{1-4}c_5^{4}p_{1-4}c_5^{5}p_{1-5}c_6^{29}p_{7-5}c_6^{5}p_{7-5}c_7^{37}p_{3-4}c_5^{42}$A14/22$c_2^{2}p_{4-7}p_{1-7}c_8^{23}p_{1-7}c_8^{3}p_{1-7}c_8p_{1-7}p_{1-3}c_4^{21}p_{3-1}c_2^{15}p_{4-1}p_{4-8}c_1p_{4-1}p_{4-8}^{2}c_1^{5}p_{4-8}c_1^{4}p_{4-1}c_2^{3}p_{4-1}p_{4-8}c_1p_{4-1}^{2}p_{4-8}c_1^{5}p_{4-1}c_2^{7}p_{4-8}p_{4-1}c_2^{2}p_{4-8}p_{4-7}c_8^{6}c_1^{20}p_{3-8}c_1^{24}p_{3-8}c_1p_{3-8}c_1^{2}p_{3-8}c_1^{8}p_{3-8}c_1p_{3-8}^{2}c_1p_{3-7}c_8^{27}p_{1-5}^{2}c_6^{3}$G14/22B14/22$c_3^{12}p_{6-7}p_{1-7}c_8^{30}p_{1-7}c_8^{4}p_{1-7}c_8^{3}p_{1-5}c_6^{2}p_{7-5}$G14/22$p_{2-5}p_{7-5}$G14/22$p_{2-5}c_6^{2}$G14/22$p_{1-5}c_6$G14/22$p_{2-5}c_6^{3}$G14/22$p_{2-5}p_{7-5}$G14/22$p_{2-5}$G14/22$p_{1-5}c_6$G14/22$p_{2-5}c_6^{2}$G14/22$p_{2-5}c_6^{3}$G14/22$p_{1-5}c_6$G14/22$p_{2-5}c_6^{3}p_{7-8}p_{2-5}c_6^{3}$G14/22$p_{2-5}c_6p_{7-5}^{2}p_{7-8}p_{2-5}c_6^{3}$G14/22$p_{2-5}c_6^{4}p_{7-5}c_6^{2}$G14/22$p_{2-5}c_6^{5}p_{7-5}c_6^{2}p_{7-5}c_6^{2}p_{7-5}c_6^{2}p_{7-5}c_6^{6}p_{7-5}c_6p_{7-5}c_6^{6}p_{7-8}c_1^{147}p_{2-7}c_8^{93}p_{1-7}c_8^{6}p_{1-7}c_8^{211}p_{1-7}^{2}c_8p_{1-7}^{2}c_8p_{1-7}c_8^{20}p_{1-7}c_8^{40}p_{1-7}c_8^{18}p_{1-7}c_8^{54}p_{1-7}^{2}c_8^{10}p_{1-7}^{2}c_8^{2}p_{1-7}c_8^{2}p_{1-7}^{2}c_8^{2}p_{2-7}c_8^{39}p_{1-7}^{2}c_8^{2}p_{1-7}c_8^{2}p_{1-7}^{2}c_8^{2}p_{2-7}c_8^{23}p_{1-6}c_7^{38}p_{1-6}c_7^{4}p_{1-6}c_7^{151}p_{1-6}c_7^{14}p_{1-6}c_7^{11}p_{1-6}c_7^{6}p_{1-6}c_7^{12}p_{1-6}c_7p_{1-6}c_7^{18}p_{1-6}c_7^{123}p_{1-6}c_7^{16}p_{1-6}c_7^{14}p_{1-6}c_7^{7}p_{1-6}c_7^{12}p_{1-6}c_7p_{1-6}c_7^{9}p_{1-6}c_7p_{1-6}c_7^{20}p_{1-7}c_8^{129}p_{1-7}p_{1-6}c_7^{3}p_{1-7}^{3}c_8^{2}p_{1-7}^{2}c_8p_{1-5}$G14/22$p_{1-7}c_8p_{1-7}c_8^{3}p_{1-7}c_8^{2}p_{1-5}$G14/22$p_{1-7}c_8p_{1-7}c_8^{3}p_{1-7}c_8^{3}p_{1-7}c_8^{6}p_{1-7}^{2}c_8^{7}p_{1-7}c_8^{3}p_{1-7}c_8^{3}p_{1-7}c_8p_{1-5}c_6p_{7-5}$G14/22$p_{1-5}c_6^{4}p_{7-5}$G14/22$p_{1-5}c_6p_{7-5}$G14/22$p_{1-5}c_6^{2}p_{7-5}c_6p_{7-5}$G14/22$p_{2-5}c_6^{2}p_{7-5}$G14/22$c_8p_{1-5}p_{7-5}^{2}$G14/22$p_{1-5}c_6p_{7-5}c_6$G14/22$p_{1-5}c_6^{5}p_{7-5}^{2}$G14/22$c_8^{2}p_{1-7}c_8^{2}p_{1-5}c_6^{5}p_{7-5}p_{7-8}p_{8-5}p_{7-5}^{2}c_6p_{7-5}c_6^{2}$G14/22$p_{1-5}c_6$G14/22$p_{1-5}c_6$G14/22$p_{1-5}c_6^{2}$G14/22$p_{1-5}c_6$G14/22$p_{1-5}c_6$G14/22$p_{1-5}c_6$G14/22$p_{1-5}c_6^{3}p_{7-5}c_6^{4}p_{7-5}c_6p_{7-5}c_6^{8}p_{7-5}c_6^{2}$G14/22$p_{1-5}c_6$G14/22$p_{1-5}c_6^{3}p_{7-5}c_6^{2}p_{7-5}c_6p_{7-5}c_6^{15}p_{7-5}c_6^{4}p_{7-5}c_6^{6}p_{7-5}c_6c_7^{5}p_{7-2}c_3c_6^{2}p_{7-5}c_6p_{7-5}c_6p_{7-5}c_6^{2}p_{7-5}c_6c_7^{2}p_{7-2}c_3c_6^{2}p_{7-5}c_6p_{7-5}c_6p_{7-5}c_6^{2}p_{7-5}c_6c_7^{5}$G14/22$c_8p_{2-8}^{2}c_1^{2}p_{2-8}c_1^{2}$B14/22$c_3$E14/22

### cenX

$p_{10-12}p_{5-6}$IXLX$p_{2-3}$EX$p_{9-12}p_{2-5}c_6^{22}p_{7-12}c_1^{3}p_{2-6}c_7^{3}p_{8-6}c_7p_{8-5}c_6^{95}p_{6-9}c_{10}^{26}$KX$c_{12}^{3}p_{1-7}c_8^{128}p_{11-7}c_8^{2}p_{11-7}c_8^{11}p_{11-7}c_8p_{11-7}c_8p_{11-7}c_8p_{11-7}c_8^{11}$KX$c_{12}^{8}p_{7-5}c_6^{2}p_{6-11}c_{12}p_{7-10}c_{11}^{50}p_{12-6}c_7^{174}p_{1-9}c_{10}^{240}p_{1-9}c_{10}^{8}p_{12-9}c_{10}^{18}p_{12-9}c_{10}^{8}p_{12-9}c_{10}^{19}p_{12-9}c_{10}^{8}p_{12-9}c_{10}^{13}p_{1-9}c_{10}p_{1-11}c_{12}^{41}p_{7-11}c_{12}^{6}p_{7-11}c_{12}^{5}p_{7-11}c_{12}^{7}p_{7-11}c_{12}^{16}p_{7-11}c_{12}^{4}p_{7-3}c_4^{13}p_{5-3}c_4^{21}$EX$c_6^{14}p_{7-3}c_4^{52}p_{5-2}c_3^{42}p_{11-5}c_6^{57}p_{6-12}c_1^{4}p_{8-11}c_{12}^{3}p_{8-11}c_{12}p_{8-12}c_1p_{8-4}c_5^{32}p_{6-2}c_3p_{5-1}c_2^{24}p_{3-4}c_5^{2}p_{6-4}^{4}p_{6-12}c_1^{94}p_{2-12}c_1^{11}p_{2-7}c_8^{14}p_{9-7}c_8^{5}p_{9-7}c_8^{21}p_{11-4}c_5^{28}p_{6-2}c_3^{87}$DX$p_{9-10}$
